# MCSS-based Predictions of Binding Mode and Selectivity of Nucleotide Ligands

**DOI:** 10.1101/622068

**Authors:** Roy González-Alemán, Nicolas Chevrollier, Manuel Simoes, Luis Montero-Cabrera, Fabrice Leclerc

## Abstract

Computational fragment-based approaches are widely used in drug design and drug discovery. One of the limitations of their application is the lack of performance of docking methods, mainly the scoring functions. With the emergence of new fragment-based approaches for single-stranded RNA ligands, we propose an analysis of an MCSS-based approach evaluated for its docking power on nucleotide-binding sites. Hybrid solvent models based on some partial explicit representation are shown to improve docking and screening powers. Clustering of the *n* best-ranked poses can also contribute to a lesser extent to better performance. The results suggest that we can apply the approach to the fragment-based design of sequence-selective oligonucleotides.

## Introduction

Fragment-based approaches are widely used in ligand design with several examples of “success stories” when applied to drug design and drug discovery (1–4) since the middle of the ‘90s (5). More than 30 fragment-based drug candidates have entered the clinic (6). Despite some hindrance related to synthetic accessibility and/or ligand-design strategies, fragment-based approaches remain very attractive while dealing more efficiently with chemical space, molecular complexity, probability of binding, and ligand efficiency (6). After high throughput screening, fragment-based approaches represent one of the three major lead generation strategies for clinical candidates (7).

Both experimental and computational approaches have been developed based on the same principles that weak-binding fragments can be converted into highly efficient ligands by covalent linking. One of the contributions to the gain of binding affinity with respect to that of the individual fragments comes from the rigid body entropic barrier, which is supposed to be independent of the molecular size. This gain is optimal when there is no energy penalty associated with the linker’s conformation and when the binding mode of each fragment is preserved in the ligand. Weak-binding fragments should still have enough favorable contacts to counterbalance the loss of rigid body entropy upon binding. In practice, the first step is to design and build a fragment library, the second is to screen the fragments, and the third to assemble them into ligands as lead compounds (8, 9).

In the experimental approaches, the fragments are validated by some screening methods, some of which are high throughput, e.g., by surface plasmon resonance (10). The screening is a critical step in the process of fragment-based design (FBD). In the computational approaches, the FBD is by default a structure-based approach like in the X-ray crystallography-based screening of fragments or in any other structural biology assisted FBD (2). However, the hits obtained *in silico* are not generally validated until the end of the process leading to the assembled ligands.

Very few published studies compare *in silico* to experimental approaches to validate virtual hits like in the screening of fragment-like inhibitors against the *N* ^5^-CAIR mutase (11). A computational screen of fragment libraries is faster and more cost-effective than in experimental approaches. However, the performance of such approaches may vary, although the case of the *N* ^5^-CAIR mutase shows a good overlap between the computational and experimental approaches. The lack of accuracy of the scoring functions is often invoked for the poor performance, i.e., the difficulty to discriminate native-like poses from false binding poses (12, 13). In the absence of validation after the screening step, sub-optimal fragments may be selected that are low-affinity binders. Thus, there is no guarantee we can identify the optimal fragments or those with a higher binding specificity by virtual screening.

Traditionally, the FBD approaches have been applied to the design of ligands assembled using small chemical groups selected from the fragment library, which is often built based on drug-like criteria. Since the fragments library should also cover some chemical space with the diversity of chemical groups and molecular properties, a good strategy is needed to assemble the fragments. The fragment merging or linking strategies consist of connecting covalently two non-competitive fragments by fusing some chemical bonds or creating some additional chemical bond(s) using a spacer to link both fragments. The alternative strategy, fragment growing (or fragment evolution), is less challenging; it can be viewed as an optimization process where one fragment is modified by adding some functional group that can make favorable contacts around the primary binding site of the fragment.

In the case of biopolymers, the chemical connectivity is well-defined, and thus the assembling strategy involves solving a distance-constraint problem to join the connecting atoms of successive residues. In the early days of computational FBD approaches, different flavors were implemented to design peptide ligands where the fragments are amino acid residues or moieties (14–18). Creating biopolymers is an easier task since the synthetic accessibility of designed ligands by *in silico* FBD is still very challenging. More recently, an FBD approach was applied using multi-residues fragments to avoid the high degrees of freedom to manage for the conformational sampling of long peptides (19).

A similar approach was applied to RNA ligands using trinucleotides to predict the binding mode of single-stranded RNAs to proteins (20, 21). In both cases, peptide or RNA ligands, the oligomer sequence is used as input for modeling the ligand-protein complex to reproduce known proteinligand interactions. Thus, some improvements are still required to progress towards the *de novo* fragment-based design of bound oligomers to protein targets.

In the case of RNA binding proteins (RBPs) that recognize single-stranded RNAs (ssRNAs), the primary binding site of the RNA binding domains from RBPs generally corresponds to an interface that can accommodate k-mers with k between 4 and 10 (22–24). However, the RNA motifs making contacts with the protein span a much shorter stretch of contiguous residues corresponding to 3-mers up to 5-mers in many structural families of RBPs (25). Such short RNA motifs, revealed by structural biology approaches, can be successive separated by short spacers and form longer bi-partite or tri-partite motifs that can easily extend to 10-mers (26). Very recent data obtained by high-throughput binding assays and sequencing on 78 human RBPs confirm this observation where the RNA motifs are composed of conserved 3-mers separated by spacers of 0 to 10 residues (27).

A preliminary study was done on the characterization of nucleotide-binding sites from ssRNAs involving contacts with the nucleic acid base moiety showing the existence of binding patterns (RsiteDB (28)). Using a knowledge-based approach, the study was extended to predict nucleotide and dinucleotide binding sites but in the perspective of screening small molecules mimicking the nucleotide-binding modes (29). Two other studies have been carried out using FBD approaches to model ssRNA-protein interactions. A method based on a coarse-grained model (RNA-LIM) was developed to model the structure of an ssRNA at the protein surface (30). However, its application is restricted to the RNA binding region surface, and the simplified representation of the nucleotides makes it impossible to distinguish between different nucleotide orientations and conformations. The more recent and advanced method was tested on a set of RBPs with RRM or Pumilio domains and could generate near-native models of RNA-protein complexes with good precision (RMSD ≤ 2Å) in most cases for chains up to 12-mers (20, 21). However, the scoring function still lacks the necessary accuracy to discriminate near-native poses robustly.

MCSS is a computational method that maps chemical functional groups at the surface of a protein target, making possible to perform virtual screening using pre-defined (15, 31, 32) or customized fragment libraries (33). Although MCSS does not include any fragment-assembly strategy, it has been widely used in FBD approaches in conjunction with fragment-linking/merging methods such as: HOOK (34), DLD (35), or CAVEAT (36) for chemical groups, and OLIGO (18) for oligopeptides or SiteMap for peptidomimetics (37). Specific applications of MCSS-based virtual screening were also reported for the identification of epitopes (38), the prediction of the displacement of water molecules upon binding (39). As with all computational FBD approaches, MCSS is generally applied using implicit solvent models, although the method also allows the docking of solvated fragments. The role of solvent is critical in the sampling and scoring of chemical fragments, but its implementation and evaluation in docking approaches remain challenging (40). The MCSS scoring function is based on the CHARMM energy function; different strategies have been applied to improve its performance using more accurate methods or solvent models. The first strategy includes post-processing of the MCSS generated fragment poses recalculating the score function by adding solvation terms (41), or by rescoring (single-point energy) using a new scheme (39, 42). The second strategy is a modification of the energy function during the MCSS calculations using, for example, a distance-dependent dielectric model (41), or an alternative charge model (43). As mentioned above, MCSS-based FBD approaches were applied repetitively to the design of peptides or peptidomimetics (15, 16, 18, 37) or to other biomolecules such as aminoglycosides (43). The CHARMM program’s development with its general force field (CGenFF (44, 45)) and solvent models makes it possible to extend the scope of applications of MCSS to any kind of chemical groups using the more appropriate solvent models.

In this study, we use an updated version of MCSS for the screening of nucleotides. We examine the ability to identify and score native poses (e.g., docking power) on an extended and representative benchmark of protein-nucleotide complexes. A clustering of the MCSS-generated poses is also proposed as a filtering process to select fewer relevant poses. We also evaluate the ability of the scoring function to identify the “true binders” (e.g., screening power) corresponding to the native nucleotide ligands as opposed to the “false binders” associated with the three other non-native nucleotide ligands. Finally, we determine the performance of various solvent models, using an implicit versus a hybrid representation (based on the initial distribution of the crystallographic water molecules) and its impact on the docking and screening powers. By using models with explicit water molecules, we can also evaluate their role and contribution in nucleotide-binding.

## Results & Discussion

Most of the docking methods and their scoring functions have been tested on different benchmarks. These benchmarks have been designed for some specific families of ligands including RNA ligands (46–51). However, the RNA-protein benchmarks include large RNAs (tRNA, rRNA, ribozyme, etc.) where single-stranded RNAs are poorly represented and mostly present in the context of single-stranded regions connected to double-stranded regions. Building the benchmark from a subset of RBPs binding ssRNAs would select optimal and sub-optimal binding sites corresponding to spacer regions. To avoid such bias, we built a benchmark based on the protein-nucleotide complexes currently available in the Protein Data Bank (RCSB PDB (52)).

A previous protein-nucleotide benchmark with 62 complexes was used to evaluate the docking power of three methods: AutoDock (4.2.3), GOLD (5.1), and MOLSDOCK (53). However, the benchmark is mostly outdated with only 40% of complexes with an atomic resolution less than 2.0Å and thus not representative of the currently available structural data. On the other hand, it was tested under biased conditions: the docked region was restricted to the native ligand pose (5Å^3^) and the high-occupancy water molecules of the binding site were preserved within a rigid receptor.

In this study, we use an updated and representative dataset of high-resolution protein-nucleotide complexes in which only nucleotide monophosphate are included (see “Protein-nucleotide Benchmark” & Methods A). The nucleotides are docked in an extended region (17Å^3^) around the binding site where all the inorganic compounds (e.g. metal ions) or organic ligands were removed. Water molecules were present in all the protein-nucleotide complexes, in particular in the binding site. Water molecules were either removed or included before energy minimization (see “MCSS Calculations” and Methods Section B). The resulting optimized proteins are then used as targets in the MCSS calculations. Two charge models (full or scaled charges), two dielectric models (constant or distance-dependent), and explicit representation of solvent or not are used. Four different solvent models were tested, excluding already tested models or irrelevant ones (implicit solvent with constant dielectric (43)). The results are first analyzed in terms of docking power, i.e., whether the method can reproduce the binding mode of the native nucleotide ligand. Then, the analysis is focused on the screening power, i.e., whether the method can discriminate the native nucleotide ligand from the other three non-native nucleotides.

### Protein-nucleotide Benchmark

The protein-nucleotide benchmark includes a non-redundant set of 121 complexes associated with 14 different known molecular functions. Despite the over-representation of proteins binding AMP in the 3D structures available in PDB (72%), all the four ribonucleotides are represented; the three other ribonucleotides are distributed almost equally (Sup. Note 1, Fig. S1). The selection criteria retained to build the benchmark are detailed in Methods A. A series of molecular and energy descriptors compose the features used to characterize the 121 nucleotide-binding sites. The benchmark covers a broad diversity of features that reflect that of the binding modes.

As shown in Fig. 1, the three nucleotide moieties: phosphate, ribose, and nucleic acid base, establish contacts. Although the base contacts are slightly more represented, they are slightly less frequent than those established by the other nucleotide moieties, especially for close contacts (Fig. 1A-B). Thus, the binding specificity dependent on the nucleic acid base’s identity might be weak in some cases (15 protein-nucleotide complexes exhibit no base contact). A similar profile applies to hydrogen bonds (Fig. 1C). The ribose moiety makes fewer contacts (H-bonds or C-C contacts), but they are present in almost all protein-nucleotide complexes (Fig. 1C-D). The stacking contacts (*π*-*π*, *π*-cation, and T stackings) are common in nucleic acid interactions; they are present in half of the benchmark with a variable number of contacts and stacking types (Fig. 1E). Similarly, salt-bridges stabilize the protein-nucleotide complexes; they are present in a bit more than half of the benchmark (Fig. 1F). The distribution of the buried fraction of nucleotide ligands indicates that a major proportion of the benchmark includes above 70% of buried atoms, meaning it corresponds to rather “closed” binding sites (Fig. 1G). On the other hand, nucleotides with a low fraction of buried atoms are more “open” binding sites. A detailed 2D representation of the contacts within the binding site (54) is provided for each protein-nucleotide complex (Sup. Note 1). Some molecular features based on those contacts’ enumeration are further used in the qualitative analysis of the method’s performance (docking and screening powers). Looking more in details at the nucleotide breakdown of the atomic contacts, it appears that A nucleotides exhibit an average number of contacts way higher than the other three nucleotides, suggesting that A binding is stronger (Fig. S2).

**Fig. 1.**
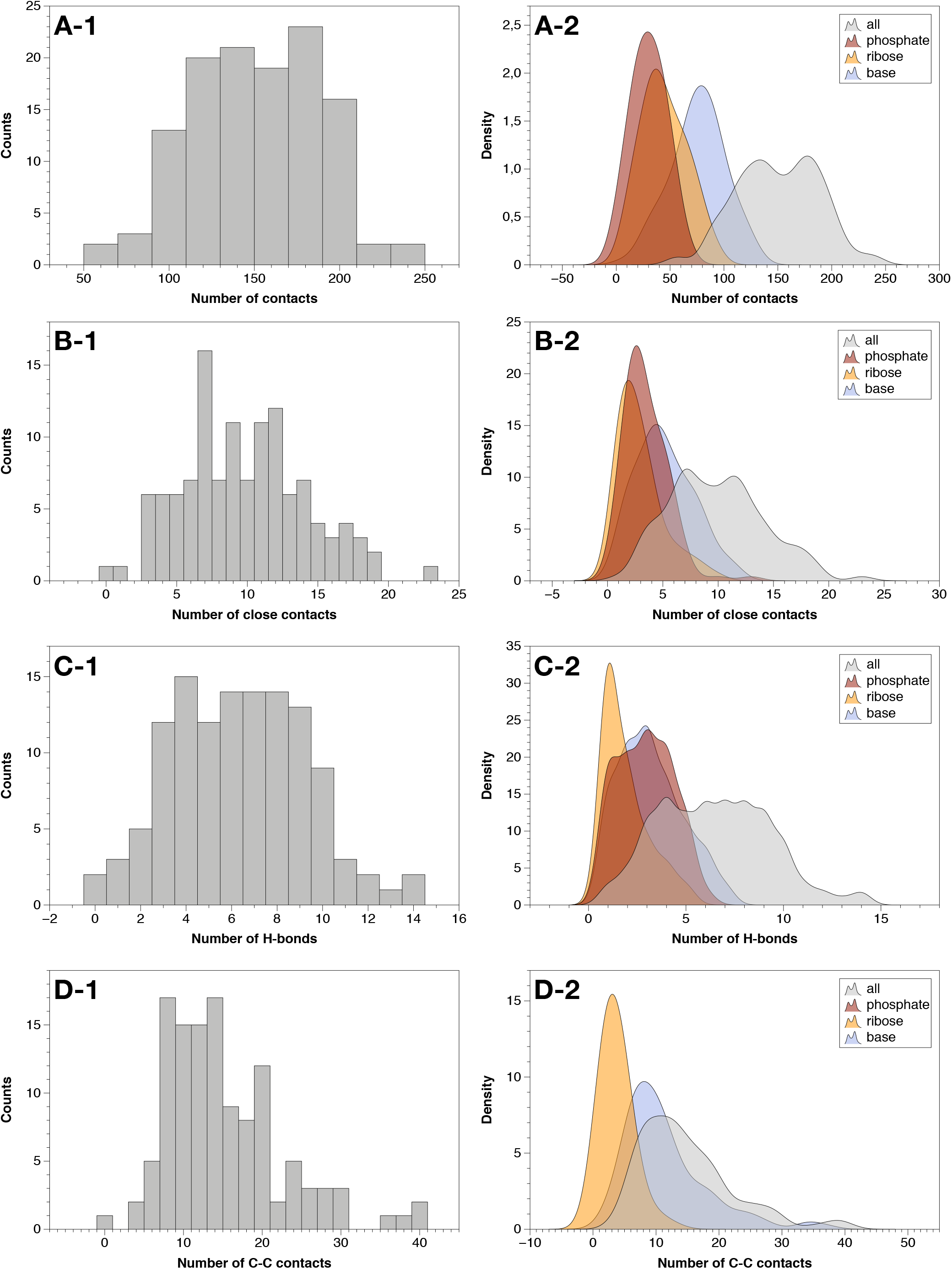
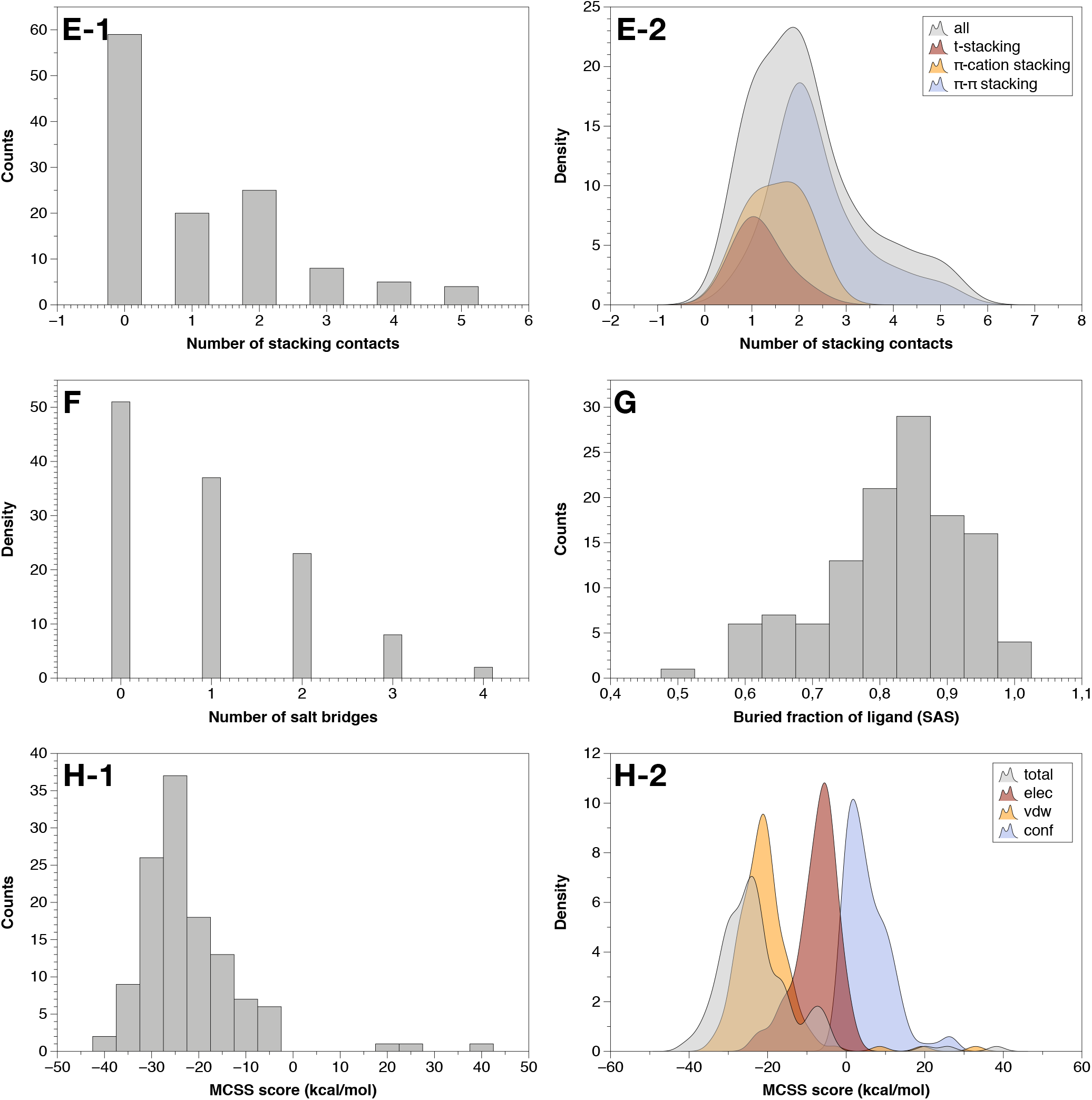
Molecular and energy features of the nucleotide-binding sites from the benchmark of 121 complexes. A-1.: Histogram of the number of contacts; A-2: Smooth histogram with decomposition per nucleotide moiety (base, ribose, phosphate); B-1.: Histogram of the number of close contacts; B-2.: Same as A-2 for close contacts; C-1.: Histogram of the number of H-bonds; C-2.: Same as A-2 for H-bonds; D-1.: Histogram of the number of C-C contacts; D-2.: Same as A-2 for C-C contacts; E-1.: Histogram of the number of stacking contacts; E-2.: Smooth histogram with decomposition per stacking types; F.: Histogram of the number of salt-bridges; G.: Histogram of the buried fraction of ligand (calculated from the solvent accessible surface); H-1.: Histogram of the MCSS scores calculated for the ligands optimized in their binding site; H-2.: Smooth histogram with decomposition per contribution types (electrostatics, van der Waals, conformational). The molecular descriptors associated with the atomic contacts are calculated by BINANA (55); the stacking contributions are calculated from OpenEye (54); the MCSS score is calculated by the scoring function derived previously (43).

The MCSS score is used as a criterion to identify peculiar cases where the interaction between the ligand and the protein is unfavorable (see Methods Section B, Equations 1-4). The score is calculated after minimization by re-inserting the nucleotide within the optimized binding site. A positive score indicates either clashes or a drift of the nucleotide from its position in the X-ray 3D structure. Three nucleotide-protein complexes do exhibit a positive score (Fig. 1H). The penalty term of the score (see Methods Section B, Equation 1) might be more than 10 kcal/mol in some cases, indicating a significant deformation of the nucleotide’s conformation. With the implicit solvent model for nucleic acids (43), the van der Waals energy contributes more to the score than the electrostatic energy (Fig. 1H).

All the high-resolution protein-nucleotide complexes of the benchmark include water molecules around the protein surface and the binding region. When these crystallized water molecules are conserved during the minimization, the protein coordinates deviate from the experimental ones by around 0.5Å on average. It is close to 1.0Å when the water molecules are removed (Fig. 2A), indicating that the presence of water molecules preserves slightly more the original coordinates from the X-ray structures. Nevertheless, the number of water molecules around the ligand does vary from one complex to another: from 1 to 18 (Fig. 2B). Base and phosphate contacts with water molecules are more frequent in the benchmark, but the ribose moiety cumulates more contacts when they are present. The minimization does induce displacements in the position of the water molecules in the binding region mostly due to the removal of the ligand; the number of water molecules within the binding region may then slightly vary after minimization (Fig. 2C-1). Because of their small molecular weight, the water molecules may also deviate significantly from their initial positions (Fig. 2C-2).

**Fig. 2.**
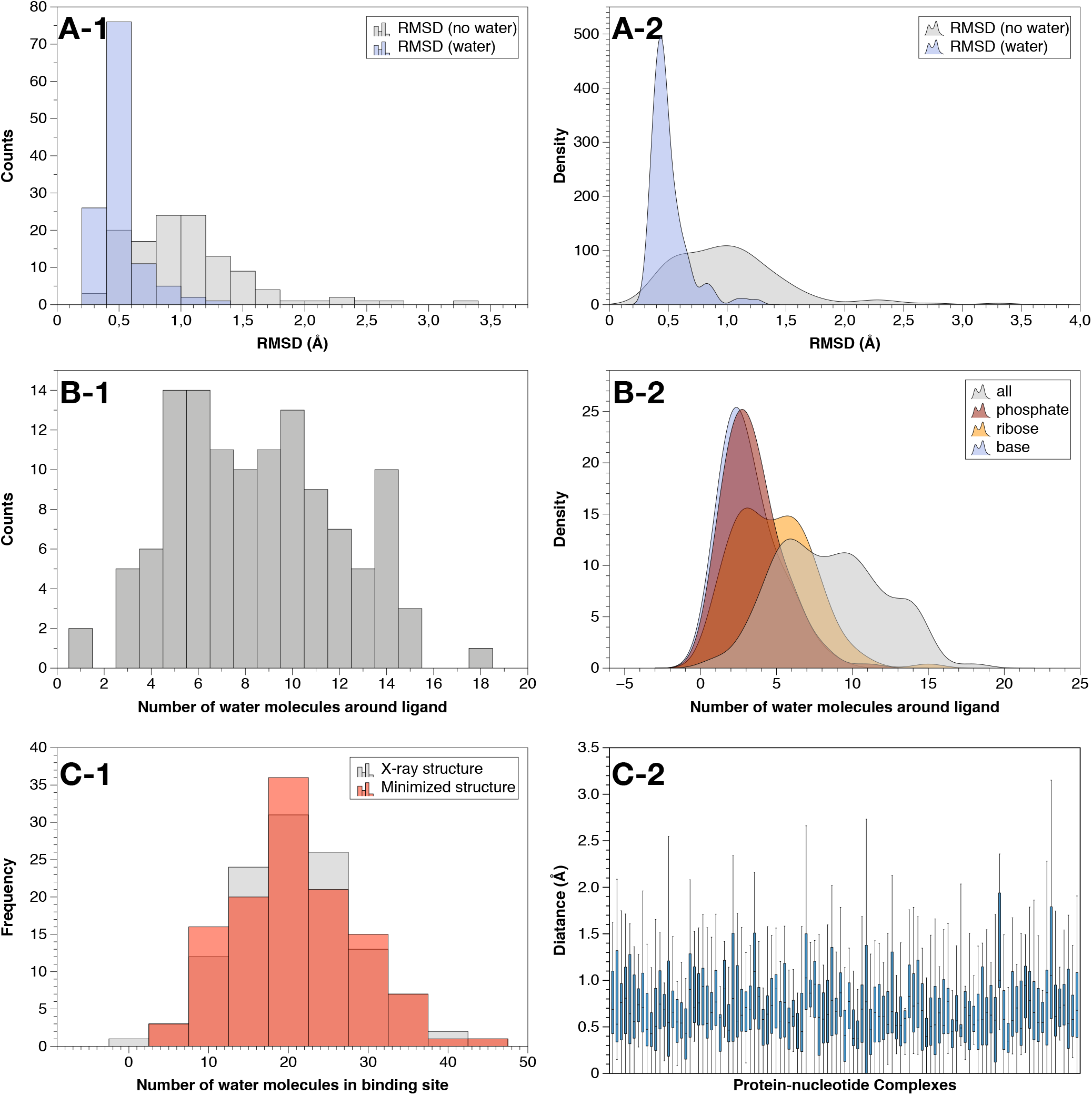
Distributions of water molecules and impact on the binding sites. A-1.: Histogram of RMSD in presence/absence of water molecules; A-2.: Same as A-1 with a smooth histogram; B-1.: the number of water molecules around the ligand (distance cutoff of 4.0Å); B-2.: Same as B-1 with decomposition per nucleotide moiety; C-1.: Number of water molecules within the binding site as defined in MCSS by the box parameters (see Methods Section B); C-2.: displacements (Å) of water molecules from their crystallized positions.

The benchmark consists of a representative set of protein-nucleotide complexes in terms of atomic contacts and binding modes with variable contributions from each nucleotide moiety. However, it includes two identified biases associated with the nucleotide type: A is overrepresented and expected to bind more strongly to the protein.

### Nonbonded Models

Several phosphate group models are used in the MCSS calculations to determine the optimal parameters for mapping nucleotides at the protein surface. We used five different phosphate models corresponding to 5’ patches (R010, R110, R210, R310, and R410) that differ by the valence and charge of the phosphate group (Fig. 3). The R010 patched nucleotide corresponds to the standard nucleotide residue defined in CHARMM, and it is the only fragment with an unfilled valence shell at the 5’ end (Fig. 3). All the partial charges on the phosphate groups are derived from the CHARMM parameters. They correspond to the original CHARMM charges or derived from them based on Manning’s theory of counterion condensation to account for the partial neutralization of the negative charges of polyelectrolytes solution (56). In this latter case, the net charge on the phosphate group is scaled down according to the implicit solvent model previously used in MCSS calculations performed on nucleic acids (43).

**Fig. 3.**
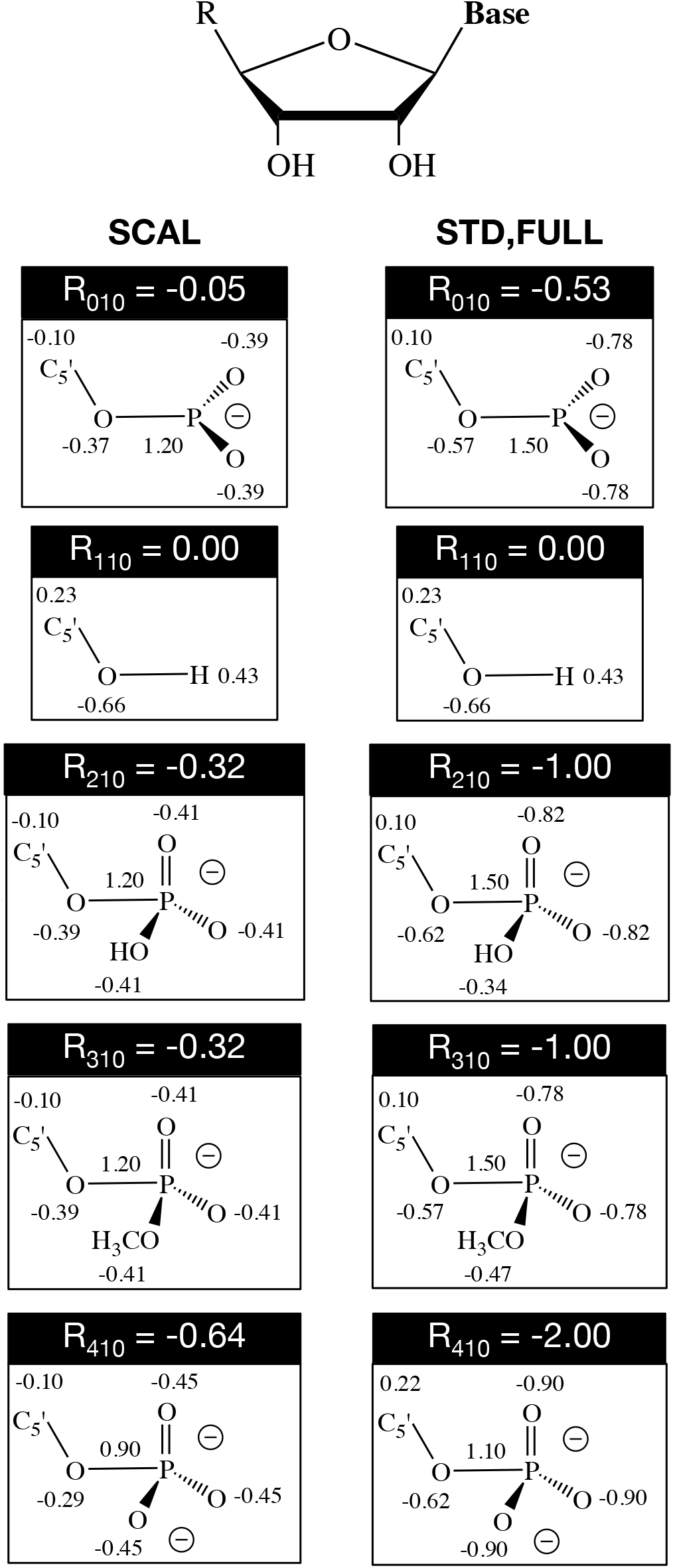
Nonbonded models used in MCSS calculations. The R group corresponding to the 5’ end of the nucleotide includes five flavors: R_010_ (standard nucleotide residue), _R110_ (5’OH patch), R_210_ (5’PO_4_ H^−^), 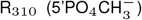, and 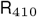 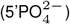. Three solvent models are used: the SCAL model is based on reduced charges on the phosphate group according to Manning’s Theory (56) and applied to nucleic acids (43); the “STD” (standard) or “FULL” models are based on standard charges. The electrostatic contribution to the interaction energy is calculated based on a constant dielectric formulation for the “FULL model”. The SCAL and “STD” models are based on a distance-dependent dielectric model. The van der Waals contribution is calculated using the standard CHARMM27 potential energy function (57).

The “SCAL” charges model (Fig. 3 - left) is combined with a distance-dependent dielectric (Equation 4) with or without water molecules: SCAL and SCALW, respectively. The default charges model “STD” or “FULL” (Fig. 3 - right) is combined with explicit solvent representation and a distance-dependent dielectric (Equation 4): STDW, or with a constant dielectric (Equation 3): FULLW.

### Poses and hits

The protein target is prepared by defining a binding region centered around the ligand and subjected to the mapping of the nucleotide fragments described above (Fig. 4). Final poses generated by MCSS (e.g. minima) are ranked by their score (Equations 1-4) in ascending order. Although a clustering step is performed iteratively during the MCSS calculation, the default RMSD cutoff value is low (0.5Å) to guarantee a fully extended search at each iteration before re-ranking the intermediate poses and the minima. As a consequence, some minima may still exhibit some degree of geometrical redundancy. Poses coming from different initial positions may converge to similar minima while still being above the RMSD cutoff value. These minima may exhibit large discrepancies in terms of score, especially when using implicit solvent models where small deviations in coordinates may significantly alter the interaction energy with the protein target. This redundancy may negatively impact further statistical analysis as very similar poses can have a drastically different score. Such a bias can be avoided through clustering analysis based on an approach similar to that already used by MCSS (see Methods C).

**Fig. 4.**
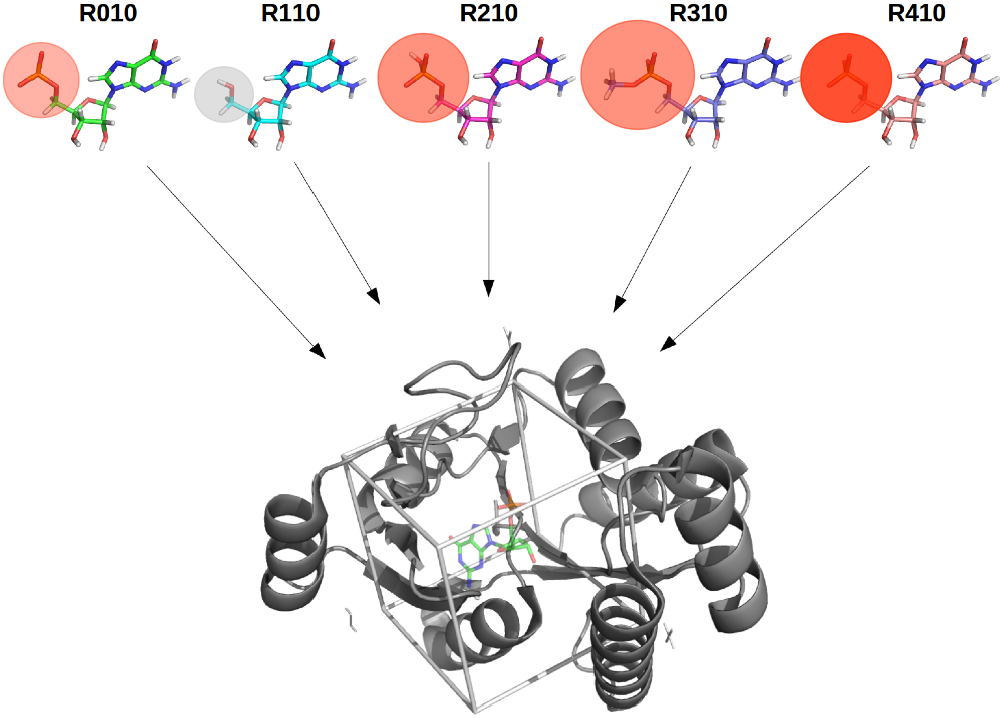
Schematic description of the series of MCSS calculations performed on each protein target. The chemical structure of each 5’ patched nucleotide is indicated: R010, R110, R210, R310, R410. The phosphate group is circled: the bigger the sphere the bigger sterically, the darker red the more negative charge (the grey color indicates a null charge). The protein target is represented in cartoon mode with the indication of the cubic box corresponding to the explored region.

The five types of patched nucleotides (R010 to R410) are mapped at the protein surface around the binding region entered on the position of the ligand in the X-ray structure (Fig. 4). The raw distributions include up to several thousands of poses. The total number of poses generated depends on the solvent model and on the phosphate patch to a lesser extent. The presence of explicit water molecules reduces in part the molecular volume accessible for nucleotides. Thus, the number of poses generated with the SCAL model is much larger than that generated with any of the models with explicit solvent: SCALW, FULLW, and STDW (Fig. 5).

**Fig. 5.**
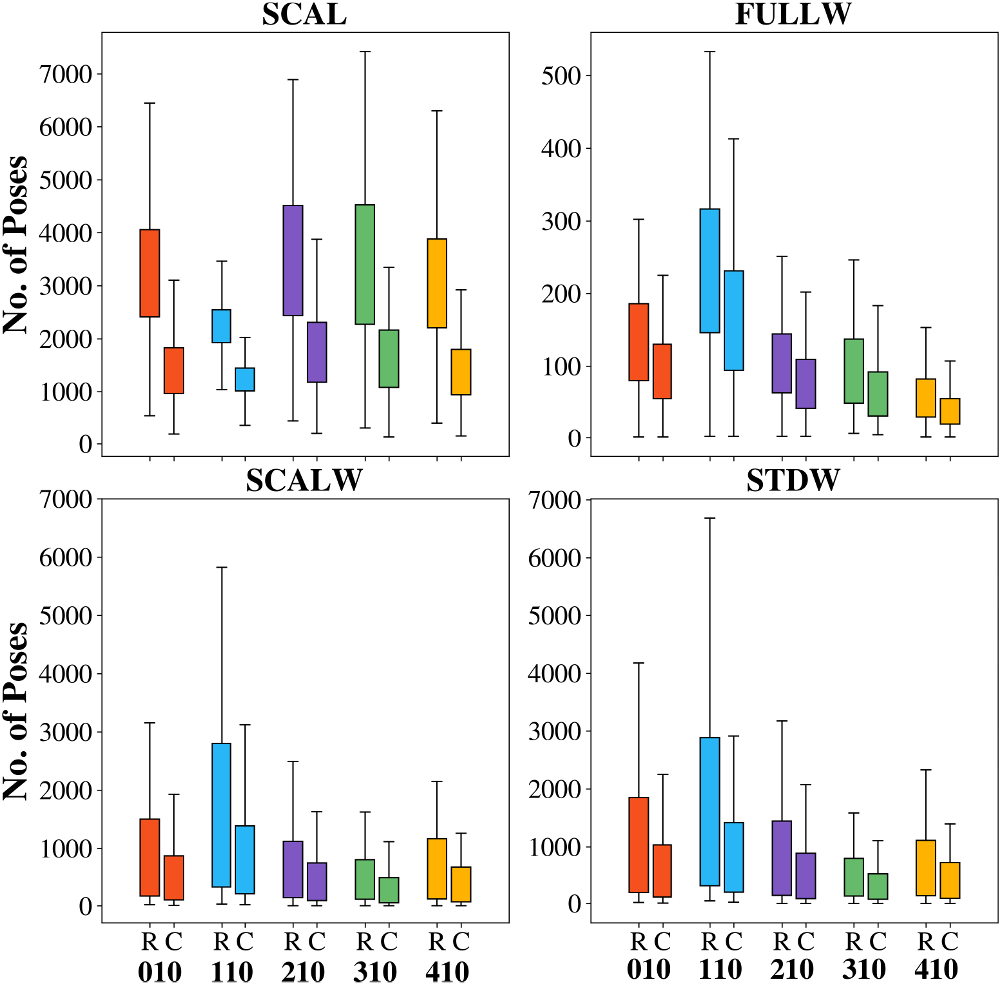
Boxplot representation of the number of poses generated for the 121 protein-nucleotide complexes for each 5’ patched nucleotide (010, 110, 210, 310, 410). Results for raw (R) and clustered (C) distributions are shown.

In the SCAL model, an average of 3000 to 3500 minima is obtained in the raw distribution (Fig. 5). The more open binding regions can accommodate 4000 poses and up to 6000-7000 poses. The clustered distribution is reduced by almost half for any patched nucleotide. The only exception is for R110, which corresponds to a nucleoside: the smallest fragment in size (Fig. 4). The more charged the nucleotide (R010 to R210, and R310), the more poses it generates except in R410, where the higher charge can produce, in several cases, unfavorable interactions with negative charges at the protein surface. As a result, more poses fail to pass the energy threshold value of the MCSS score. This trend is valid for both the raw and clustered distributions. R110 generates a less number of poses, even if it is the smallest fragment. Since it has a zero charge, it has a lower electrostatic contribution to the MCSS score. In this particular solvent model, there is a delicate balance between the charge and the energy threshold. A zero charge will cancel most of the electrostatic contribution and eliminates many poses. On the other hand, a high negative charge will induce unfavorable interactions, also leading to poses removal. The R210 and R310 are the patched nucleotides that generate more poses (raw distribution) and more non-redundant poses (clustered distribution).

In the SCALW model, we see the direct effect of including explicit water molecules. As expected, the number of poses is significantly reduced in both the raw and clustered distributions by 4 to 7 times. The only exception is the R110 patched nucleoside, where the number of generated poses is almost equivalent in the raw and clustered distributions between the SCAL and SCALW models (Fig. 5). Although the presence of water molecules tends to reduce the accessible volume of the binding region (SCALW/SCAL), the total number of poses is equivalent between both models. We can attribute this specificity to the small size of the R110 fragments. As a consequence, it appears that the lower number of poses for the other patched nucleotides is mostly due to their size, even though the accessible volume is reduced with respect to the SCAL model. Accordingly, the number of poses for R010 (the second smallest nucleotide fragment in size) is slightly higher in both distributions compared to that of R210, R310, and R410. On the opposite, the biggest patched nucleotide R310 exhibits a lower number of poses among all nucleotides. The STDW model shows a very similar profile for the absolute values in the number of poses and the difference in the number of poses depending on the nucleotide patch. Here, the choice of the charge model SCALW/STDW has no significant impact on the total number of poses. Instead, the dielectric model does alter downward the number of poses in the FULLW model; the profile in the relative number of poses between the patched nucleotides is very similar to those of the other models with explicit solvent (SCALW and STDW) except for R410, which is the patch associated with the lowest number of poses. The electrostatic contribution to the interaction energy accounts for this massive decrease in the number of poses. The FULLW model is the only one in which this term is based on a constant dielectric formulation, meaning that no screening effect is included between charges. As mentioned above, for the R410 patched nucleotide, an excessive charge density can make the electrostatic contribution less favorable. It is associated with the charge in the SCAL model (specifically for R410) while related to the absence of screening effect in the FULLW model. Consistently, the lowest number of poses is obtained with the FULLW model and the R410 patched nucleotide which carry the higher unscreened charge.

The clustering method applied to the raw distribution of poses makes it possible to eliminate a geometrical redundancy. This redundancy varies depending on the choice of the solvent model and patched nucleotide to a lesser extent. It is much higher in the SCAL model, where the absence of water molecules multiplies similar poses. Among the other solvent models that include water molecules (SCALW, STDW, and FULLW), the redundancy is very alike. However, the dielectric model has a strong impact on the number of poses, which is reduced by 5 to 35 times in the FULLW model; less than 100 poses are present in the clustered distributions. The clustered distributions in the SCALW and STDW models include around 500 poses, while the SCAL model retains more than 1000 poses. However, the lone number of poses does not give any indication about the number of native-like poses.

We define as a native-like pose or hit those poses generated by MCSS with an RMSD less than 2Å to the ligand’s experimental coordinates in the reference PDB (see Methods Section B). The fraction of hits over the entire MCSS distribution for all solvent models and patches is shown in Figure 6. This fraction is similar for all patches in each of the four models, except for R310. The patch R310 carries a methyl group in one of the phosphate oxygen. This group confers the ability to establish more hydrophobic contacts than other patches. The SCAL model shows a significantly lower fraction of hits than solvated models despite a much larger number of generated poses (Fig. 5); both the raw and clustered distributions are more scattered in the absence of water molecules. In the case of solvated models, SCALW and STDW hit fractions are very similar. On the other hand, the FULLW model has more cases where no hits are found for some proteins; it is revealed by the displacement to zero of the first interquartile section for the boxplots.

**Fig. 6.**
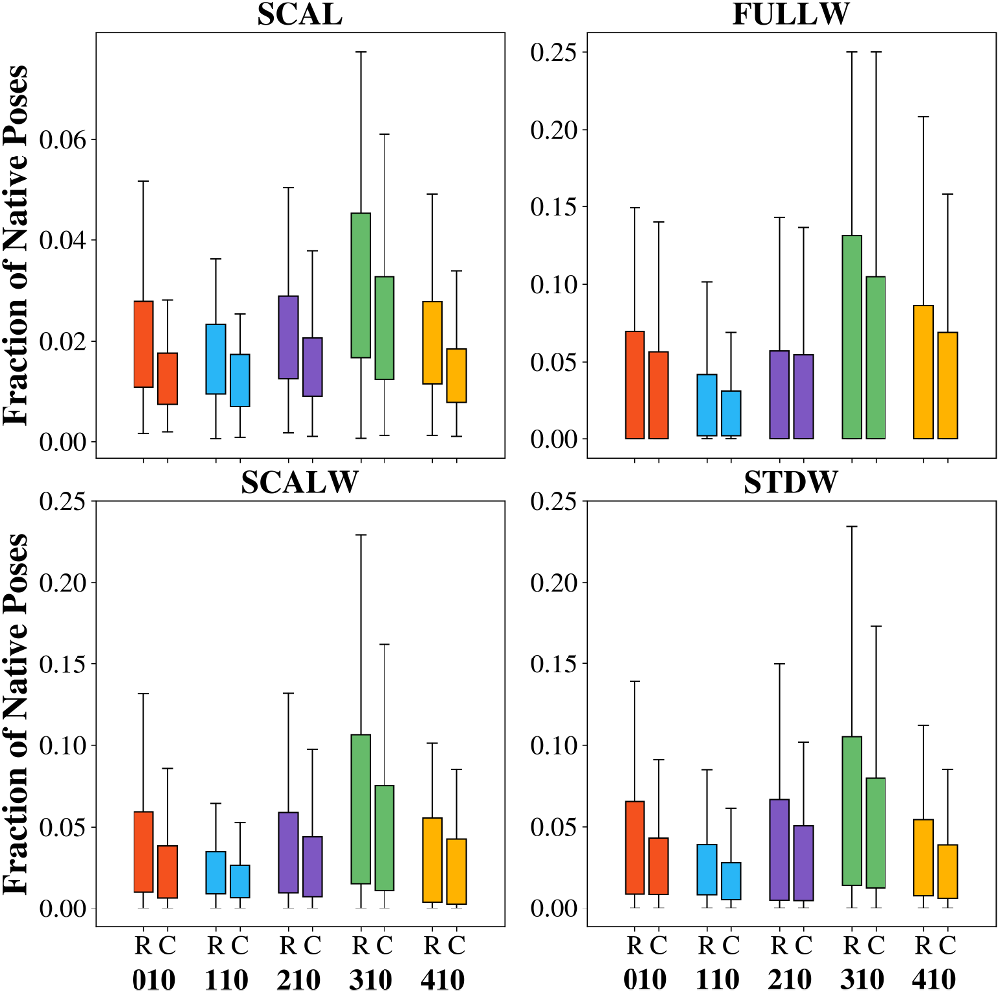
Boxplot representation of the fraction of native poses generated for the 121 protein-nucleotide complexes for each 5’ patched nucleotide (010, 110, 210, 310, 410). Results for raw (R) and clustered (C) distributions are shown.

### Docking Power

The MCSS predictions are ranked according to the success rate for the identification of at least one native pose obtained on the full benchmark in the Top-*i* (Top Native in the best ranked i poses) with *i* in a range from 1 to 100. The results are shown for the range of Top-1 to Top-100 with the intermediate ranks: Top-5, Top-10, and Top-50 (Fig. 7). For each patch (from R010 to R410), the number of protein-nucleotide complexes which are predicted with a native pose in the Top-*i* follows a similar trend between the different models.

**Fig. 7.**
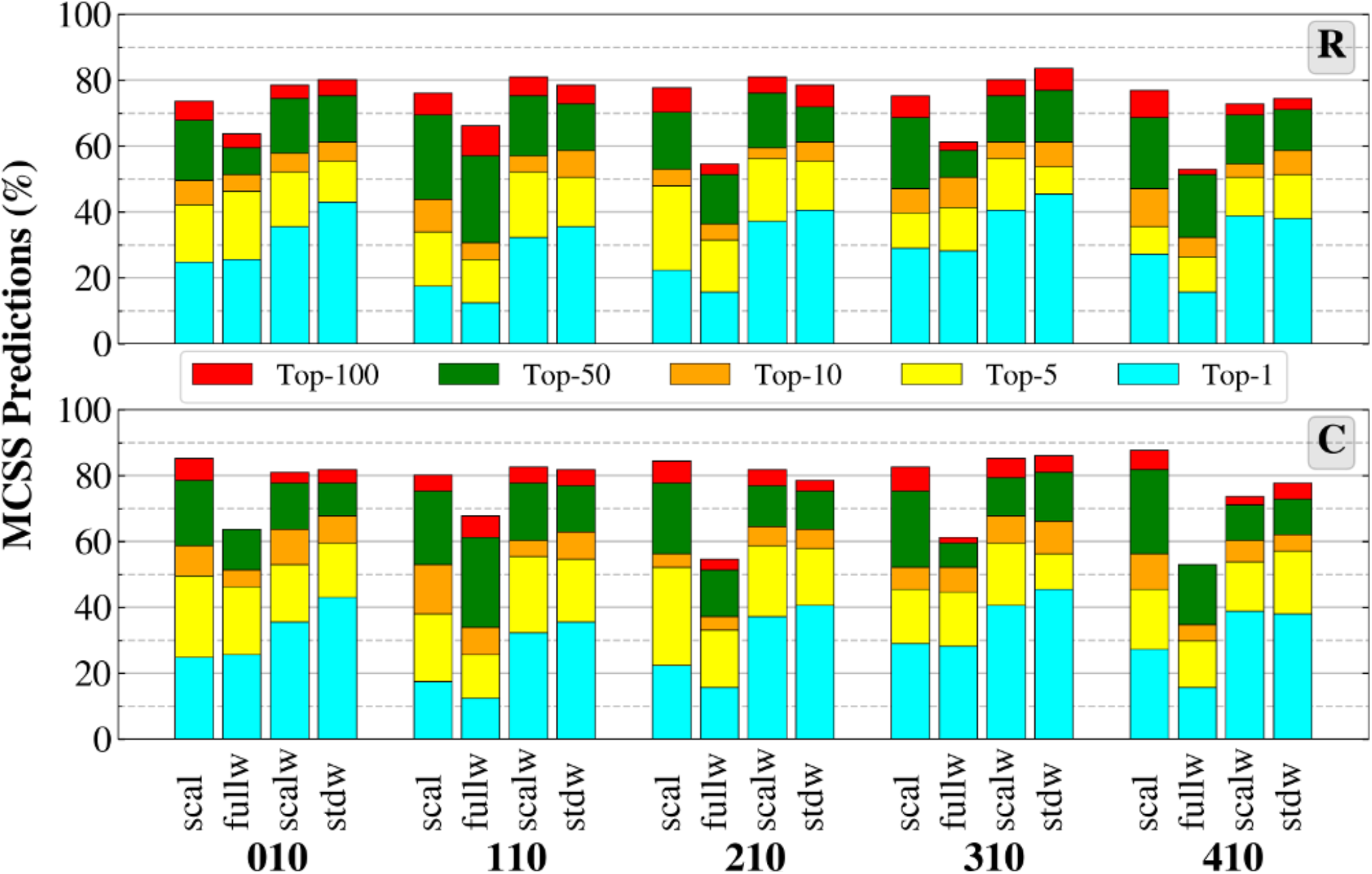
Stacked histogram representation of the Top-*i* ranked native poses generated for the 121 protein-nucleotide complexes for each nucleotide patch. Result on raw (upper) and clustered (bottom) distributions are shown.

The SCAL and FULLW models exhibit a lower success rate from Top-1 to Top-100 except for the patch R410, where the SCAL model performs slightly better for a few cases in the Top-100 but underperforms in all the other Top-*n*. The SCALW and STDW models outperform for all the patches; the FULLW systematically underperforms in almost any Top-*n* except in the Top-1 to Top 10 for R010 and R310, where it slightly outperforms the SCAL model. Among the two best models, the STDW model slightly outranks the SCALW model in the Top-1 and Top-10 for all the patches (except for R310 where the performance is equivalent for the Top 10), while the performance is pretty similar for the Top-50 and Top-100. The best performance is obtained for R310 with a success rate of 45% in the Top 1, a bit more than 60% for the Top 10, and more than 80% in the Top-100. The clustering does not change the general trends observed in the raw distributions. It slightly increases the performance in the Top-100 and, to a lesser extent, the lower Top-*i*. The use of R310 improves a bit the performance of the SCALW model over the STDW model in the Top-10.

Additional nonbonded and water models (available within CHARMM) can be tested as well as external scoring functions. External methods (e.g., Vina) are not implemented into MCSS, but they can be tested through single-point calculations on the MCSS distributions. Some of the tested scoring functions are specialized on some class of ligands, e.g. nucleic acids for ITscorePR (58), or Δ_*vina*_*RF*_20_ (59) or more generic for Autodock Vina (60), or Vinardo (61). The results of the comparison show that the standard MCSS scoring function corresponding to the SCAL model performs at the same level than Δ_*vina*_*RF*_20_ and Vina and slightly better in the Top-10 to Top-100 for the clustered distribution (Sup. Note 3). The Vinardo scoring function performs slightly better than the three others on both MCSS distributions (Fig. S4). However, the MCSS scoring function associated with the STDW model outperforms all of the external scoring functions in the Top-1 to Top-10 in both raw and clustered distributions (Fig. 7). In the CASF-2016 benchmark, the docking power ranges from around 30% to 90% for a variety of scoring functions (62). The docking power is around 90% for both Vina and Δ_*vina*_*RF*_20_. On the current benchmark, their performance is only 33%, indicating the challenging task to score charged ligands such as nucleotides. Vinardo performs slightly better (42%) and also MCSS-STD (45%).

One of the contributions to the MCSS score is the conformational penalty term (Equation 1) corresponding to the deformation of the fragment from its optimal conformation. Although this term is generally a minor contribution, it may vary depending on the nonbonded model. We can compare the torsion angles observed in the MCSS minima with respect to the known ideal values and values observed in the native bound conformations of the nucleotides from the benchmark (Sup. Note 2, Fig. S3). The absence of water molecules in the SCAL model reveals a few biases where, for example, the syn conformation is more populated than expected as compared with the experimental or the ideal values collected from the experimental structures of nucleic acids (63, 64). The SCALW model is also biased, and the STDW but to a lesser extent; only the FULLW model is exempted. Another common bias to all models (except for the FULLW model) is the overrepresentation of the C2’-endo conformation for the ribose while the initial conformation is always a C3’-endo conformation. It is partly due to the nonbonded model and the absence of a full solvation of this nucleotide moiety. In FULLW, the C3’-endo/C2’-endo representation is more balanced, but the phosphodiester backbone (torsion angles *α* and *β*) deviates from the optimal values because of some distortion of the phosphate group, which is highly charged and tend to stick closely to the protein surface in the absence of any screening effect (constant dielectric model).

The performance was then analyzed by nucleotide type. Since the adenosine is over-represented in the benchmark, it generally follows the global trend described above (Fig. 8). However, the performance for guanosine decreases for the larger patches R210 to R410, whatever the model used. Only the smaller patches R010 and R110 give a similar performance or better in some cases; the success rate with R110 is even better from Top-1 to Top-50, indicating the existence, as discussed before, of a size effect that drives down the perfor mance (guanine is slightly more voluminous than adenine). Consistently, the performance generally improves for pyrimidines (C or U), which are smaller than purines. On the other hand, the performance is degraded in the smaller nucleoside fragments (R110) that do not carry any phosphate group. However, in the case of U nucleosides and only for the STDW model, the performance is better than that for A nucleosides. The pyrimidic nucleotides are better predicted, especially for the two best models SCALW and STDW with R310. The predictions are equivalent or degraded for the more highly charged patch R410, especially with U. The analysis of the clustered distributions confirms the observed trends of the raw distributions, with improved performances reaching 90% to 100% for the Top-100 in a larger number of models and patches (Fig. S5).

**Fig. 8.**
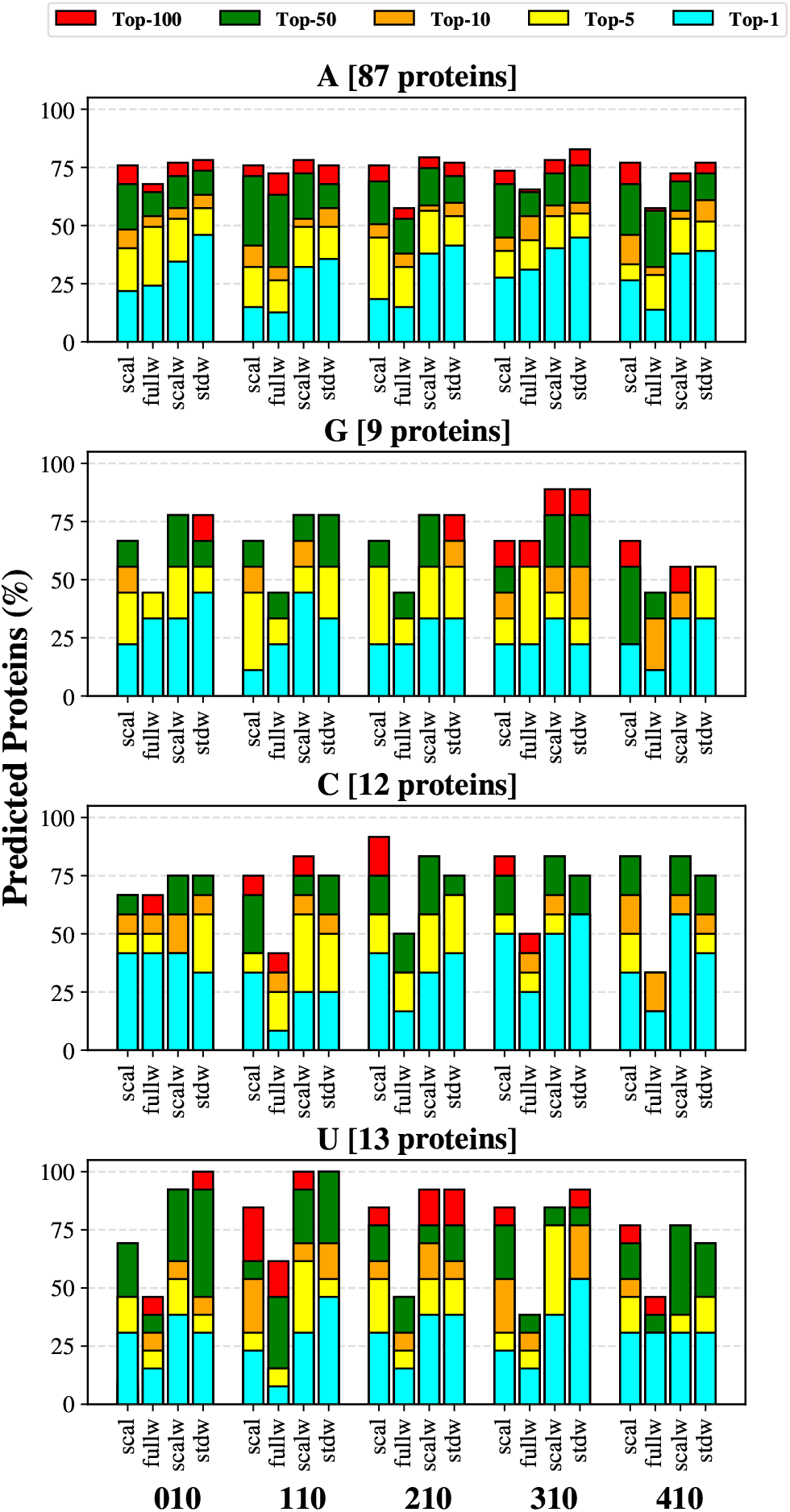
Nucleotide breakdown of the success rates obtained for each nonbonded model and patches combination. The data are shown for the raw distribution (without clustering) and each Top-*i*.

### Screening Power

In the benchmark, we assume that the crystallized nucleotide is always the native and more specific nucleotide, i.e. it is the only nucleotide ligand with a detectable affinity or the best binder among the four nucleotides. Based on this assumption, we can define a screening power as the ability to rank the native nucleotide ahead of the other three nucleotides. In that case, we will refer to optimal predictions as the native pose is identified and the native nucleotide is ranked first. Since the scoring function is still an estimate and raw approximation of the relative binding energy, we consider as good predictions the cases where the native nucleotide is predicted within a 2 kcal/mol range from the best ranked non-native nucleotide. This threshold value corresponds to a maximum offset of 2 kcal/mol in 90% of the benchmark (STDW model) where the offset is defined as the difference between the best-ranked pose whatever the nucleotide type and the best-ranked pose for the nucleotide corresponding to the native ligand (Fig. S6).

The other predictions are considered poor predictions even if native poses are found for the native nucleotide. As an illustration, we show the results obtained for one protein-nucleotide complex (PDB ID: 1KTG) for both SCAL and STDW models (Fig. 9). The best-ranked nucleotide is the native one (A) in the STDW model; other poses of the native nucleotide are also identified (Top-5, Top-10, etc.) but only one is within the 2 kcal/mol score range (good prediction). Some of the poses corresponding to non-native G nucleotides are within the MCSS score range of 2 kcal/mol. In the STDW model, the prediction is optimal since the best-ranked pose does correspond to the native nucleotide. In the SCAL model, the pose with the best score corresponds to a non-native G nucleotide, but the Top-1 for the native nucleotide is within the 2 kcal/mol range; it is not considered as an optimal prediction but as a good prediction. The other poses for the native nucleotide, which lie out of the 2 kcal/mol range (Top-5, Top-10, etc.), correspond to poor predictions.

**Fig. 9.**
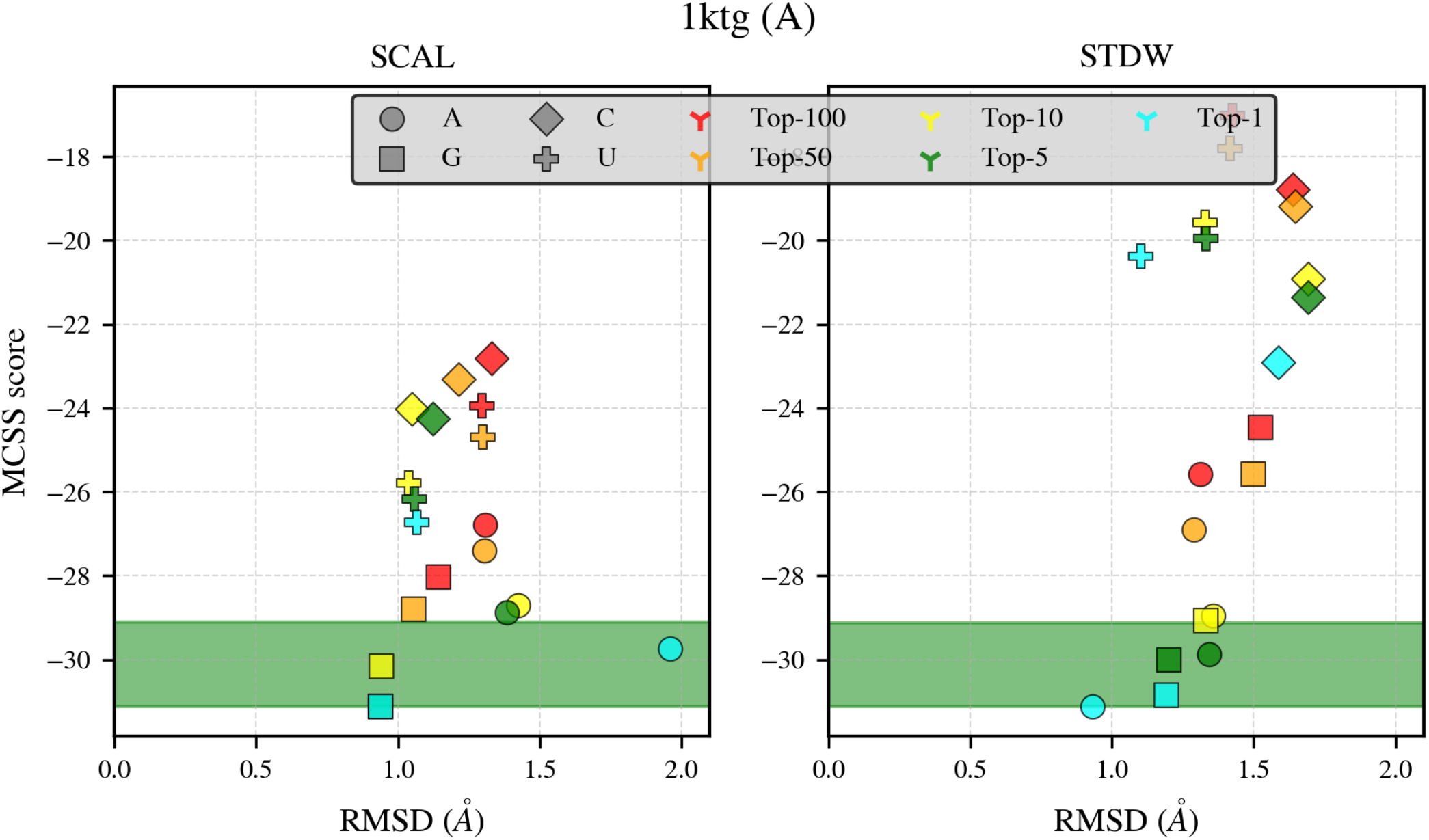
Binding selectivity predictions for 1KTG. Left: SCAL model (R310); right: STDW model (R310); the interval of MCSS scores corresponding to a 2 kcal/mol range is indicated by the green bar. Each Top-*i* for i > 1 is represented by a single point that corresponds to the average RMSD and score of all its members.

We selected the best combination between the nonbonded model and patch for comparison to the standard SCAL model (without explicit solvent) to evaluate the impact of includ ing water molecules. The STDW model with the R310 patch gives the best performance, slightly better than the SCALW model, particularly in the Top-1 predictions (Fig. 7). The comparison between the SCAL and STDW models shows a significant gain of performance with explicit solvent (Fig. 10).

**Fig. 10.**
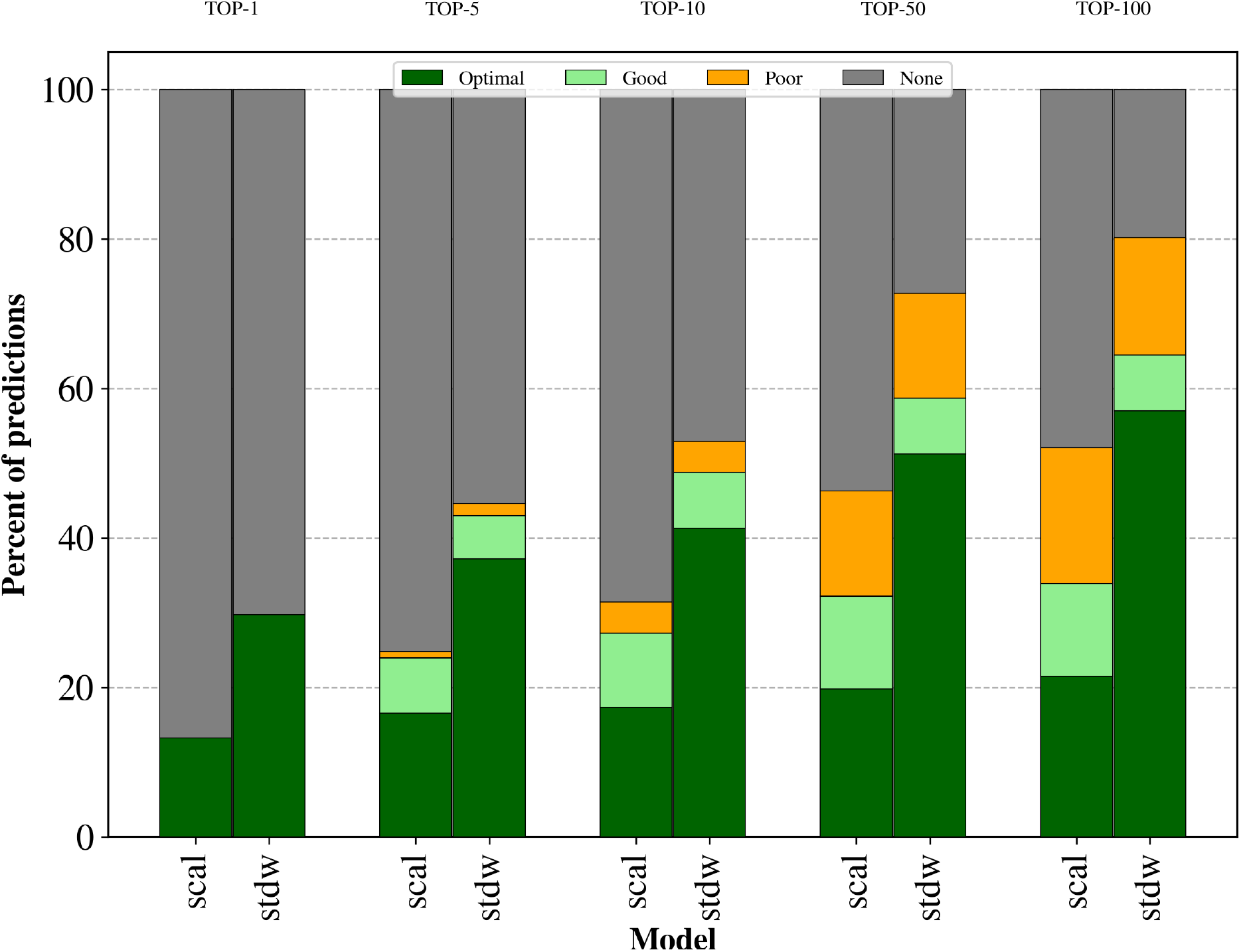
Binding Selectivity Predictions. Optimal: native nucleotide as the best ranked; good: native nucleotide ranked within a 2 kcal/mol range from the best ranked non-native nucleotide; poor: native nucleotide ranked out of the 2 kcal/mol range.

In the optimal predictions, the STDW outperforms by 15 to more than 30% from the Top-1 (Fig. 10A) to Top-100 (Fig. 10E), respectively. In all Top-*i*, the STDW optimal predictions always exceed the SCAL total predictions. Moreover, the ratio of optimal/good predictions is always much higher in STDW. The proportion of poor predictions is tiny up to Top-10. These results indicate that the STDW model has a better screening power, i.e., it predicts more protein-nucleotide complexes and more accurately the native nucleotide than the non-native ones. This better performance is not only imputable to an unbalanced number of predictions between the two models (cases where the STDW provides prediction while the SCAL model does not provide any) but to a more discriminatory power when comparing only the cases which generate predictions in both models (Fig. S7). The distributions of scores between the two models show two different profiles. In the SCAL model, G nucleotides tend to be slightly better scored, then A and pyrimidines with lower scores. However, the average score deviation is within a small range (Fig. S8). In contrast, A nucleotides are better scored in the STDW model while the other three nucleotides have similar distributions. Another difference is the much more extensive range of scores for all four nucleotides. The more favorable scoring of A is consistent with more tightly binding modes, a known bias of the benchmark as mentioned previously (Fig. S2). The nucleotide breakdown of the screening power does show there is no significant difference in performance between A and the other three nucleotides, although it is slightly better in the Top-100 (Fig. S9). In the SCAL model, G nucleotides are better scored in all Top-*i* than pyrimidines in particular. It suggests that the bias towards a better scoring of G and to a lesser extent A contributes to a degraded screening power of the SCAL model. The current scoring functions (tested on the CASF-2016 benchmark) do not exhibit high screening powers, which reach 30% or a bit more than 40% for the highest success rates in the Top1% and a bit more than 60% in the Top10% (62). The comparison with the results of this study (Fig. 10) is risky because the Top1% or Top10% would represent a two- or three-fold number of poses (Fig. 5) with respect to the approximate 1000 poses generated in the CASF-2016 scoring benchmark (62). Furthermore, the molecular diversity of the four nucleotides is limited to the few atoms of the nucleic acid base, making the discriminatory scoring much more challenging.

### Molecular Features

To better understand the role of solvent and other molecular properties on the predictions, we define a series of features that are used to assess the performance or lack of performance for some specific associations of nonbonded models and features qualitatively. We classify the features into three main groups related to:

1. The binding site properties (volume, number of water molecules, metals, other nucleotidic fragments).
2. The conformational properties (purine/pyrimidine, syn/anti).
3. The interaction properties (contacts, clashes, stacking, salt bridges).

The intrinsic properties of the binding site (evaluated from the experimental structure) make it more prone to get good or poor predictions or none. Several protein-nucleotide complexes fail to be predicted at all in the Top-100, representing from 16 to 45% for the raw distributions depending on the nonbonded model and patch used, and a minimum of 12% for the clustered distributions (Figs. 7 & 8). However, only two complexes do not generate any predictions in the Top-100 for any model and patch. To have more examples of negative predictions, we analyze the features based on the presence/absence of predictions in the Top-10. Without any distinction from the model and patch, 17 complexes do not generate any prediction in the Top-10. We consider that a given feature has a significant impact on the prediction when it is found associated with the absence of prediction at a higher frequency than that in the benchmark (Sup. Note 4, Table S1). The feature that correlates significantly with the absence of prediction is a low volume of the binding site (Fig. 11), as calculated by PyVOL (65) (see Methods Section E). Metal ions usually stabilize the phosphate group and occupy some volume in the binding site (it is correlated with a low volume of the binding site and a low number of water molecules). Although it is removed from each protein target, its absence in the calculations is not a negatively impacting feature.

**Fig. 11.**
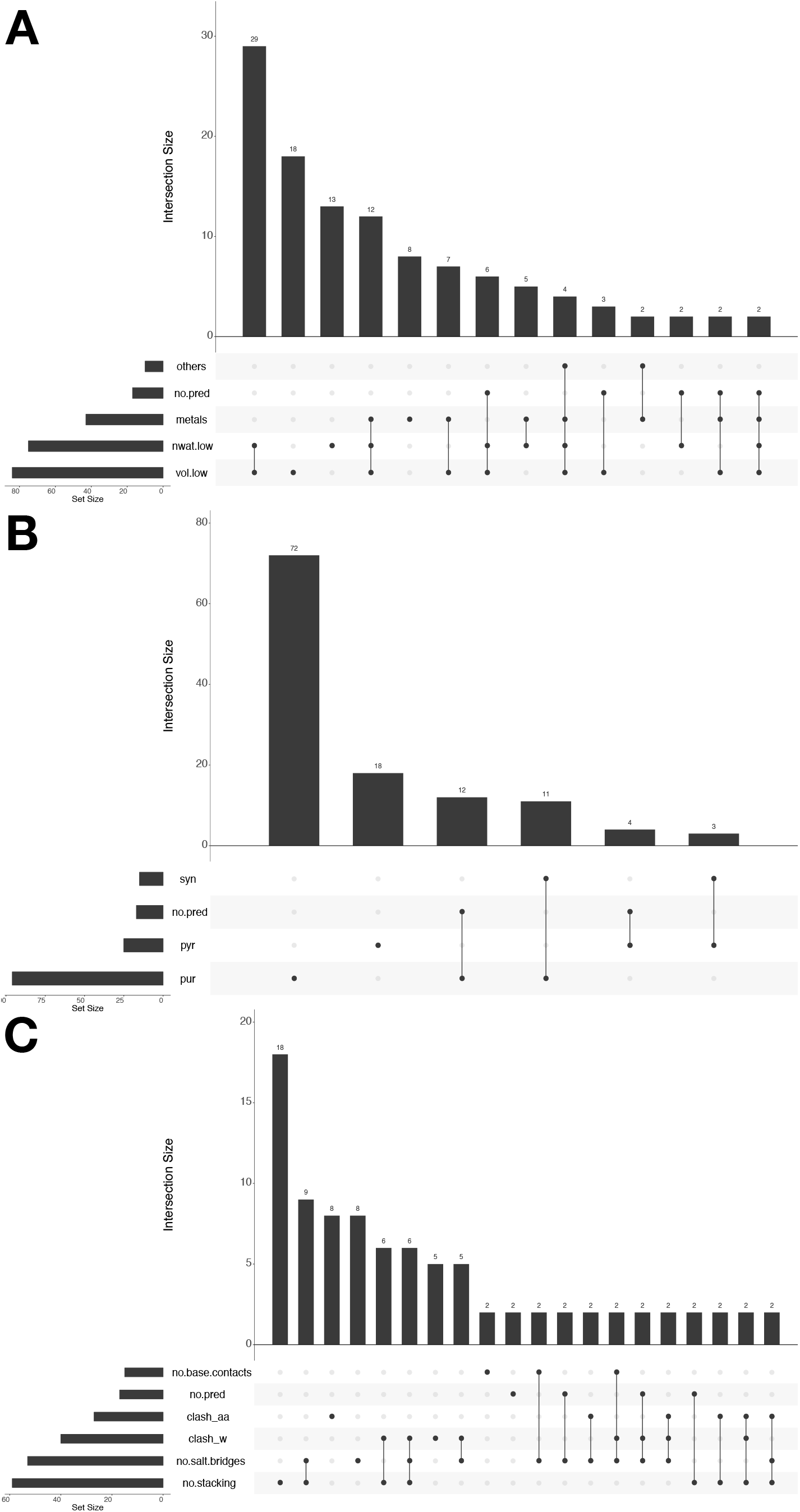
UpsetR diagram of features for the Top-10 predictions. A. binding site features. B. conformational features. C. interaction features. others: presence of additional nucleotidic (nucleic acid) fragment in the binsing site; no.pred: no prediction; metals: presence of metal(s) in the binding site; nwat.low: presence of number of water molecules below the threshold value; vol.low: volume of the binsing site below the threshold value; syn: syn conformation of the nucleic acid base; pyr: pyrimidine; pur: purine; no.base.contacts: absence of contacts with the nucleic acid base; clash_aa: clash(es) with amino-acid residues; clash_w: clash(es) with water molecules; no.salt.bridges: absence of salt-bridge; no.stacking: absence of stacking. Only the intersections with more than one member are shown.

Among the conformational features, none is an impacting feature. Only the pyrimidines have a very tiny overrepresentation in the non-predicted cases (Sup. Note 4, Table S1). Even if the pyrimidic nucleotides are generally better predicted than purinic ones (Top-1, Top-5, Top-50, and Top-100), the non-predicted cases in the Top-10 are slightly more represented. The predictions in the Top-10, in particular for C nucleotides are more sparse in a number of model-patch combinations (Fig. 8). Three interaction features are negatively impacting the performance: the absence of salt bridges, the presence of clashes with water molecules, and to a lesser extent, the absence of stacking contact (Fig. 11). Among these latter contacts, the *π*-*π* interactions are those which more contribute to the negative impact on the predictions (Fig. S10). The presence of clashes with water molecules might induce some deviations of water molecules in the binding site during the protein target’s preparation (minimization).

If we focus on the non-predicted cases specific to the STDW model with the R310 patch, the observations described above remain valid with very similar trends for all the molecular features (Sup. Note 4, Table S2). In the case of the non-optimal predictions which fail to score the native nucleotide as the best ranked (i.e., good predictions, Fig. 10), similar trends are again observed but with two specificities associated with the metals and stacking contacts (Sup. Note 4, Table S3). First, metals’ presence does impact negatively the performance suggesting that metals contribute directly or indirectly to the nucleotide selectivity. Second, the absence of stacking contacts makes it more challenging to score the native nucleotide properly; the binding selectivity of purines versus pyrimidines, in particular, can be easier to identify in the presence of stacking contacts.

As described above, a low volume of the binding site is detrimental *per se* to the prediction performance. Once the experimental structure is optimized after removal of the ligand (and the water molecules in the SCAL model), the volume can be modulated in a decreasing or increasing way (Sup. Note 4, Fig. S11). The average variation shrinks the binding site by 27 to 30Å^3^ for the SCAL and STDW models, respectively.

In two-thirds of the benchmark, the binding site shrinks by an average of 87 (SCAL) to 92Å^3^ (STDW). In one-third of the benchmark, the binding site expands by an average of 92 (STDW) to 95Å^3^ (SCAL). Thus, a similar trend of variations is observed for both SCAL and STDW models. However, only the STDW is significantly impacted in the performance for the prediction of the Top-10 (Sup. Note 4, Table S4); the shrinking of the binding site combined with the presence of water molecules prevents the identification of any hit in the Top-10 in the concerned cases. This is confirmed by the fact that 9 of the 17 proteins in the subset with no predictions in the Top-10 exhibit recovered predictions in the upper Top-*i* with a smaller patch such as R110 (Sup. Note 4, Table S5). In 6 other cases, the absence of predictions with the STDW model can be imputed to the presence of water molecules (Table S5). Finally, only two cases do not provide any prediction in the Top-*i*.

### Case Studies

The analysis of the molecular features that impact the docking and screening powers shows that different factors are responsible for the general lack of performance of all the nonbonded models and, more specifically, that of the purely implicit model (SCAL). We illustrate the impacting features through a series of case studies, looking particularly at those contributing to the improved performance of the hybrid models, including explicit water molecules. Since all protein-nucleotide complexes in the benchmark include crystallized water molecules, we should expect that the water-mediated contacts will be detrimental to the SCAL model. The presence of water contacts involving the base or the phosphate group has a powerful impact (Fig. S12). We refer to each case using the PDB ID.

In the 1S68 case where the native nucleotide is A, a single water molecule and only one is involved in two close contacts with the nucleotide (Fig. 12). These two water-mediated contacts involve the Watson-Crick face of the adenine. The SCAL model does not provide any prediction within the Top-100 for any nucleotide (Fig. 12A). A few native-like poses exist, but they are not ranked within the Top-100, i.e., their MCSS score is higher than any of the first 100 non-native poses (Fig. 12B,D-E). On the contrary, the STDW model generates several native-like poses within the Top-1, Top-5, and Top-50 corresponding to optimal and good predictions for the native nucleotide (Fig. 12A,C,E-F). Excluding the water-mediated contact with the base, all the other native contacts are found in the native-like poses for both models (Fig. 12D-F). Both water molecules and the nonbonded model used are responsible for the differential scoring between the two models even if the water molecules are not considered in the scoring (Methods B). In all the native-like poses in both SCAL and STDW models, the syn conformation of the nucleotide is preferred (or a high-syn conformation), although the starting conformation in the initial distributions is C3’-endo anti. It is indicative that the ligand’s flexibility allows to switch from anti to syn during the MCSS calculations without any hindrance.

**Fig. 12.**
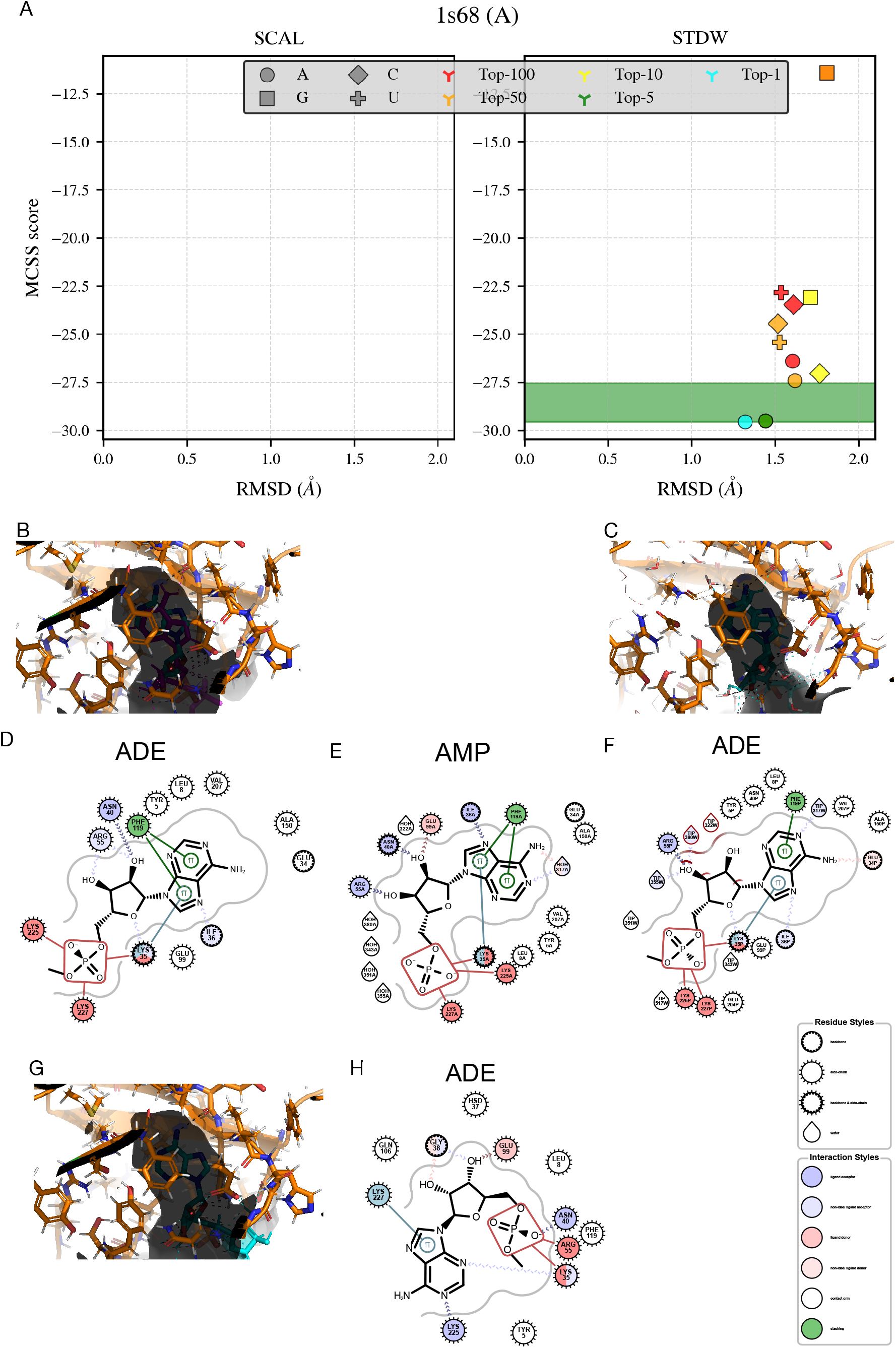
Case Study 1S68. A. Schematic representation of the nucleotide selectivity in the SCAL (left) and STDW (right) models; B. 3D representation of the native-like pose for the SCAL model (128th scored pose); C. 3D representation of the native-like pose for the STDW model (Top-1); D. Diagram of the binding site and nucleotide contacts for the SCAL model (see B); E. Diagram of the binding site and nucleotide contacts for the native binding mode; F. Diagram of the binding site and nucleotide contacts for the STDW model (see C); G. 3D representation of the native-like pose for the SCAL model (Top-1); H. Diagram of the binding site and nucleotide contacts for the SCAL model (see G);

In the SCAL model, the best-ranked poses for the native nucleotide (Top-1 to Top-10) exhibit alternative positionings of the phosphate groups; it interact closely on the opposite side of the binding site with residues Lys35, Asn40, and Arg55 instead of Lys35, Lys225, and Lys227 (Fig. 12G-H). Although the net charge is reduced by two-thirds in the SCAL model (but not in the STDW model), the absence of explicit solvation around the phosphate group leads to an alternate positioning of the nucleotide (Fig. 12H), which is incompatible with the native binding mode. 1S68 is associated with a low volume of the binding site, a feature that impacts the performance negatively (Fig. 11A). Furthermore, the binding site is shrunk for both models and slightly more pronounced for the SCAL model (Sup. Note 4, Fig. S11).

The 3EWY case is specific with a pyrimidic ligand: U, which adopts a syn conformation. Both models exhibit a similar performance with predicting the U native nucleotide (Fig. 13). However, the native contacts are better reproduced by the SCAL model (Fig. 13A-B,E-F) and only the 10th pose reproduces all the native contacts with the residues of the binding site in the STDW model (Fig. 13A,D,F-G). Besides, several non-native nucleotides have very similar MCSS scores: A in the case of the SCAL model, G in the case of the STDW model. There is no global shrinking of the binding site in the protein structures optimized with or without water molecules in the case of 3EWY (Sup. Note 4, Fig. S11). However, there is a local contraction in some parts of the binding site, which is not equivalent between the two models. It is more pronounced on the Hoogsteen and Watson-Crick faces of the base in STDW, making it more challenging to reproduce the native contacts with the base (Fig. 13C). On the other hand, non-native nucleotides can fit into the remodeled binding site with scores which are within the 2 kcal/mol range from the native one (Fig. 14). The native contacts which are specific to the base are lost in the case of A with the SCAL model (Fig. 14A-B,D), but native-like and isosteric contacts are found in the case of G with the STDW model (Fig. 14C,E-F).

**Fig. 13.**
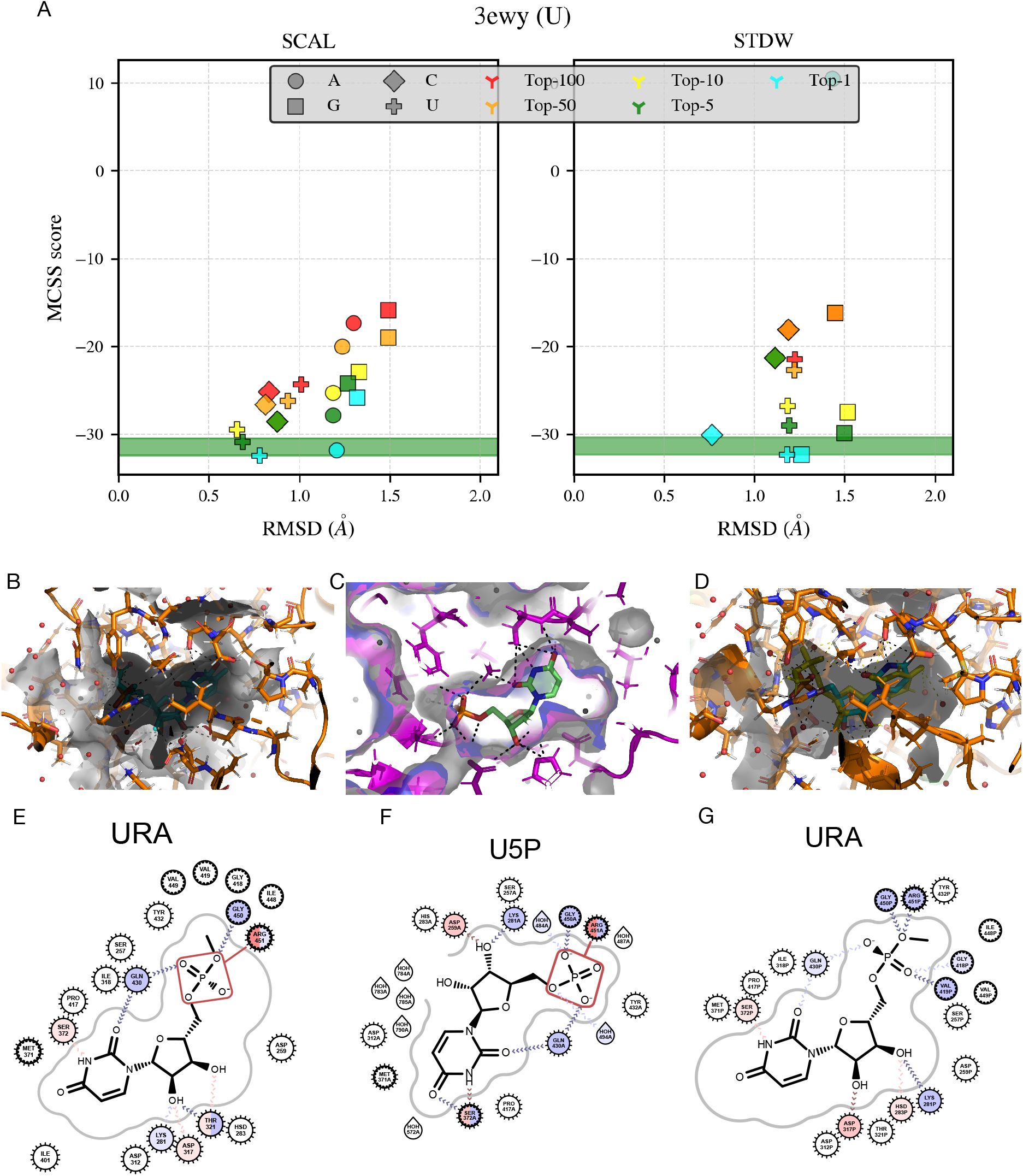
Case Study 3EWY. A. Schematic representation of the nucleotide selectivity in the SCAL (left) and STDW (right) models; B. 3D representation of the Top-1 native pose for the SCAL model; C. 3D representation of the native ligand in the binding site as seen: in the experimental structure (grey), in the optimized structure without water molecules (magenta), in the optimized structure with water molecules (blue); D. 3D representation of the Top-10 native pose for the STDW model; E. Diagram of the binding site and nucleotide contacts for the SCAL model (see B); F. Diagram of the binding site and nucleotide contacts for the native binding mode; G. Diagram of the binding site and nucleotide contacts for the STDW model (see D).

**Fig. 14.**
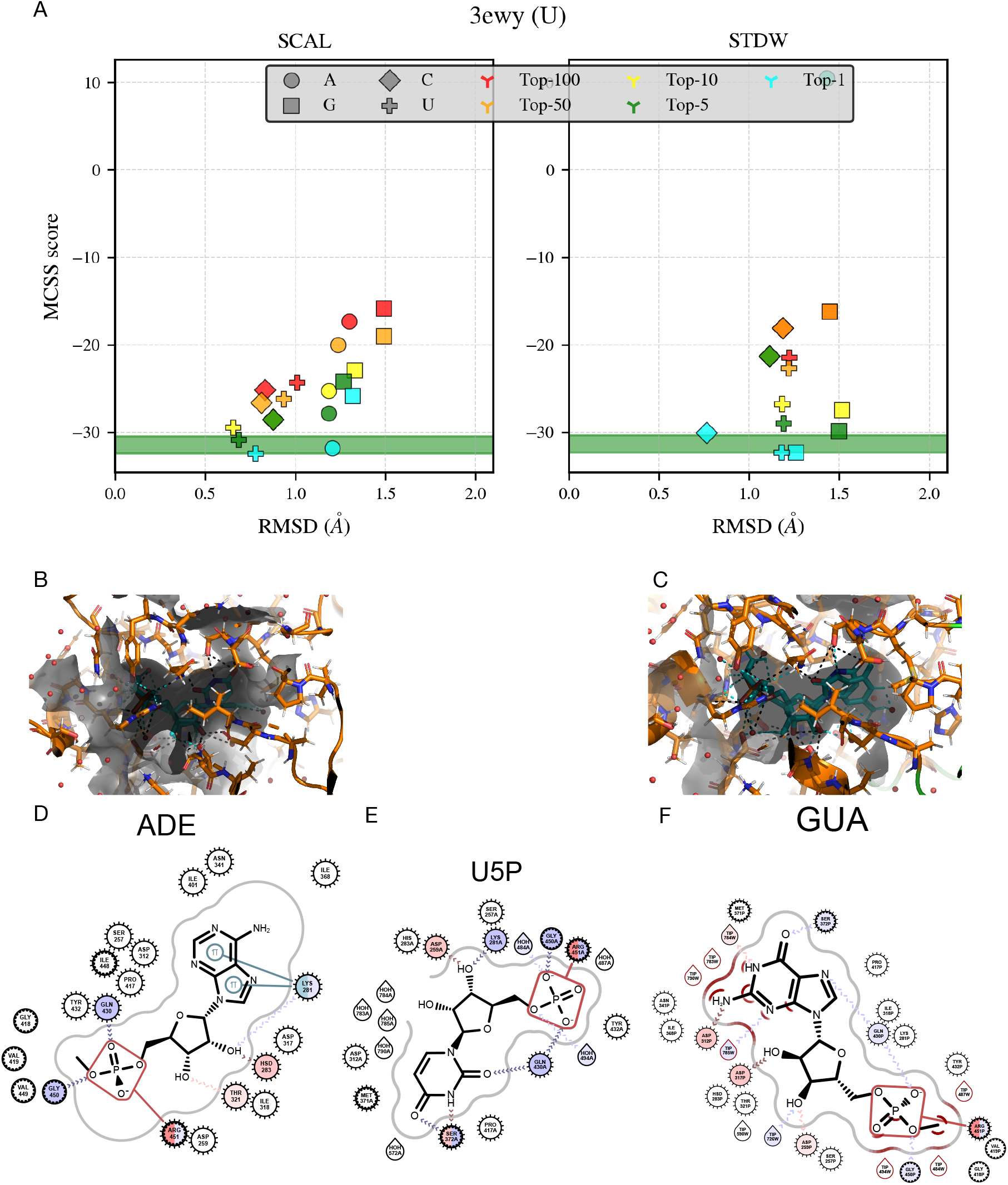
Case Study 3EWY. A. Schematic representation of the nucleotide selectivity in the SCAL (left) and STDW (right) models; B. 3D representation of the Top-1 native-like pose for A (syn conformation) in the SCAL model; C. 3D representation of the Top-1 native-like pose for G (anti conformation) in the STDW model; D. Diagram of the binding site and nucleotide contacts for the SCAL model (see B); F. Diagram of the binding site and nucleotide contacts for the native binding mode; G. Diagram of the binding site and nucleotide contacts for the STDW model (see C).

In the 2XBU case, the cavity of the binding site is well conserved in the SCAL model but slightly shrunk in the STDW model (Sup. Note 4 and Fig. S11). The binding site’s volume is low because it is quite open with only the base moiety within a well-defined cavity. Only the STDW model provides a good prediction (Top-5) while the native poses generated by the SCAL model are all over the Top-100 scores (Fig. 15A). The first poses in the STDW model (Top-1 to Top-4) are all located in the binding site. However, their RMSD is over 2Å and are thus excluded from the native poses. Independent of the scores, the native poses reproduce the native contacts with the base in both models (Fig. 15B-G). The phosphate group establishes very close contacts with hydrogen-bond donors from the peptide backbone, but those contacts are not retrieved in the native poses except for one residue (Thr115). The presence of a terminal methyl group in the phosphate patch used: R310 (Fig. 4 and Fig. 3) prevents a native positioning of the phosphate group. In the SCAL model, its positioning is more in agreement with the experimental structure (Fig. 15E-F). However, the Top-1 pose and the other best-scored poses are completely off-site (Fig. 16).

**Fig. 15.**
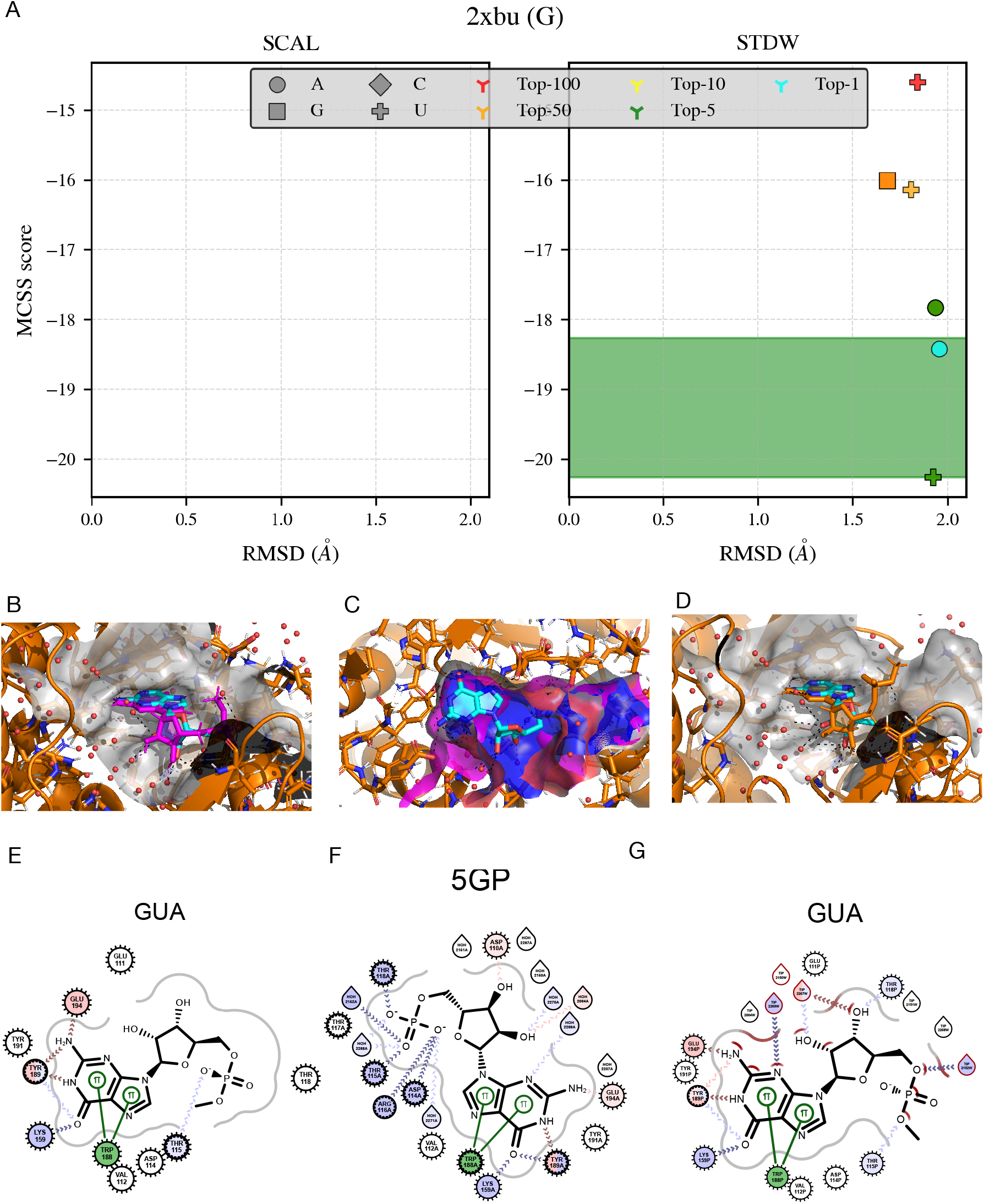
Case Study 2XBU. A. Schematic representation of the nucleotide selectivity in the SCAL (left) and STDW (right) models; B. 3D representation of a native-like pose for G in the SCAL model (169th scored pose); C. 3D representation of the native ligand in the binding site as seen: in the experimental structure (grey), in the optimized structure without water molecules (magenta), in the optimized structure with water molecules (blue); D. 3D representation of a native-like pose for G in the STDW model (Top-50, 12th scored pose); E. Diagram of the binding site and nucleotide contacts for the SCAL model (see B); F. Diagram of the binding site and nucleotide contacts for the native binding mode; G. Diagram of the binding site and nucleotide contacts for the STDW model (see D).

**Fig. 16.**
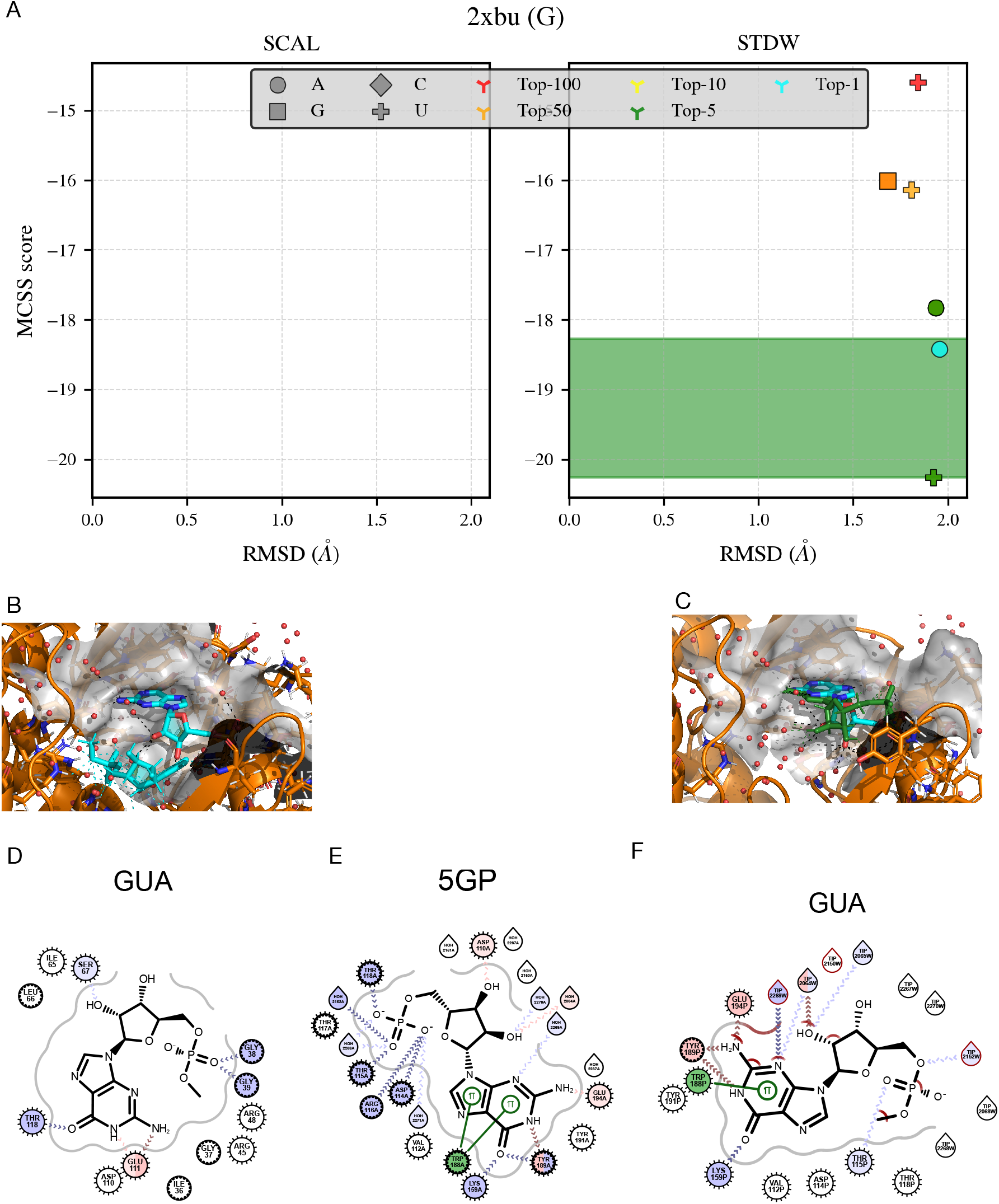
Case Study 2XBU. A. Schematic representation of the nucleotide selectivity in the SCAL (left) and STDW (right) models; B. 3D representation of Top-1 pose for G in the SCAL model; C. 3D representation of Top-4 pose for G in the STDW model (RMSD > 2.0Å); D. Diagram of the binding site and nucleotide contacts for the SCAL model (see B); E. Diagram of the binding site and nucleotide contacts for the native binding mode; F. Diagram of the binding site and nucleotide contacts for the STDW model (see C).

Combined with the shrinking of the binding site, the large phosphate group largely deviates from the expected position in the STDW model, and the only contact with Thr115 is weaker. This deviation on the phosphate group increases the global RMSD of the pose to the native coordinates and leads to exclude the Top-1 to Top-4 G poses from the list of native poses (Fig. 16). However, the Top-4 G pose reproduces almost all the native contacts (Fig. 16C,E-F).

## Conclusions

MCSS was evaluated for the docking of nucleotides on a benchmark of 121 protein complexes. Different solvent and phosphate models were tested to optimize the success rate for identifying native poses (docking power) and the true native nucleotide (screening power). As a result, the STDW model, which is a hybrid implicit and explicit solvent model, appears to give the best performance slightly ahead of the SCALW model based on partially reduced charges on the phosphate group. A clustering procedure was set up that allows a slight increase of the success rates, especially in the high Top-*i* (Top-50 and Top-100). Among the different phosphate models, the more voluminous one that carries a terminal methyl group: R310, is slightly better in the Top-1 predictions. It is also the phosphate model that facilitates the linking of nucleotide fragments in the perspective of fragment-based design of oligonucleotides (unpublished data). The combined STDW-R310 model outperforms despite the few cases where the lack of predictions in the Top-10 could be correlated to a size effect that prevents the phosphate group from fitting correctly in the binding site. The presence of water molecules in the preparation and optimization of the protein structure allows the minimized structure to deviate less from the experimental structure. On the other hand, the water molecules generally induce a more pronounced shrinking of the binding site with respect to the experimental structure, which is responsible for some degradation of the performance. The inclusion of water molecules gives a more realistic description of the binding site, whether they are involved in water-mediated contacts with the ligand or just solvating the phosphate group or ribose.

We have identified some pitfalls that contribute to degrade the performance of prediction in all models. From the intrinsic features of the binding site, a low binding volume is the more impacting factor. It can be seen as a low accessible volume for close contacts that typically occurs when the binding site is open with few contacts with the ligand or when the close contacts are just present in some part of the nucleotide (small binding cavities). Among the conformational features, the syn conformation does not have any negative impact although the docking is performed using an initial C3’-endo anti conformation. This confirms that the flexibility of the ligand during the docking allows a proper conformational sampling. Among the interaction features, the presence of salt bridges makes it a bit easier to get good predictions. On the other hand, the presence of clashes with water molecules in the experimental structure has a slight negative impact. More specific to the STDW-R310 model, the negative effect of the low binding volume is smoothed.

The quality of the scoring explains, to some extent, the better performance of the hybrid model STDW over the implicit model SCAL. First of all, the SCAL includes a slight bias in the scoring, favoring G over A and the other nucleotides, while A is the nucleotide that generally establishes the stronger contacts in the binding site. Furthermore, the STDW model based on the original parameters from the CHARMM forcefield describes better, in the presence of water molecules, the bonded contributions associated with the nucleotides’ torsion angles. Thus, the penalty term of the MCSS score from the conformational distortions of the bound ligand is more accurate. The STDW model outperforms not only in docking power with more predictions but also in screening power even if we just consider the common predictions for both models. The STDW model has a much stronger discriminatory power between very similar ligands. It is also consistent with the broader range of score distributions for each type of nucleotide. The native poses scored as optimal reproduce most (if not all) of the native contacts as well as the good predictions, although they are not ranked first among the four nucleotides.

Both free and bound conformations for the same protein are not available on a large set of 3D structures with nucleotide ligands. Thus, the protein targets correspond to some unbound forms where the ligands were extracted from the binding site. Consequently, the optimized binding site is usually shrunk, making the identification of native poses and native binders more challenging. The method’s performance is then degraded both in terms of docking and screening powers because of missing native contacts in the shrunk areas of the binding sites. The four standard nucleotides are very similar from the chemical viewpoint and thus harder to discriminate in terms of binding selectivity. Chemical modifications would increase the dissimilarity between nucleotidic fragments, which would likely be easier to discriminate. Many modified nucleotides are already used experimentally in the synthesis of oligonucleotides (66) or modified aptamers (67) for medical or biotechnological applications to improve specific properties such as the therapeutic index (68).

From the perspective of designing oligonucleotides, MCSS provides a reasonable performance to predict native poses and identify the binding preference(s) of nucleotidic fragments. More accurate or complete descriptions of the solvent open the possibility to improve its performance. Other improvements may come from using more relevant and more diverse conformations of the protein targets and increasing the chemical diversity of the nucleotidic fragments.

## ACKNOWLEDGEMENTS

NC was supported by the French Ministry of Higher Education, Research and Innovation. RGA is supported by the Excellence Eiffel Ph.D. Program. We thank the French Ministry of Foreign Affairs for financial support (PHC Carlos J. Finlay, 41814TM).

## Methods

### A. Protein-nucleotide Benchmark

The PDB was filtered out to select a set of protein-nucleotide complexes based on different structural criteria associated with the atomic resolution and the structural similarity. A first query was carried out to find protein complexes with each of the four nucleotides as ligands and annotated in the PDB by the following labels: AMP, C5P, 5GP, U5P. An additional criterion based on a cut-off value of 2Å resolution was also used to select only high-resolution X-ray structures. The resulting complexes were then clustered according to their sequence similarities in order to remove the redundancy. If any chain in the protein of a complex has at least 30% sequence identity with a chain in the protein from another complex, the two complexes were grouped into the same cluster. The crystal structure with the best resolution in each cluster was selected as the cluster’s representative. The 188 complexes thus selected by pulling down the results from the four queries (AMP-bound: 123, C5P-bound: 18, 5GP-bound: 21, U5P-bound: 27) were then manually curated to retain those that exhibit a known binding preference for the crystallized ligand. This feature was established based on the literature and/or the annotation of the protein, e.g., a C nucleotide for CMP-kinase, etc. After curation, the dataset was reduced to 132 complexes. An additional curation was performed to eliminate some potential redundancy associated with the presence of identical binding sites for different types of nucleotides. The followed procedure consists of superimposing all the protein structures using the program TM-align (69) and review all the structures that are similar based on the TM-score (TM-score ≥ 0,8). Two binding sites were considered non-redundant if they differ by only one amino acid residue in direct contact with the ligand. According to this criterion, only one complex was removed from the dataset in the case of the proteins corresponding to the PDB IDs: 3DXG (U5P ligand) and 3DJX (C5P ligand); the latter complex was conserved in the dataset to compensate for the minor under-representation of C5P. The full procedure ends up with a dataset of 131 protein-nucleotide complexes.

After a review of the MCSS calculations, ten protein-nucleotide complexes resulted in non-productive (see below) and were then removed from further analyses. The resulting benchmark is thus composed of 121 protein-nucleotide complexes (Sup. Note 1). Their binding features were characterized by the number of contacts between the protein and its ligand, the fraction of buried surface area, the number of H-bonds in the binding site, and the energy of interaction as calculated by the MCSS scoring function (Sup. Note 2). The contacts are calculated using the program BINANA (55). The full tables, including the molecular features of the protein-nucleotide complexes, are provided in the supplementary materials (Sup. Note 4).

### B. MCSS

All the proteins are prepared using the CHARMM-GUI interface (70) to convert the PDB files into CRD and PSF formats. After removal of all heteroatoms, hydrogens are added to the protein using the HBUILD command from CHARMM. Histidine residues are considered as neutral. The protein targets are then submitted to an energy minimization (tolerance gradient of 0.1 kcal/mol/Å^2^). The average deviation between the experimental structure and the minimized structure is around 1.0Å for the structures optimized without water molecules and 0.5Å for the structures optimized with the crystallized water molecules (Fig. 2).

The nucleotide library of fragments include multiple conformations, 5’ and 3’ patches (see MCSS documentation: https://www.mcss.cnrs.fr/MCSSDOC/ Welcome.html. The initial default conformation used in the calculations is a C3’-endo/anti ribonucleotide. A set of five different patches on the 5’ end is used in the current study with this nucleotide conformation: R010, R110, R210, R310, R410. Each binding region is defined by a 17Å^3^ cubic box centered on the ligand centroid (Fig. 4). MCSS sample files are provided for the input and nonbonded parameters (Sup. Note 2).

Ten protein-nucleotide complexes (PDB IDs: 1HXP, 2CFM, 2Q4H, 3L9W, 3REX 4OKE, 4XBA, 5ERS, 5M45, and 5DJH) resulted as non-productive because of a significant conformational change of the binding site after minimization (see protocol for energy minimization above) that prevented the identification of native-like poses. They are excluded from post-docking analyses due to 3 main reasons: (1) no native pose (RMSD ≤ 2.0Å) could be generated because of a nucleotide-binding site too buried to be accessible after minimization (PDB ID: 5M45, 5DJH); (2) no native pose could be identified consistent with a huge deviation (RMSD > 2.0Å) of the crystallized ligand minimized within the optimized protein binding site (PDB IDs: 1HXP, 2CFM, 4OKE, 4XBA, 5ERS); (3) the native poses identified showed highly unfavorable energies indicating the presence of steric clashes between the nucleotide and the minimized binding site (PDB IDs: 2CFM, 2Q4H, 3L9W, 3REX, 4XBA, 5ERS). After the removal of those ten non-productive protein-nucleotide complexes, the resulting benchmark includes 121 protein structures. The reference coordinates of the ligand used to evaluate the poses correspond to those of the experimental X-ray structure.

The initial distributions of fragments are generated using 2000 groups distributed randomly and repeatedly among 25 iterations. These parameters guarantee that fragments fully saturate the binding region of all the protein-nucleotide complexes in the benchmark, i.e., the atomic density of the fragments mapped into the box is at least twice that of the maximum carbon density.

In the models that include explicit solvent (SCALW, STDW, and FULLW), the water molecules are treated independently from the fragments, which are replicated from their initial distribution during each iteration. The number of water molecules is conserved during the calculations, and they are free to move around without any constraint. However, they are not considered in the scoring as described below.

The MCSS score is defined by the electrostatic and van der Waals contributions to the interaction energy plus a penalty term corresponding to the deviation of the fragment’s conformation from its energy minimum:

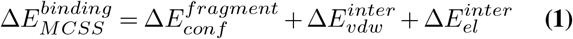

The van der Waals contribution to the score is calculated in the same way for all models:

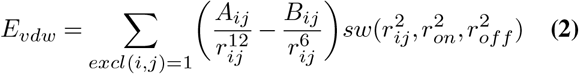

while the electrostatic contribution depends on the solvent model used. In the case of the “FULL” model, it is calculated using the standard charges as follows:

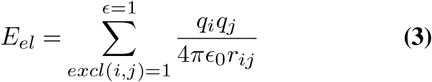

In the case of the other models using either scaled charges (i.e. “SCAL”) or standard charges (i.e. “STD”) (Fig. 3), it is calculated this way:

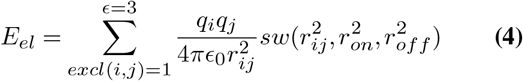

where the dielectric constant is set up according to some previous work (43).

The MCSS software may be obtained after signing a license agreement upon request to Martin Karplus. The source code can be obtained from a Git repository on the I2BC software forge https://forge.i2bc.paris-saclay.fr).

### C. Clustering

A fast and straightforward clustering procedure (orthogonal clustering from now on) is performed on the MCSS distributions; the first pose (best ranked) is taken as the seed of the first cluster, and all other poses in the exploration with an RMSD less equal than 1Å to the seed (redundant poses) are removed from the dataset. The seed is preserved, and the process resumes taking as seed the next available pose and performing the same comparison against remaining poses. At the end, a set of geometrically non-redundant seeds is obtained. The MCSS results presented include the distributions’ analysis: the raw (R) and clustered (C) distributions.

### D. Screening power

To evaluate the screening power, the MCSS distributions from the four nucleotides are merged and sorted according to their score in increasing order as in the nucleotide-specific distributions (from the more negative to the less negative or positive). In each Top-*i*, a prediction is considered as optimal if both conditions are met: (1) it corresponds to a native pose (RMSD ≤ 2.0Å), (2) the native nucleotide is ranked ahead of the three other non-native nucleotides. For example, an optimal prediction in the Top-1 means a native pose is found with the best score from the merged distributions. In the cases where a non-native is ranked ahead of the native nucleotide in the native-like poses, a prediction is considered as good if the score difference does not exceed 2 kcal/mol. On the contrary, the prediction is considered as poor.

### E. Molecular Features

The volume calculation of the binding site is performed using the PyVOL python package (65). PyVOL is used with the pocket corresponding to the nucleotide-binding site as input (coordinates of the nucleotide ligand of interest). The threshold value to discriminate between high or low binding volume is set to 635Å^3^.

The other molecular features include the number of water molecules around the nucleotidic ligand, the presence of metals, and the presence of other nucleotidic ligands (nucleic acid or cofactor) in close vicinity to the binding site. The threshold value for the number of water molecules between nwat high and low is set to: 6 (nwat.low ≤ 6 & nwat.high > 6).

The interaction features (base contacts, clashes, salt bridge, stacking) are extracted from the analysis of the binding site (54) (Sup. Note 1).

## Supplementary Note 1: Benchmark of 121 protein-nucleotide complexes

Attached Supplementary Data 1 (Data-S1.csv): a list of PDB IDs including the ligand ID, the atomic resolution, functional classification, and EC number.

**Fig. S1.**
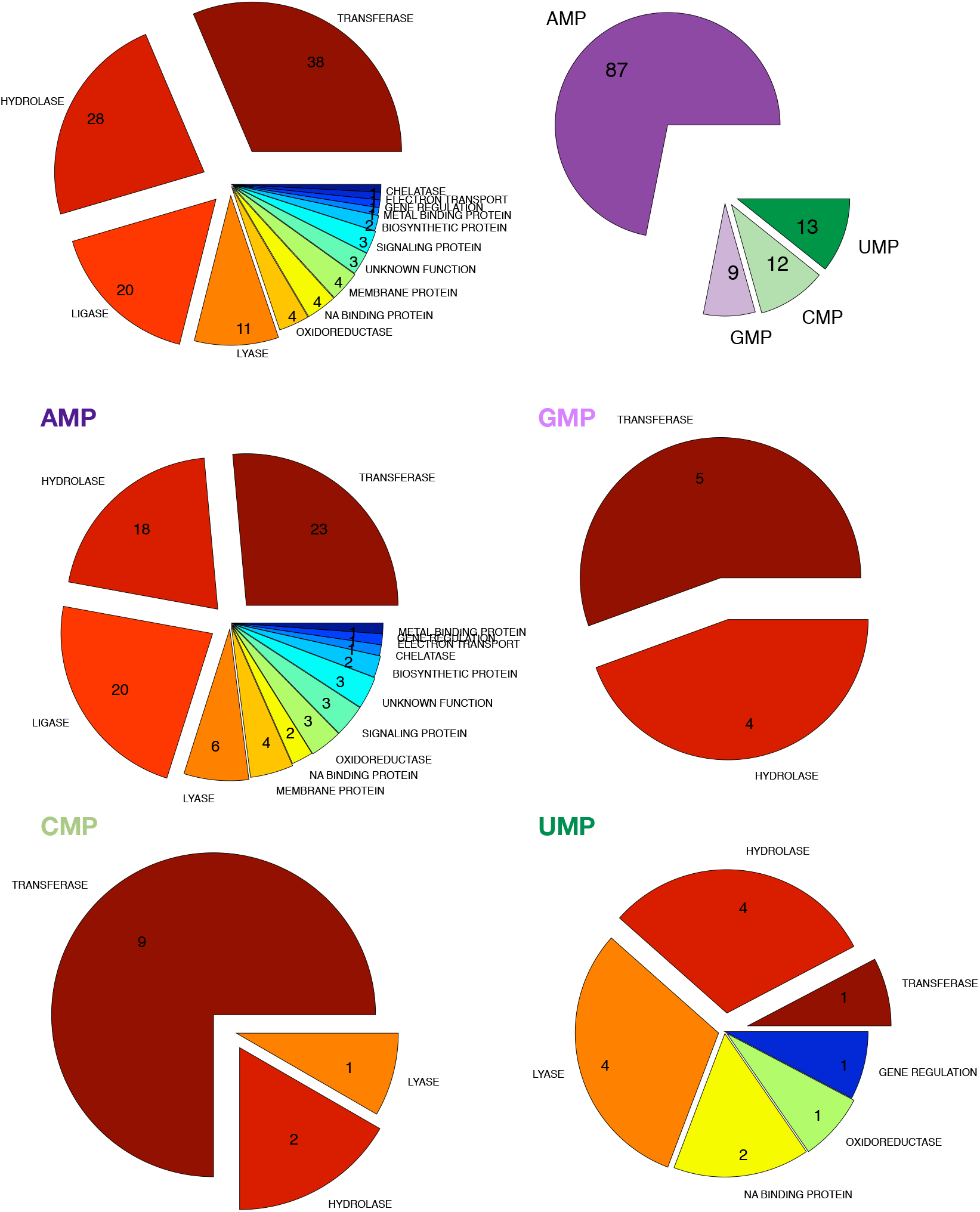
Distribution of molecular functions and nucleotide types in the protein-nucleotide benchmark. Top: General distribution of molecular functions (left) and nucleotide type (right). Middle-Bottom: Nucleotide-specifc distributions (AMP, GMP, CMP, UMP).

Attached Supplementary Data 2 (Data-S2.csv): calculations of the BINANA features (number of contacts, number of H-bonds, the buried fraction of ligand, etc)

Attached Supplementary Data 3 (Data-S3.csv): calculations of the NACCESS surface terms for the fraction of buried surface of the ligand

Attached Supplementary Data 4 (Data-S4.tar.gz): 2D diagrams of the contacts within the binding sites (SVG format).

**Fig. S2.**
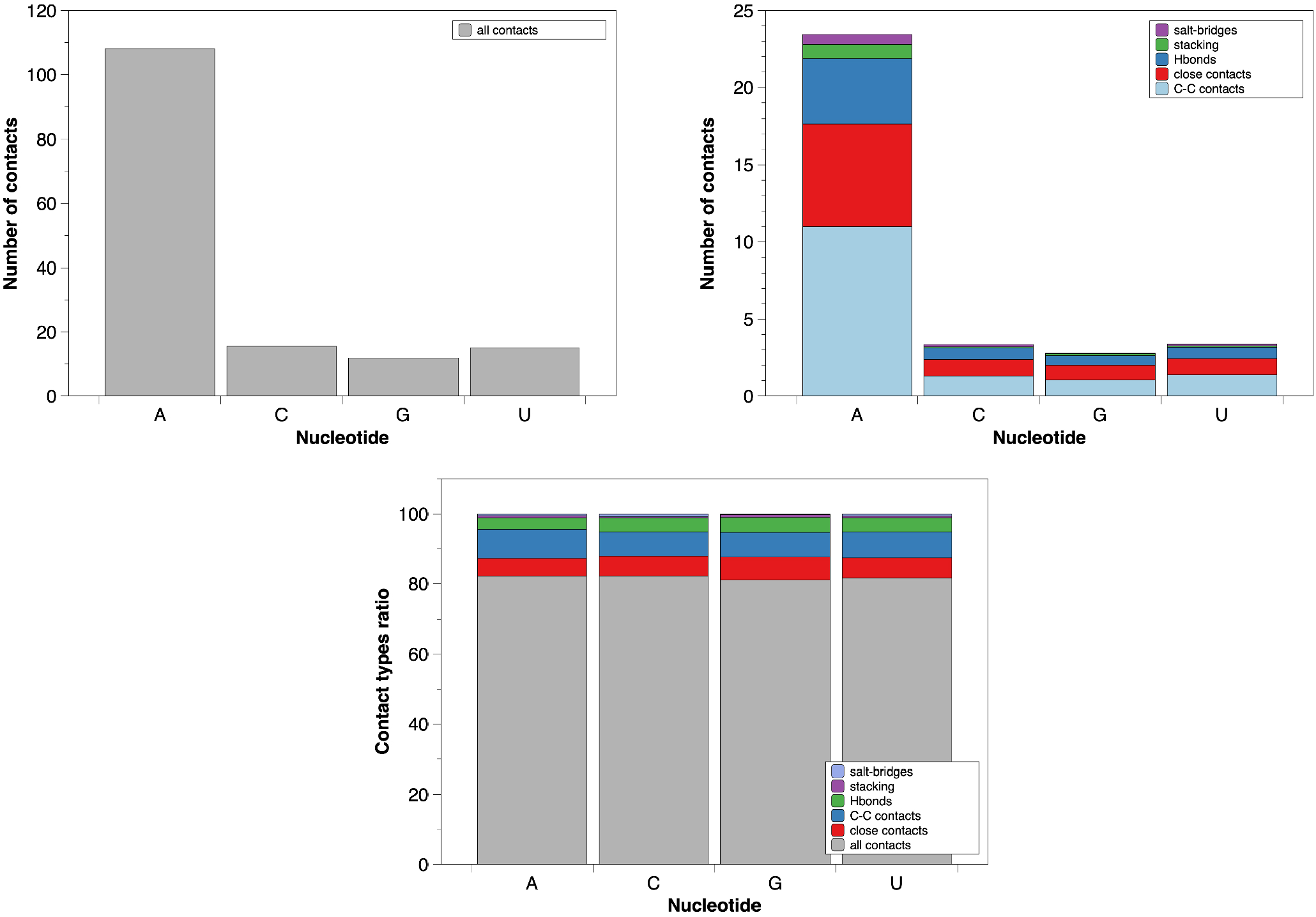
Nucleotide breakdown of atomic contacts. Top-left: all contacts; top-right: specific contacts (C-C contacts, close contacts, Hbonds, stacking contacts, salt-bridges); bottom: ratio of each type of specific contacts. The number of contacts correspond to the average value over the full benchmark.

## Supplementary Note 2: MCSS

Attached Supplementary Data 5: MCSS input sample (Data-S5.txt)

Attached Supplementary Data 6: MCSS nonbonded parameters sample (Data-S6.txt)

Attached Supplementary Data 7 (Data-S7.csv): MCSS score (including its VdW and elec terms) and RMSD values for each protein-nucleotide complex

**Fig. S3.**
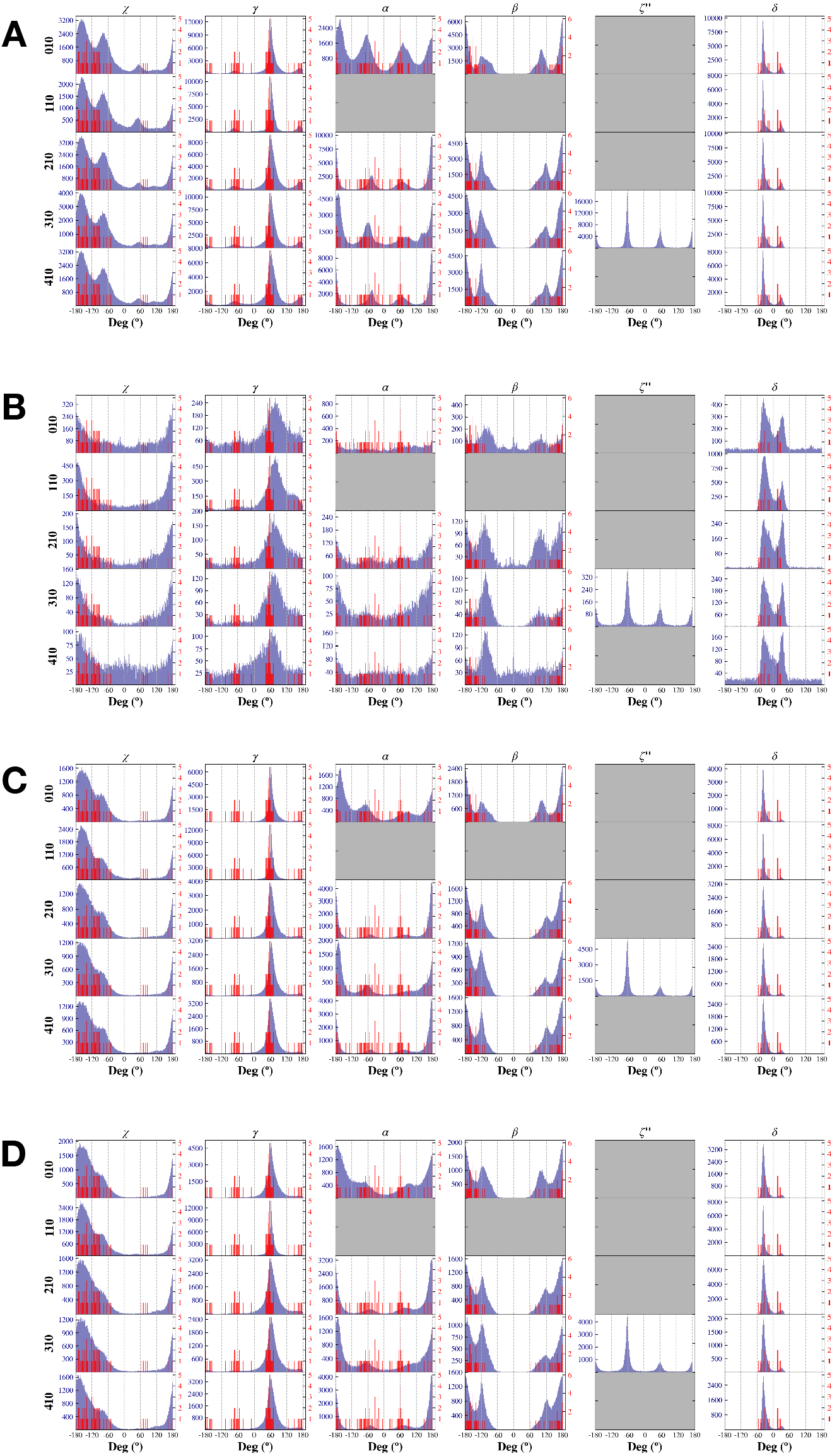
Torsions angles. Nonbonded models and associated patches (R010 to R410): A. SCAL, B. FULLW, C. SCALW, D. STDW. In blue: the distribution of the torsions angles observed in the MCSS minima; In red: the distribution of the torsions angles observed in the bound ligands.

## Supplementary Note 3: Scoring

Autodock Vina is a well-known docking method used for virtual screening; the associated scoring function is pretty robust, having regularly been used in the comparative assessment of scoring functions (CASF) challenges (71). Vinardo and Δ_*vina*_*RF*_20_ were both derived from Vina and tested in the CASF-2013 challenge. Vinardo was optimized and validated on large datasets (61). It was tested in particular on the DUD library that contains, among other proteins, kinases with nucleotide ligands or nucleotide analogs (72). Δ_*vina*_*RF*_20_ was derived more recently from Vina with a new parametrization based on random forest. The performance of Δ_*vina*_*RF*_20_ was superior to that of Vina when tested on the CASF-2007 and CASF-2013 challenges benchmarks. Finally, ITscorePR was included since it has been specifically developed for protein-RNA interactions. The scores calculated with all the scoring functions: ITscorePR (58), Δ_*vina*_*RF*_20_ (59), Autodock Vina score (60), and Vinardo (61), except MCSS (43) correspond to single-point calculations on the MCSS-generated poses.

Attached Supplementary Data 8 (Data-S8.tar.gz): selectivity diagrams SCAL/STDW for the native poses for each protein-nucleotide complex of the benchmark.

**Fig. S4.**
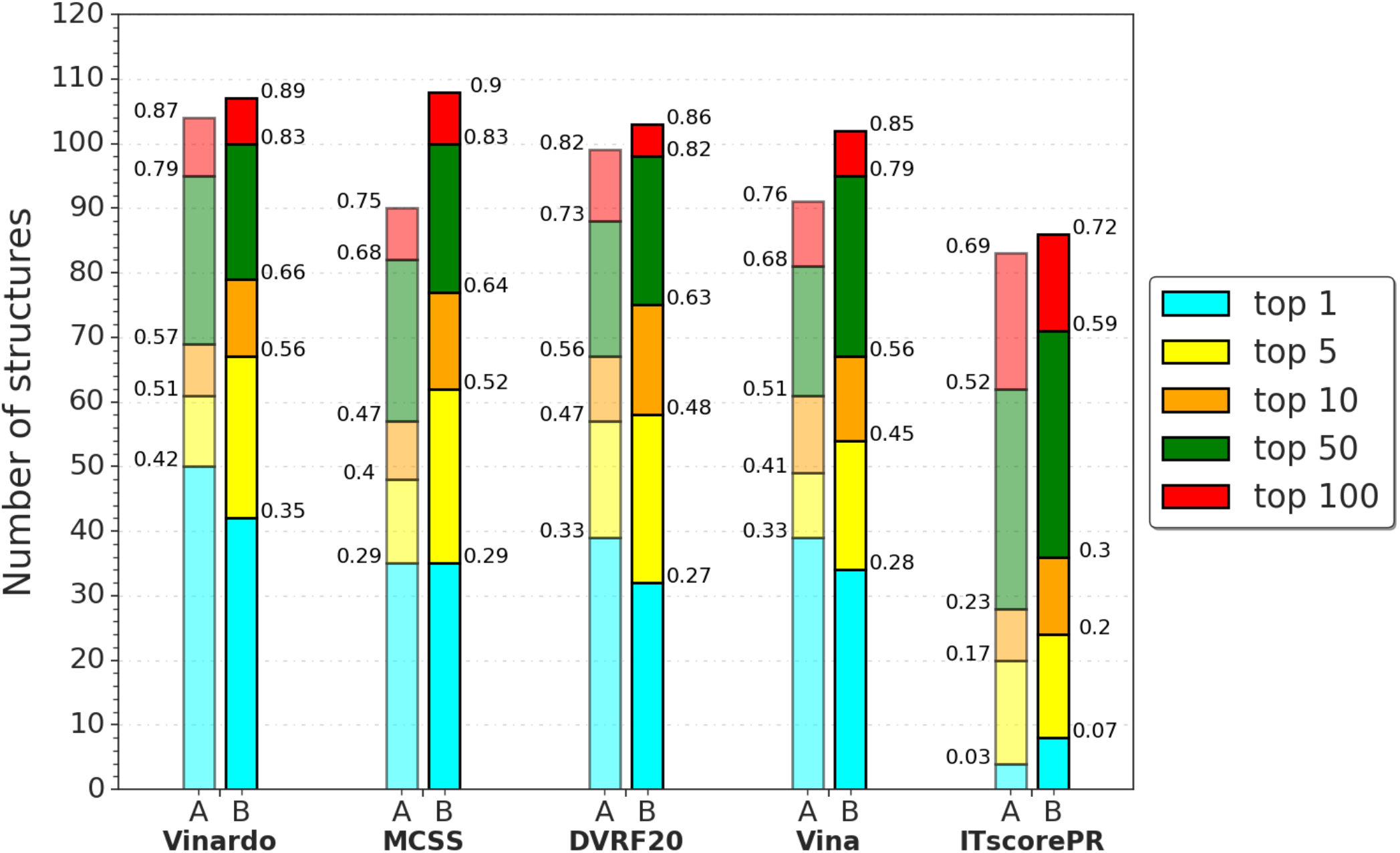
Stacked histogram representation of the native poses in the top1 to top100 as scored by Vinardo, MCSS, Δ_*vina*_*RF*_20_, Vina, and ITscorePR. A. no clustering; B. clustering.

**Fig. S5.**
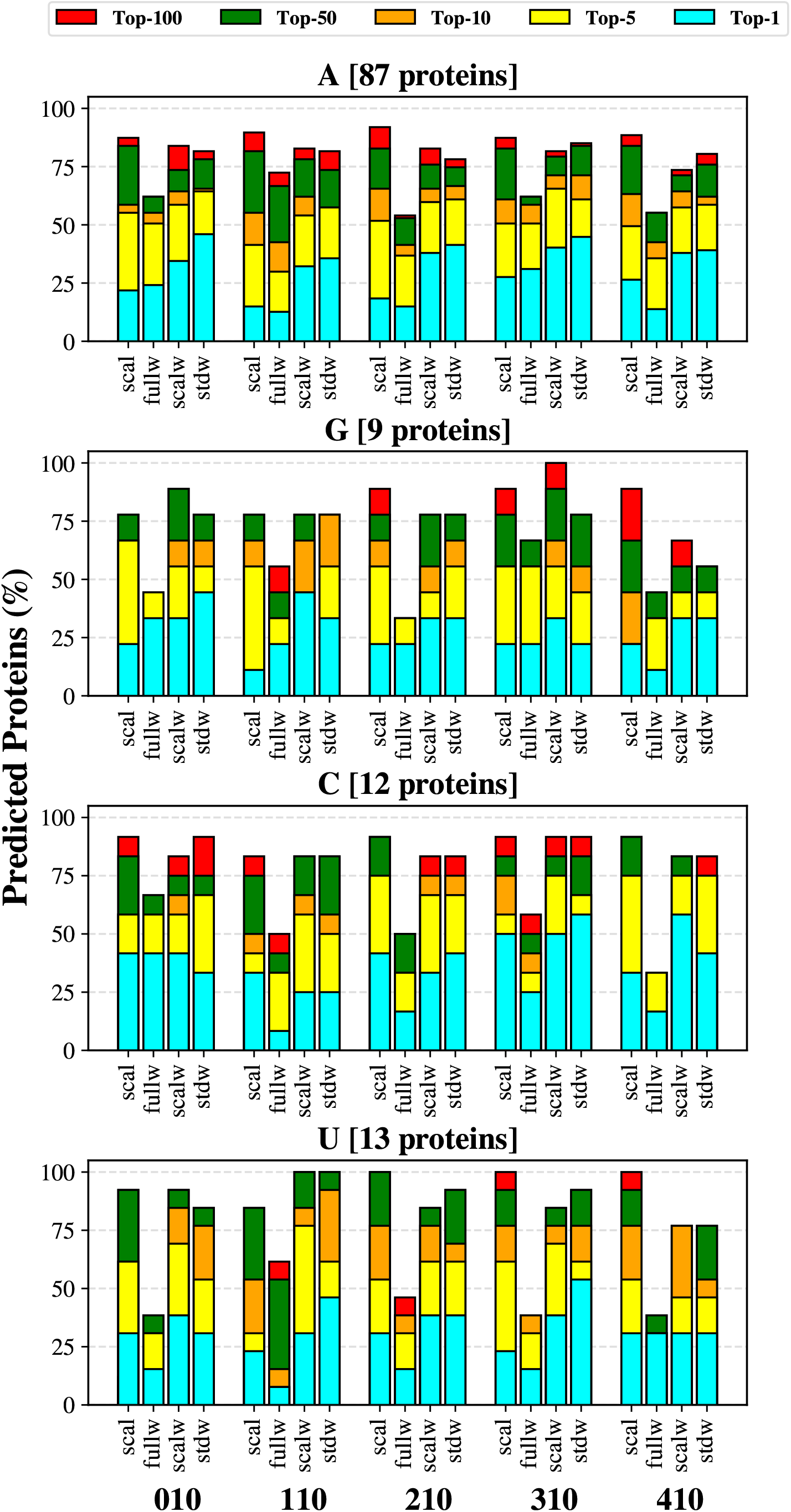
Nucleotide breakdown of the success rates obtained for each nonbonded model and patches combination. The data are shown for the clustered distribution and each Top-*i*.

**Fig. S6.**
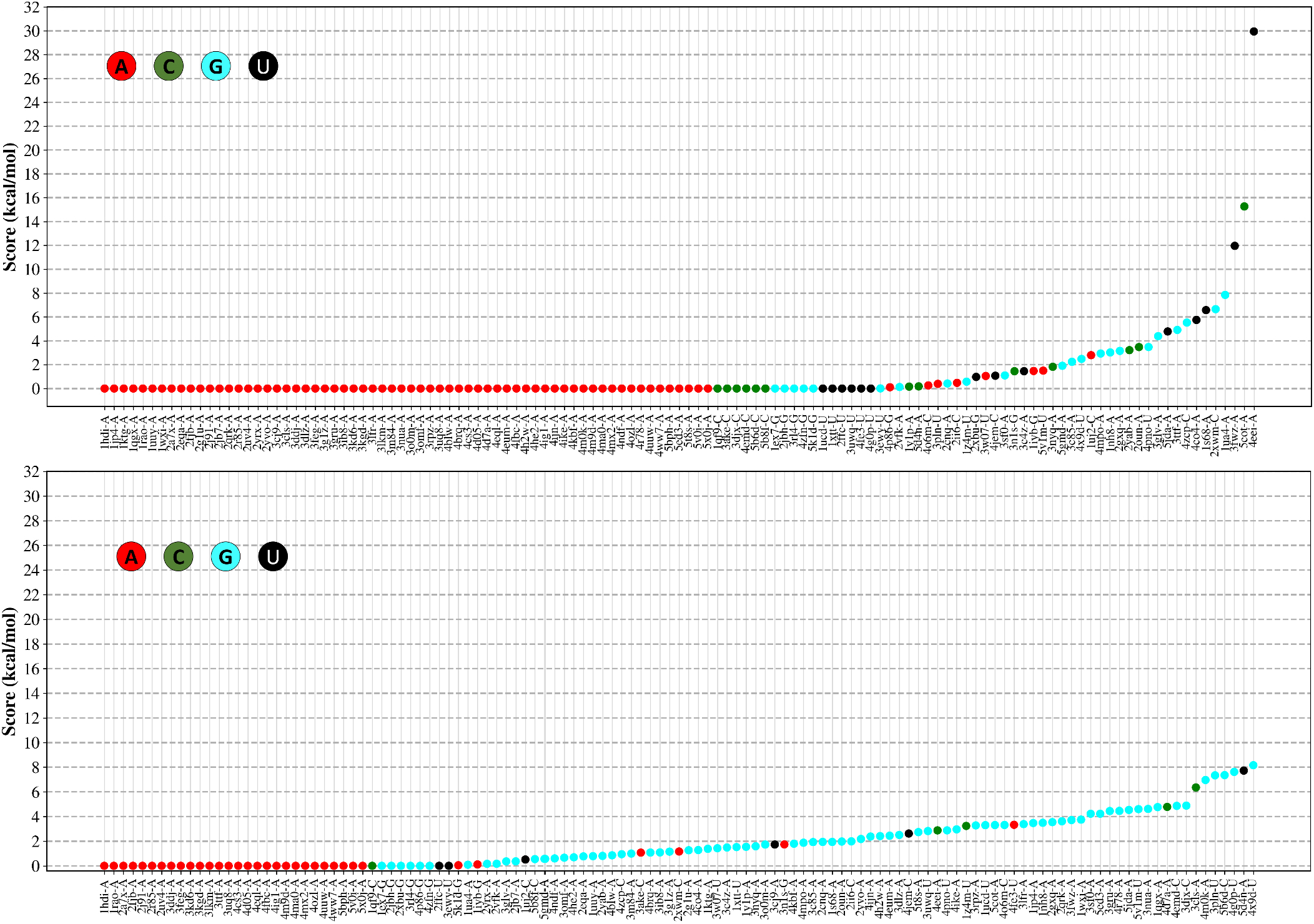
Scoring differences (offset) between the best-ranked pose whatever the nucleotide type and the best-ranked pose for the nucleotide corresponding to the native ligand. Top: STDW model; bottom: SCAL model. The color code indicates the nucleotide type.

**Fig. S7.**
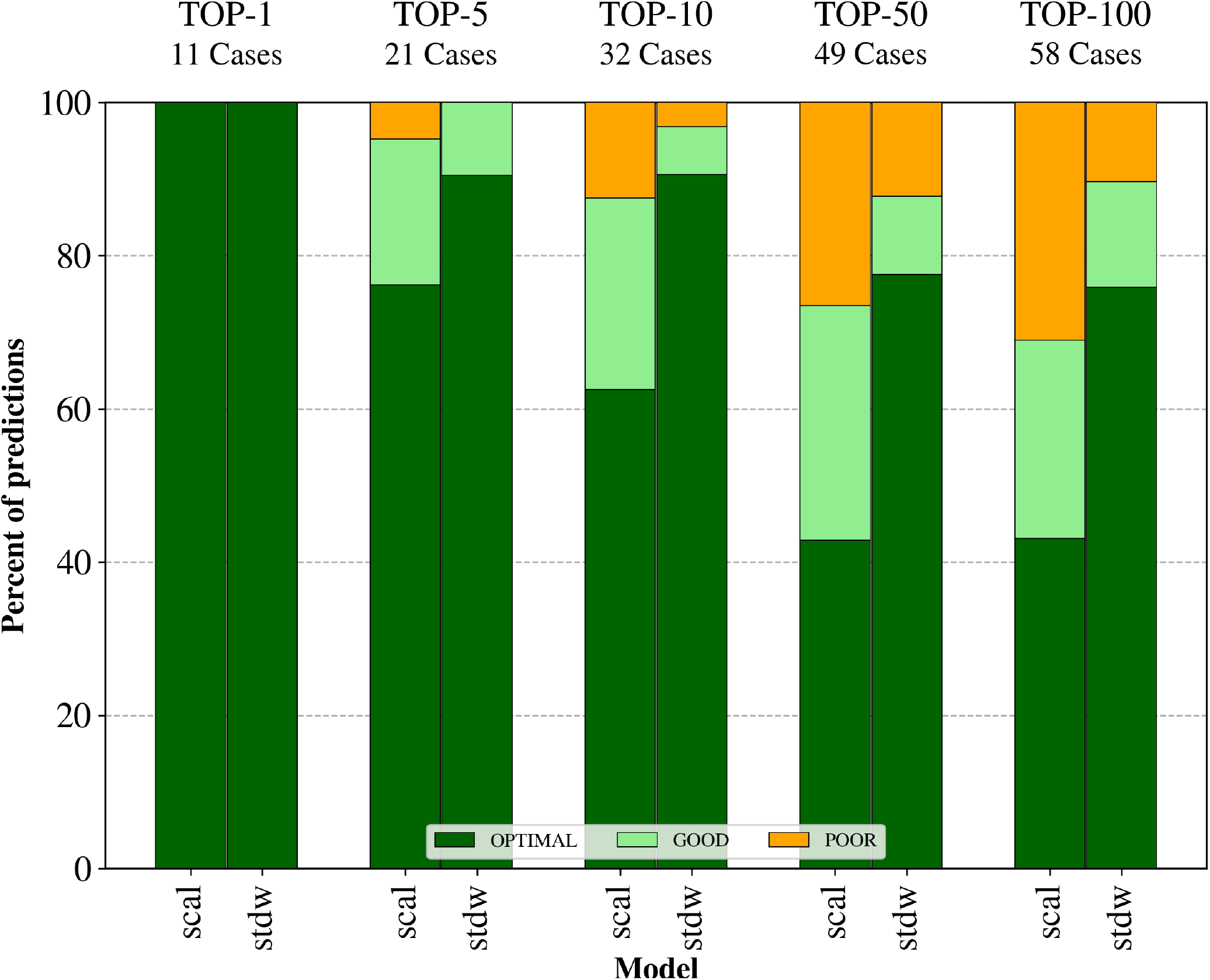
Binding Selectivity Predictions. Optimal: native nucleotide as the best ranked; good: native nucleotide in the ranked within a 2 kcal/mol range from the best ranked non-native nucleotide; poor: native nucleotide ranked out of the 2 kcal/mol range. Only a subset of the benchmark is considered where both SCAL and STDW models do provide predictions.

**Fig. S8.**
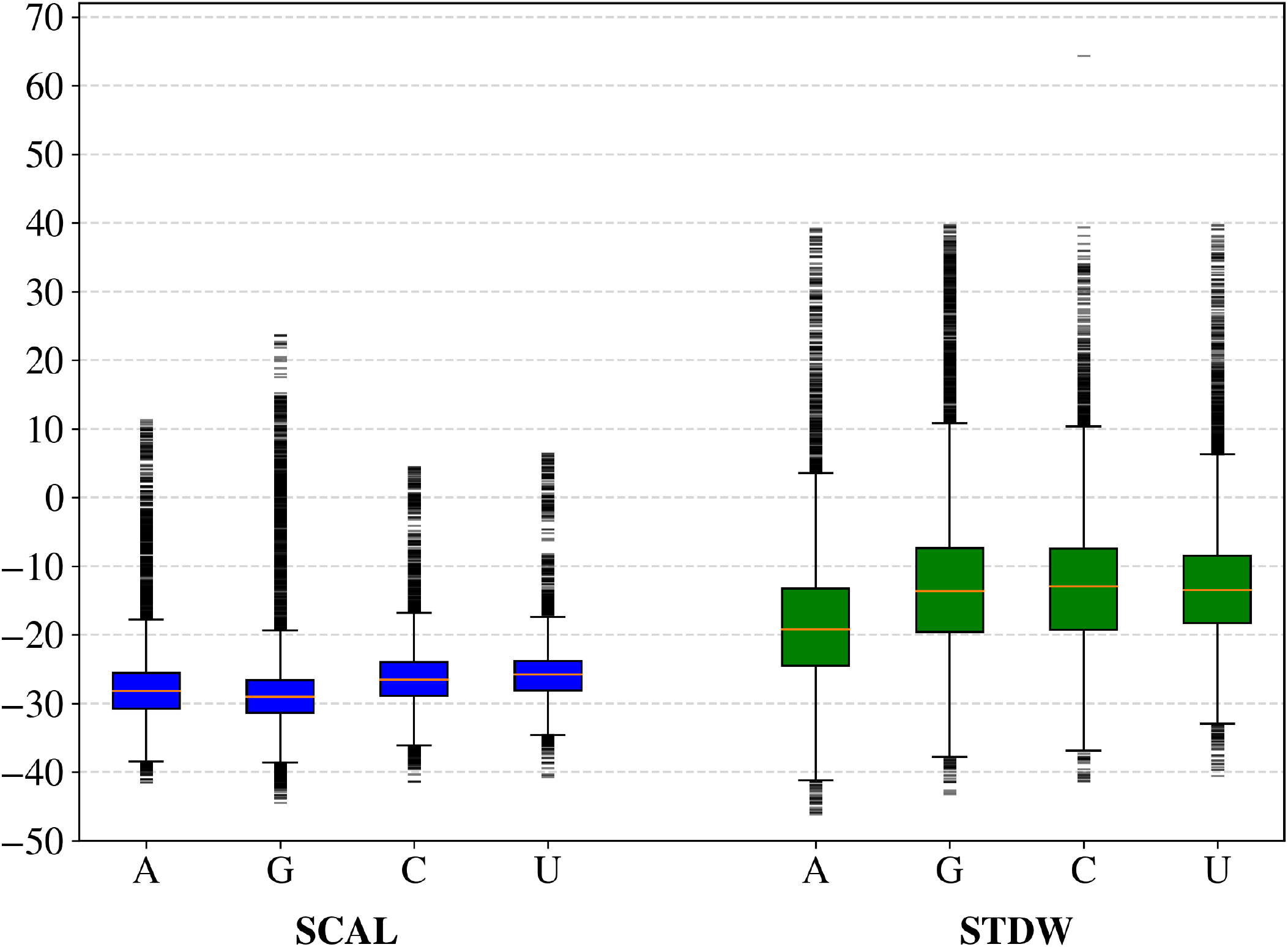
Boxplots for the nucleotide breakdown of the MCSS score. The distributions correspond to the full benchmark with the SCAL or STDW model (R310).

**Fig. S9.**
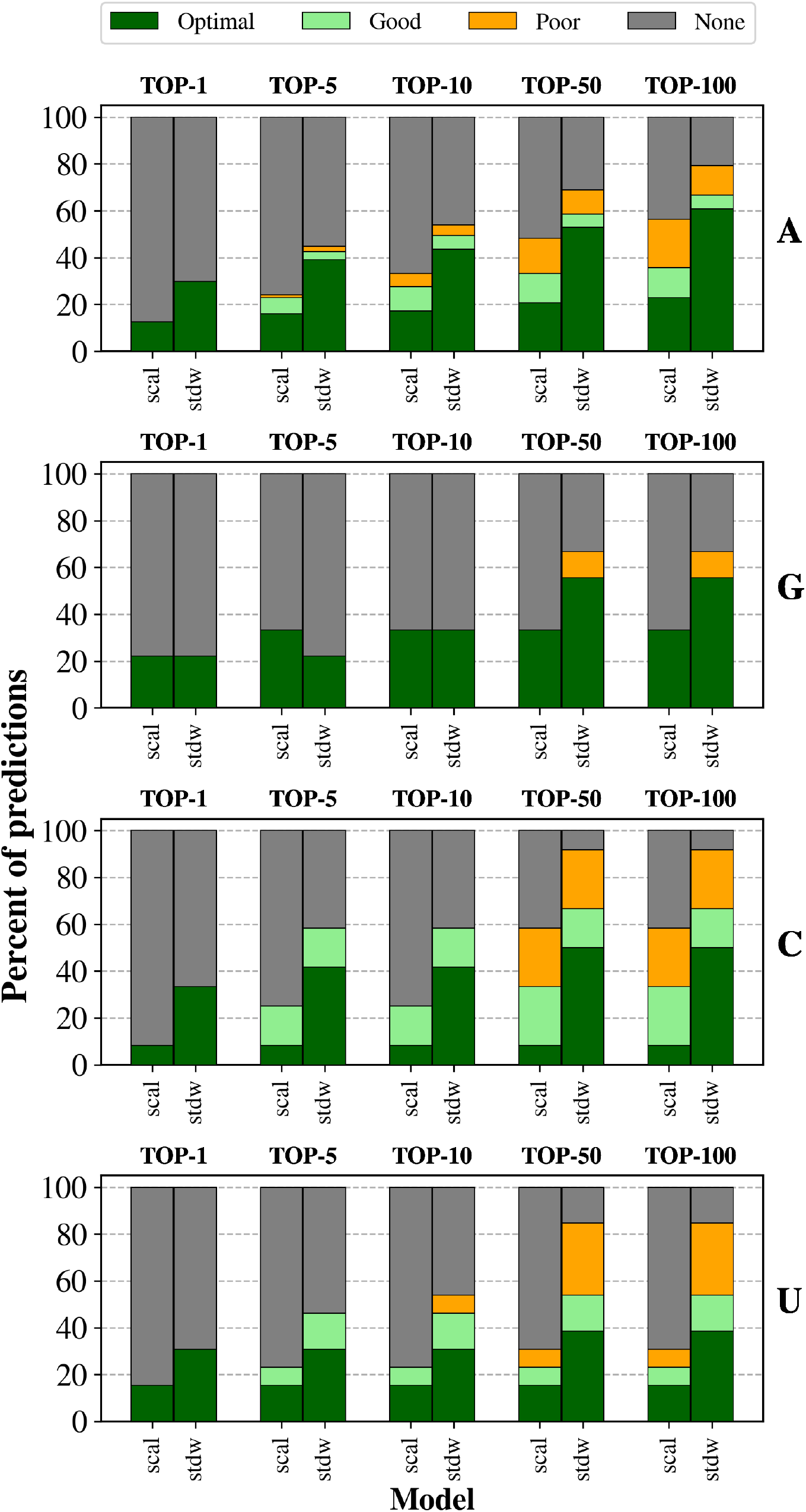
Nucleotide breakdown of the Binding Selectivity Predictions. Optimal: native nucleotide as the best ranked; good: native nucleotide in the ranked within a 2 kcal/mol range from the best ranked non-native nucleotide; poor: native nucleotide ranked out of the 2 kcal/mol range.

## Supplementary Note 4: Molecular features

**Fig. S10.**
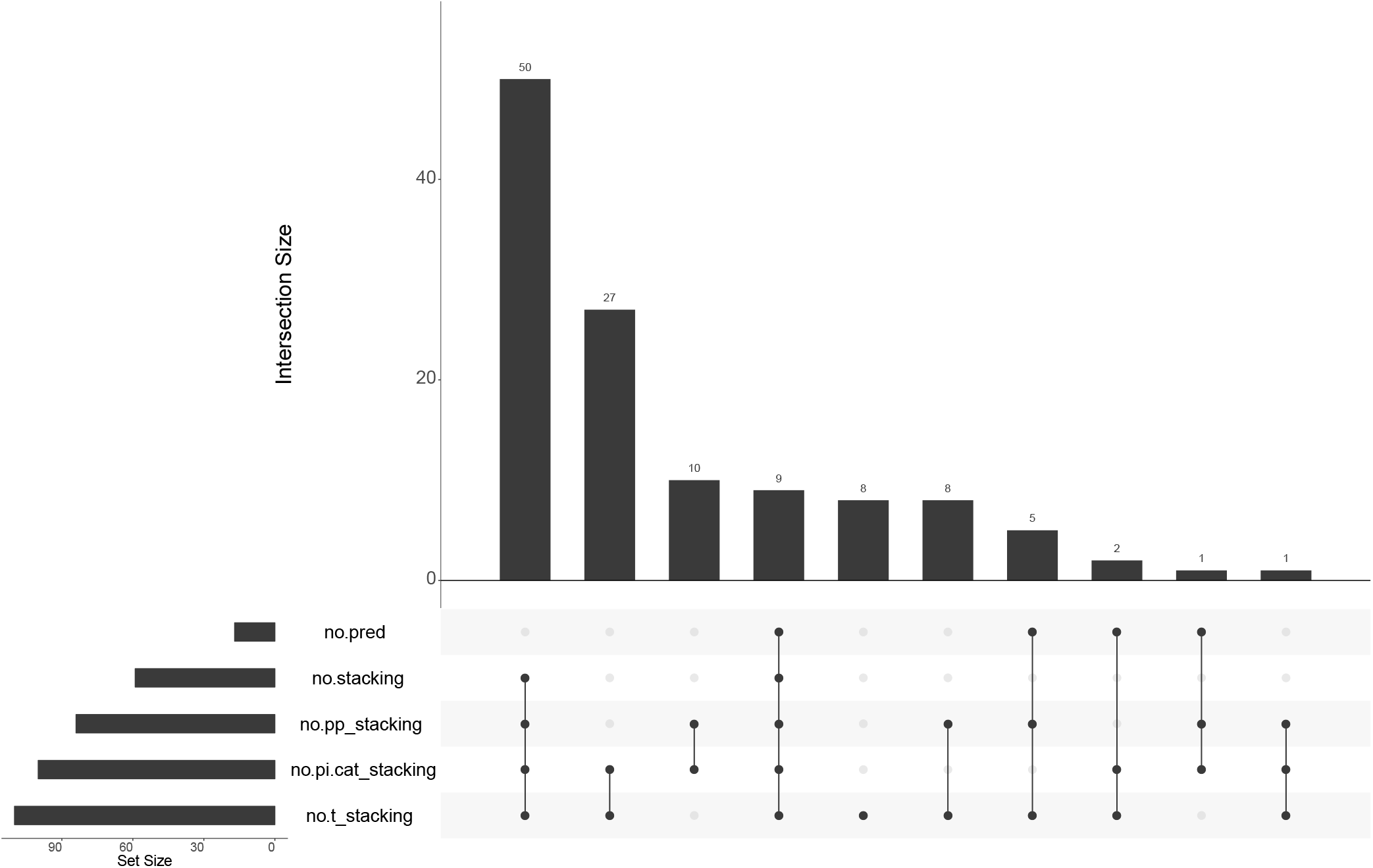
UpsetR diagram of stacking contributions for the Top-10 predictions. no.pp_satcking: no *π*-*π* stacking; no.pi.cat_stacking: no *π*-cation stacking; no.t_stacking: not stacking

**Table S1.** Frequencies of occurences for molecular features in the Top-10 non-predicted cases versus benchmark. Others: presence of additional nucleotidic (nucleic acid) fragment in the binsing site; metals: presence of metal(s) in the binding site; nwat.low: presence of number of water molecules below the threshold value; vol.low: volume of the binding site below the threshold value; syn: syn conformation of the nucleic acid base; pyr: pyrimidine; pur: purine; no.base.contacts: absence of contacts with the nucleic acid base; clash_aa: clash(es) with amino-acid residues; clash_w: clash(es) with water molecules; no.salt.bridges: absence of salt-bridge; no.stacking: absence of stacking.

| Features |  | Freq. Benchmark | Freq. no.pred |
| --- | --- | --- | --- |
| binding site | nwat.low | 62 | 59 |
|  | vol.low | 69 | 82 |
|  | others | 12 | 6 |
|  | metals | 36 | 24 |
| conformational | syn | 12 | 0 |
|  | pur | 79 | 71 |
|  | pyr | 21 | 23 |
| interaction | no.base.contacts | 12 | 12 |
|  | no.salt.bridges | 44 | 59 |
|  | no.stacking | 49 | 53 |
|  | clash aa | 22 | 18 |
|  | clash w | 33 | 41 |

**Table S2.** Frequencies of occurences for molecular features in the Top-10 for non-predicted cases of STDW-310 versus benchmark. Others: presence of additional nucleotidic (nucleic acid) fragment in the binsing site; metals: presence of metal(s) in the binding site; nwat.low: presence of number of water molecules below the threshold value; vol.low: volume of the binding site below the threshold value; syn: syn conformation of the nucleic acid base; pyr: pyrimidine; pur: purine; no.base.contacts: absence of contacts with the nucleic acid base; clash_aa: clash(es) with amino-acid residues; clash_w: clash(es) with water molecules; no.salt.bridges: absence of salt-bridge; no.stacking: absence of stacking.

| Features |  | Freq. Benchmark | Freq. STDW(R310) |
| --- | --- | --- | --- |
| binding site | nwat.low | 62 | 51 |
|  | vol.low | 69 | 72 |
|  | others | 12 | 6 |
|  | metals | 36 | 30 |
| conformational | syn | 12 | 17 |
|  | pur | 79 | 83 |
|  | pyr | 21 | 17 |
| interaction | no.base.contacts | 12 | 11 |
|  | no.salt.bridges | 44 | 62 |
|  | no.stacking | 49 | 49 |
|  | clash aa | 22 | 21 |
|  | clash w | 33 | 40 |

**Table S3.**
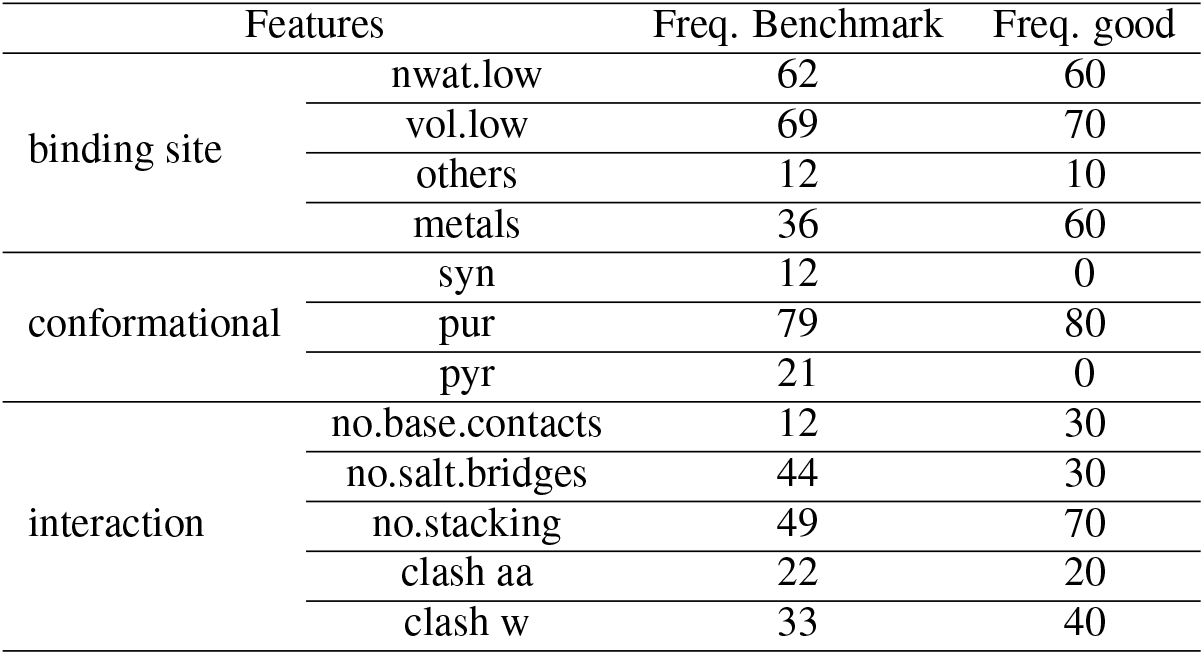
Frequencies of occurences for molecular features in the Top-10 for non-optimal (good) predictions. Others: presence of additional nucleotidic (nucleic acid) fragment in the binsing site; metals: presence of metal(s) in the binding site; nwat.low: presence of number of water molecules below the threshold value; vol.low: volume of the binding site below the threshold value; syn: syn conformation of the nucleic acid base; pyr: pyrimidine; pur: purine; no.base.contacts: absence of contacts with the nucleic acid base; clash_aa: clash(es) with amino-acid residues; clash_w: clash(es) with water molecules; no.salt.bridges: absence of salt-bridge; no.stacking: absence of stacking.

Attached Supplementary Data 9 (Data-S9.txt): raw data corresponding to the number of water molecules around the ligand at a distance up to 4Å.

Attached Supplementary Data 10 (Data-S10.csv): raw data corresponding to the variations of the binding site’s volume for each protein of the benchmark in three conditions: experimental, SCAL, and STDW models.

**Table S4.** Variations in the binding site’s volume for the subset of protein-nucleotides complexes with no prediction in the Top-10. The volume of reference corresponds to that of the experimental structure; the modified volumes are calculated for both the SCAL and STDW models. Only the cases where the variation equals or exceeds 100Å^3^ are considered. UP: increase of the binding site’s volume. DOWN: decrease of the binding site’s volume.

| Volumes |  | Freq. Benchmark | Freq. nopred. |
| --- | --- | --- | --- |
| SCAL | UP | 12 | 0 |
|  | DOWN | 19 | 18 |
| STDW | UP | 13 | 0 |
|  | DOWN | 21 | 35 |

**Fig. S11.**
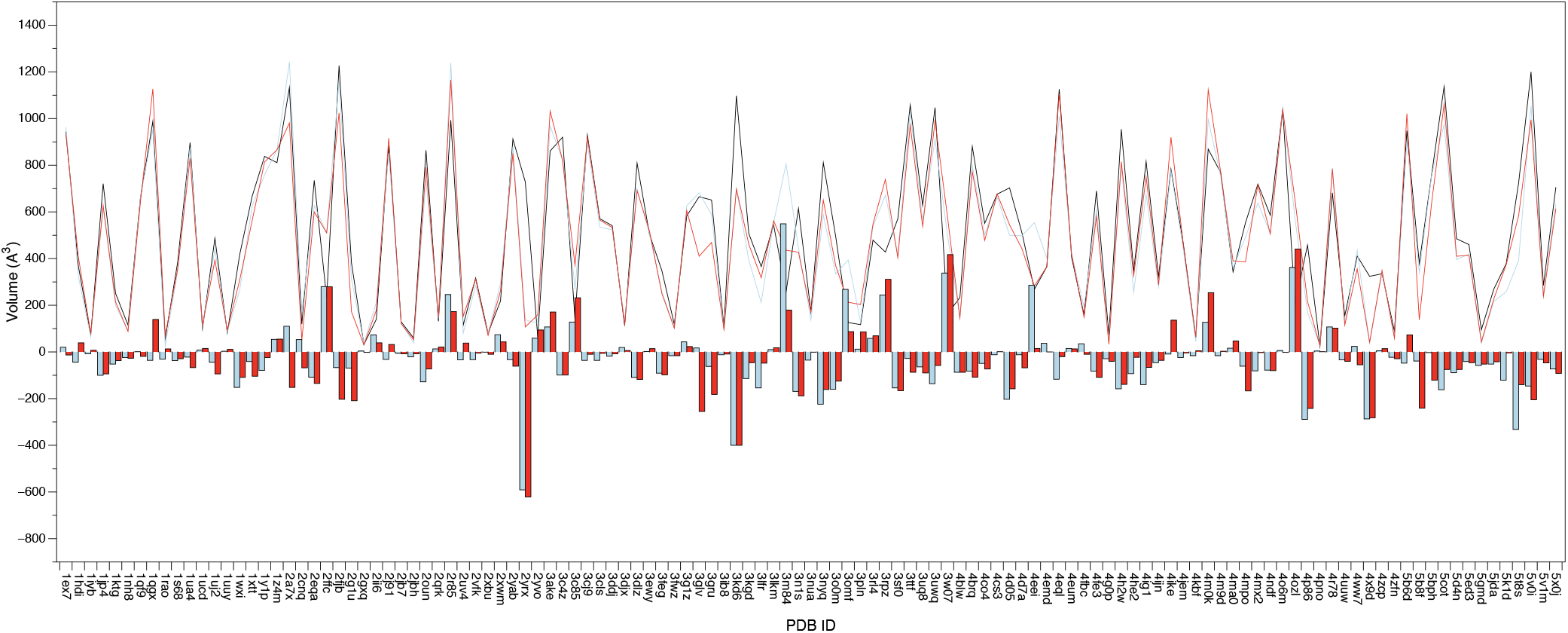
Variations in the volume of the binding site. Black line: experimental structure; Blue line: optimized structure for the SCAL model; Red line: optimized structure for the STDW model. The histograms indicate a decreasing of the volume for the negative values and an increasing for the positive values. The calculation of volume does not take into account the water molecules.

**Table S5.**
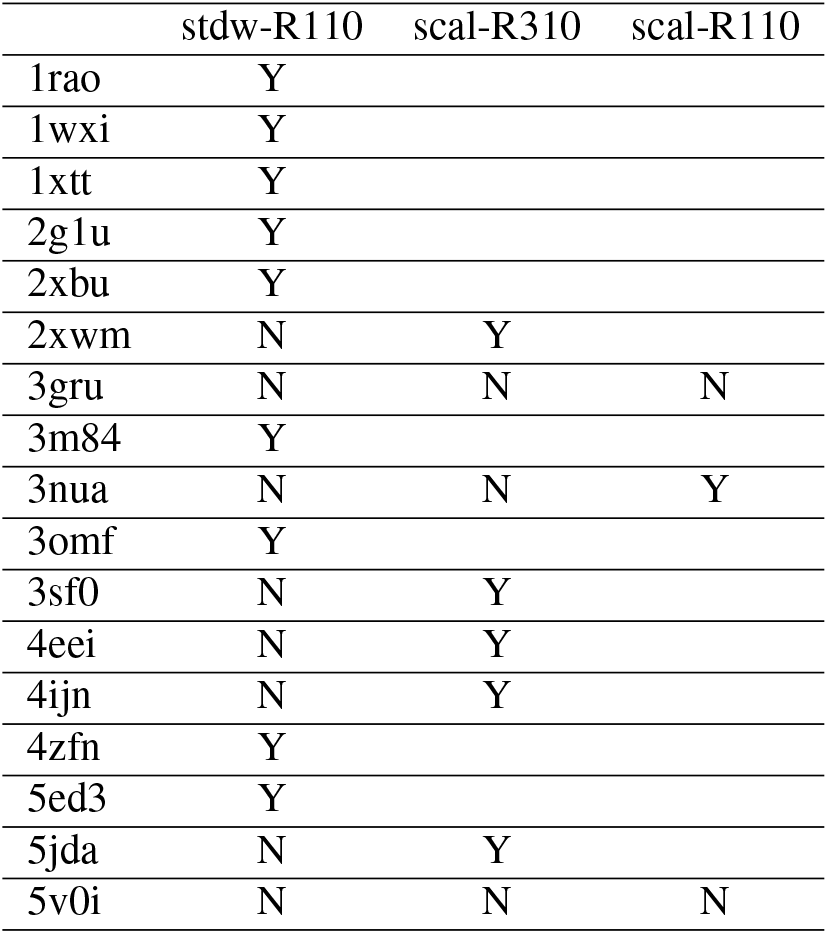
Impact of the nonbonded model and phosphate patch on the recovery effect of the Top-10 no-prediction subset. Y: recovered prediction using a different model and patch; N: no recovered prediction with the given model and patch.

**Fig. S12.**
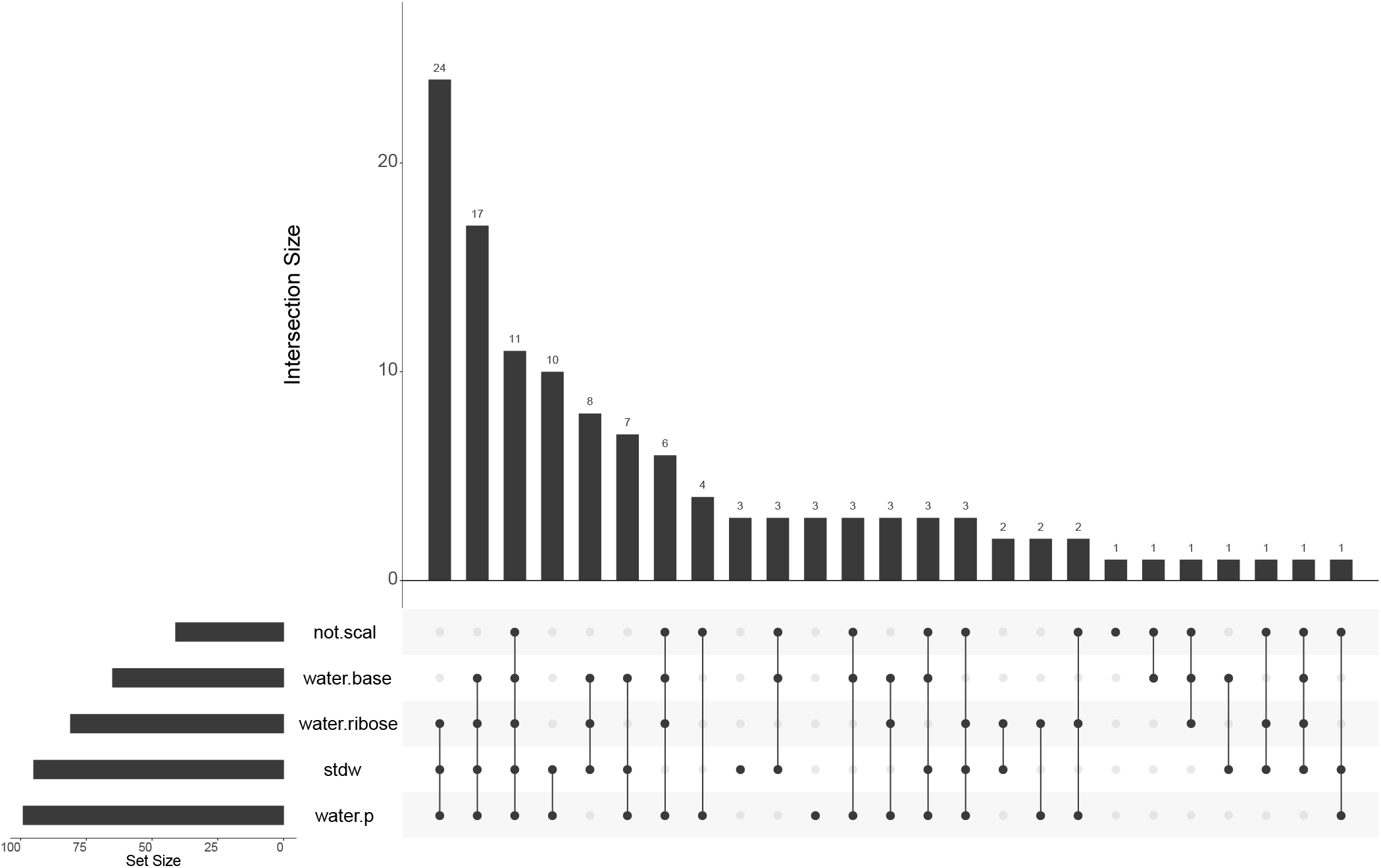
UpsetR diagram of water-mediated contacts for the Top-10 predictions. not.scal: no prediction with the SCAL model; stdw: predictions with STDW model; water.base: presence of water-mediated contacts with the nucleic acid base; water.ribose: presence of water-mediated contacts with the ribose; water.p presence of water-mediated contacts with the phosphate group.

Attached Supplementary Data 11 (Data-S11.csv): raw data corresponding to the molecular features associated with the Top-10 predictions.

