## Supplementary figures and images for "MCSS-based Predictions of Binding Mode and Selectivity of Nucleotide Ligands"

### 1wxi_benchmark-121_selectivity_details.png

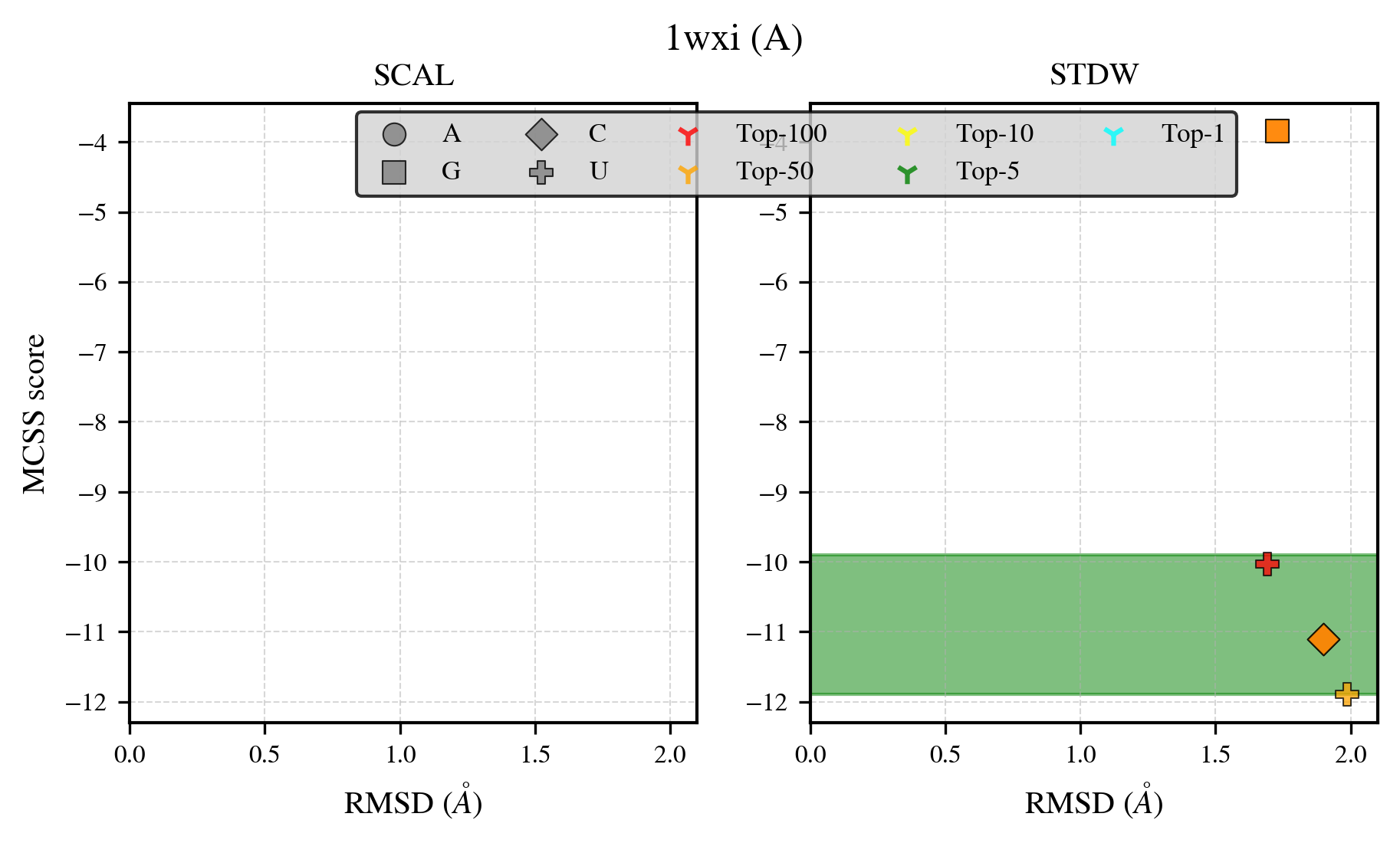

### 2g1u_benchmark-121_selectivity_details.png

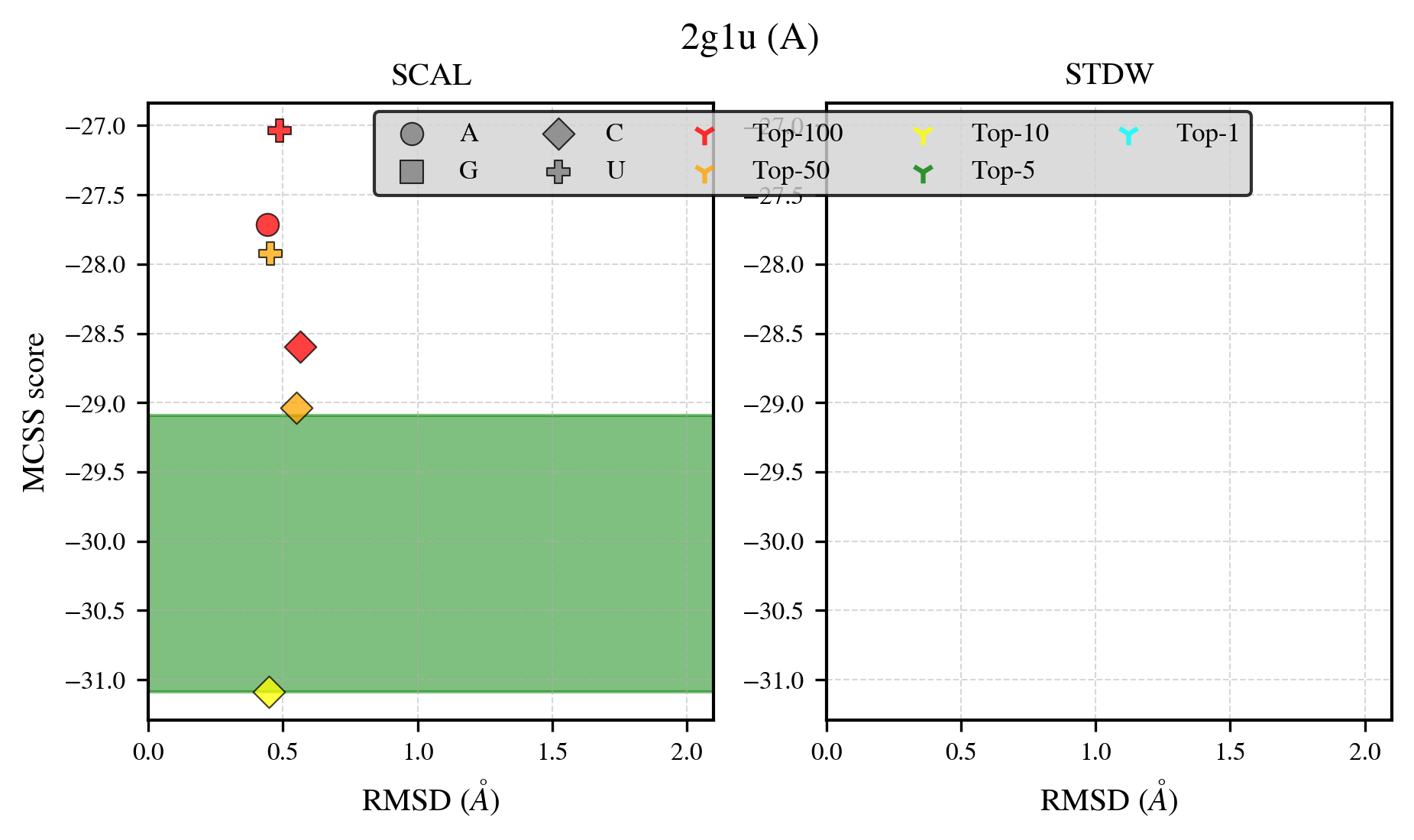

### 2jb7_benchmark-121_selectivity_details.png

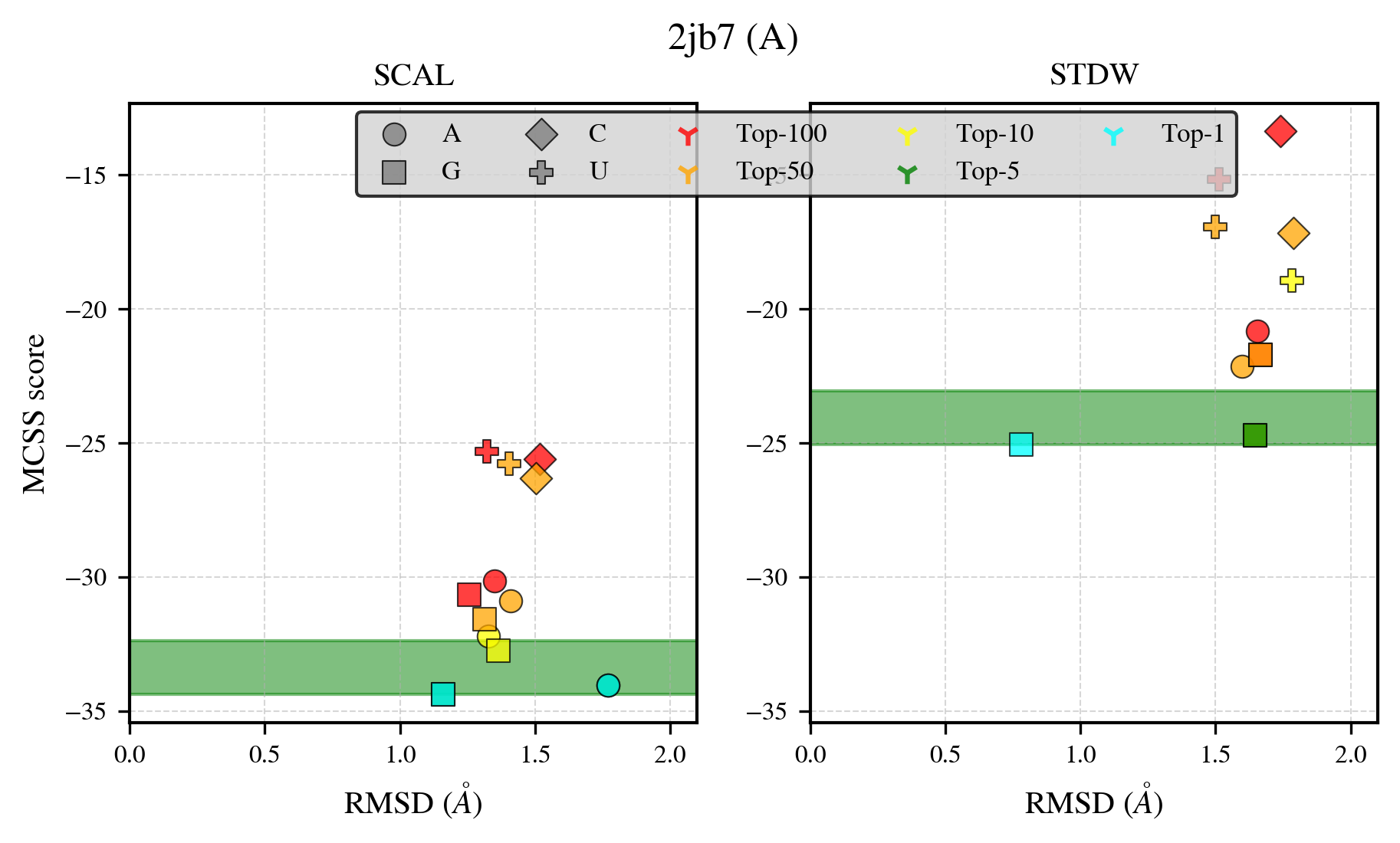

### 2vfk_benchmark-121_selectivity_details.png

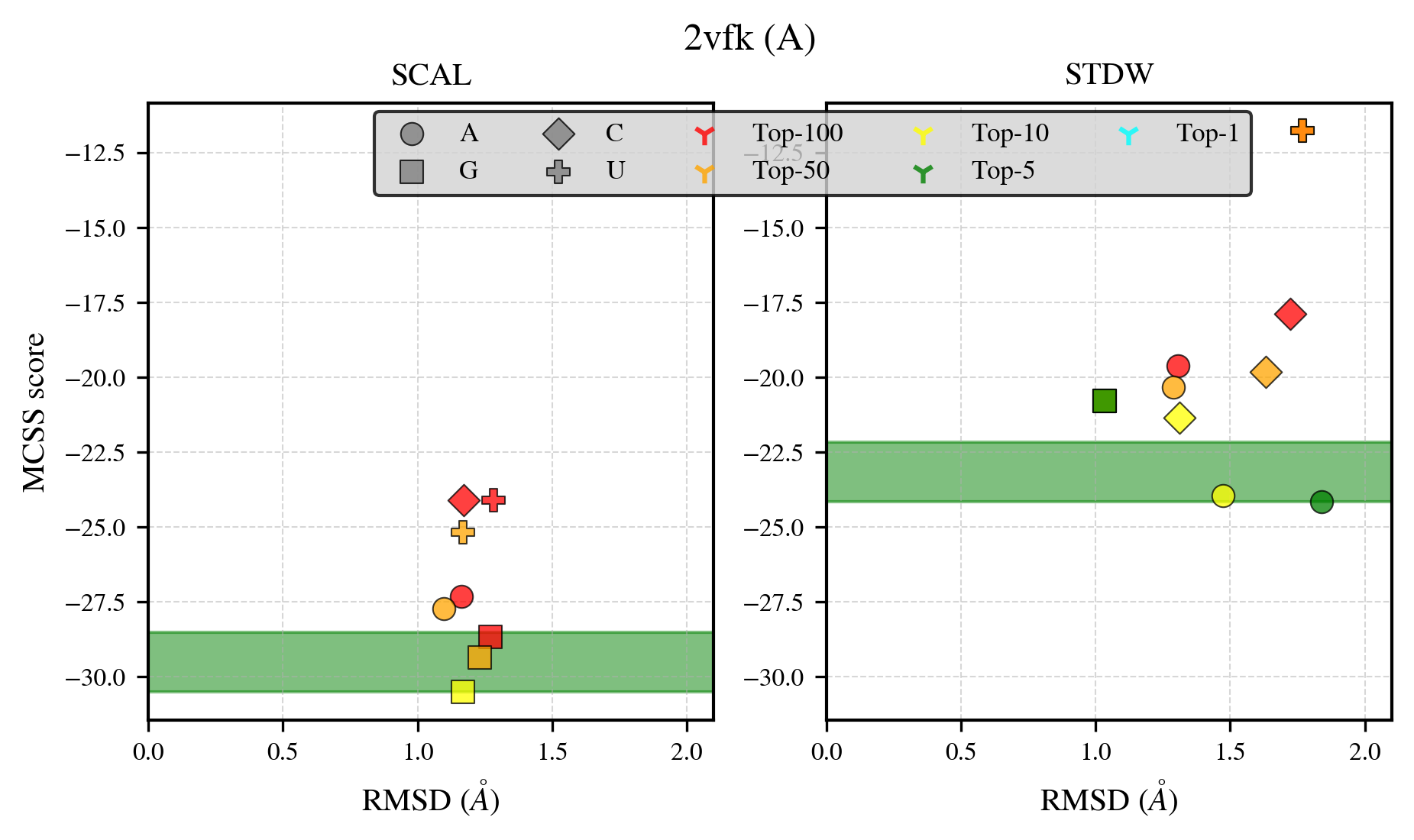

### 2xwm_benchmark-121_selectivity_details.png

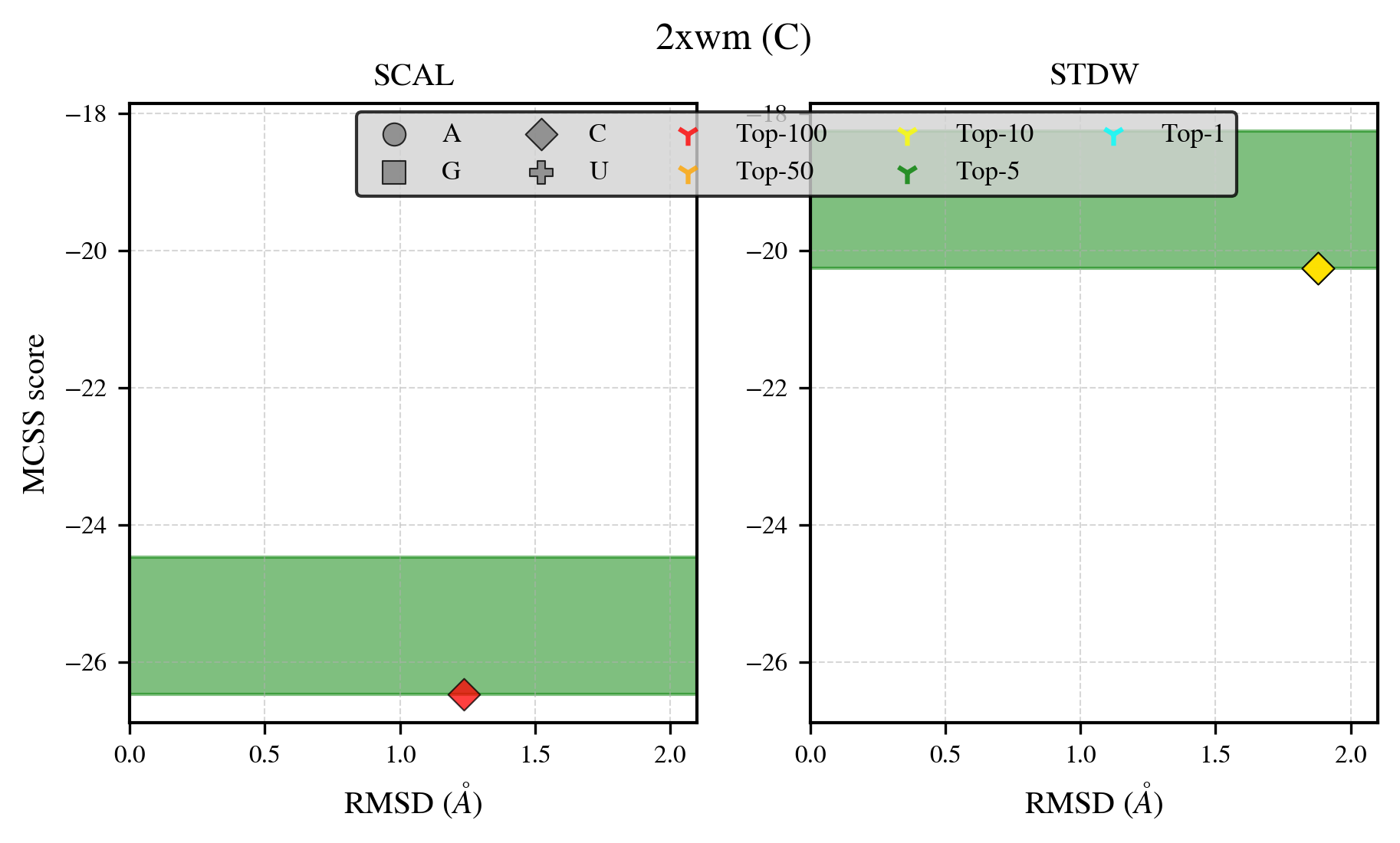

### 3feg_benchmark-121_selectivity_details.png

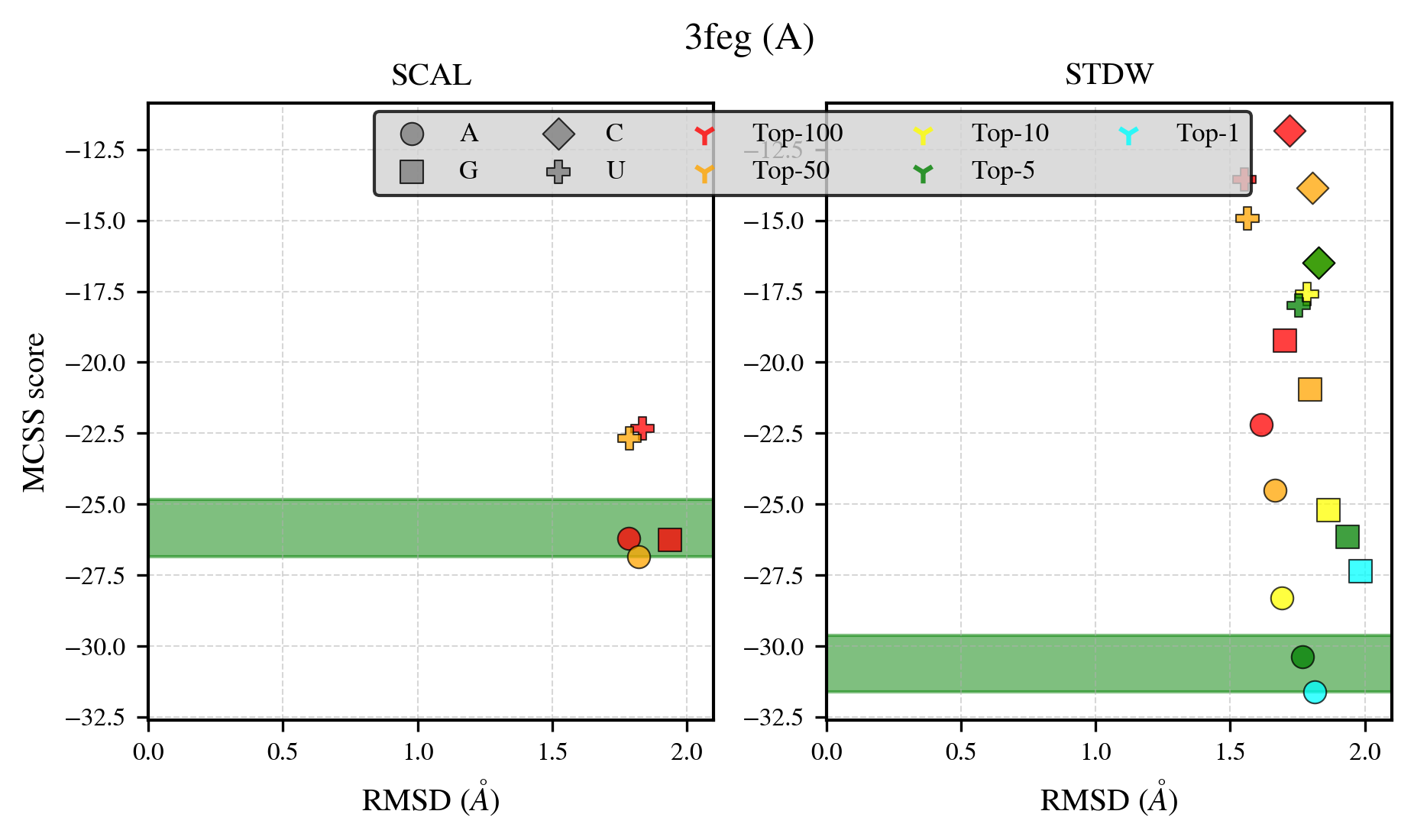

### 3fwz_benchmark-121_selectivity_details.png

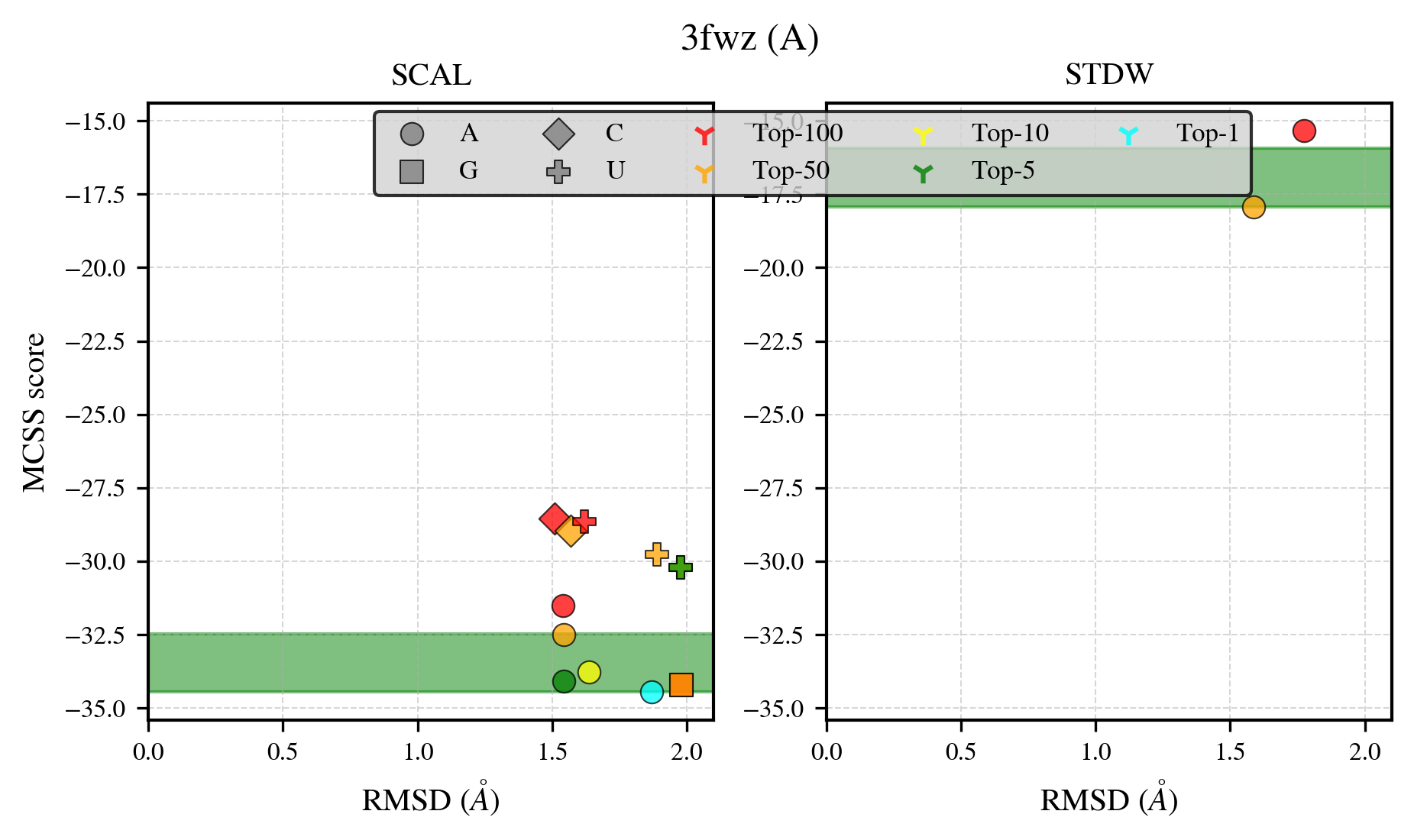

### 3g1z_benchmark-121_selectivity_details.png

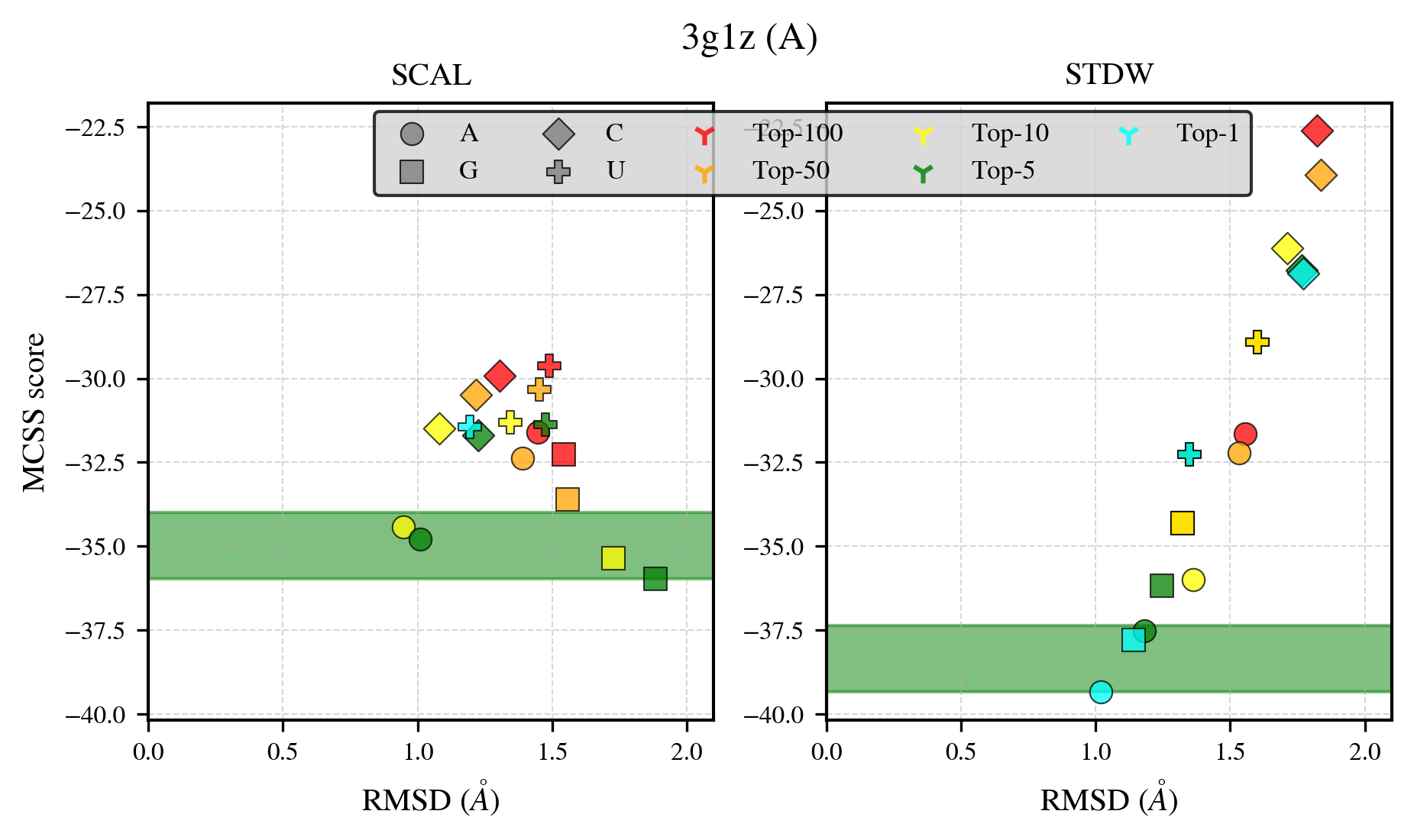

### 3ib8_benchmark-121_selectivity_details.png

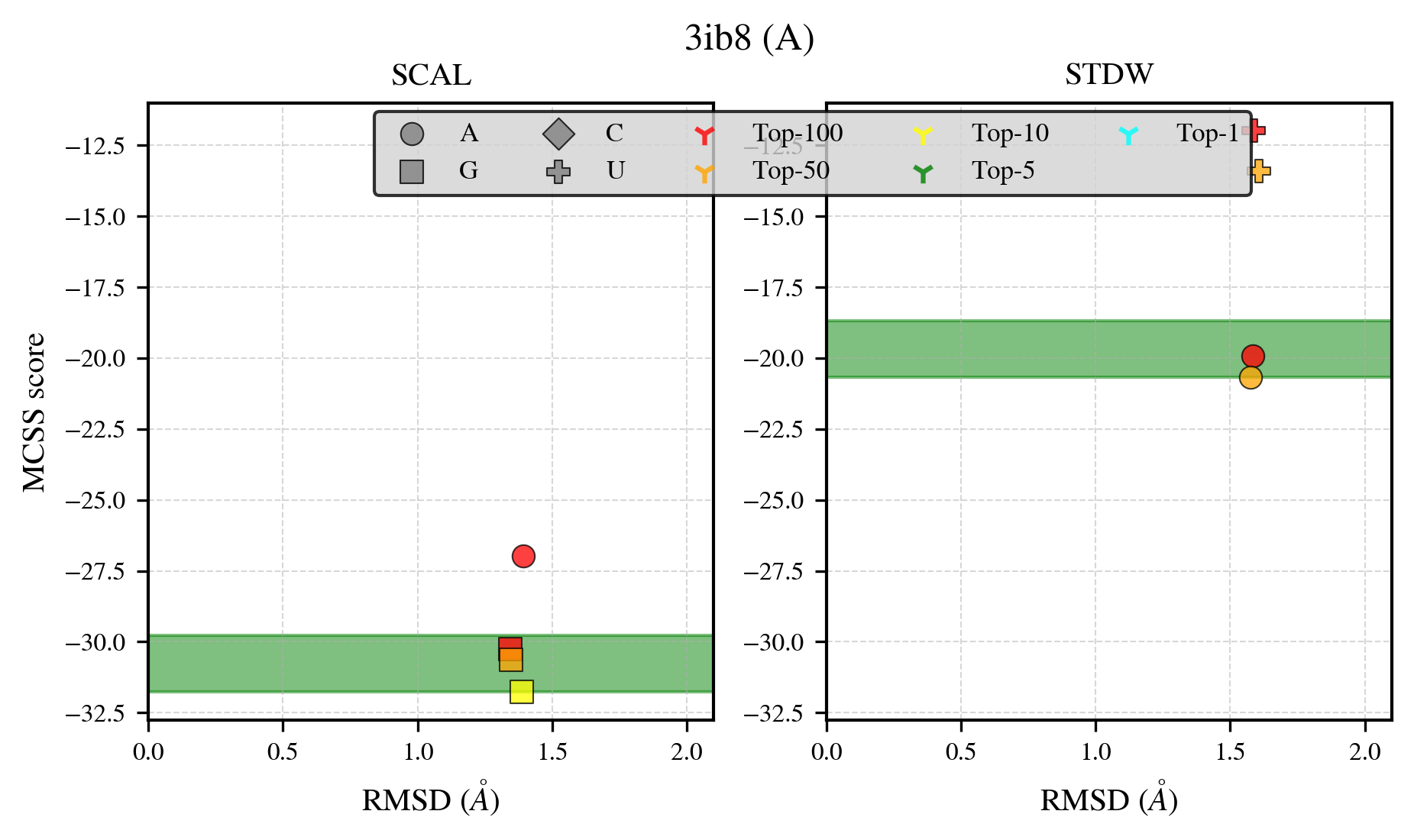

### 3o0m_benchmark-121_selectivity_details.png

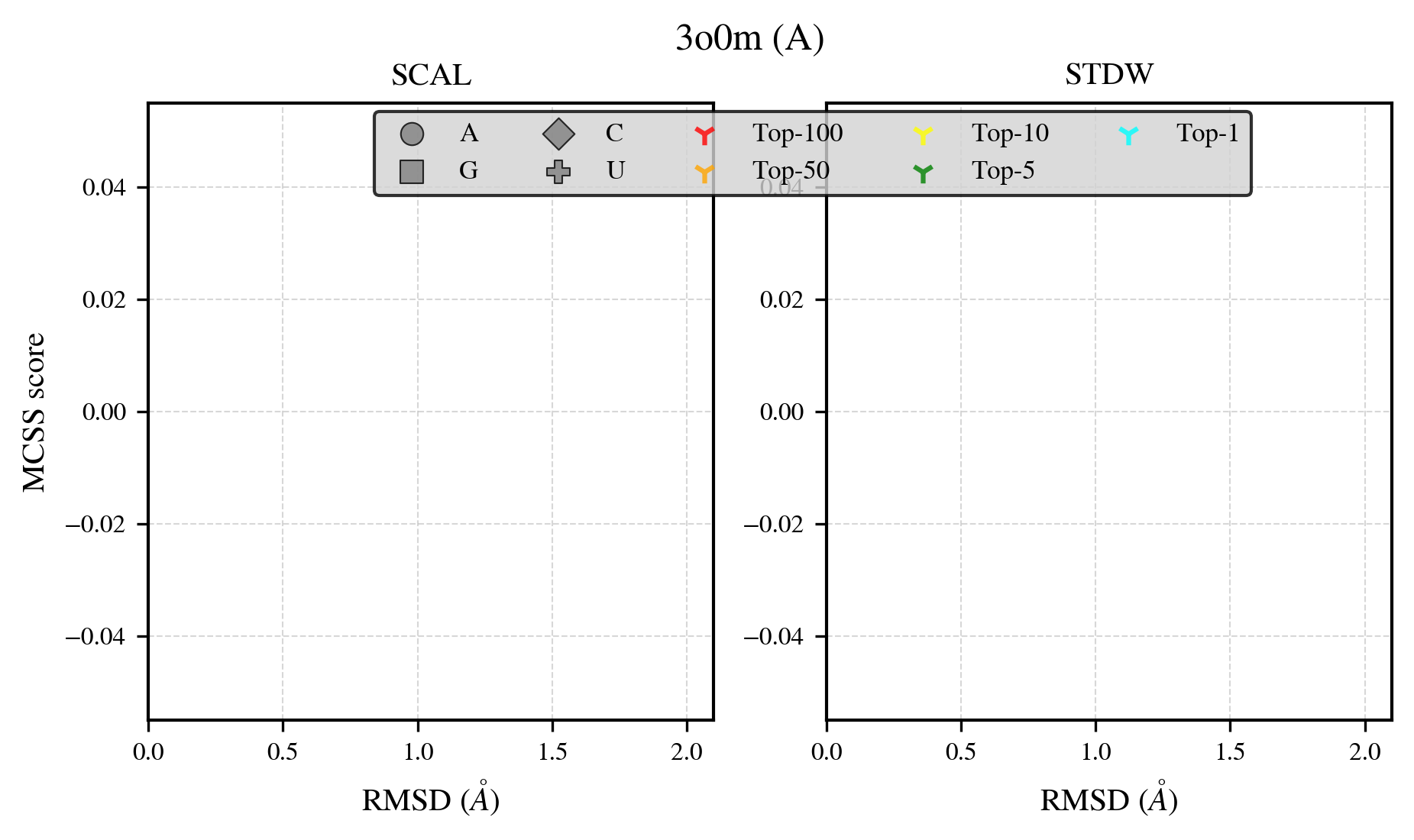

### 3sf0_benchmark-121_selectivity_details.png

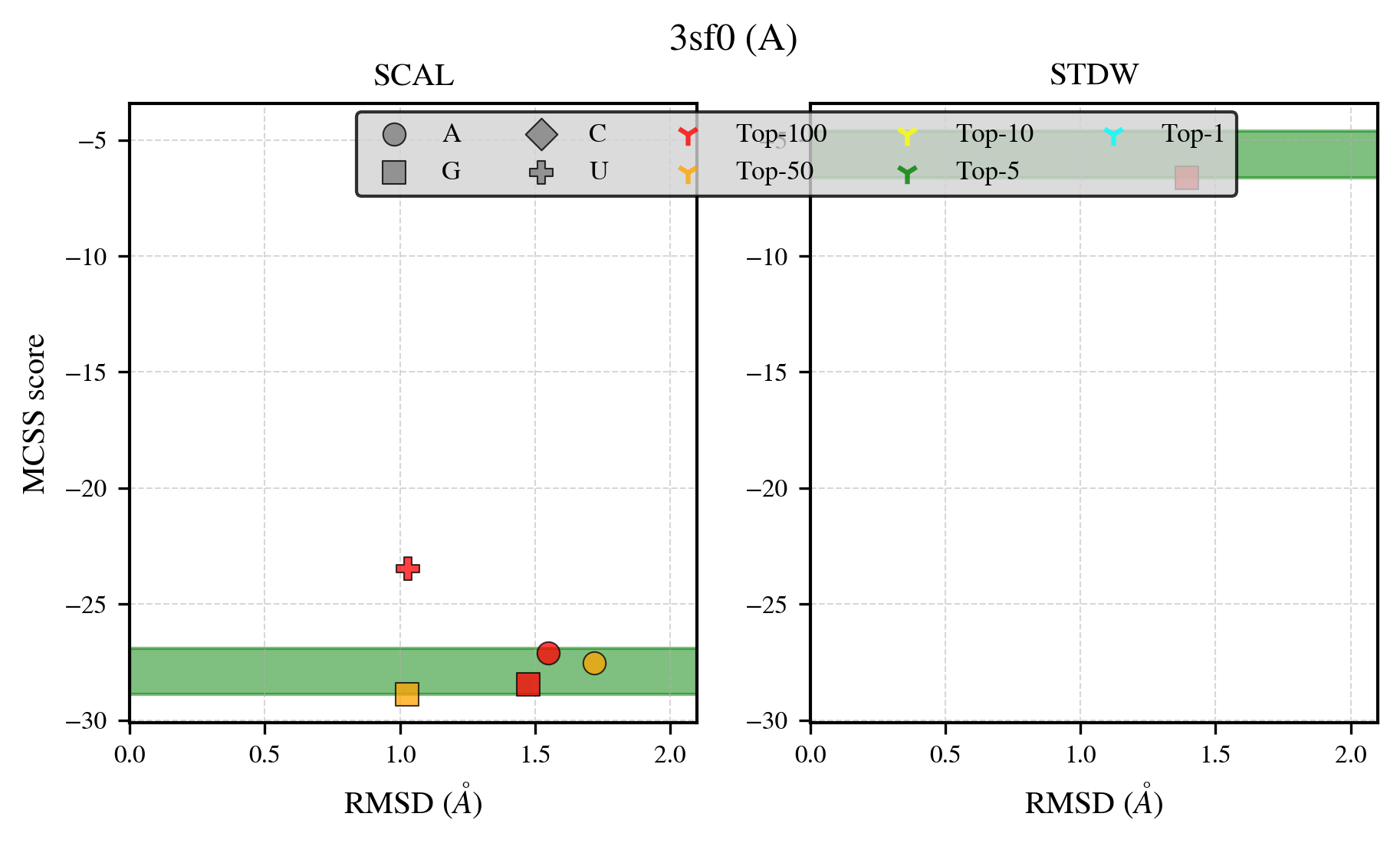

### 3ttf_benchmark-121_selectivity_details.png

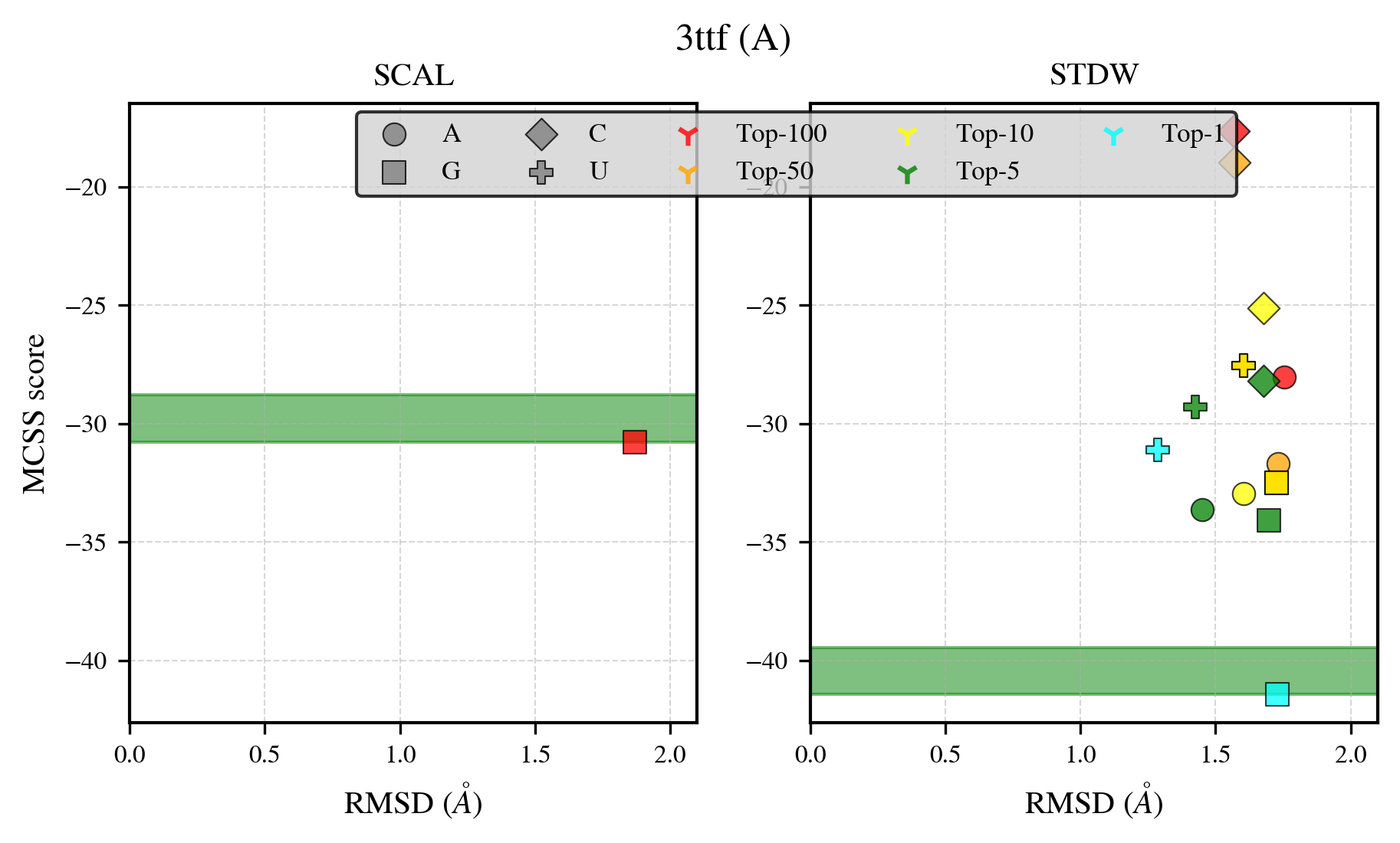

### 4co4_benchmark-121_selectivity_details.png

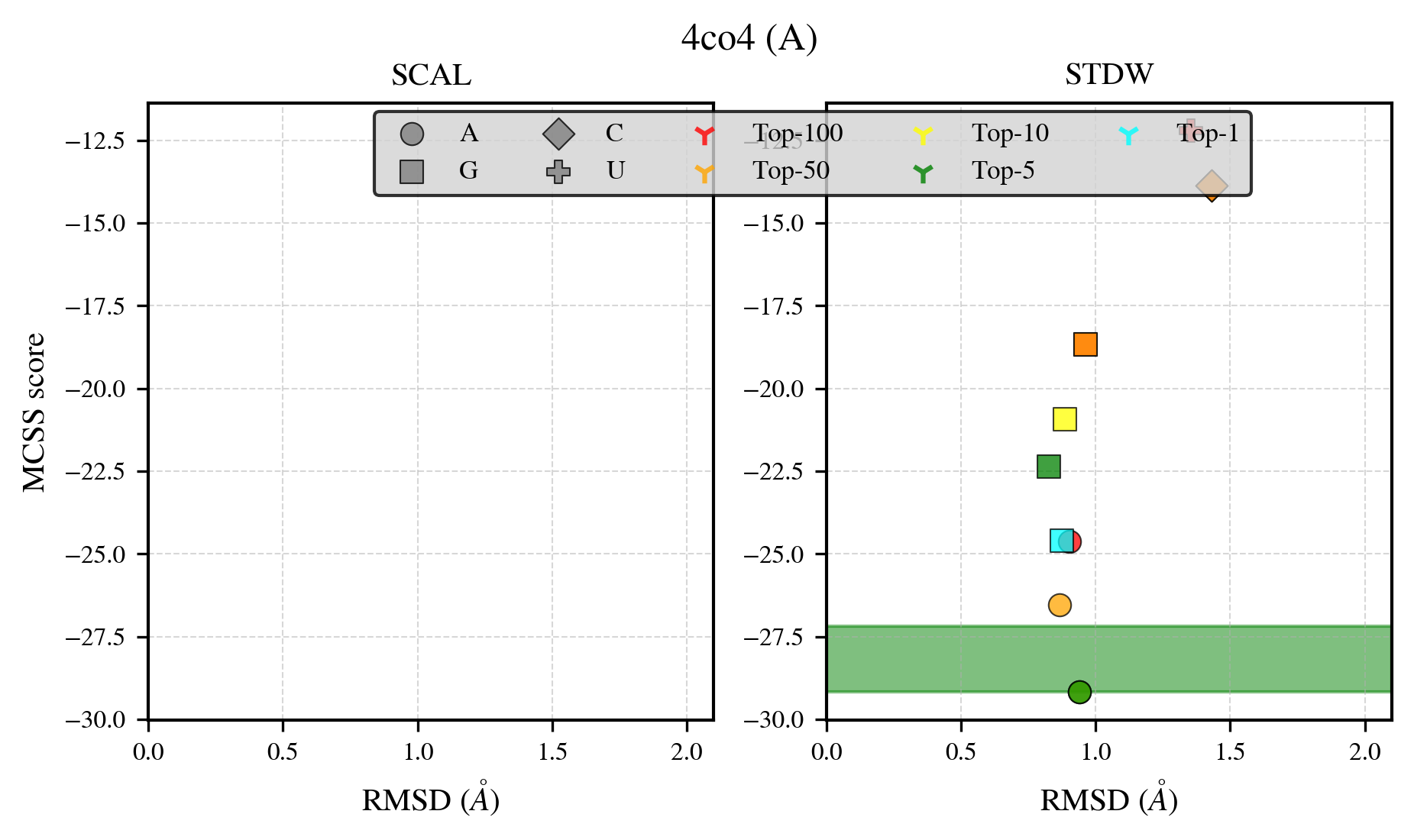

### 4eum_benchmark-121_selectivity_details.png

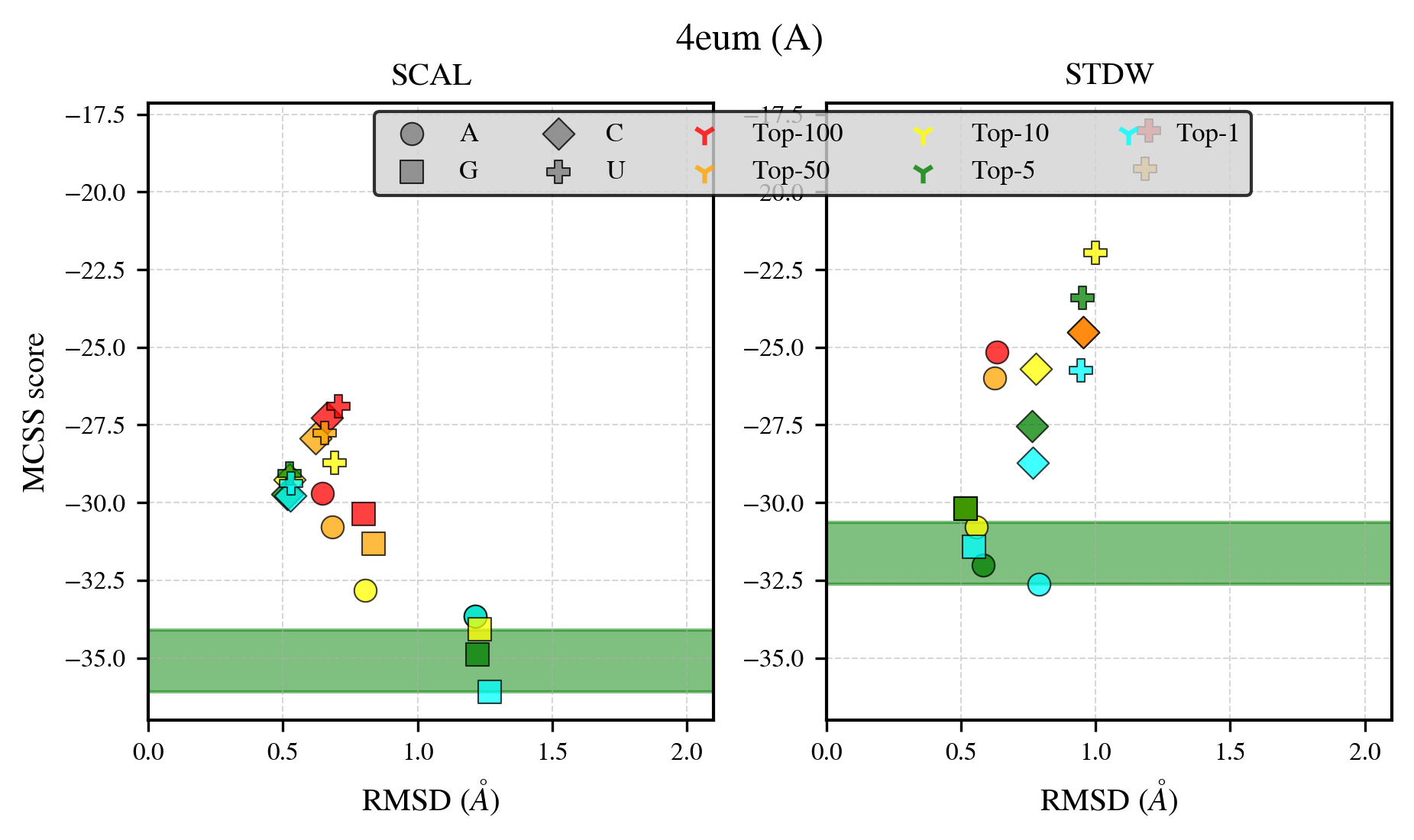

### 4g0p_benchmark-121_selectivity_details.png

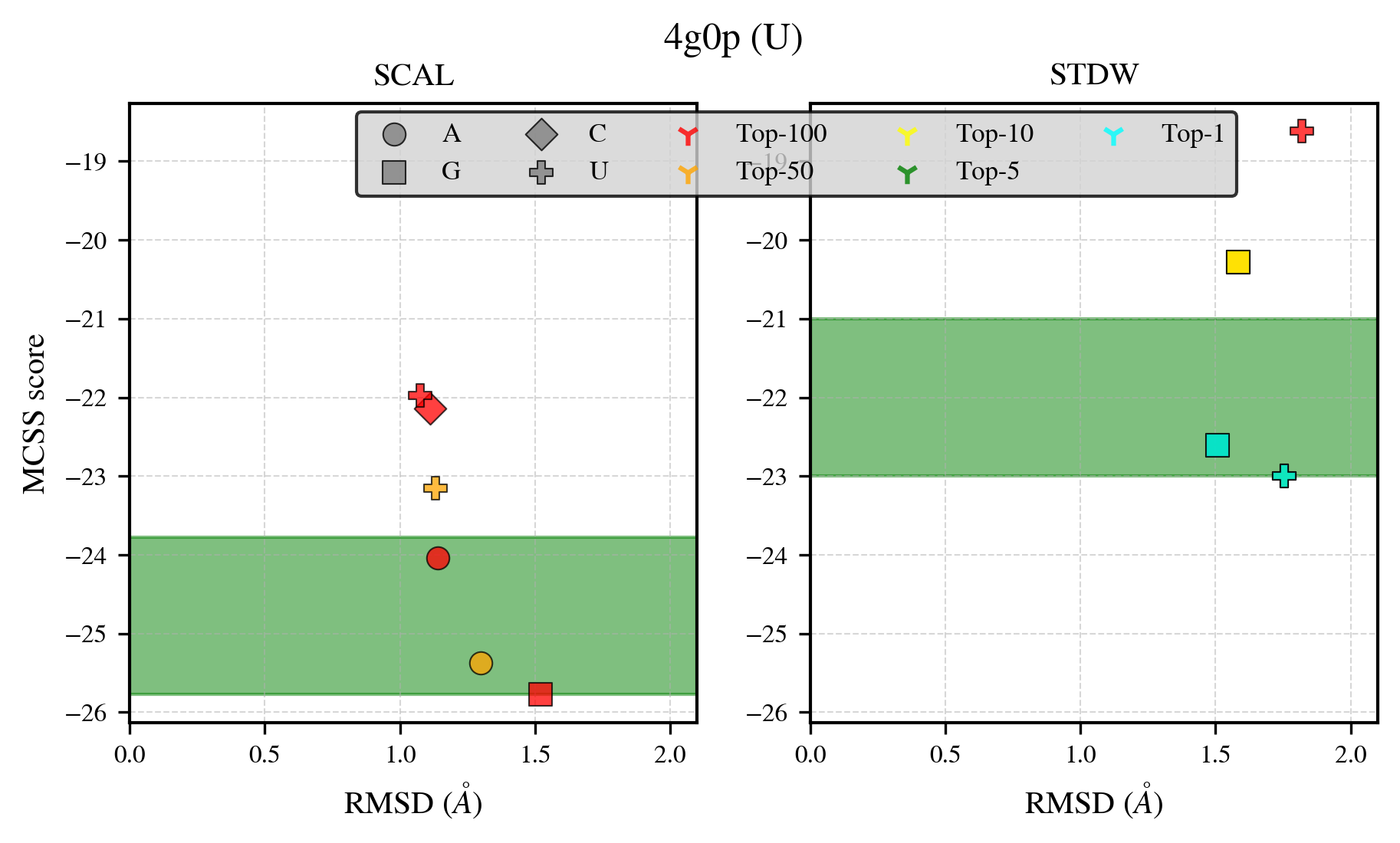

### 4mx2_benchmark-121_selectivity_details.png

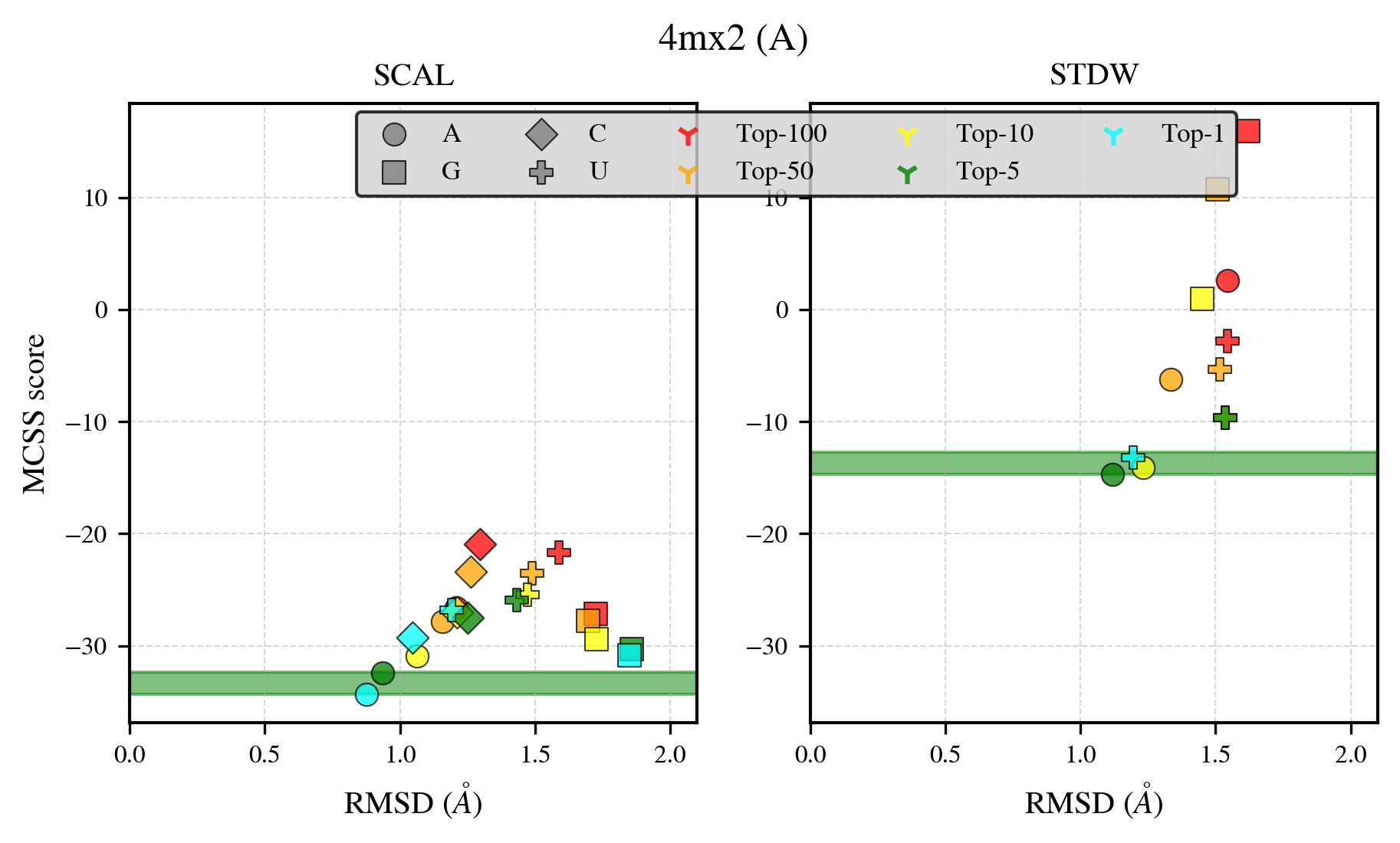

### 4p86_benchmark-121_selectivity_details.png

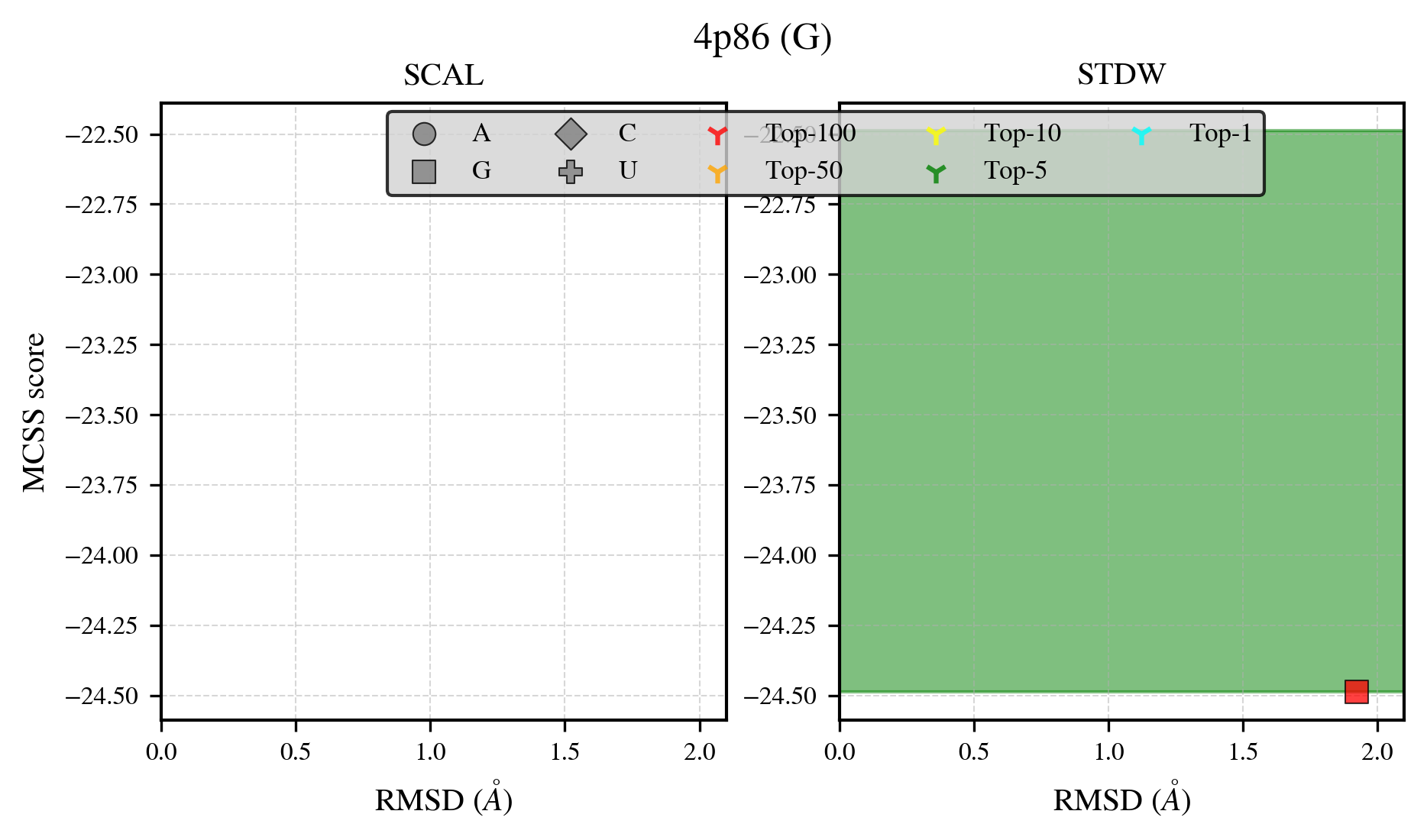

### 4pno_benchmark-121_selectivity_details.png

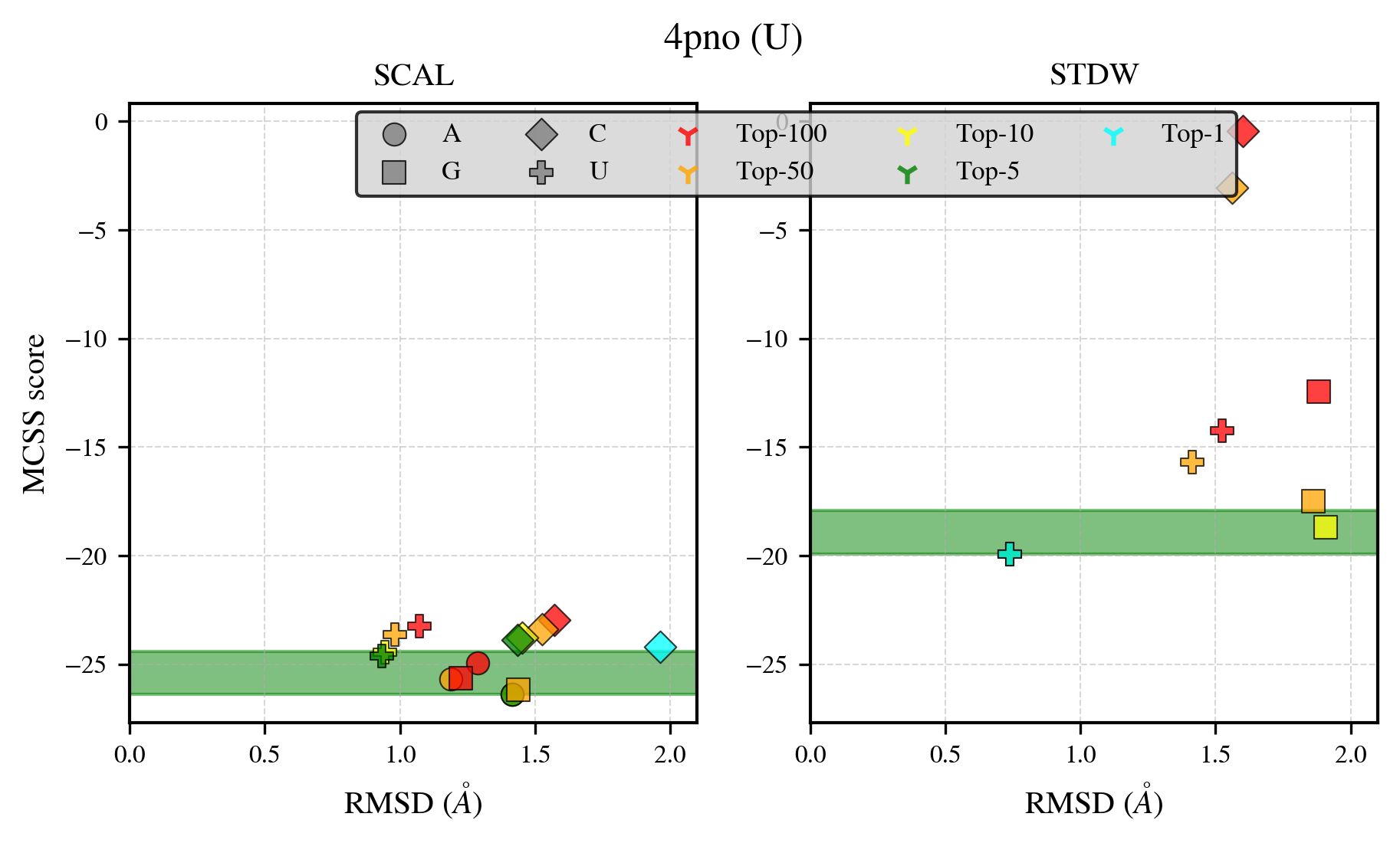

### 4uuw_benchmark-121_selectivity_details.png

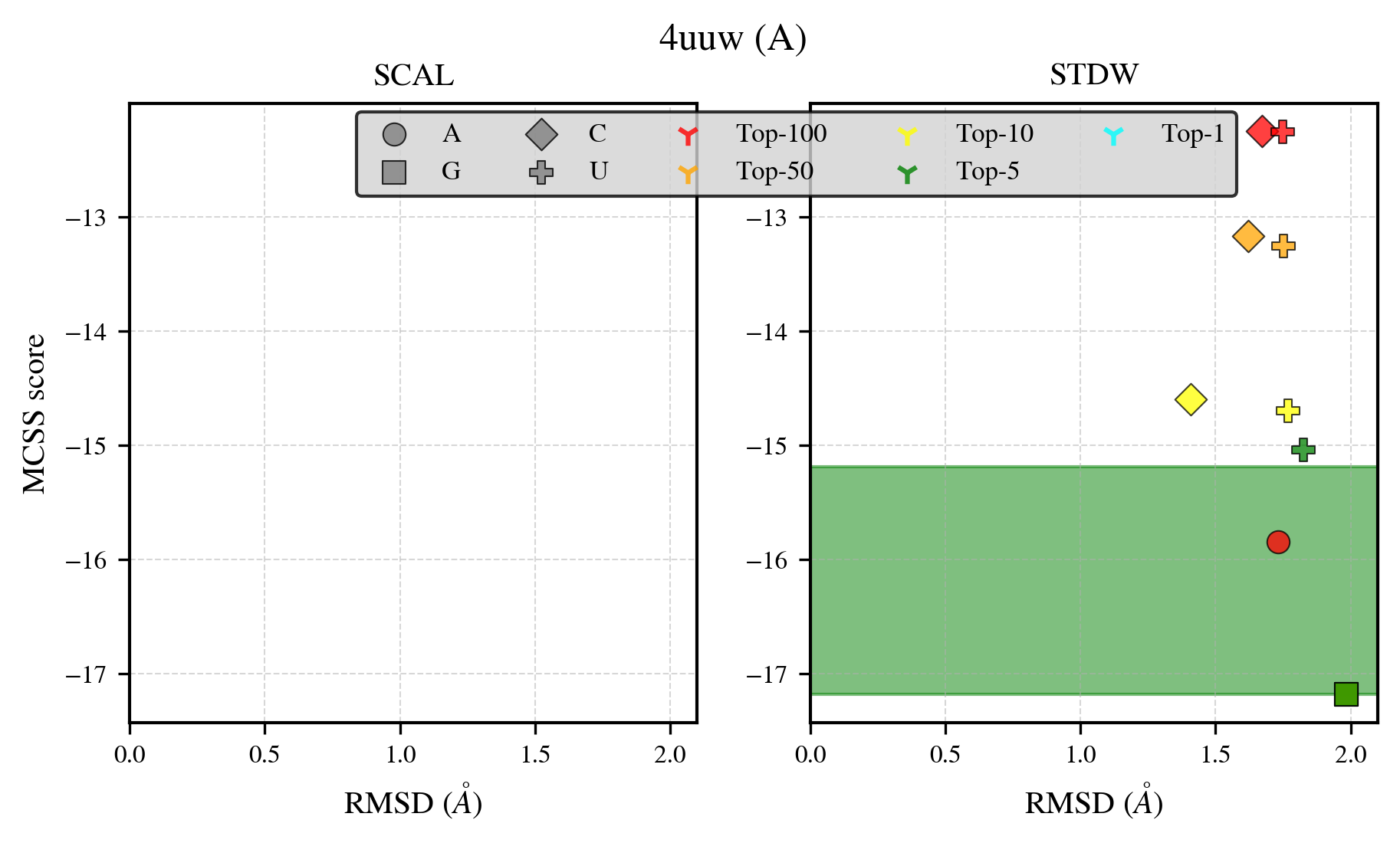

### 5jda_benchmark-121_selectivity_details.png

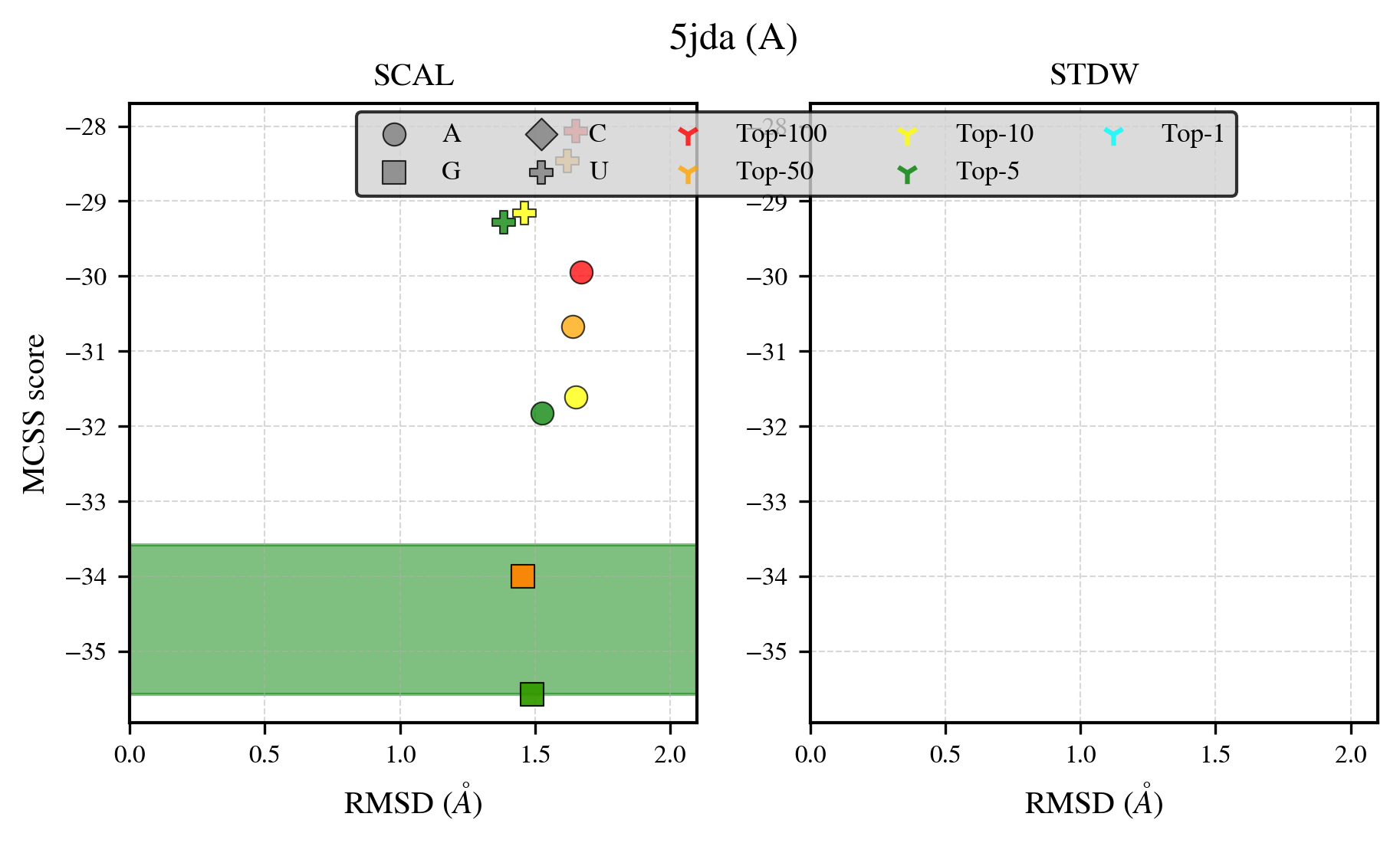

### 5t8s_benchmark-121_selectivity_details.png

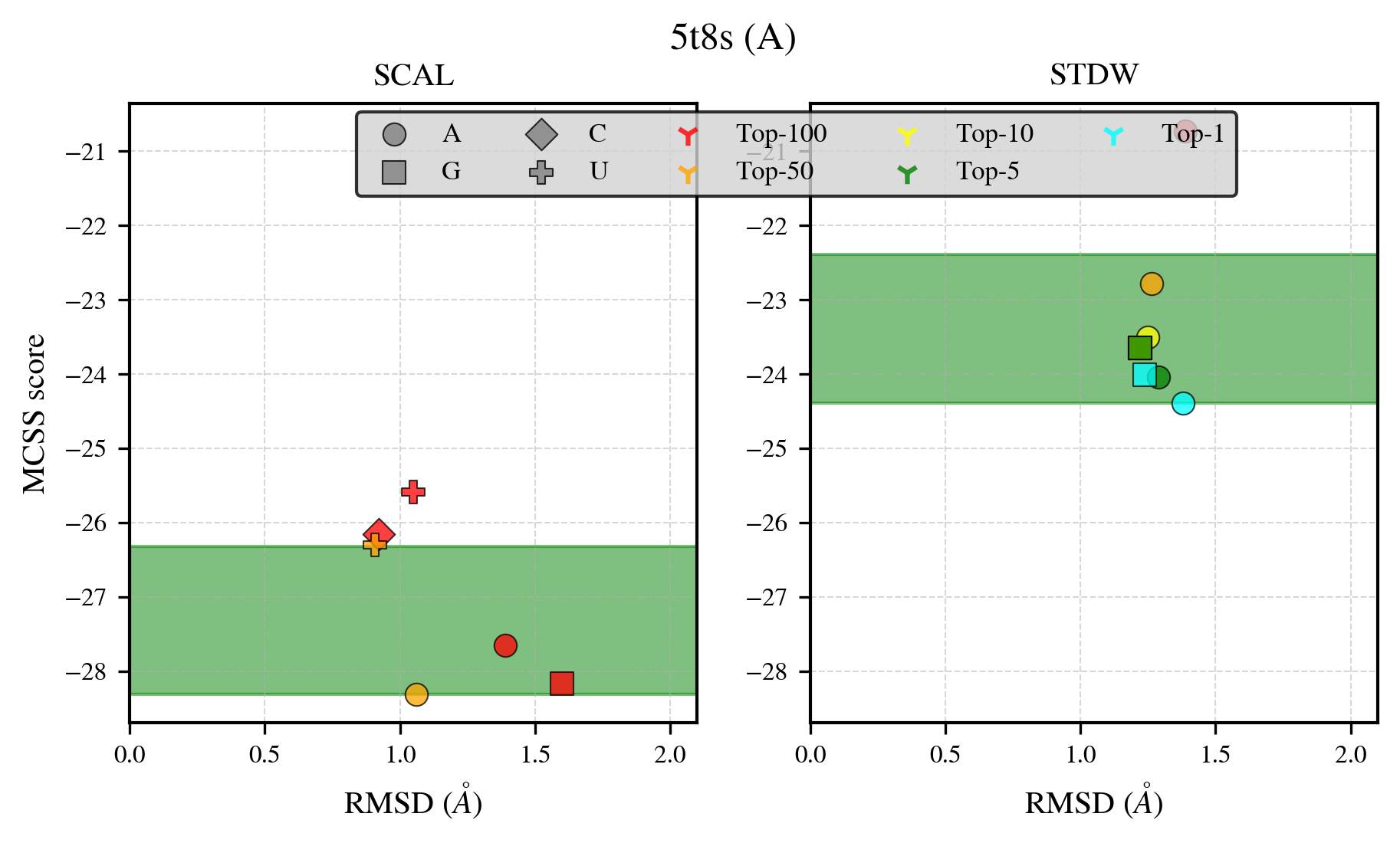
