## Supplementary figures and images for "MCSS-based Predictions of Binding Mode and Selectivity of Nucleotide Ligands"

### 1ex7_benchmark-121_selectivity_details.png

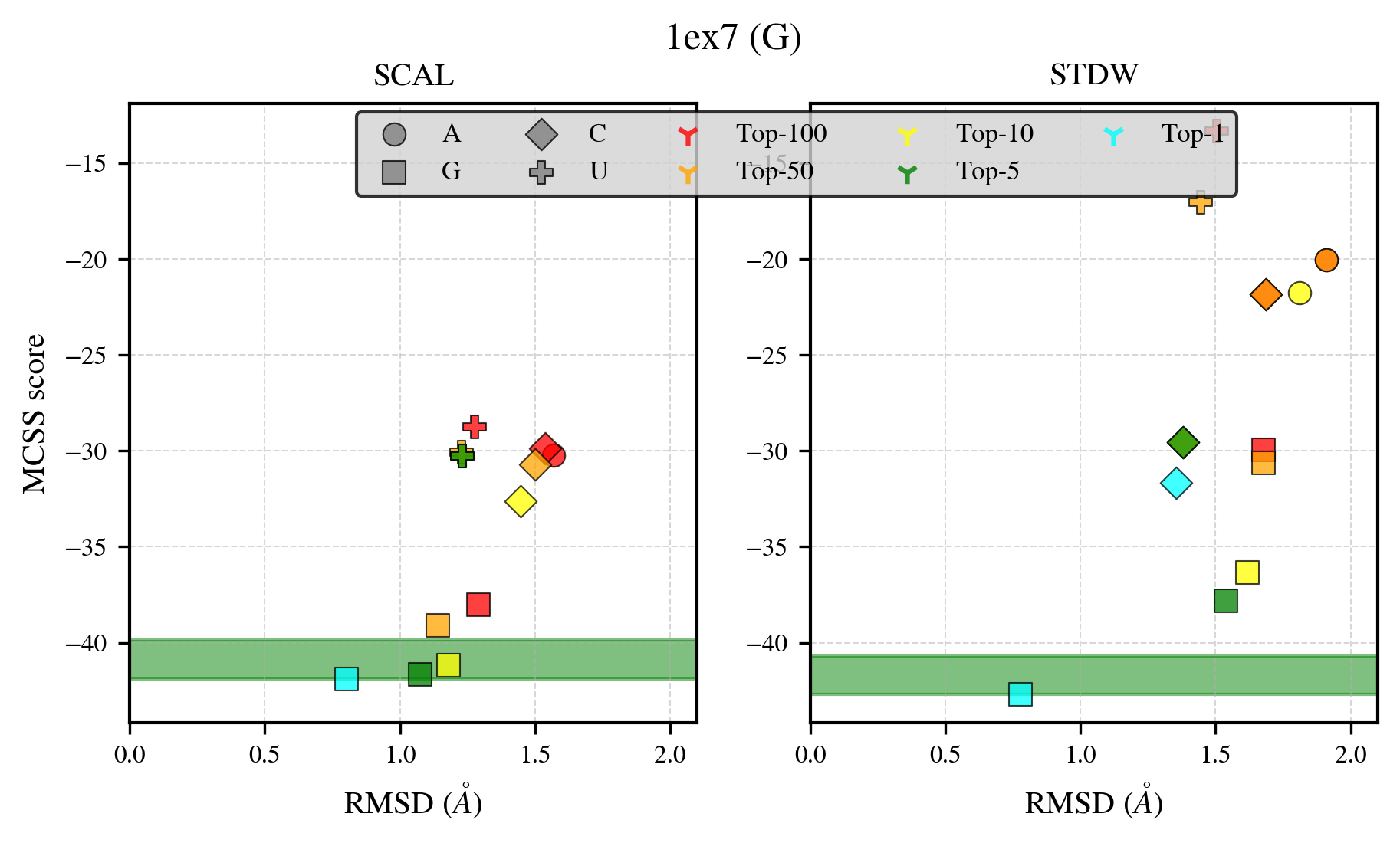

### 1hdi_benchmark-121_selectivity_details.png

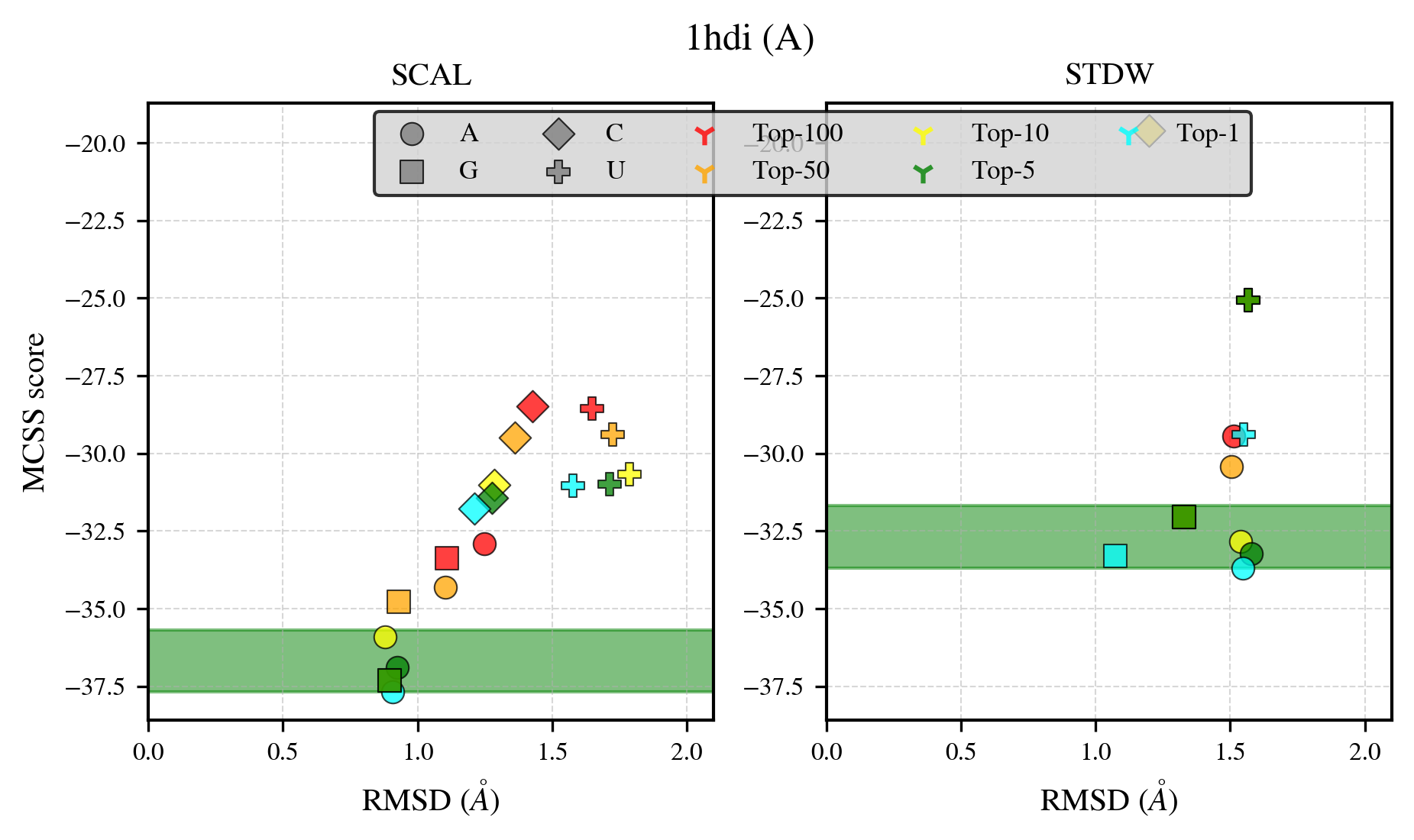

### 1iyb_benchmark-121_selectivity_details.png

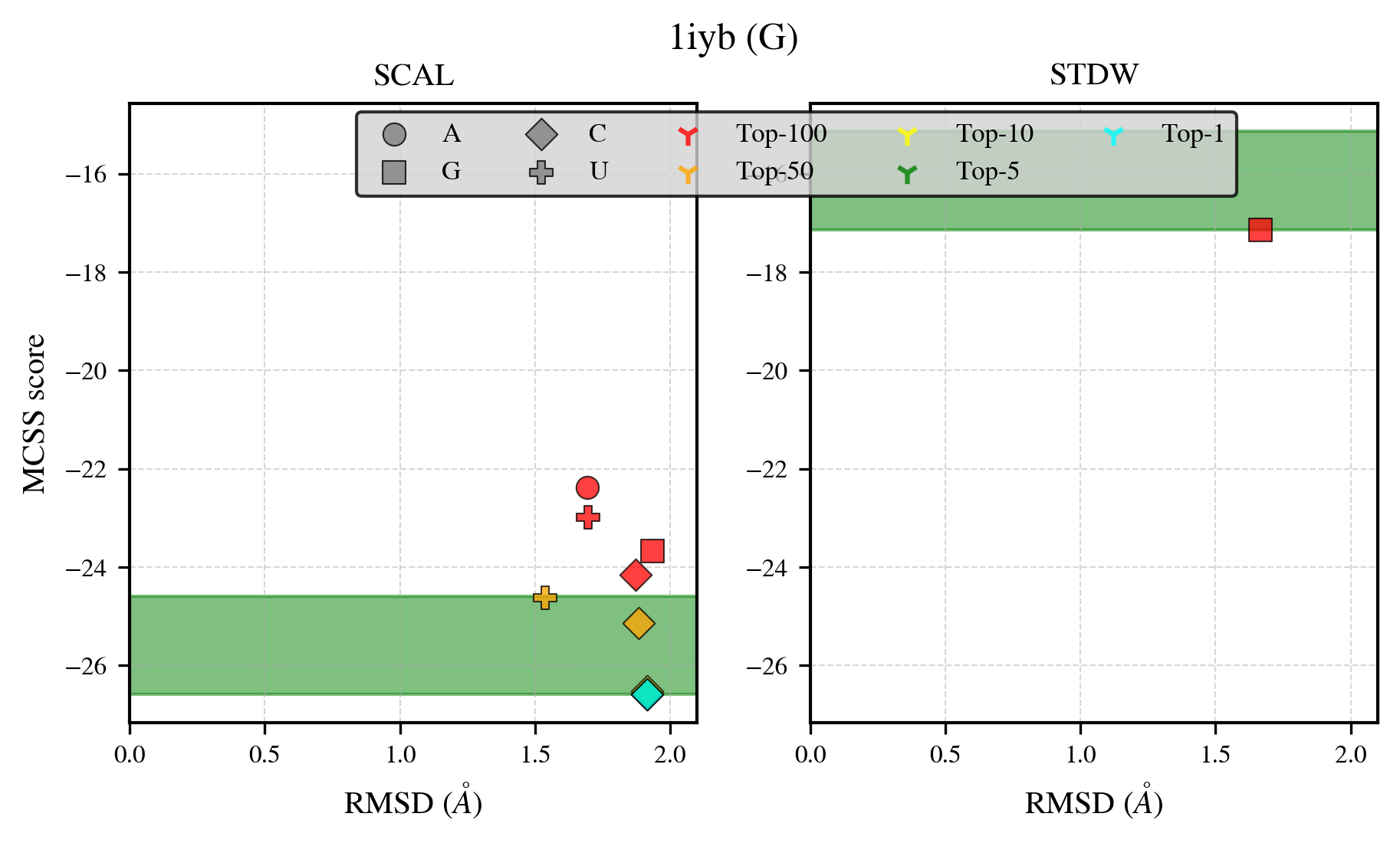

### 1jp4_benchmark-121_selectivity_details.png

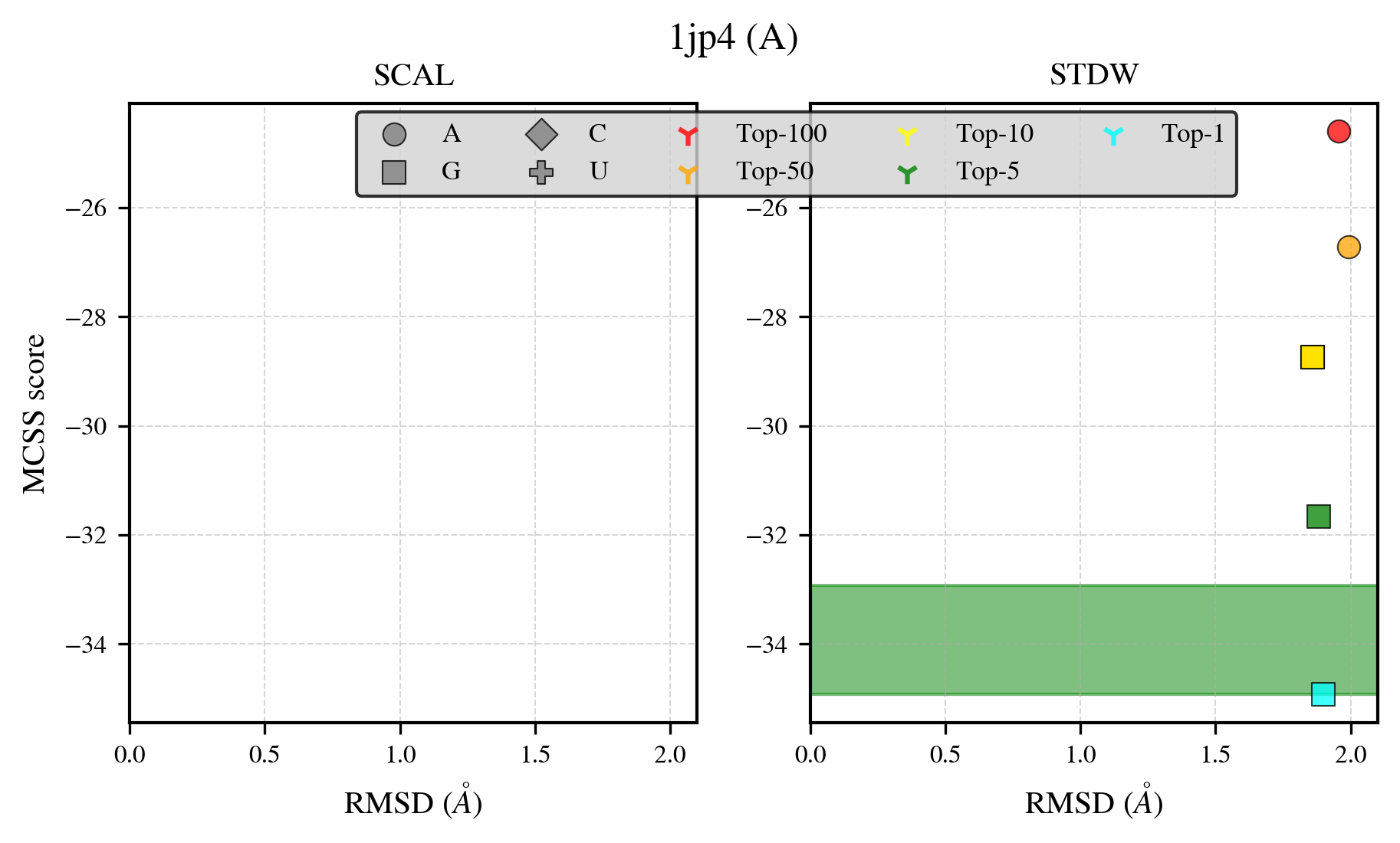

### 1ktg_benchmark-121_selectivity_details.png

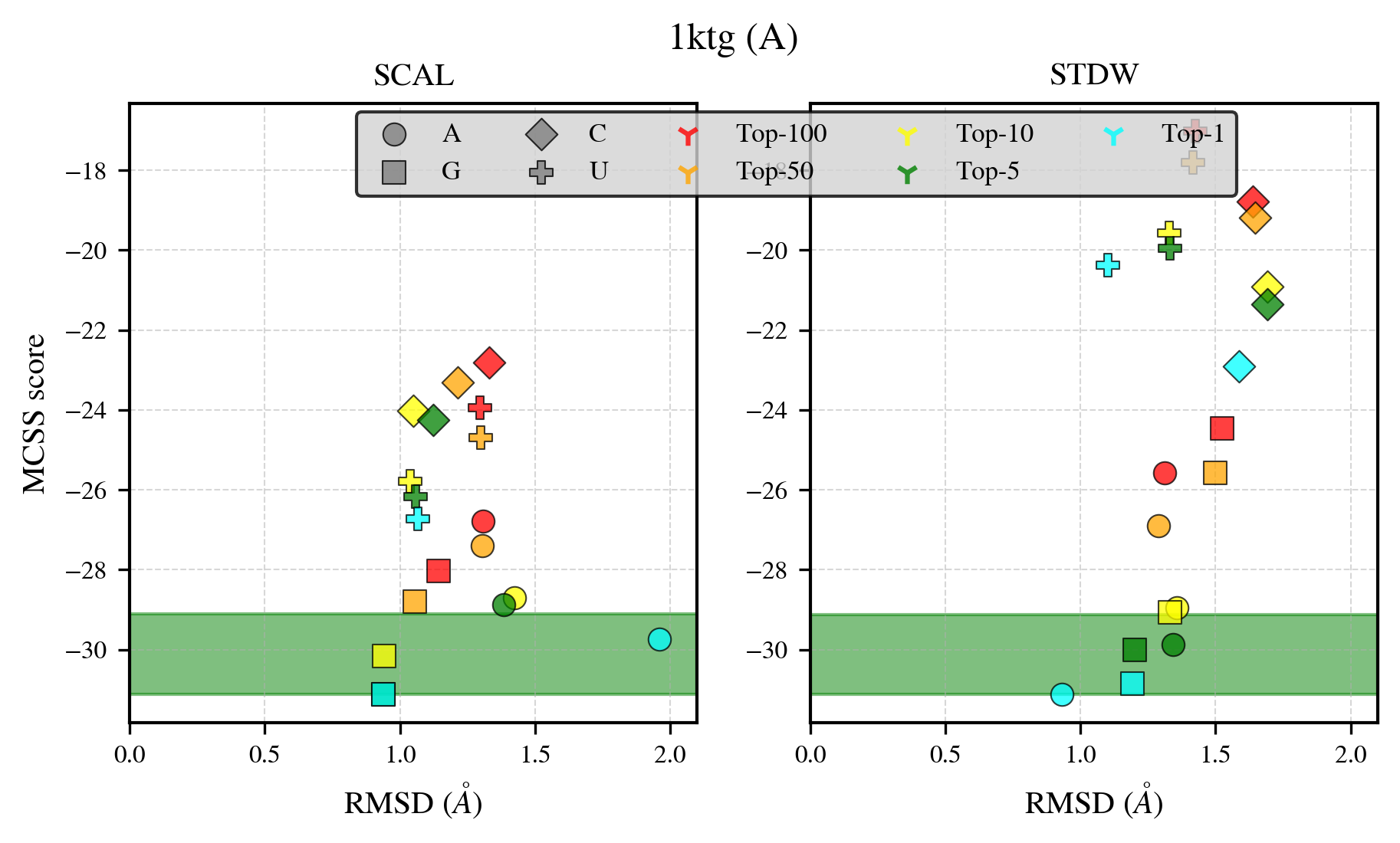

### 1nh8_benchmark-121_selectivity_details.png

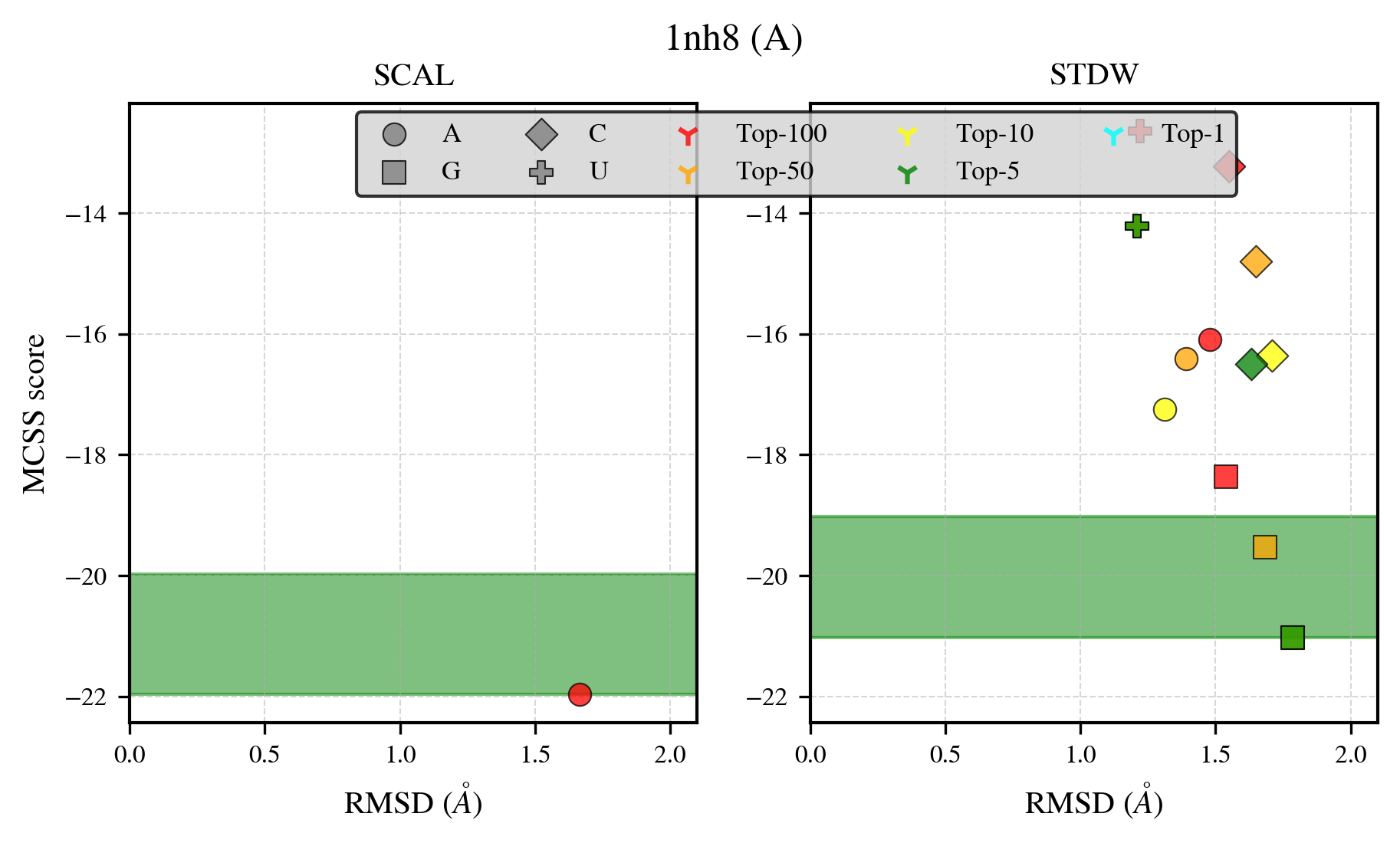

### 1qf9_benchmark-121_selectivity_details.png

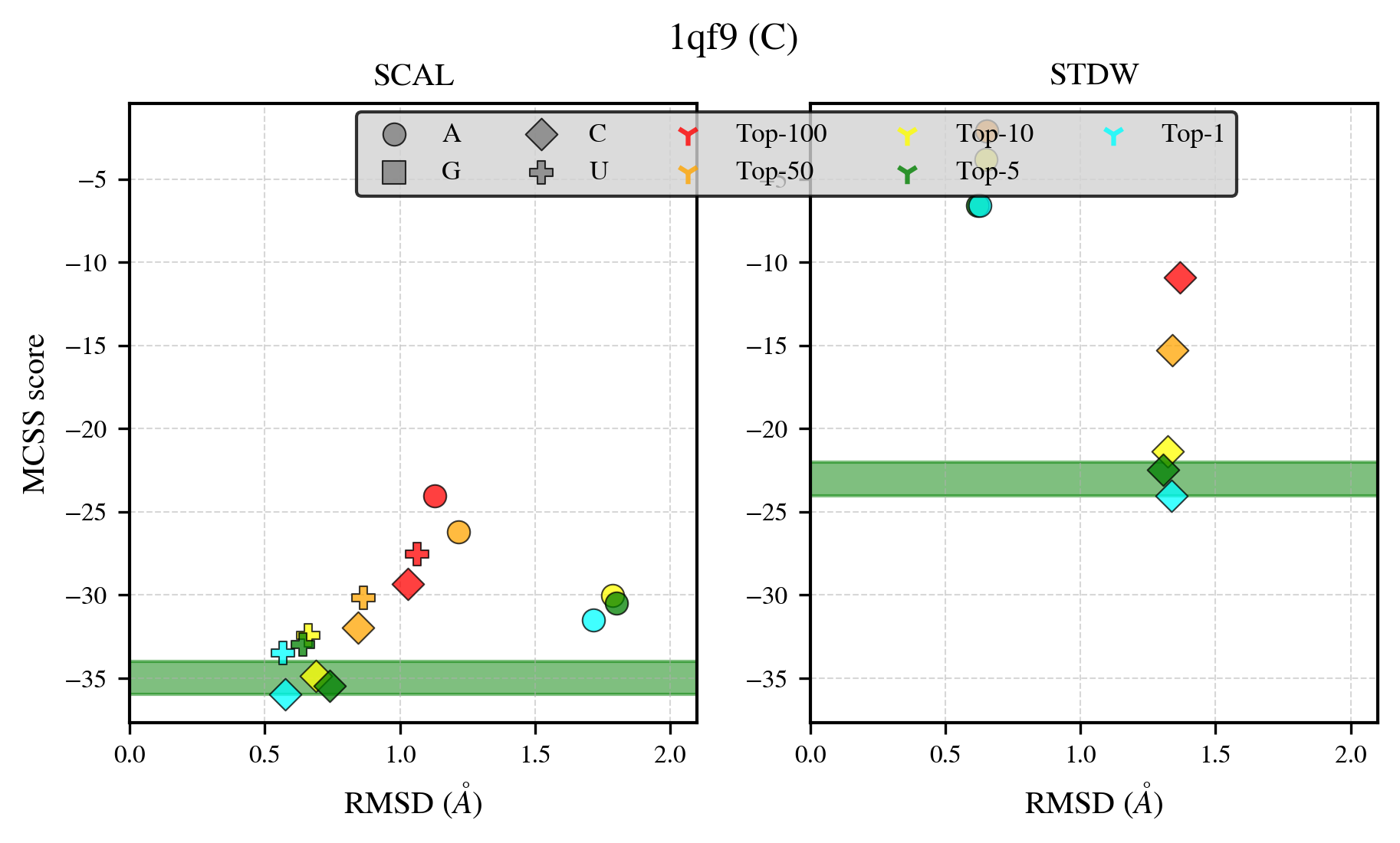

### 1qgx_benchmark-121_selectivity_details.png

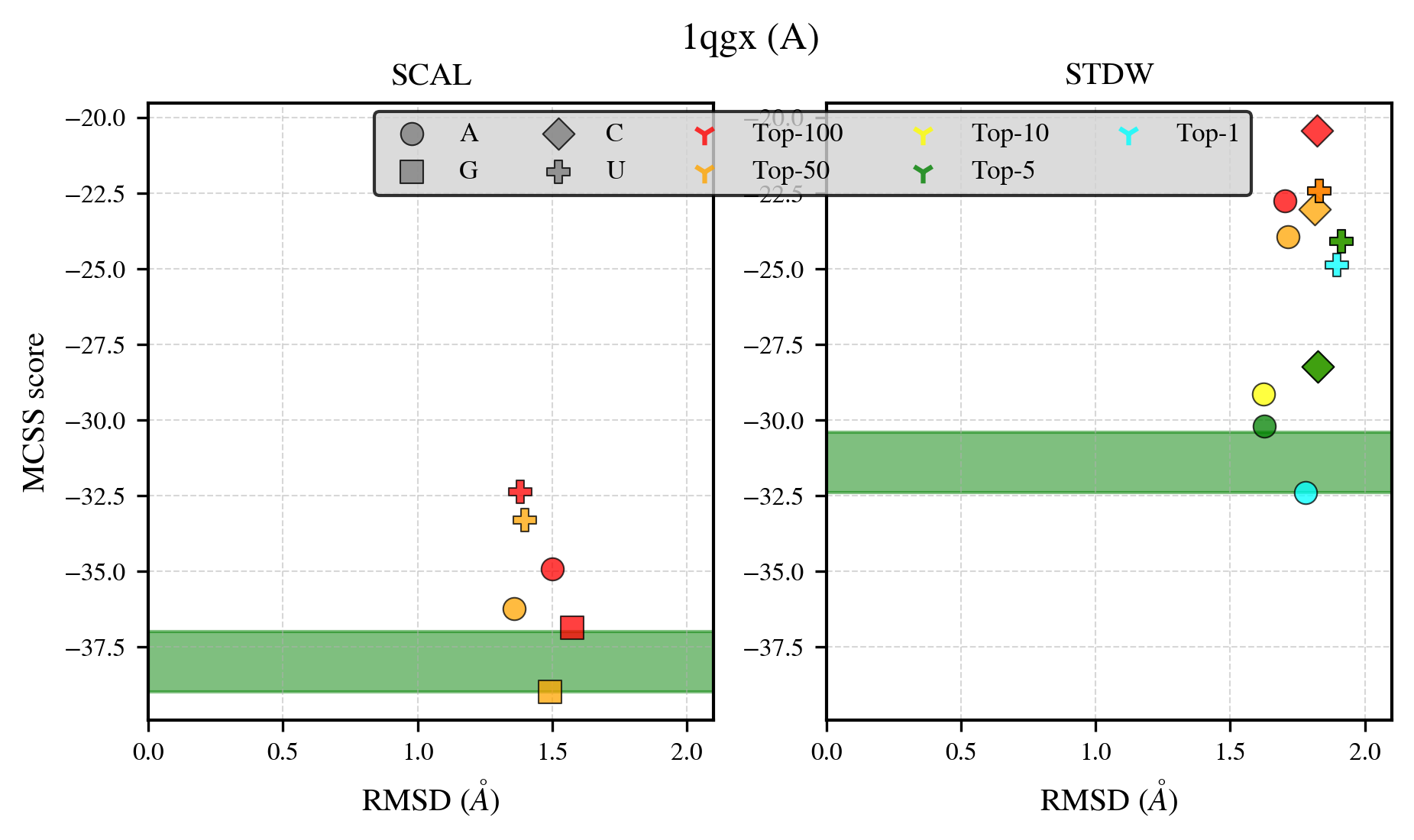

### 1rao_benchmark-121_selectivity_details.png

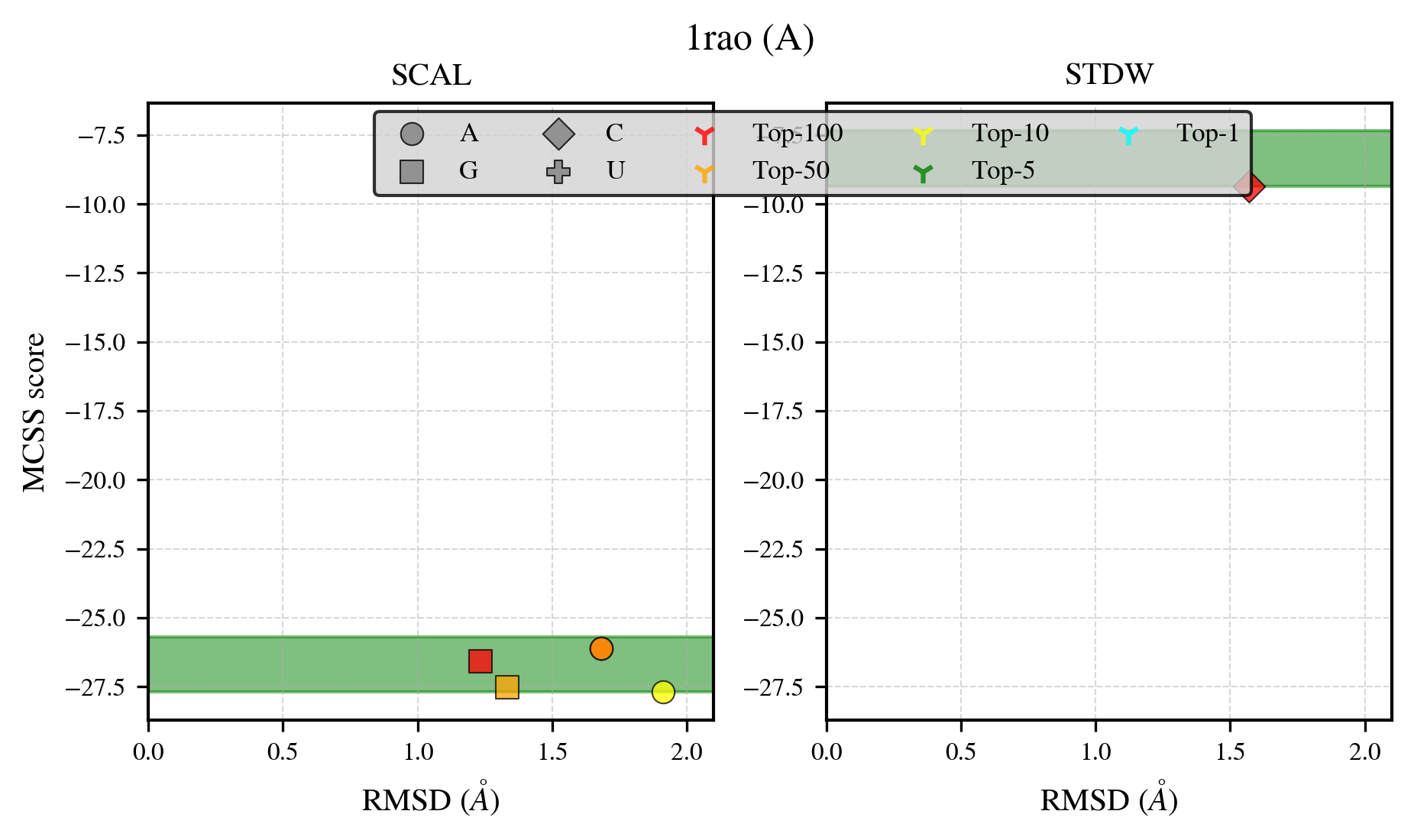

### 1s68_benchmark-121_selectivity_details.png

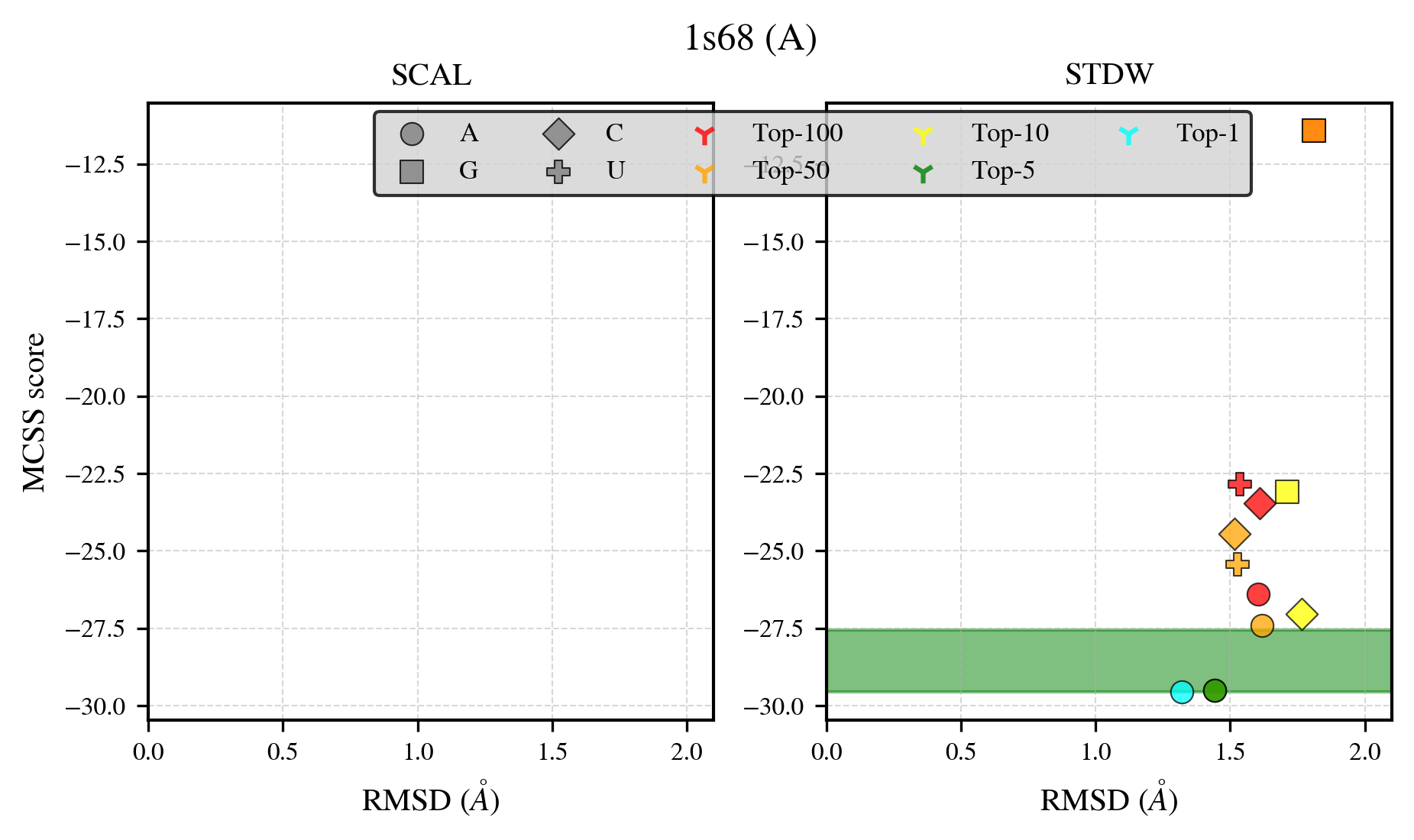

### 1ua4_benchmark-121_selectivity_details.png

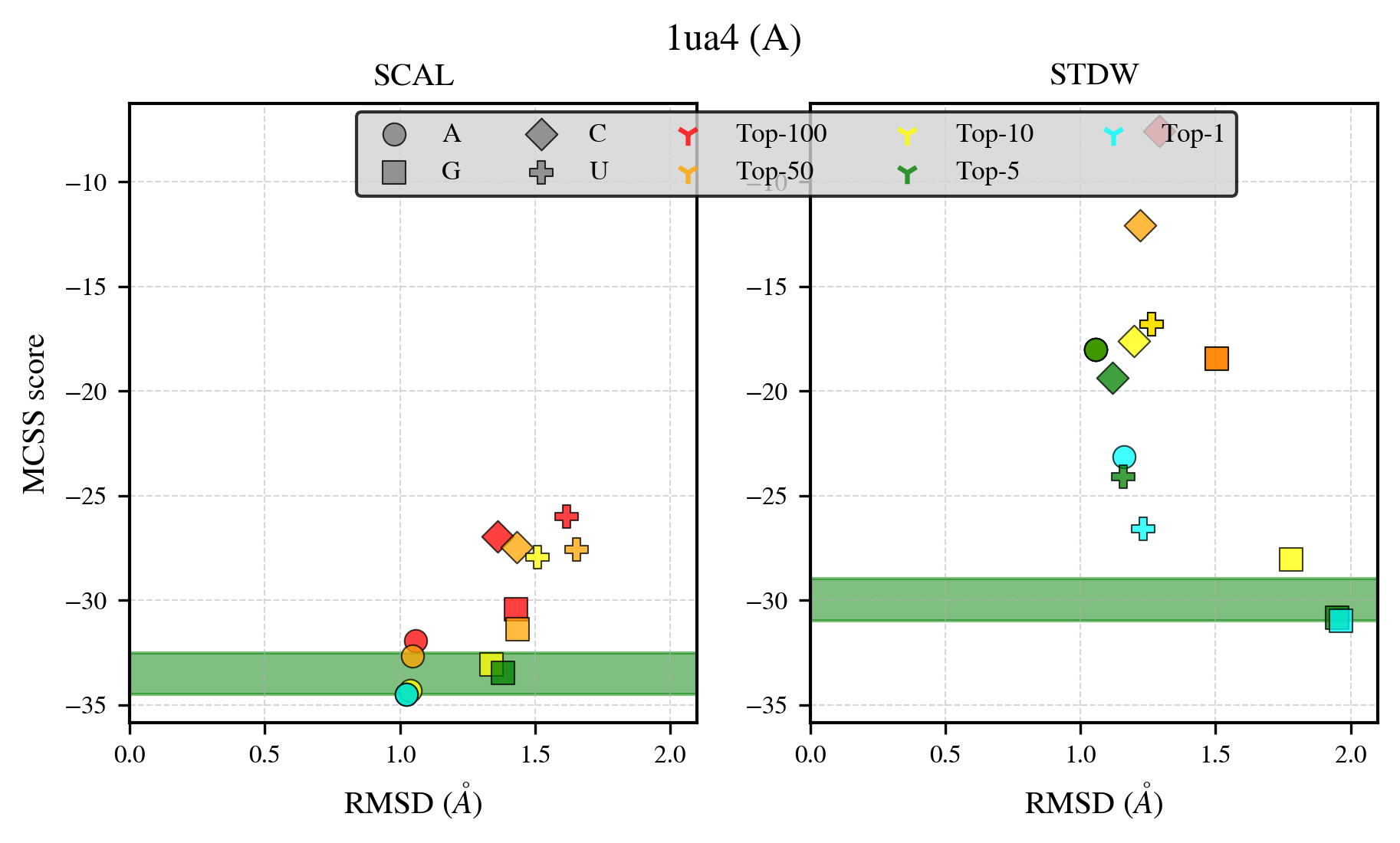

### 1ucd_benchmark-121_selectivity_details.png

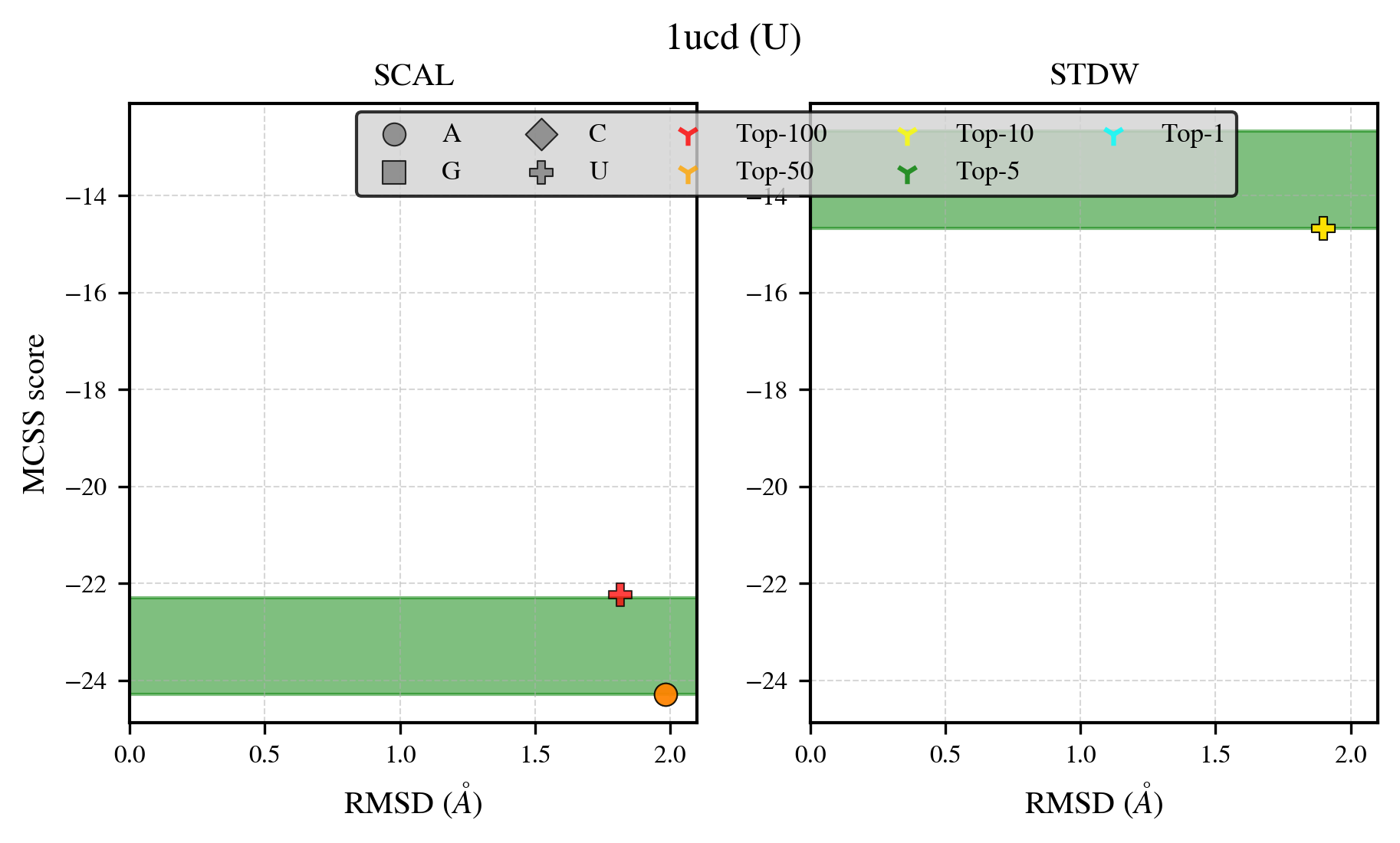

### 1uj2_benchmark-121_selectivity_details.png

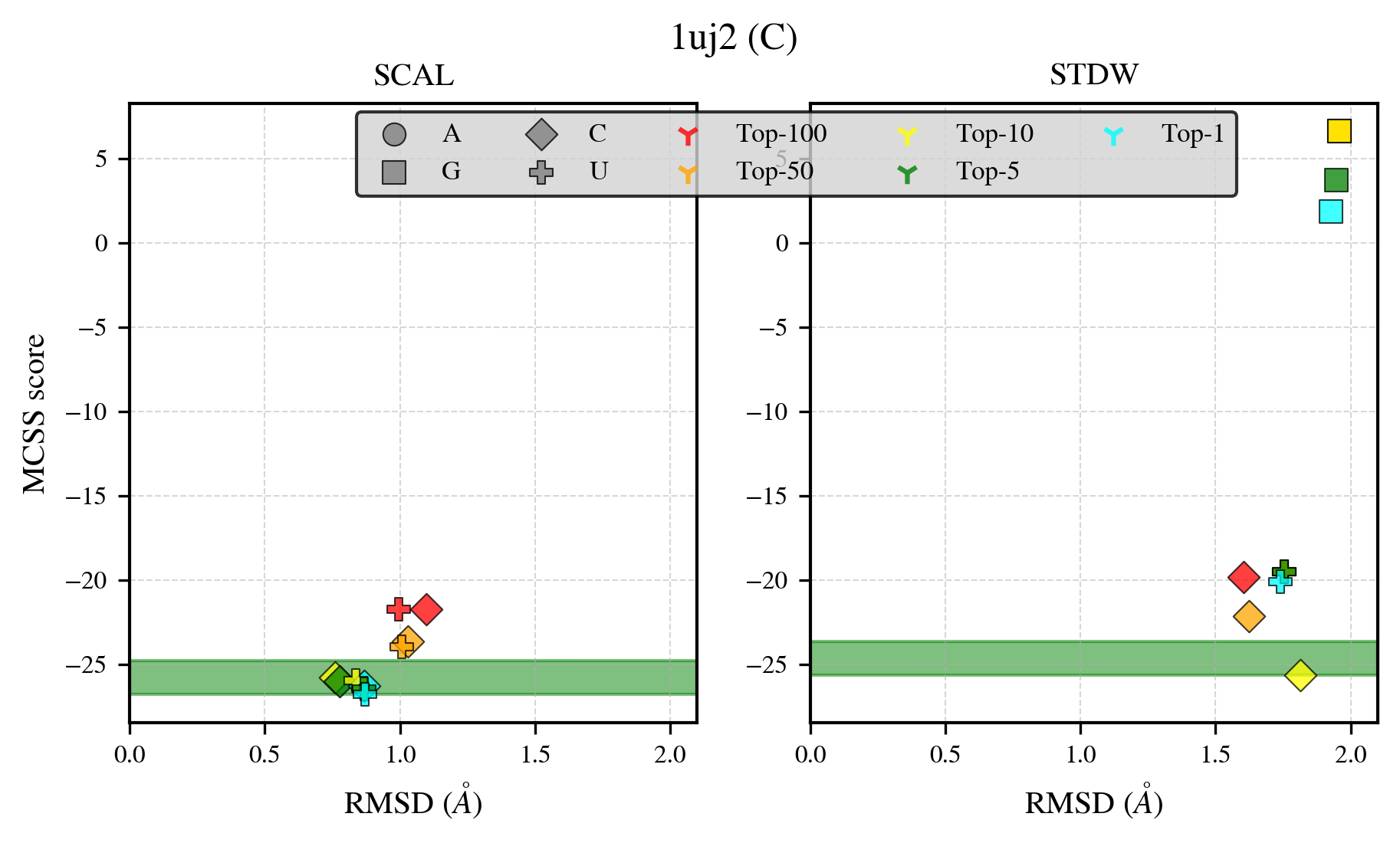

### 1uuy_benchmark-121_selectivity_details.png

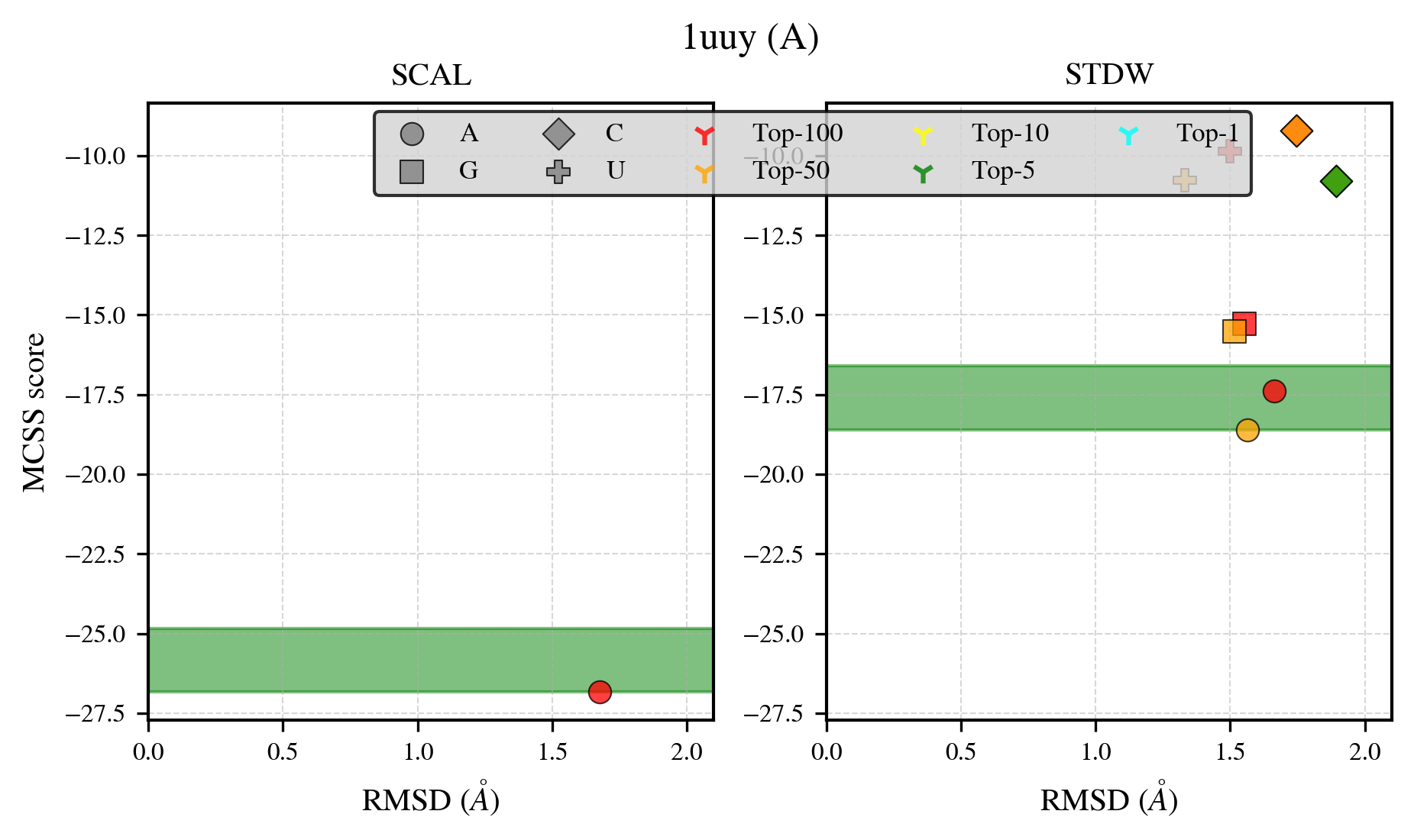

### 1xtt_benchmark-121_selectivity_details.png

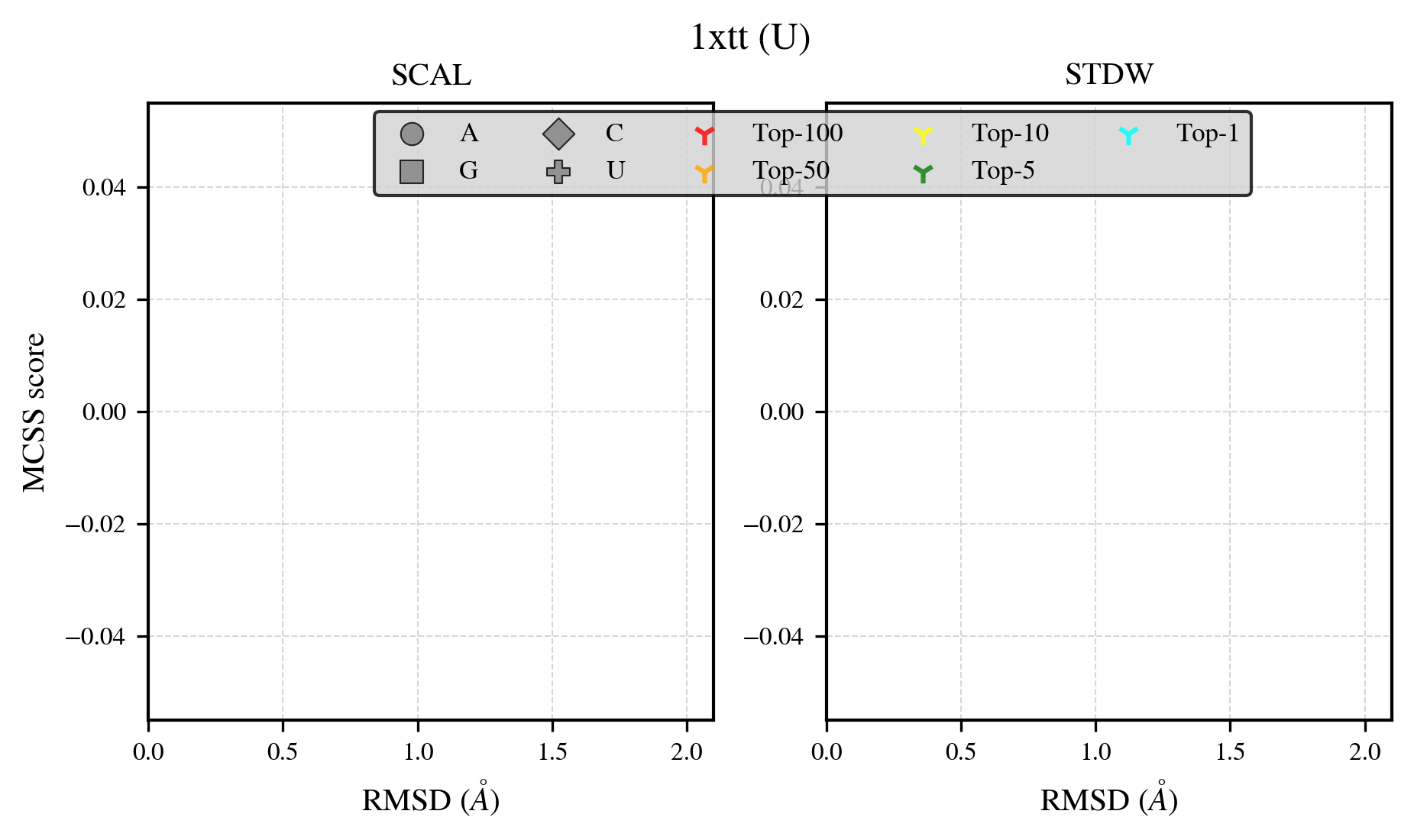

### 1y1p_benchmark-121_selectivity_details.png

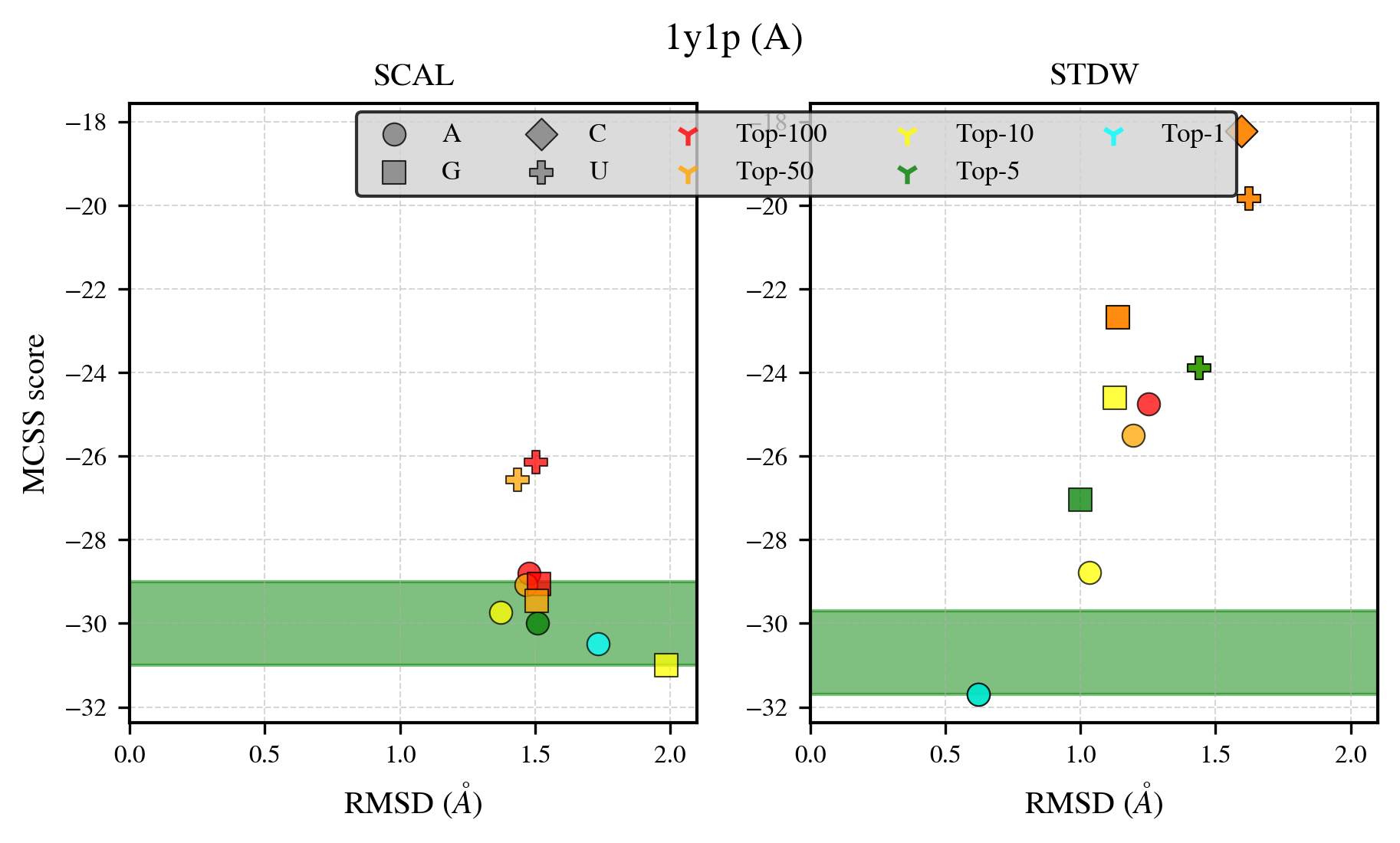

### 1z4m_benchmark-121_selectivity_details.png

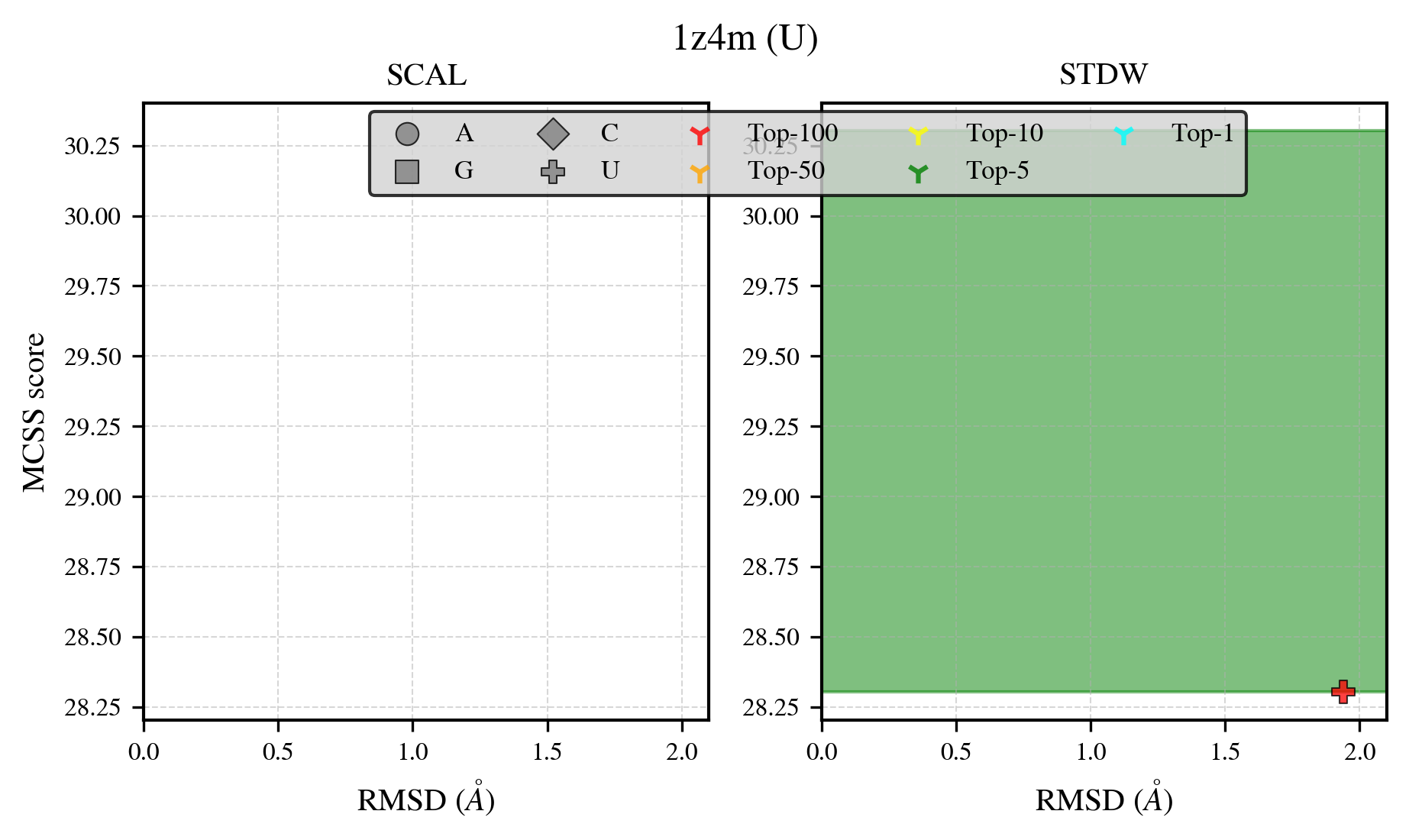

### 2a7x_benchmark-121_selectivity_details.png

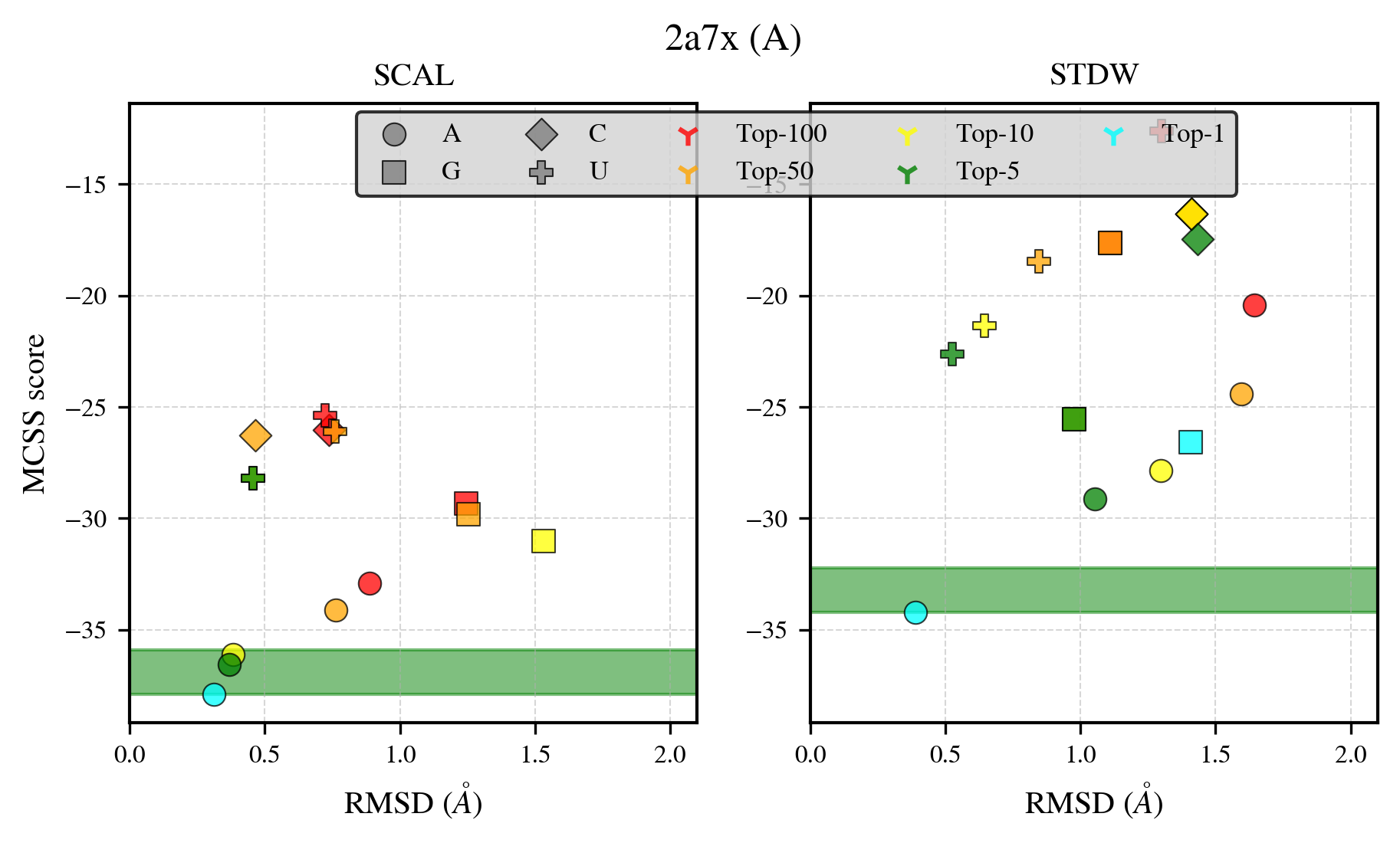

### 2cnq_benchmark-121_selectivity_details.png

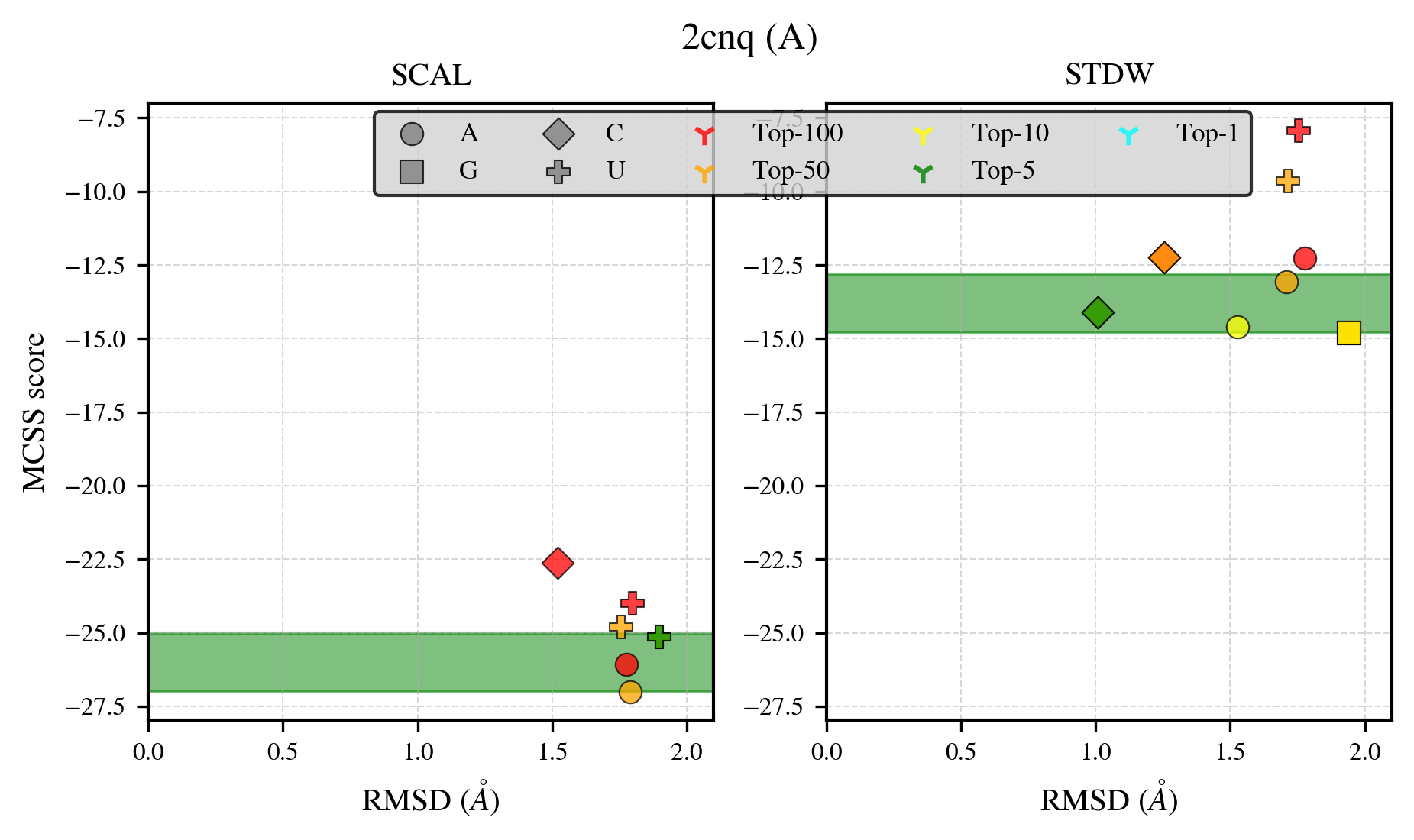

### 2eqa_benchmark-121_selectivity_details.png

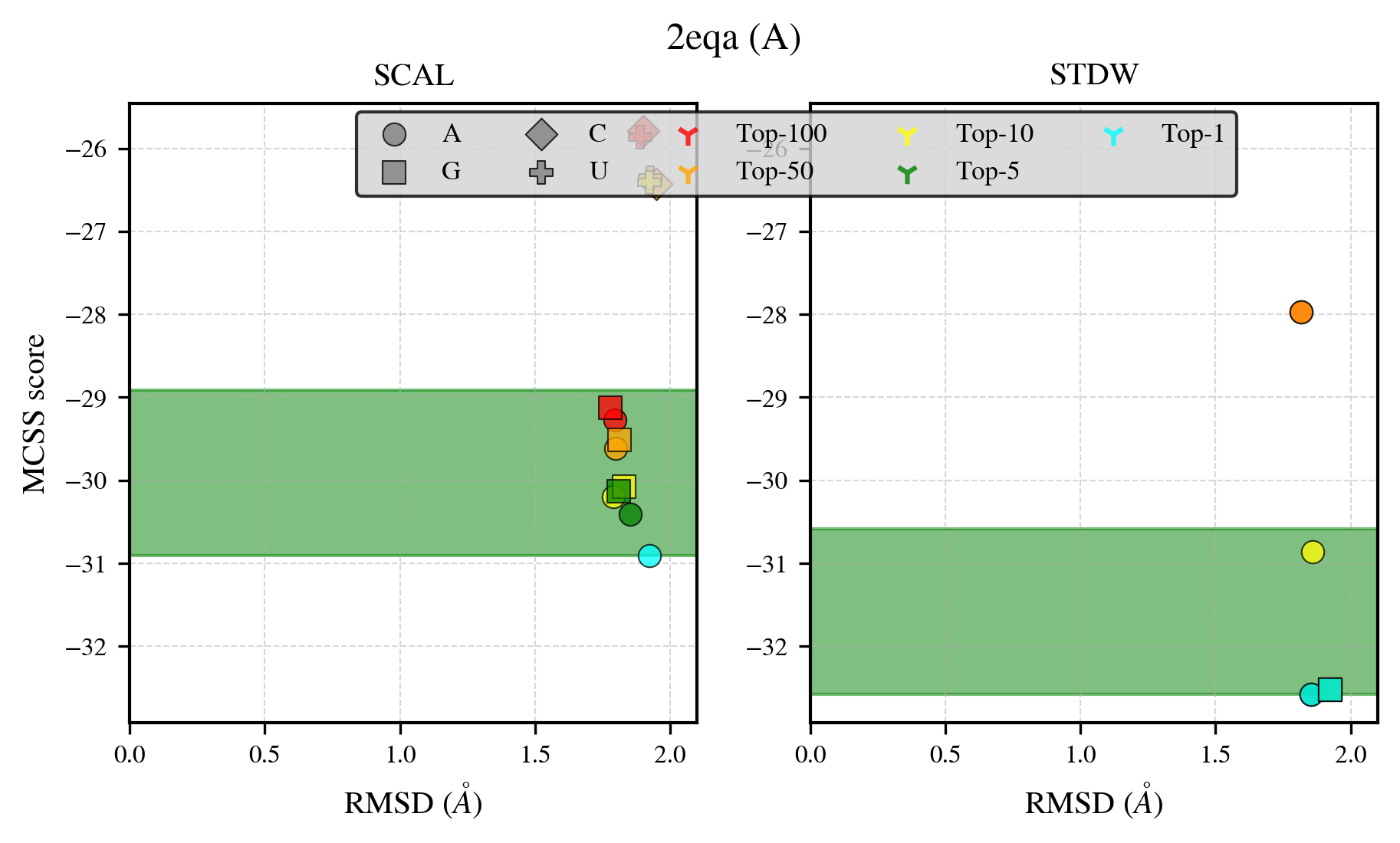

### 2ffc_benchmark-121_selectivity_details.png

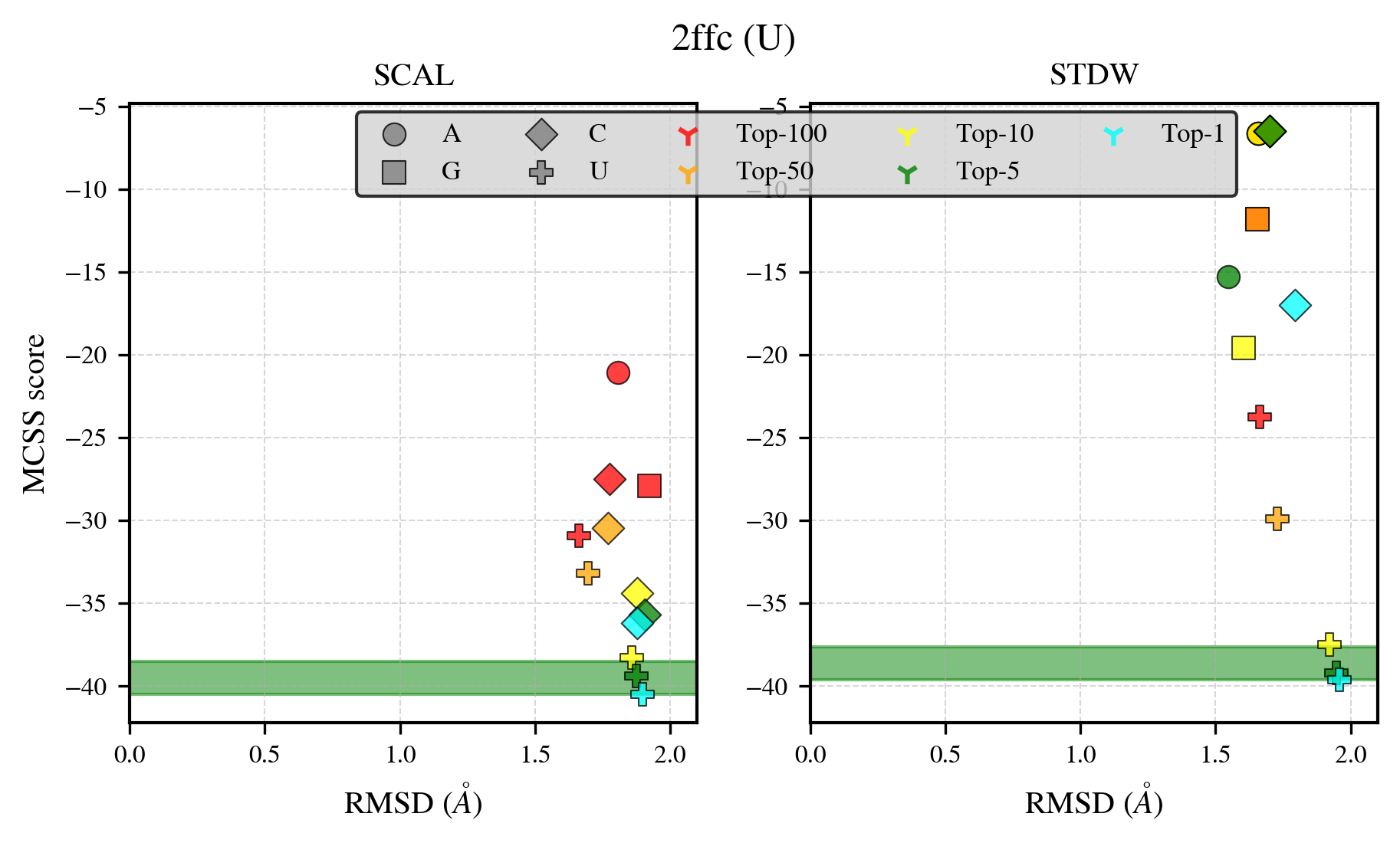

### 2fjb_benchmark-121_selectivity_details.png

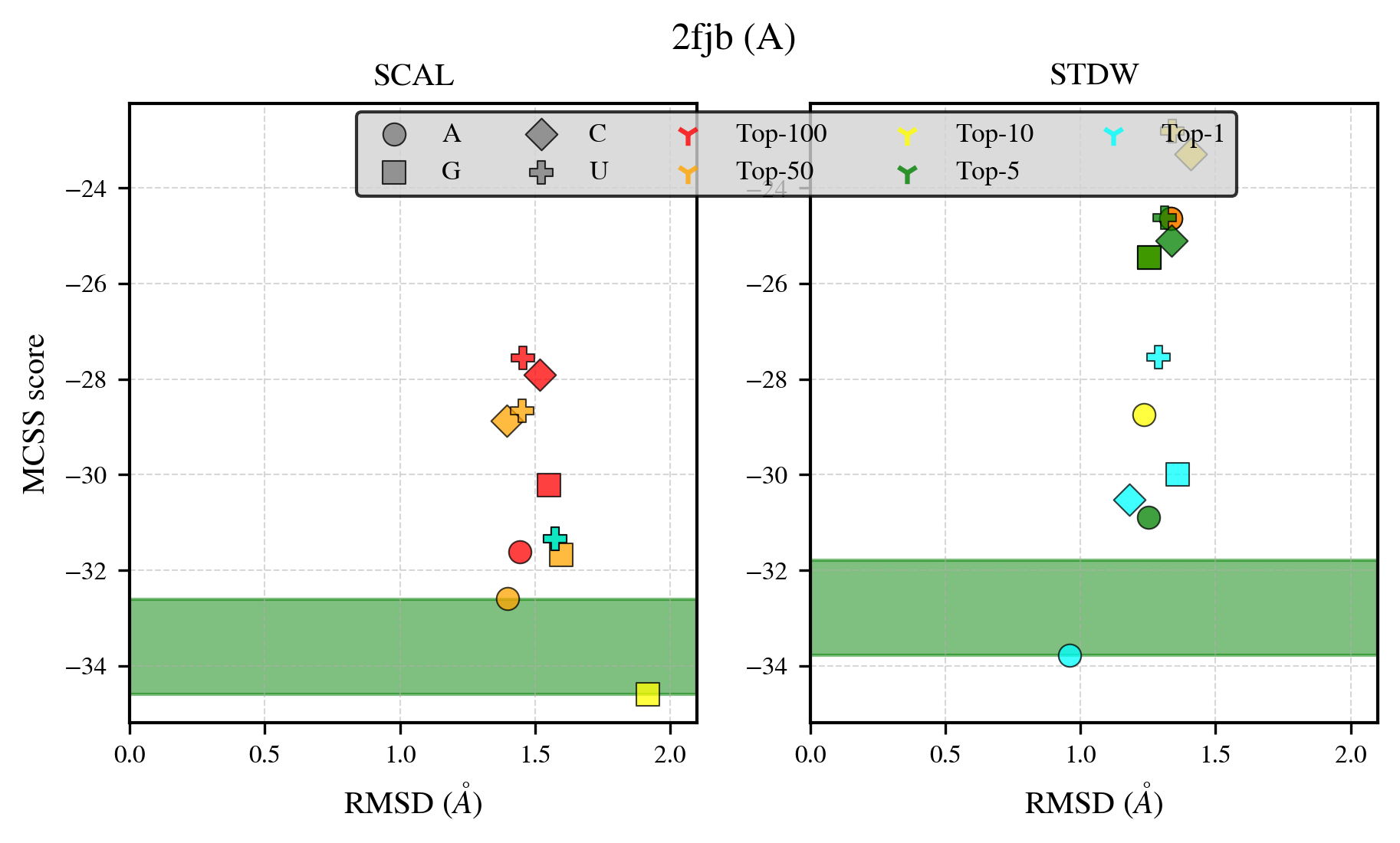

### 2gxq_benchmark-121_selectivity_details.png

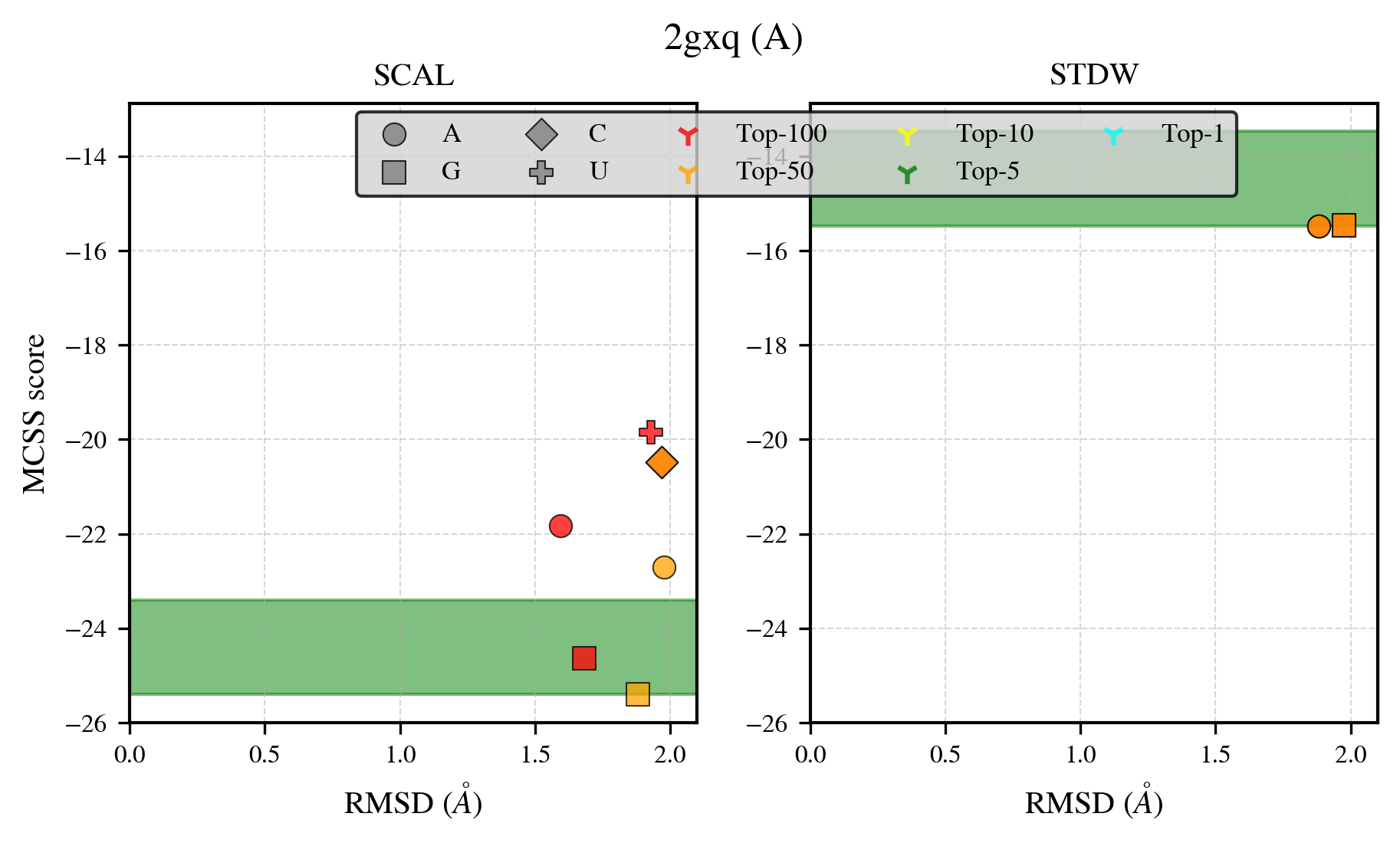

### 2ii6_benchmark-121_selectivity_details.png

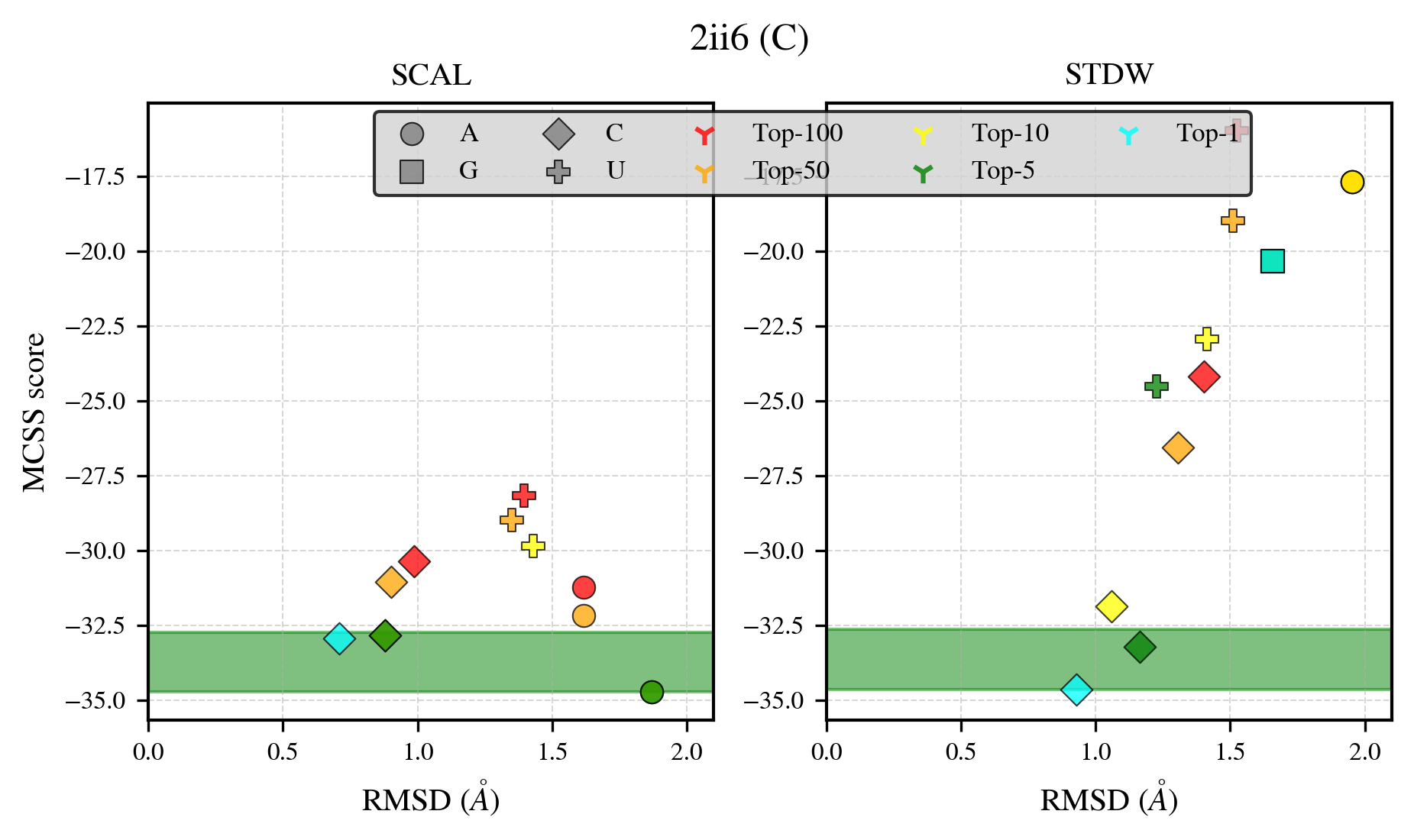

### 2j91_benchmark-121_selectivity_details.png

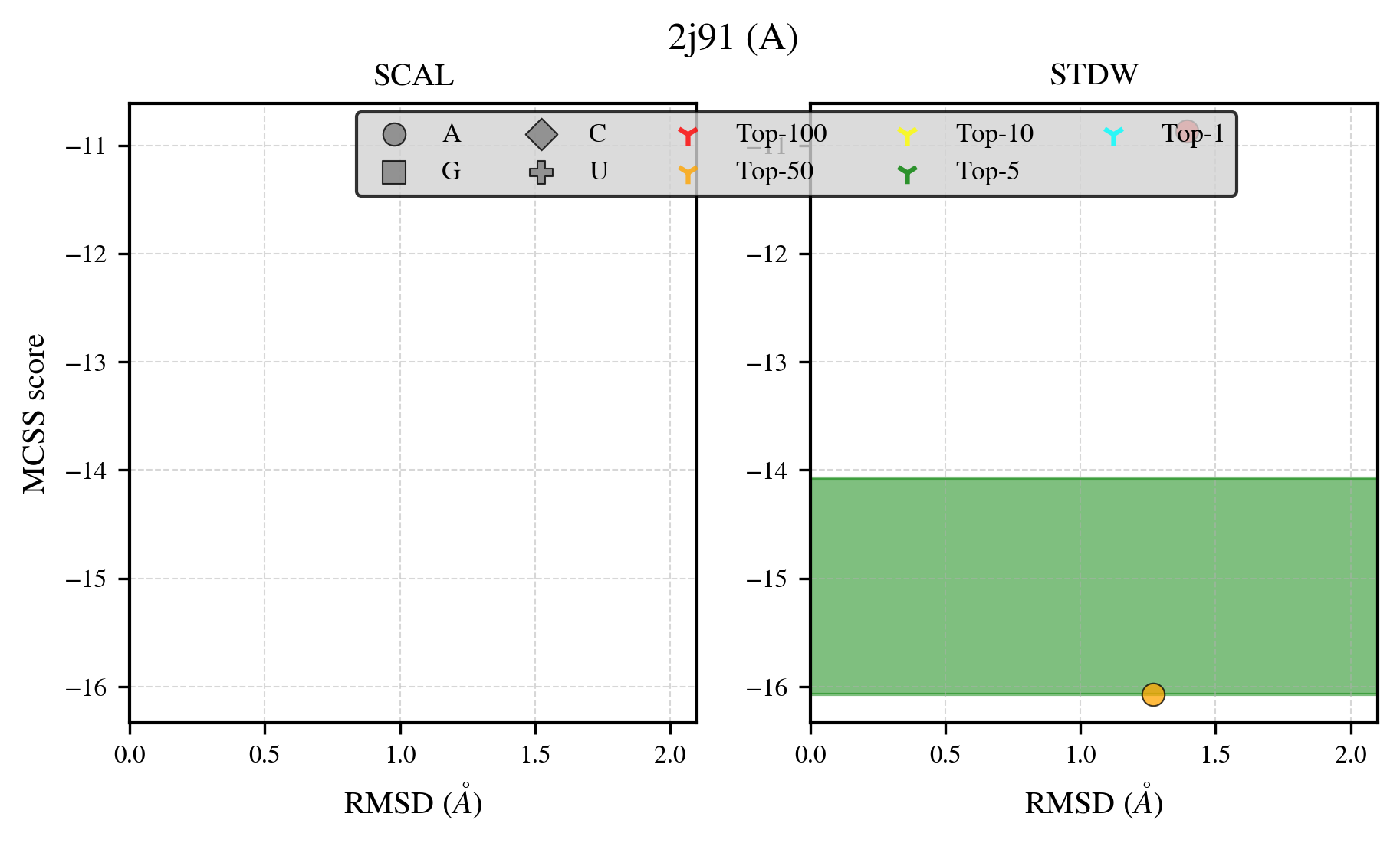

### 2jbh_benchmark-121_selectivity_details.png

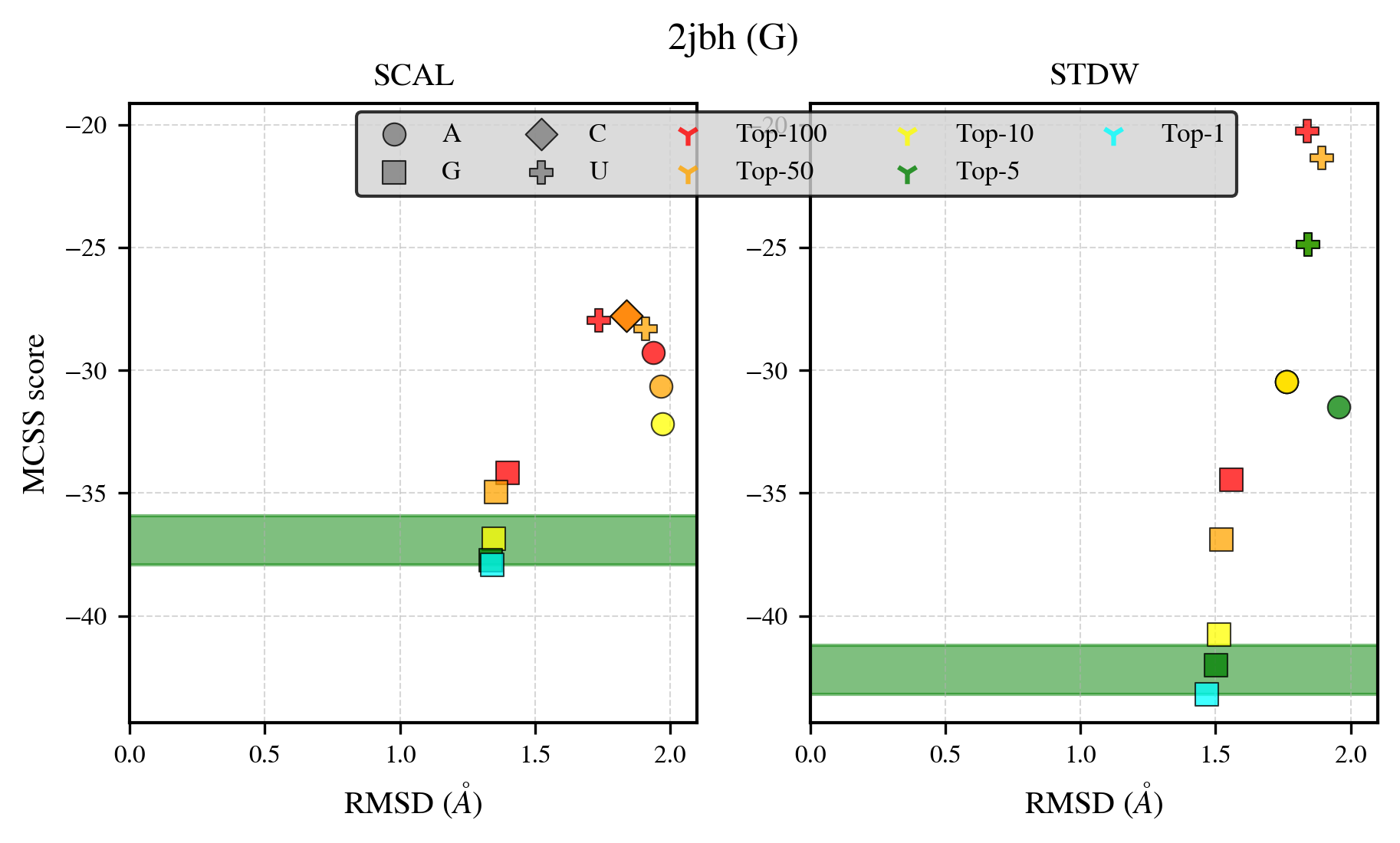

### 2oun_benchmark-121_selectivity_details.png

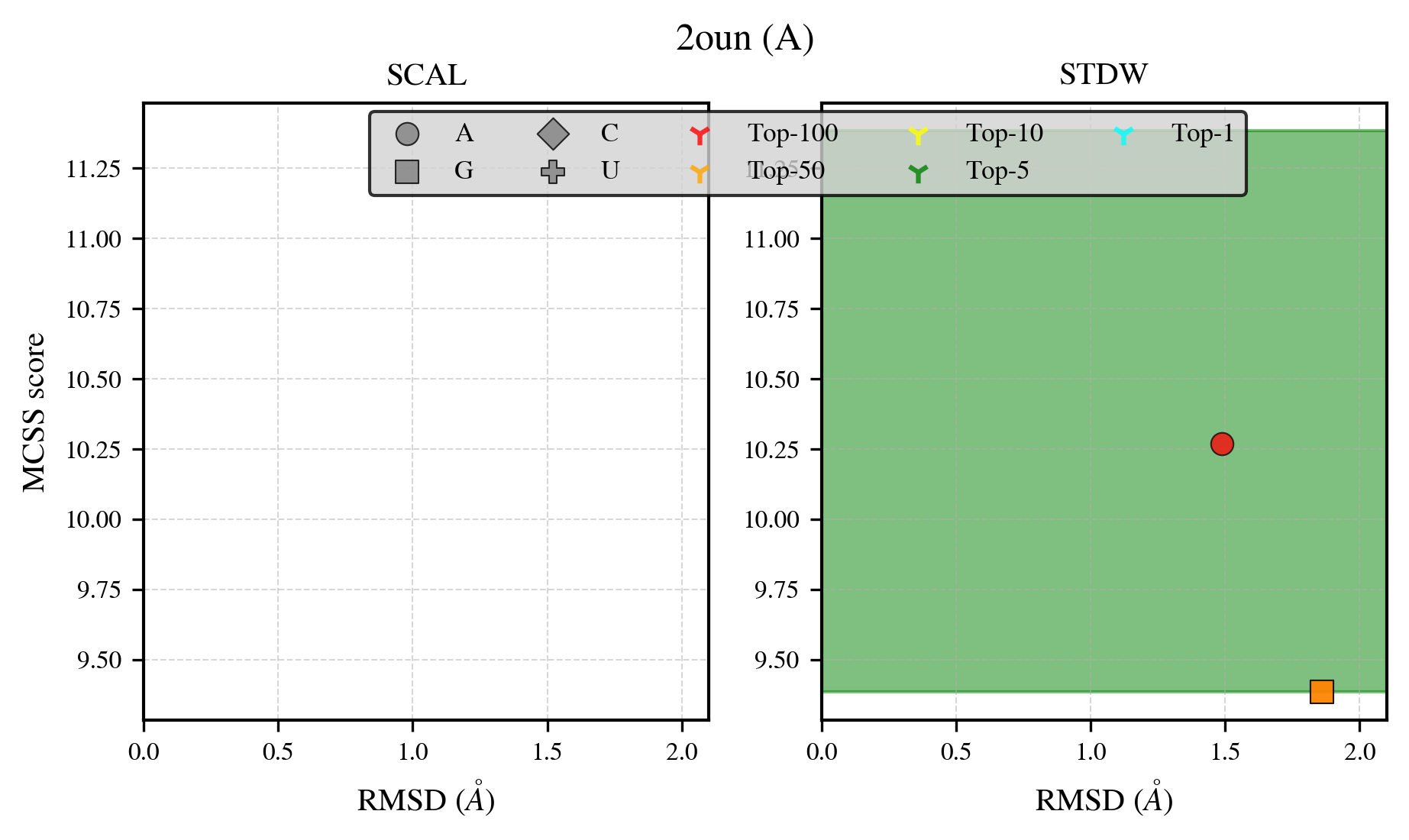

### 2qrk_benchmark-121_selectivity_details.png

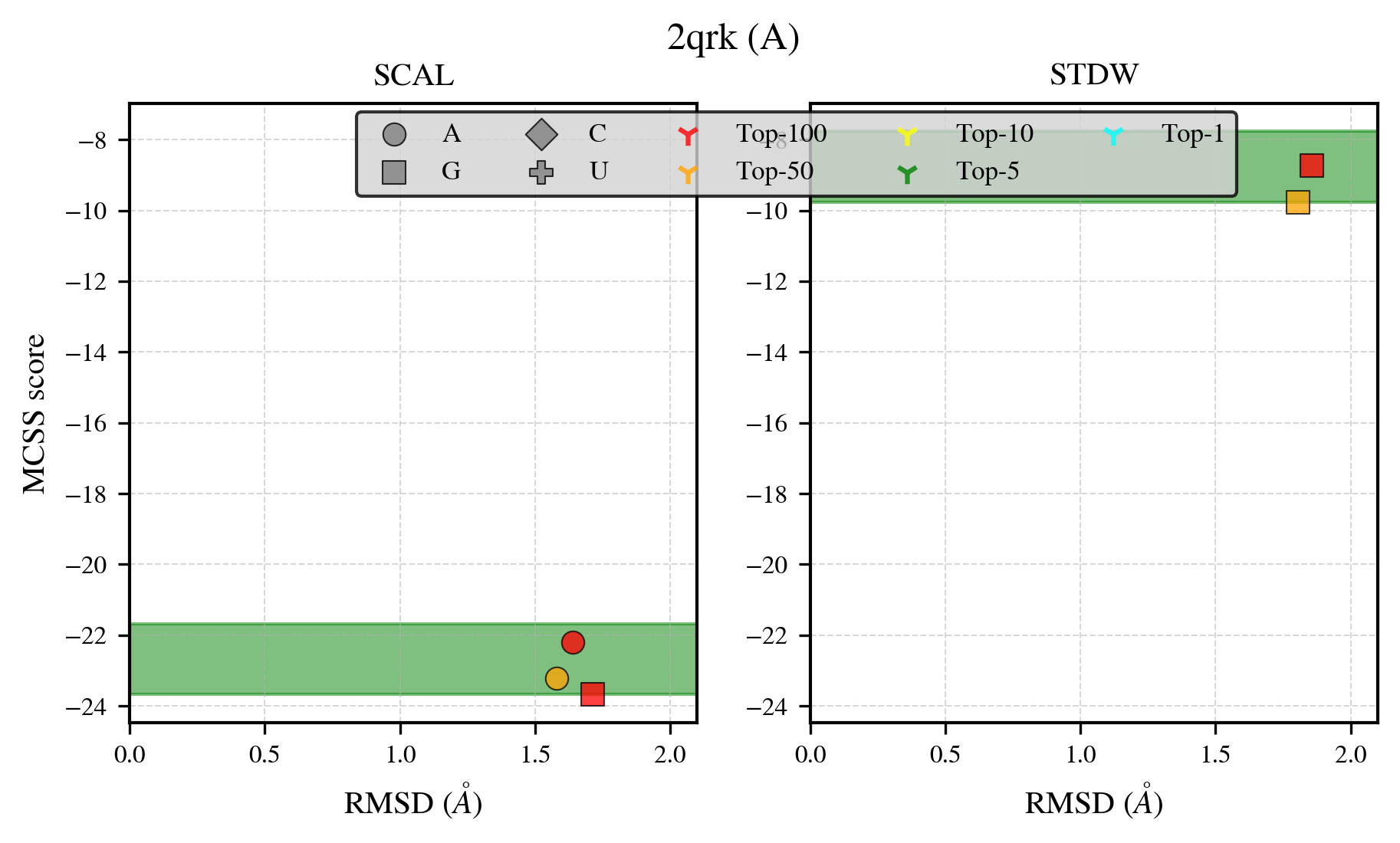

### 2r85_benchmark-121_selectivity_details.png

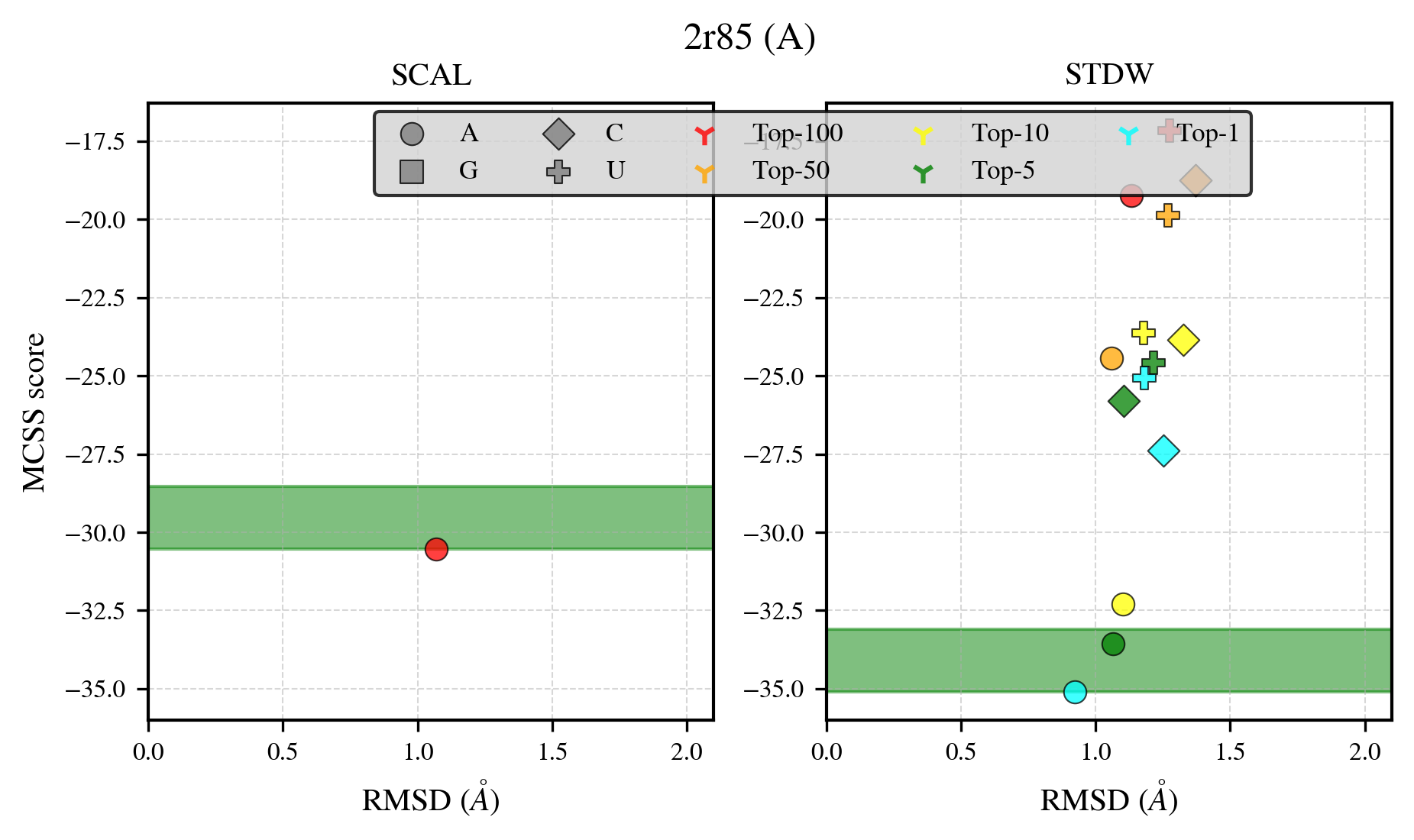

### 2uv4_benchmark-121_selectivity_details.png

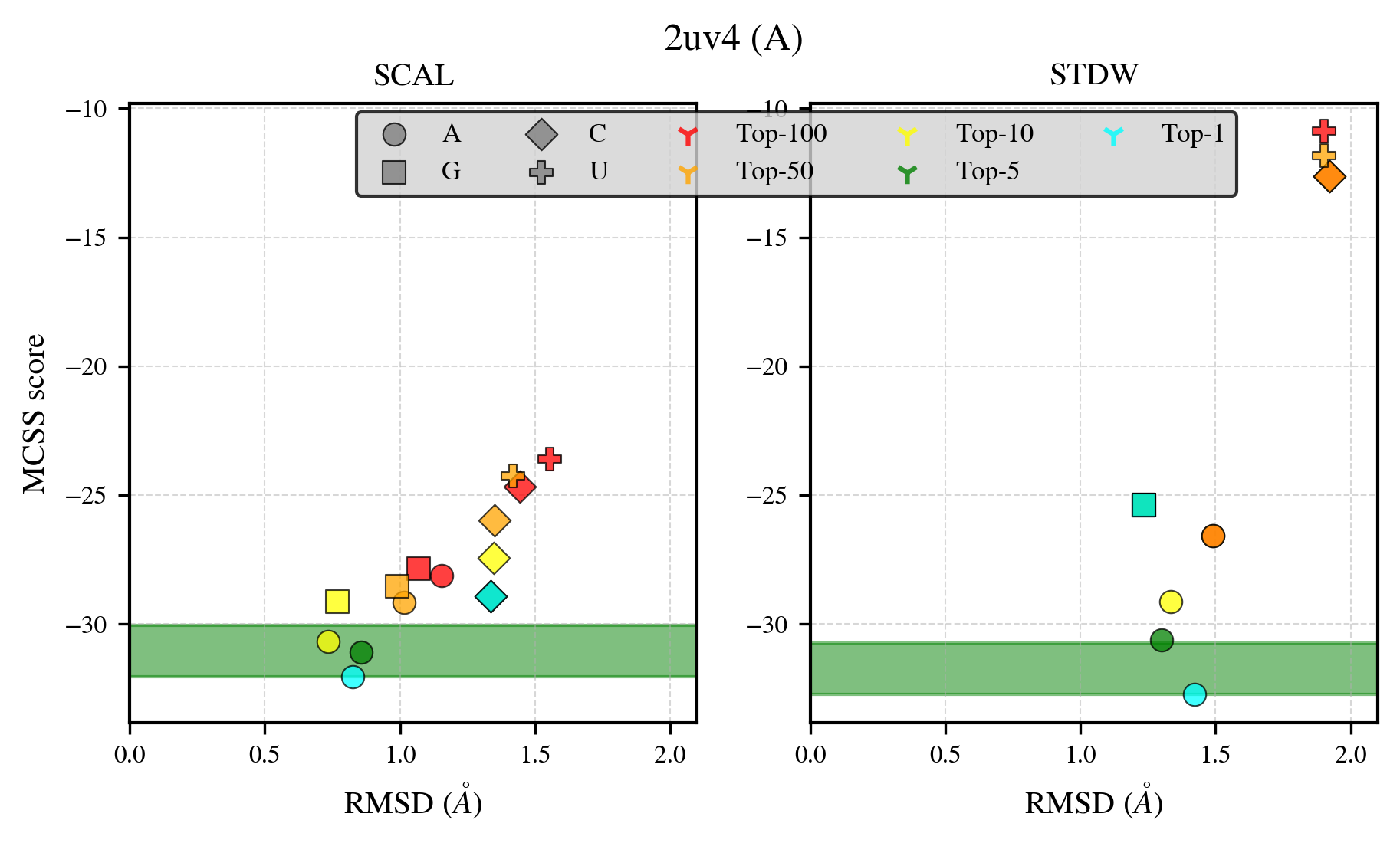

### 2xbu_benchmark-121_selectivity_details.png

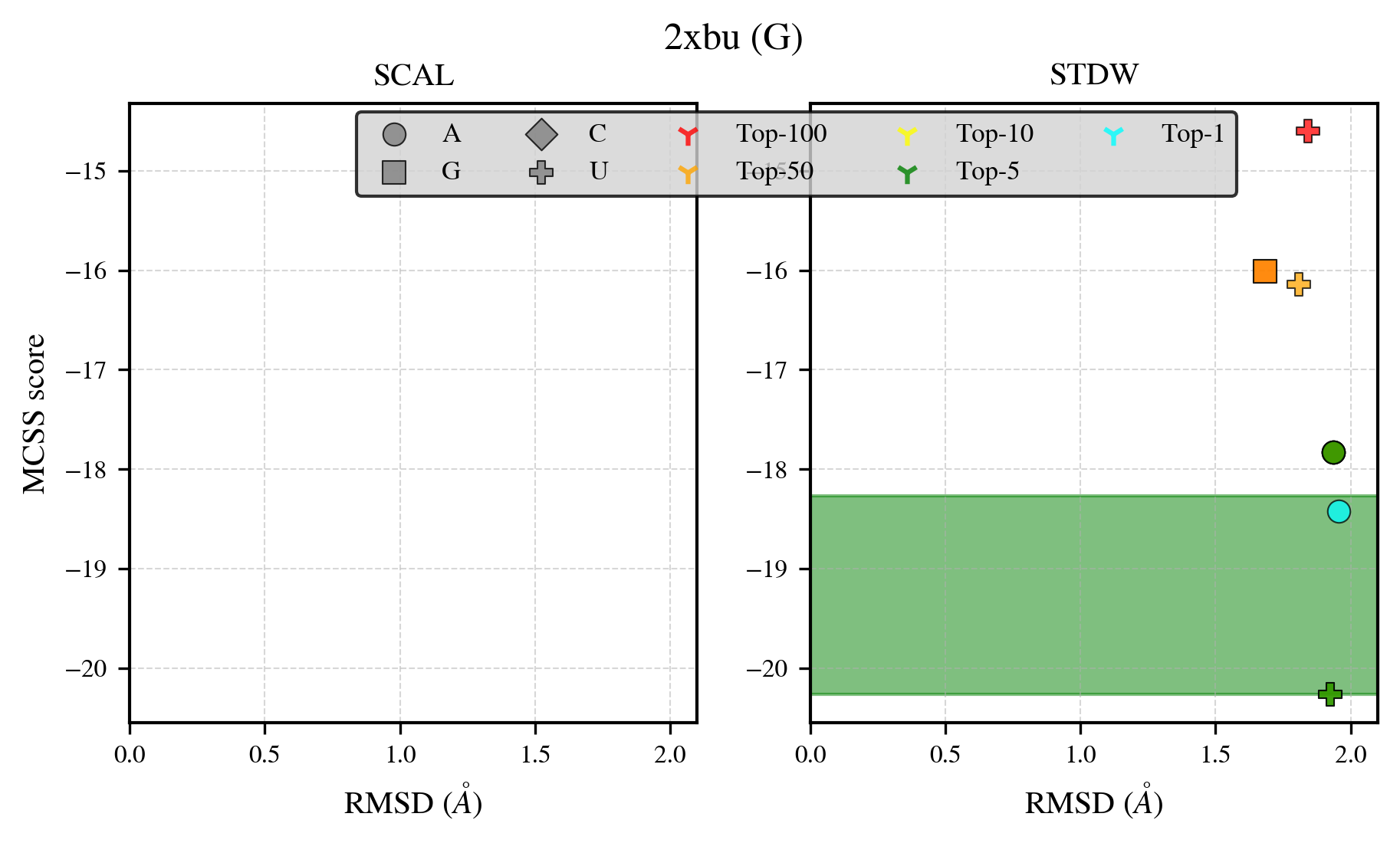

### 2yab_benchmark-121_selectivity_details.png

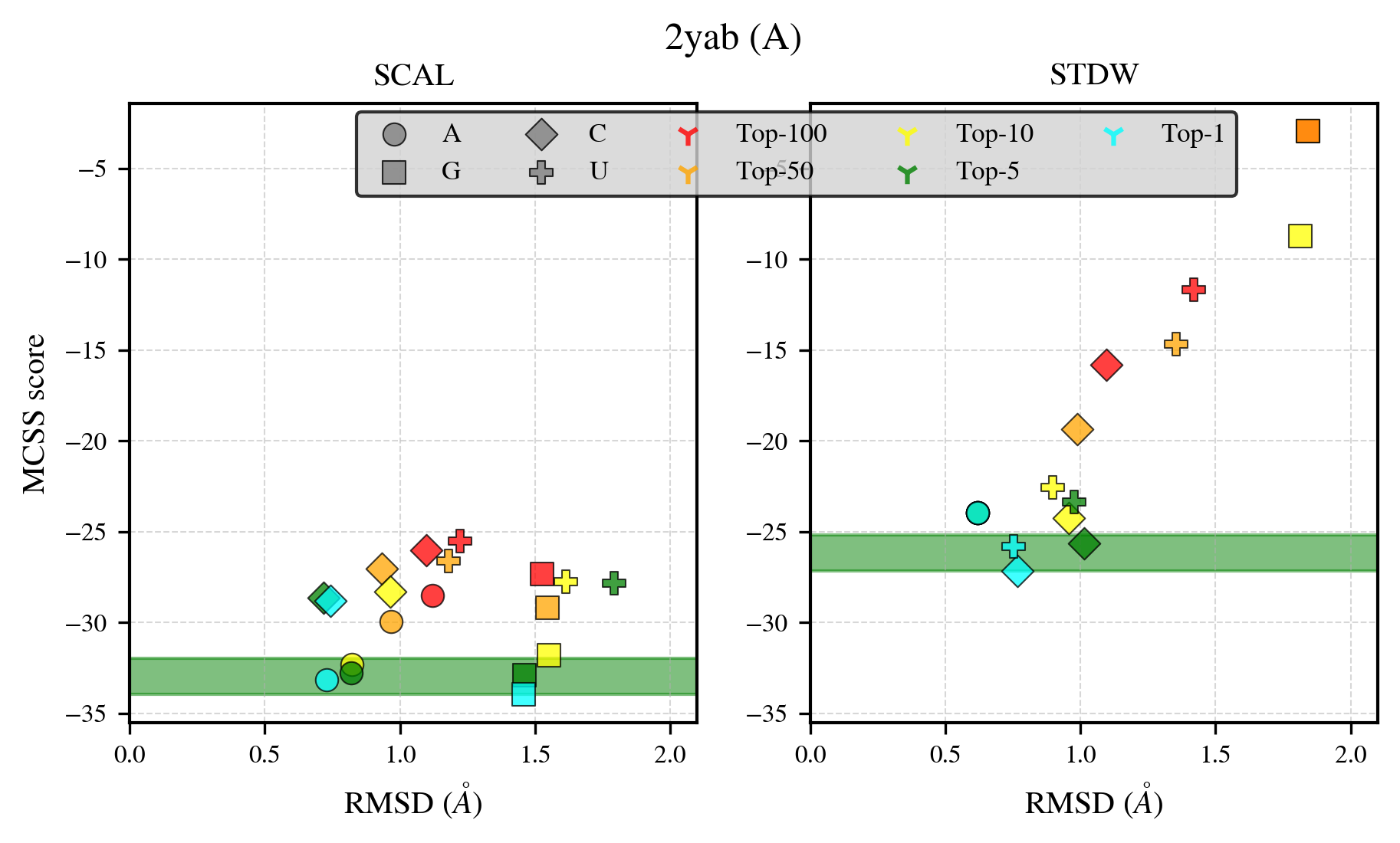

### 2yrx_benchmark-121_selectivity_details.png

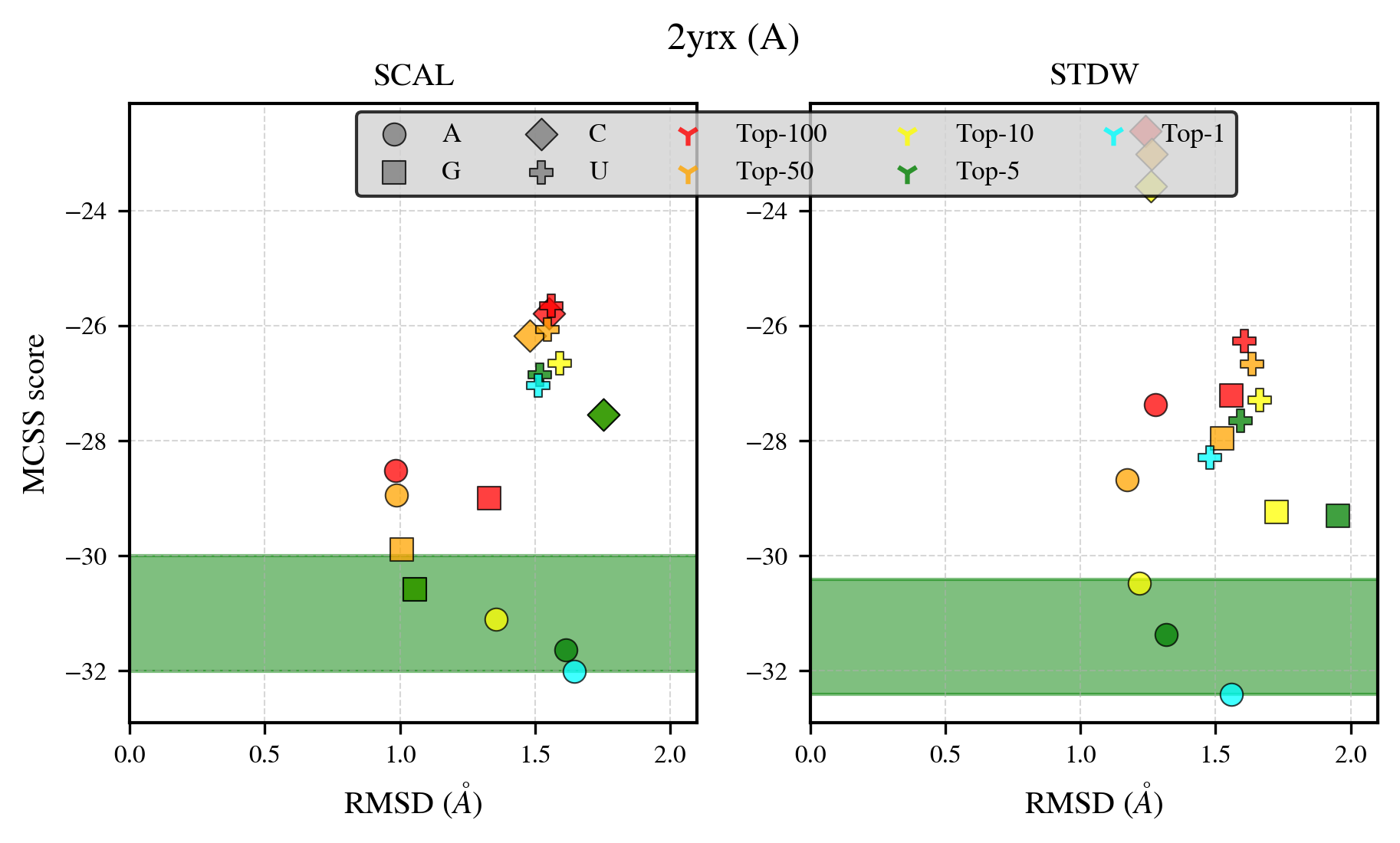

### 2yvo_benchmark-121_selectivity_details.png

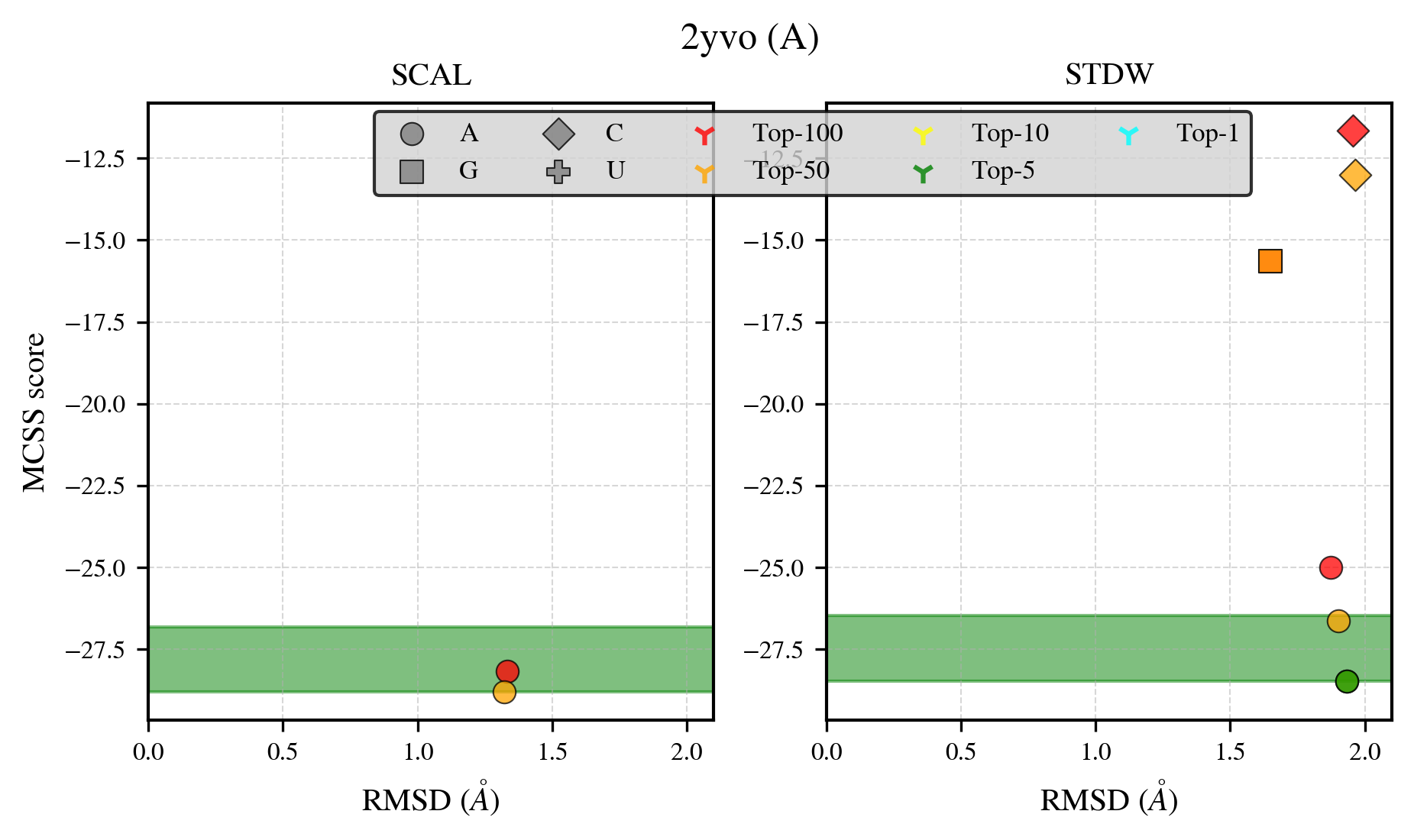

### 3ake_benchmark-121_selectivity_details.png

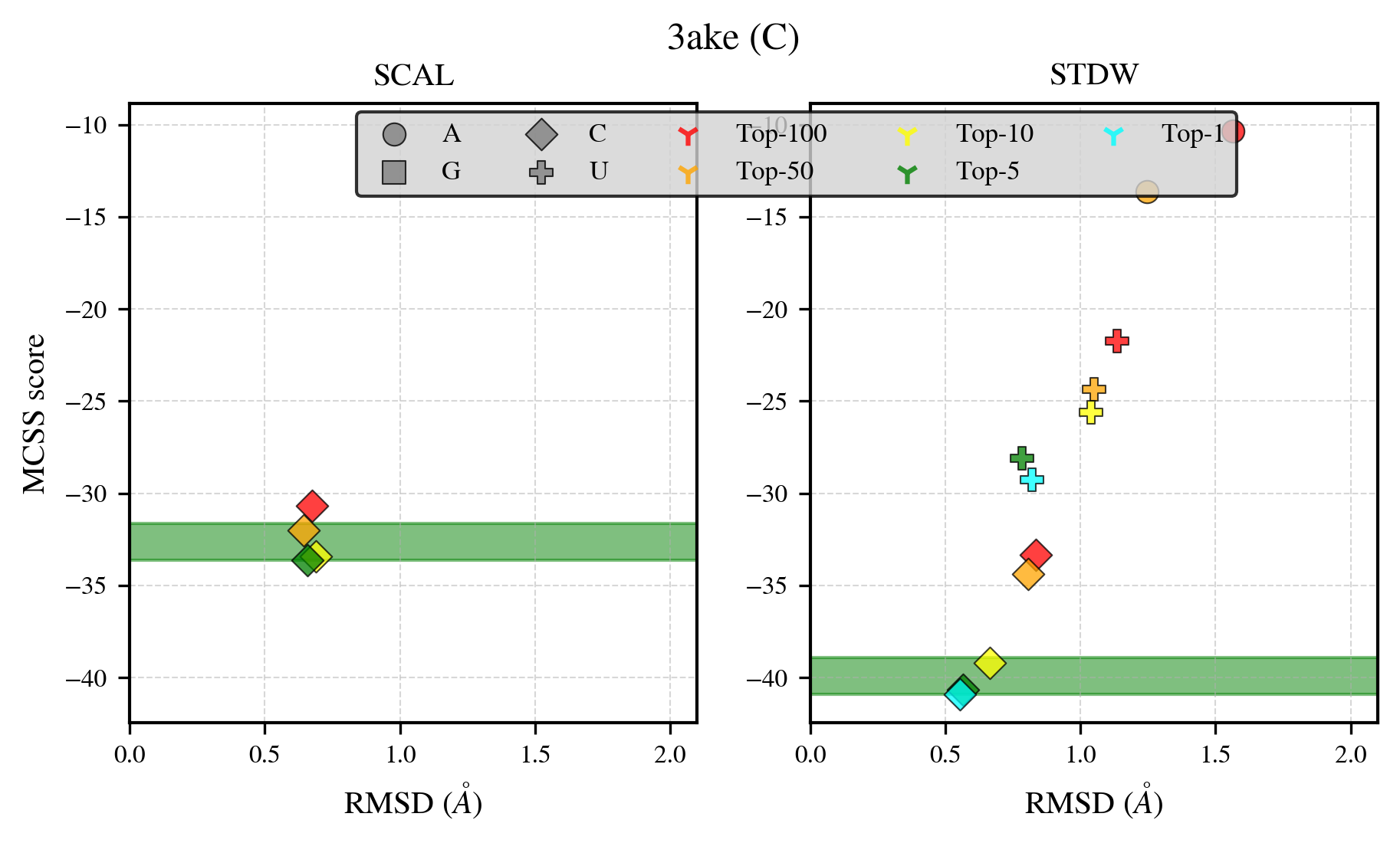

### 3c4z_benchmark-121_selectivity_details.png

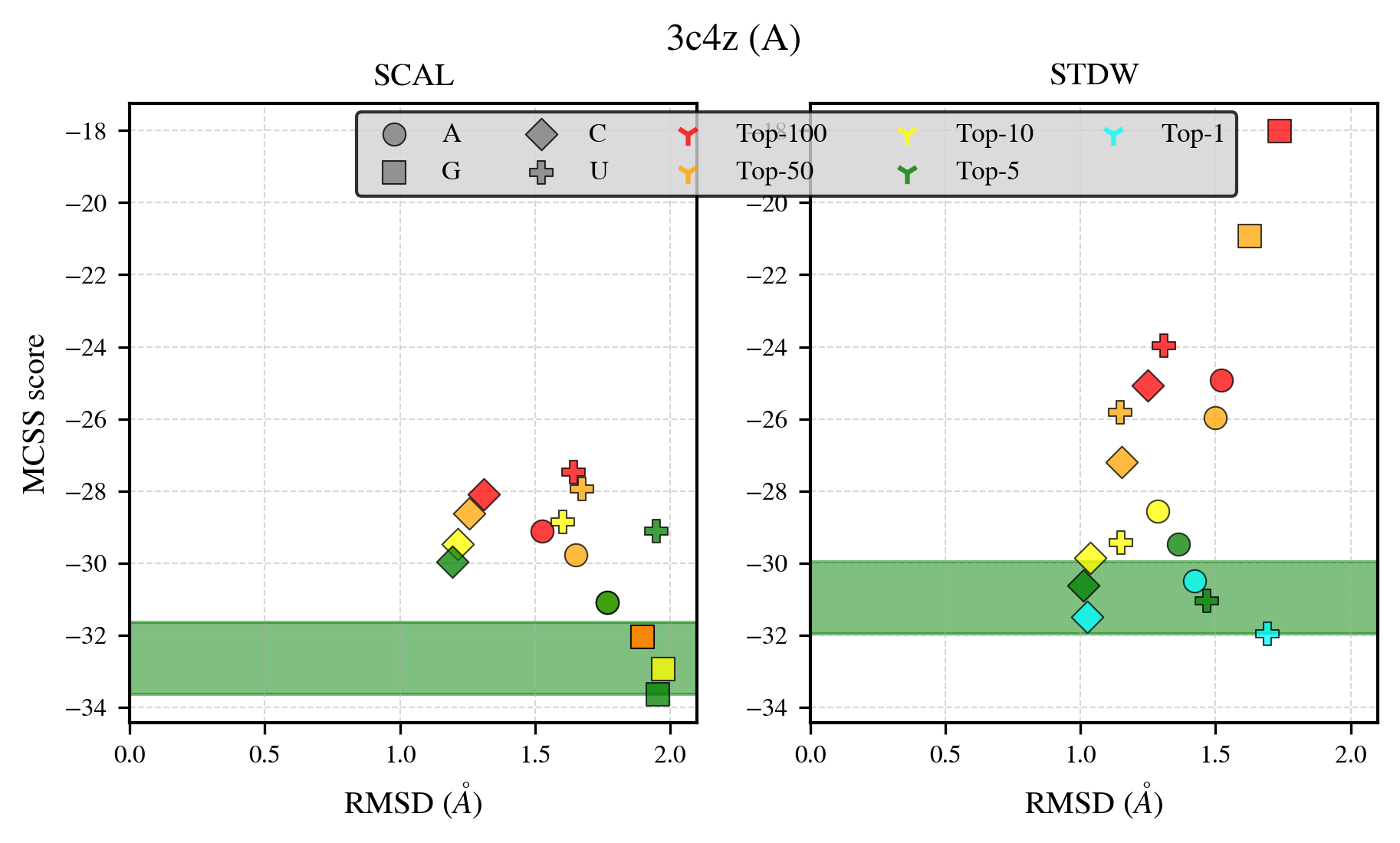

### 3c85_benchmark-121_selectivity_details.png

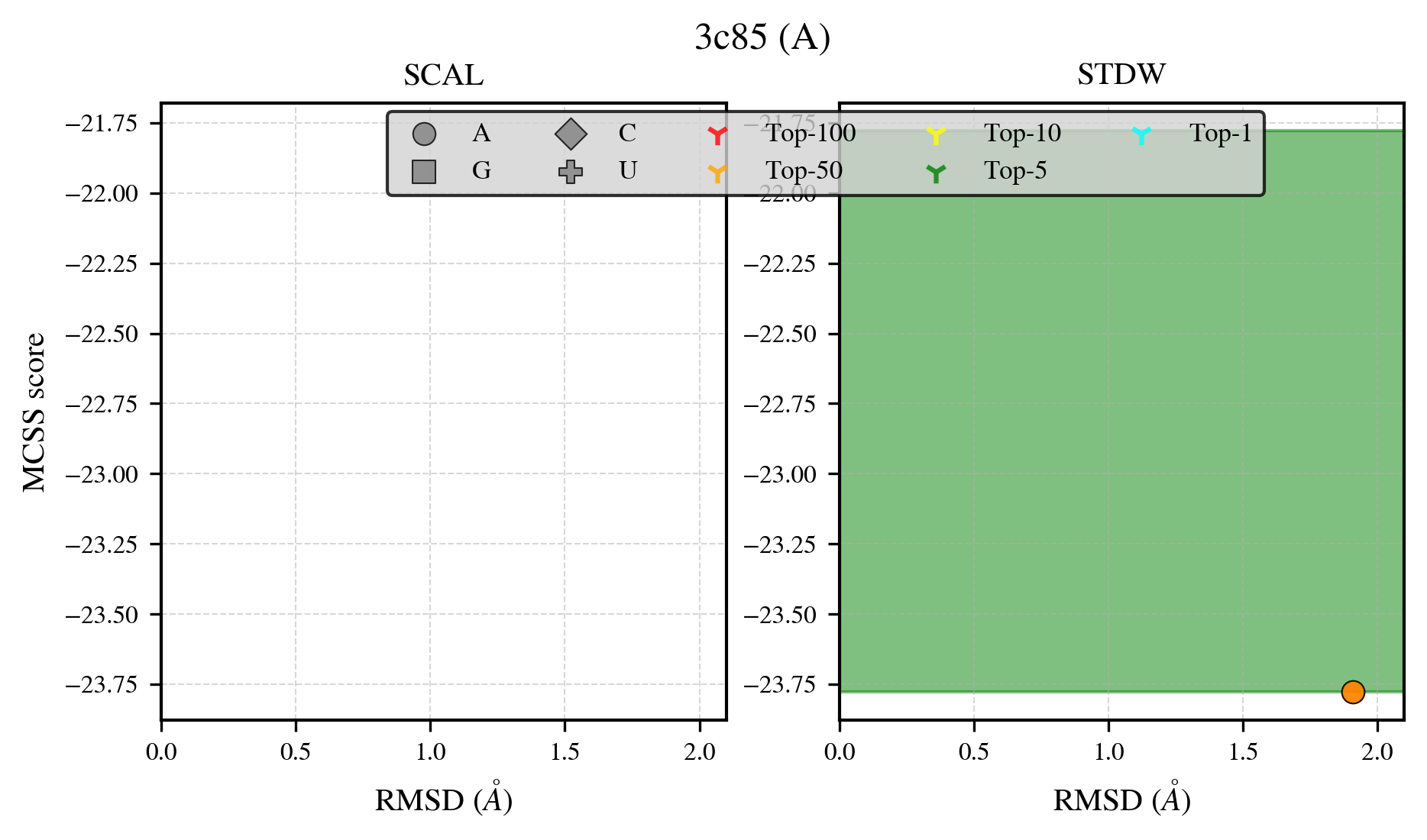

### 3cj9_benchmark-121_selectivity_details.png

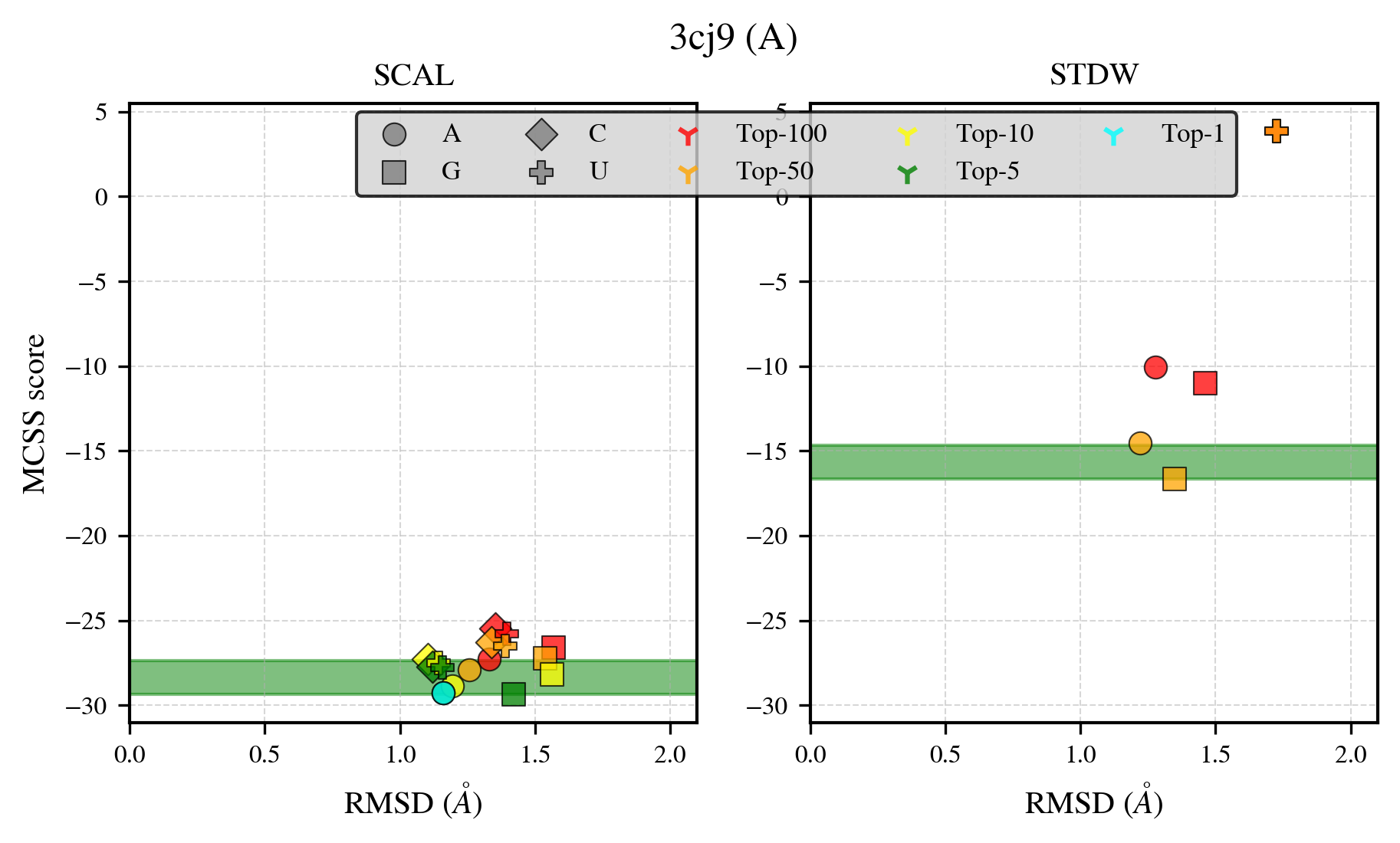

### 3cls_benchmark-121_selectivity_details.png

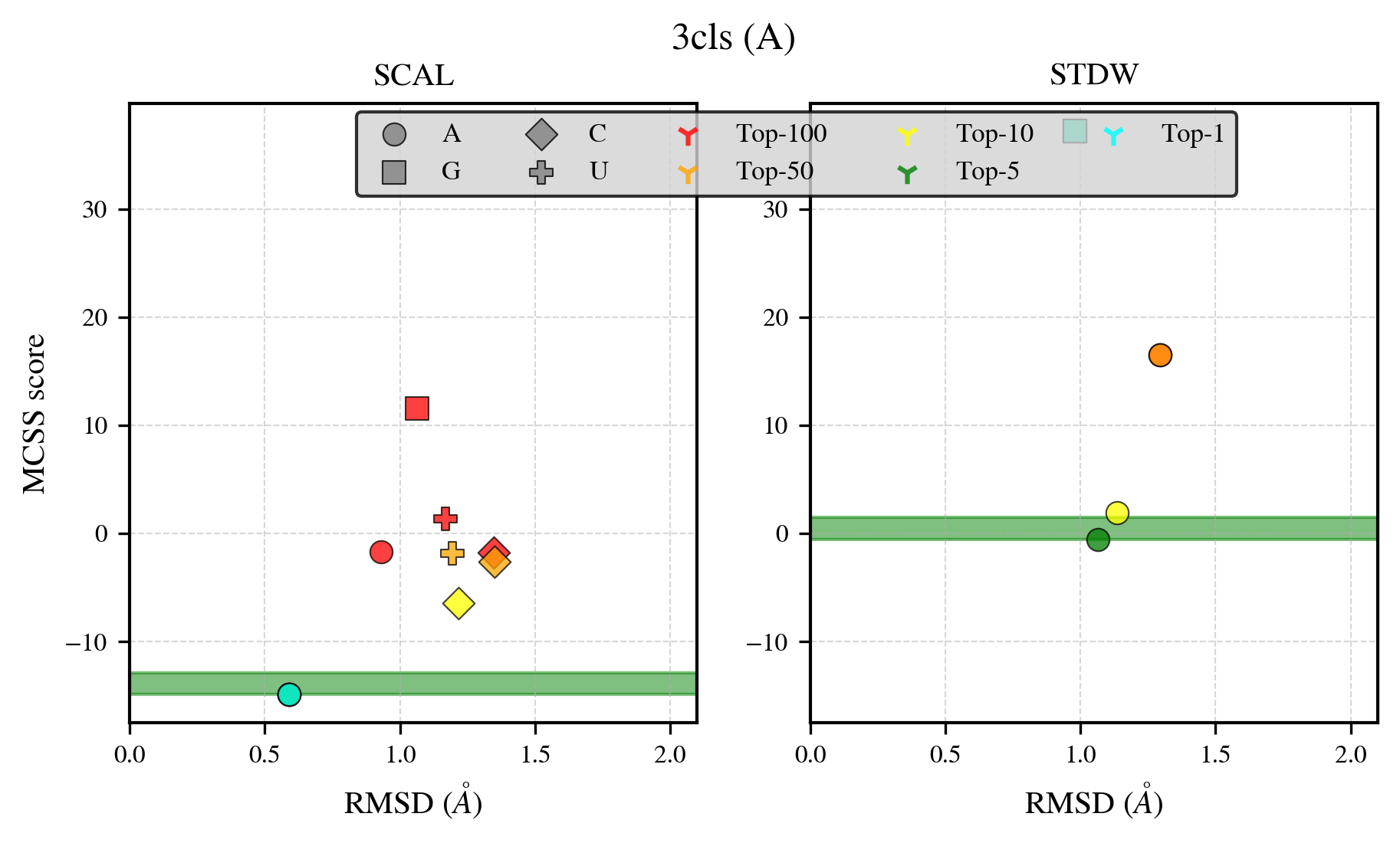

### 3ddj_benchmark-121_selectivity_details.png

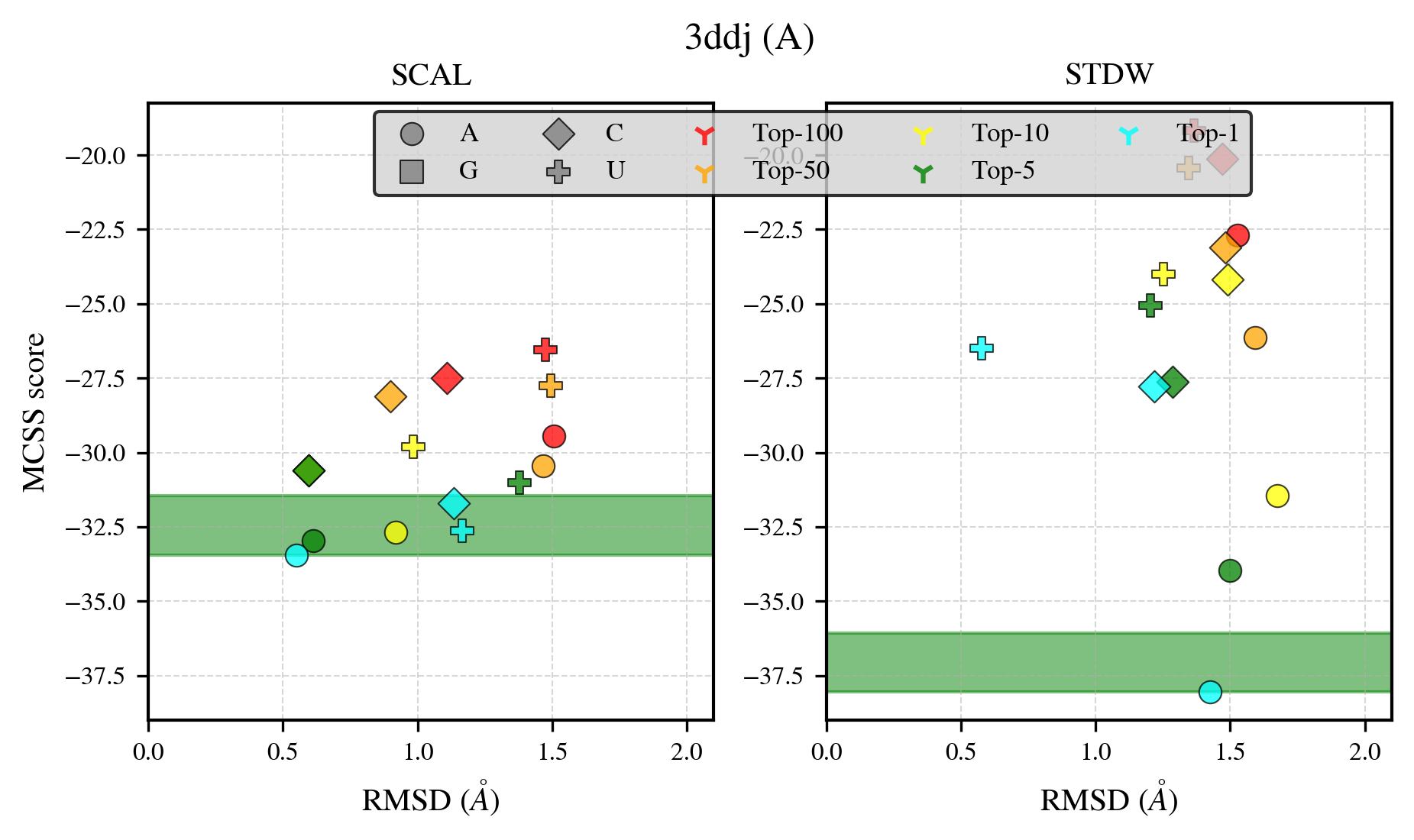

### 3djx_benchmark-121_selectivity_details.png

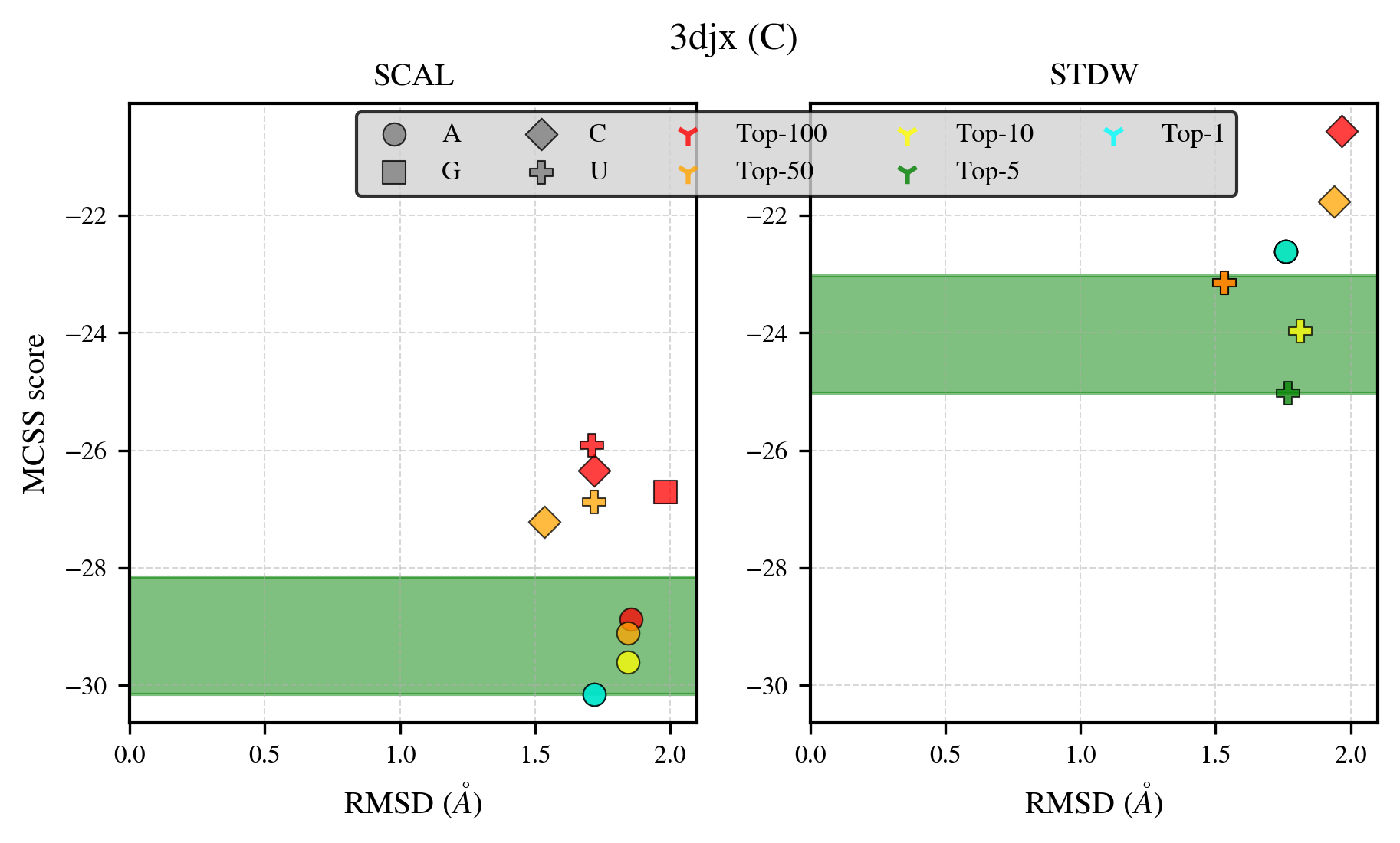

### 3dlz_benchmark-121_selectivity_details.png

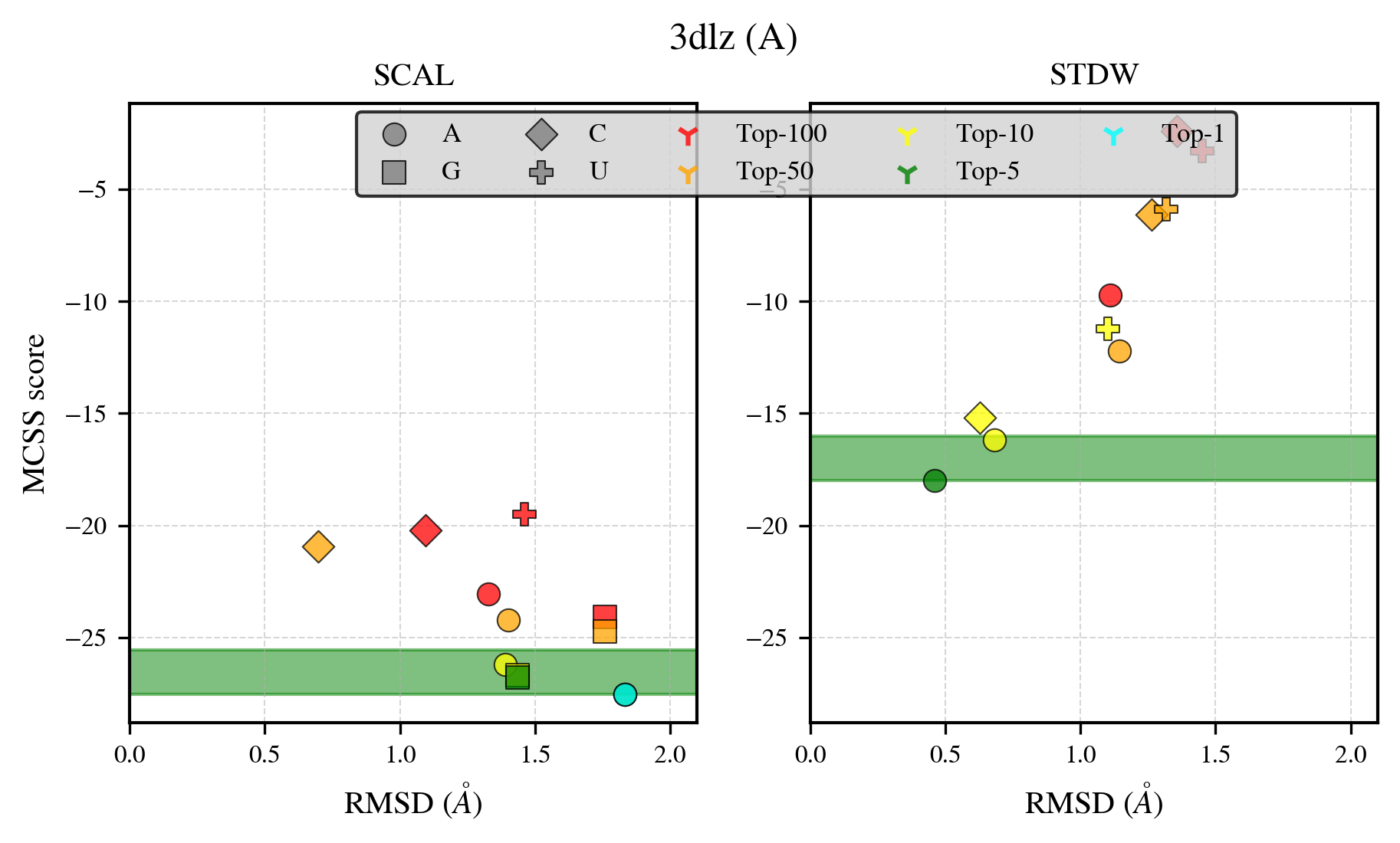

### 3ewy_benchmark-121_selectivity_details.png

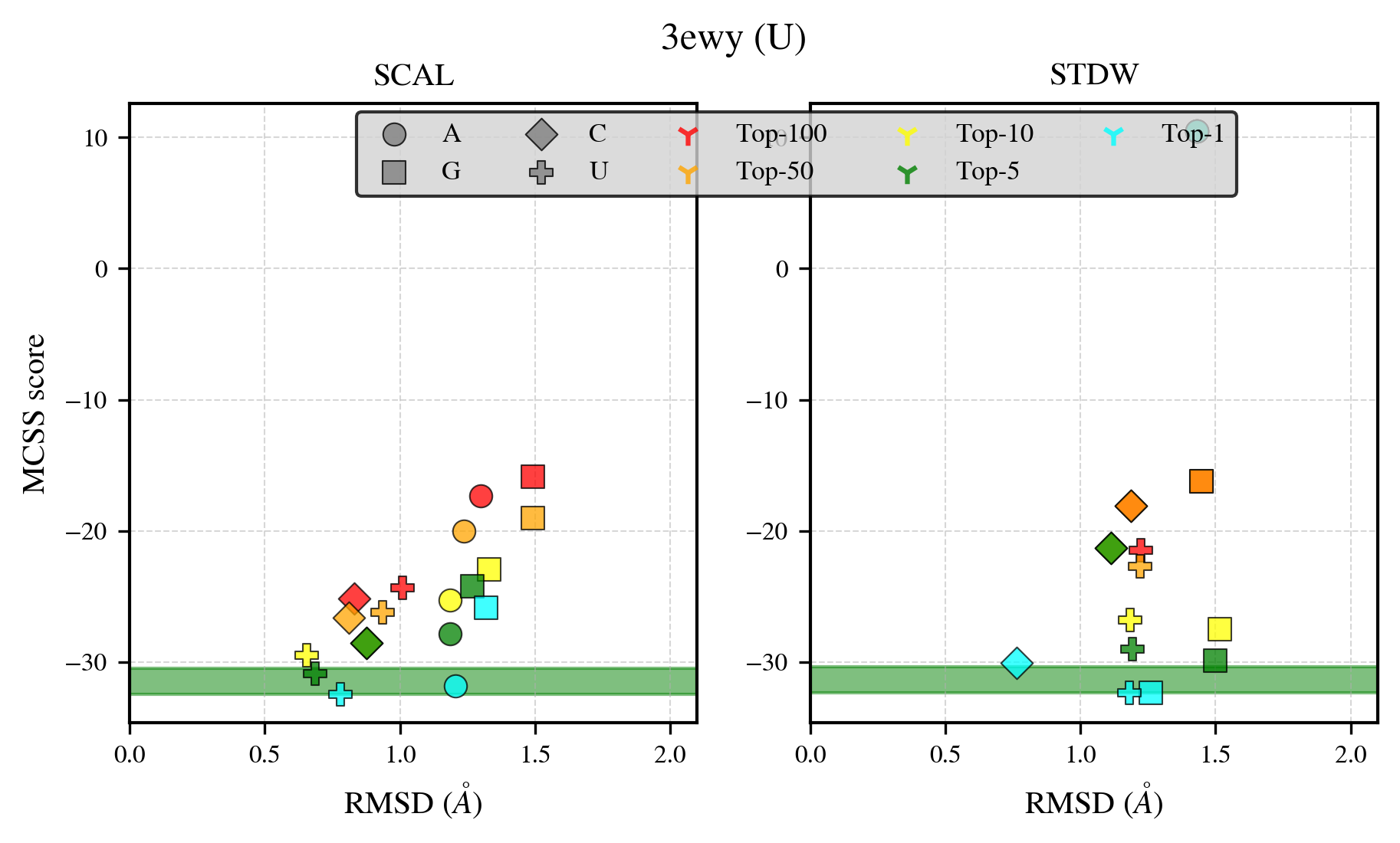

### 3glv_benchmark-121_selectivity_details.png

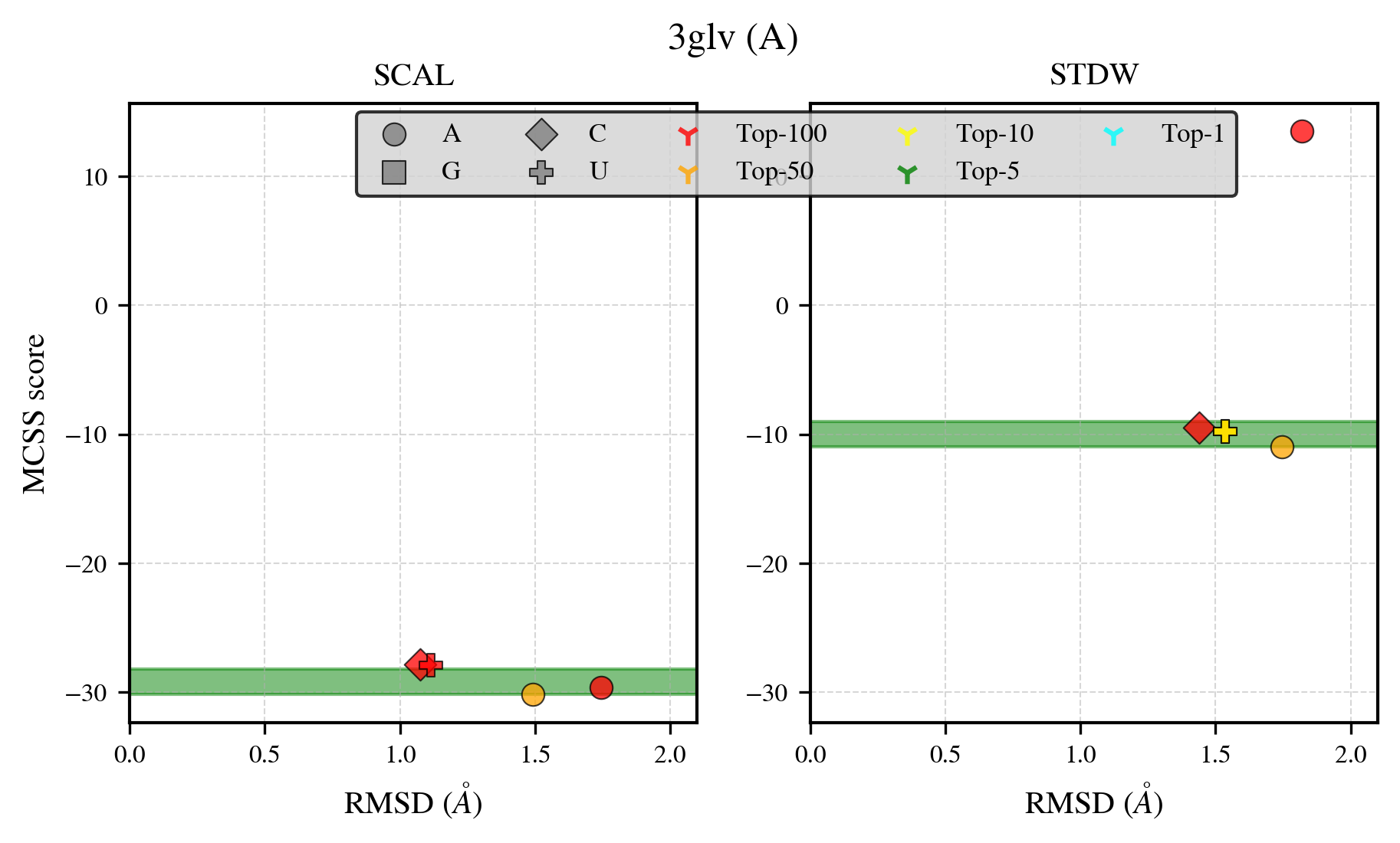

### 3gru_benchmark-121_selectivity_details.png

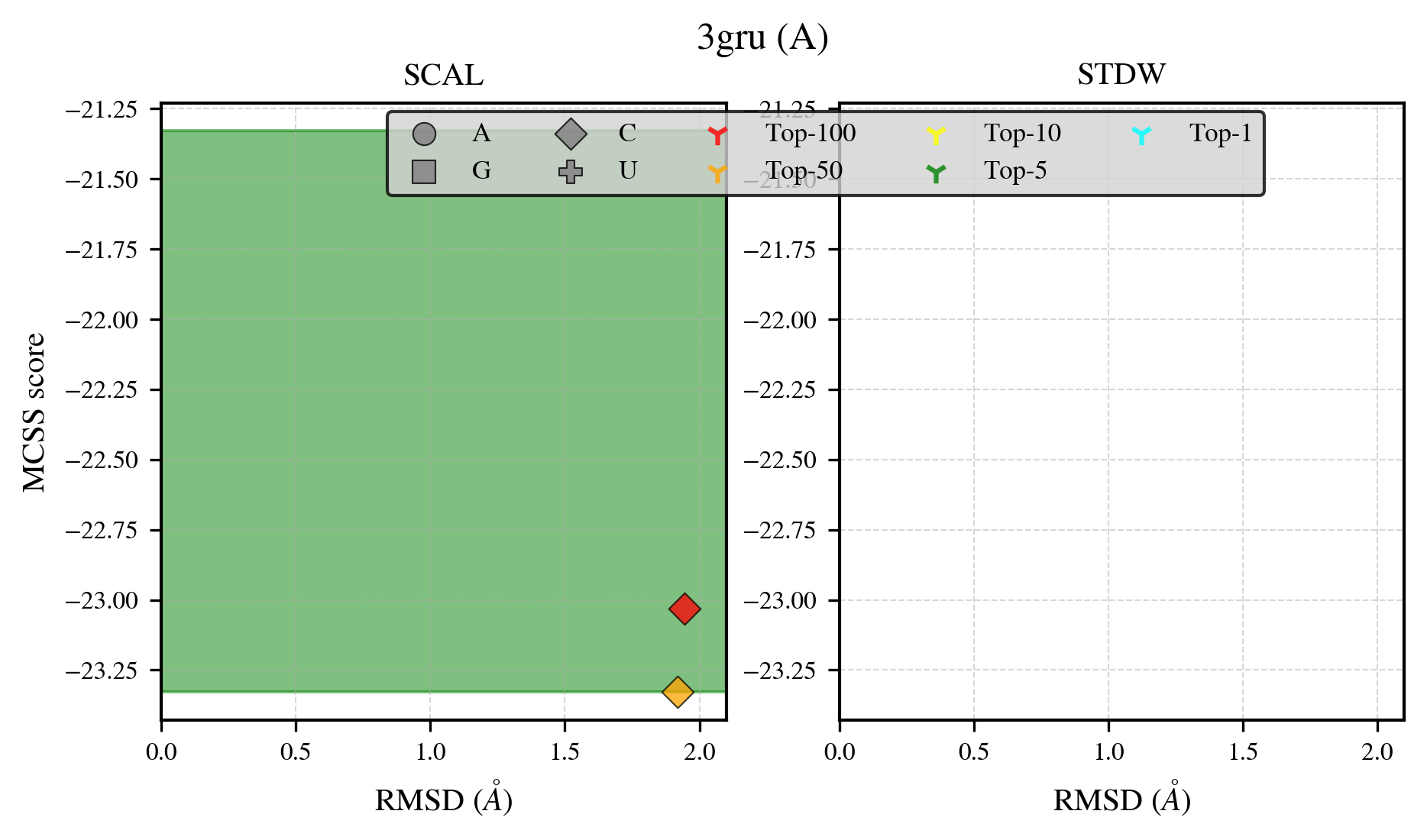

### 3kd6_benchmark-121_selectivity_details.png

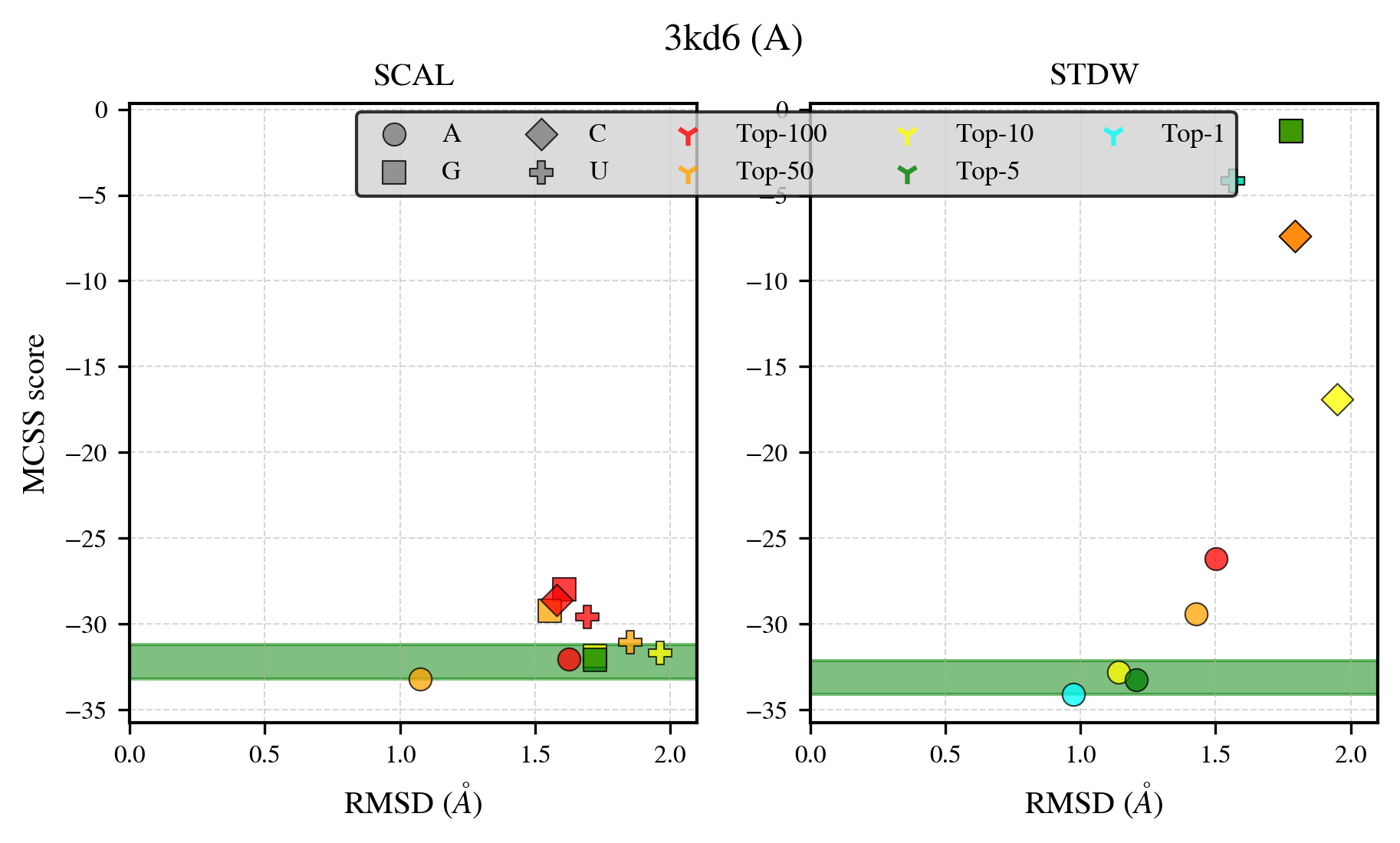

### 3kgd_benchmark-121_selectivity_details.png

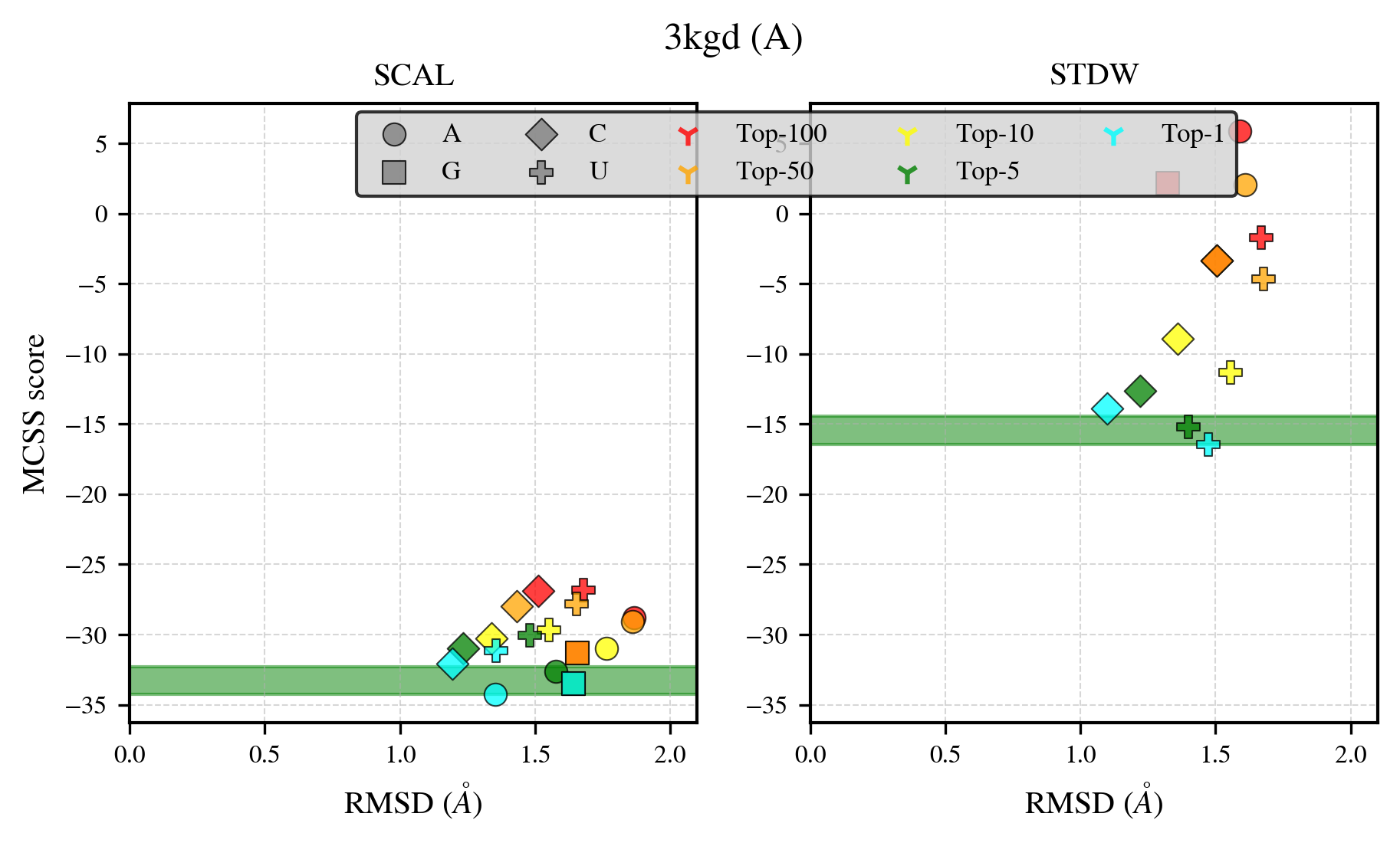

### 3lfr_benchmark-121_selectivity_details.png

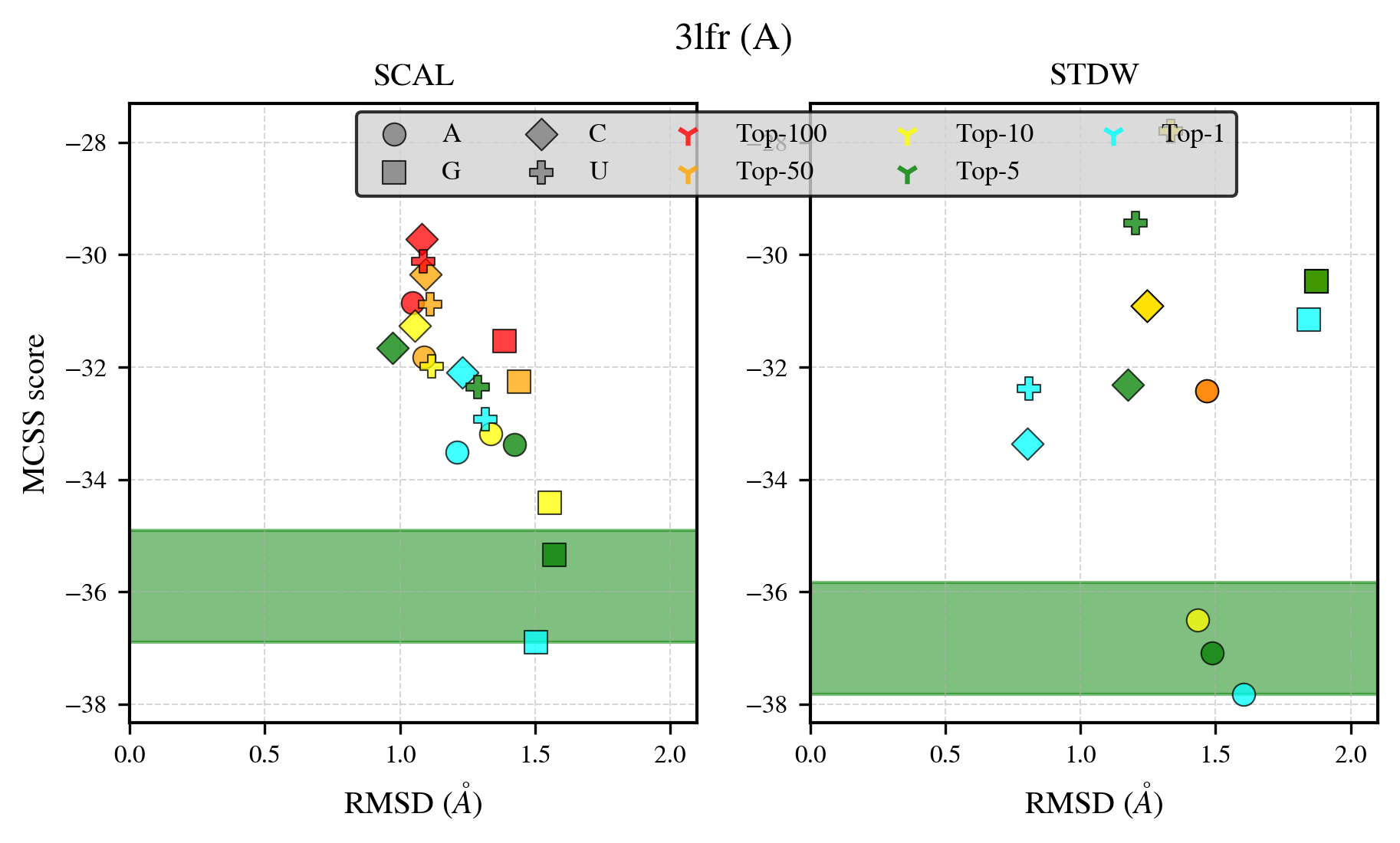

### 3lkm_benchmark-121_selectivity_details.png

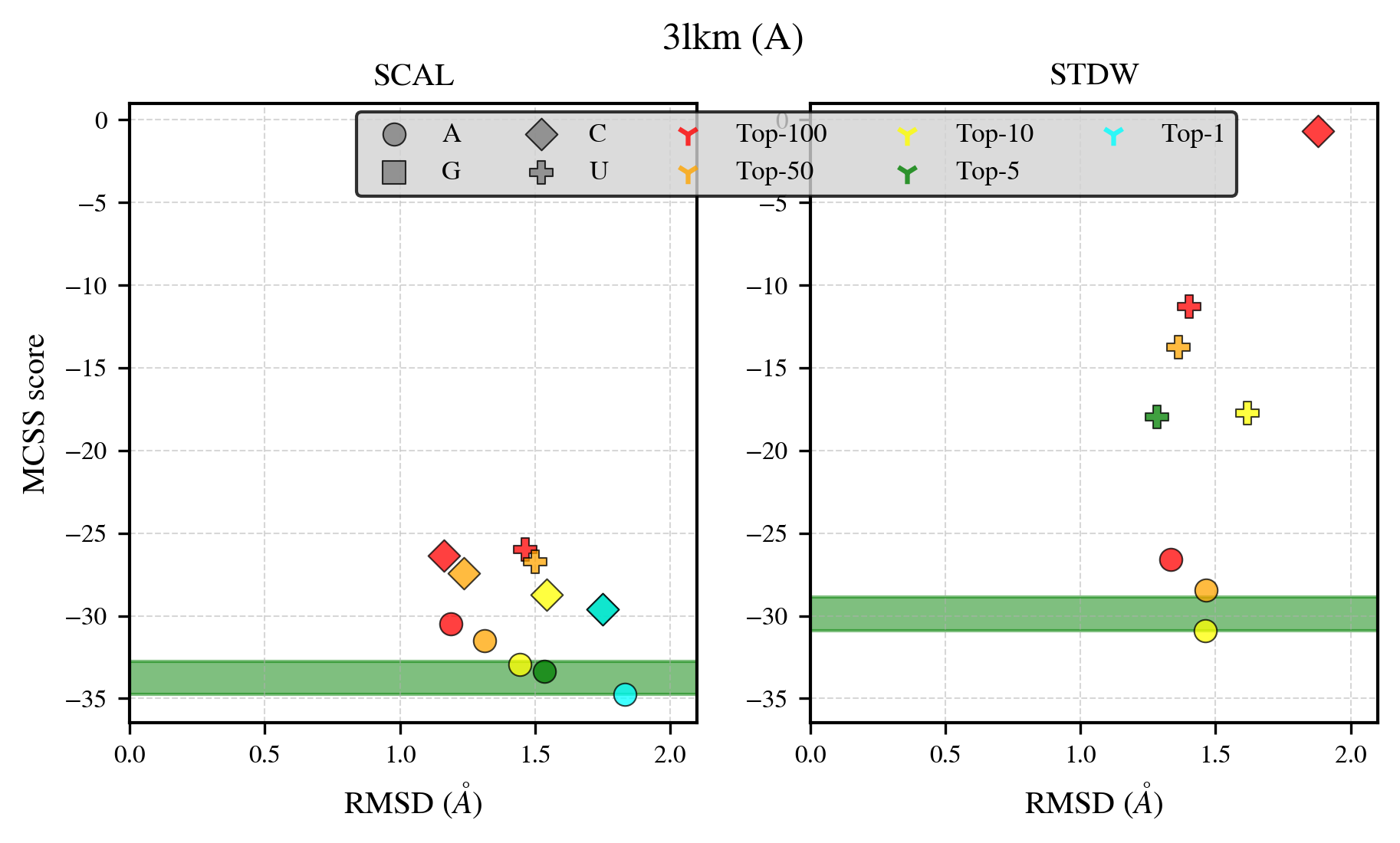

### 3m84_benchmark-121_selectivity_details.png

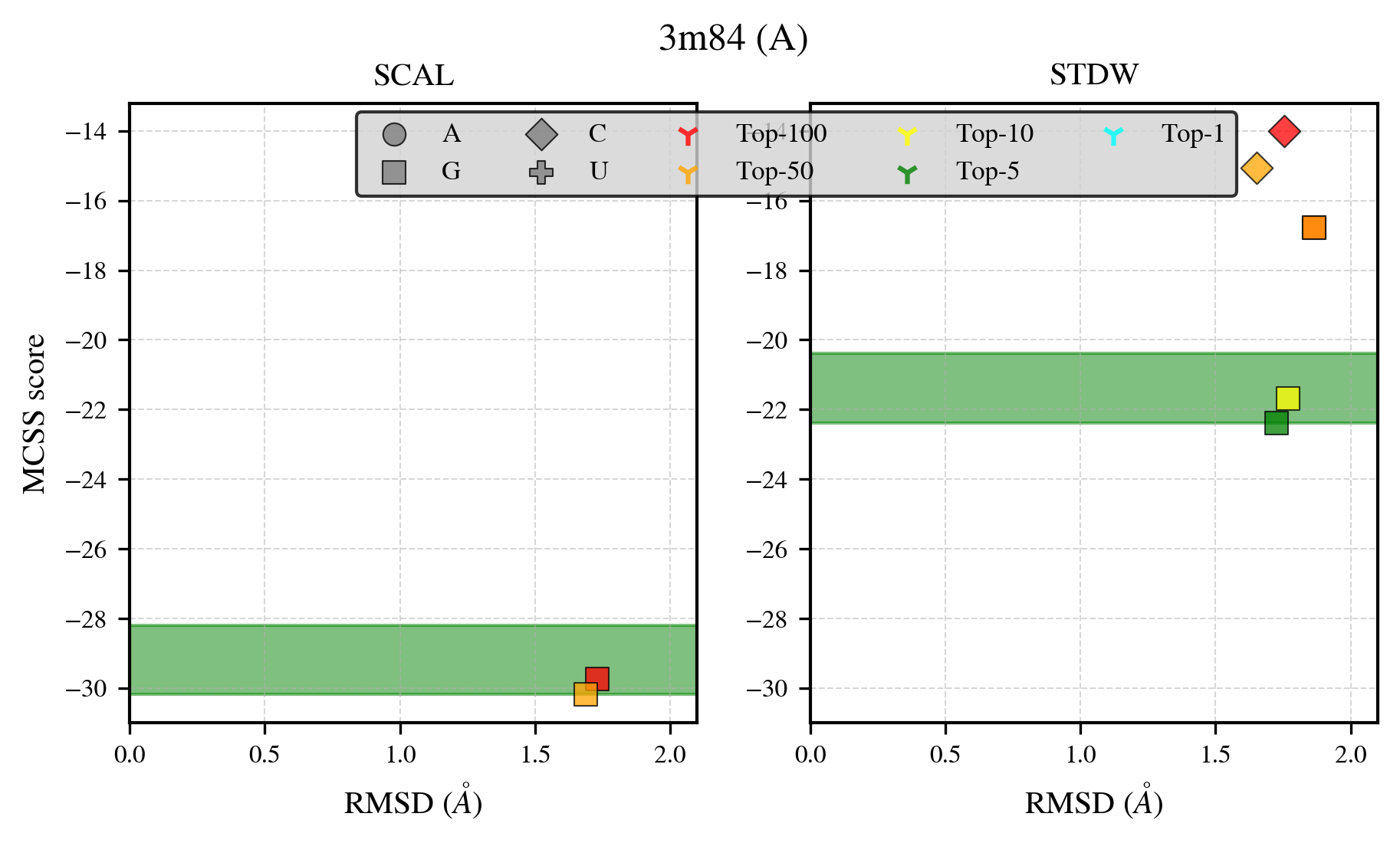

### 3n1s_benchmark-121_selectivity_details.png

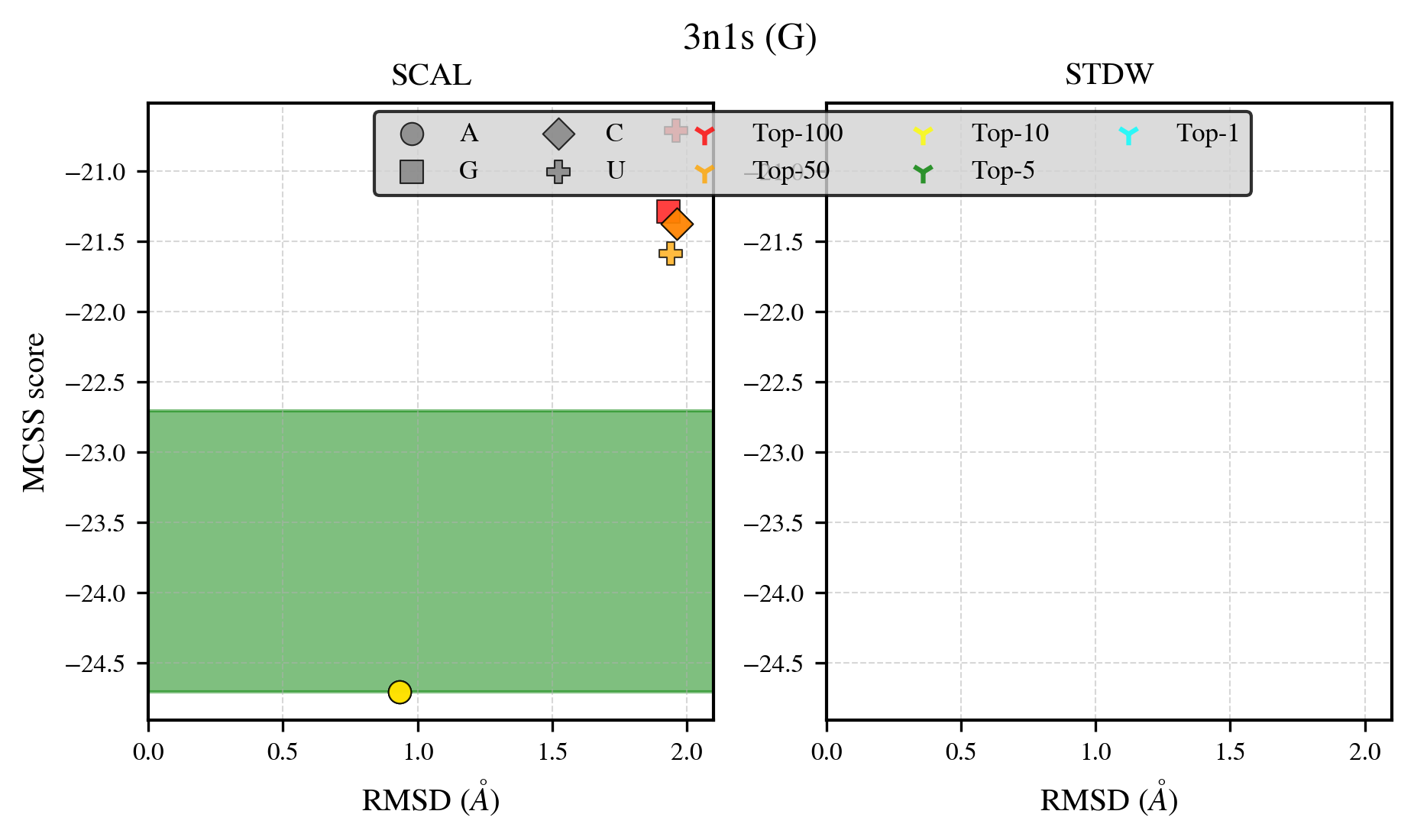

### 3nua_benchmark-121_selectivity_details.png

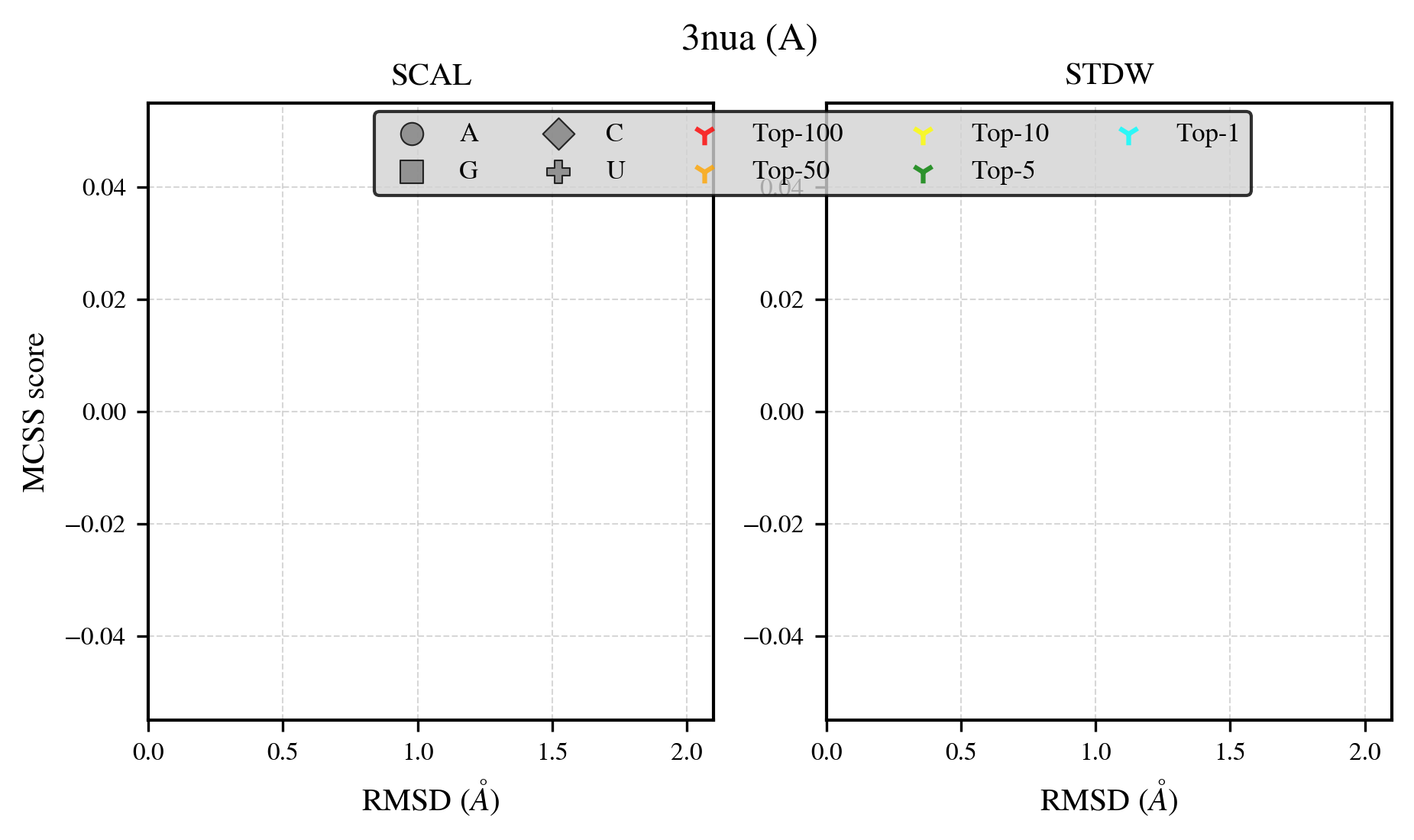

### 3nyq_benchmark-121_selectivity_details.png

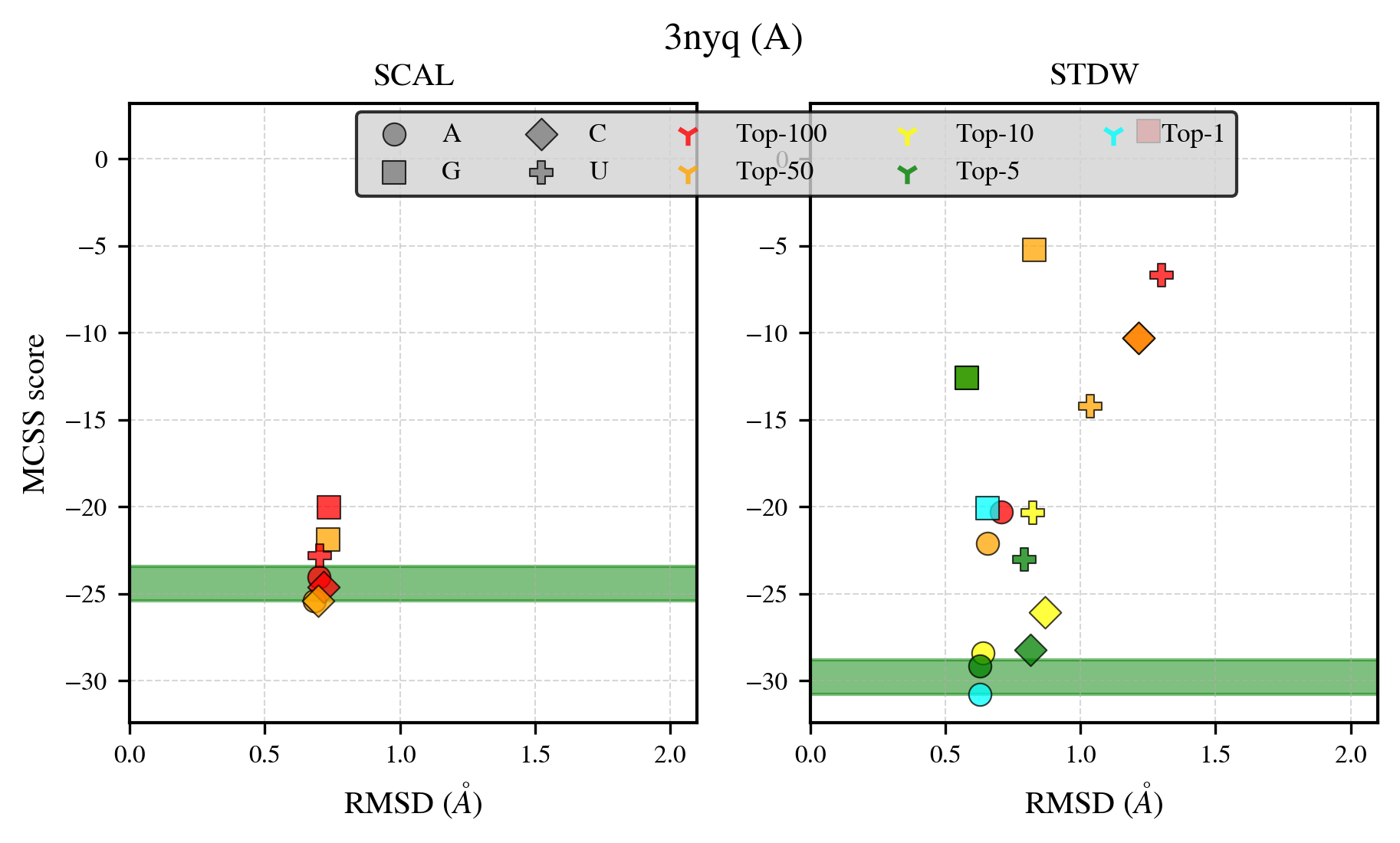

### 3omf_benchmark-121_selectivity_details.png

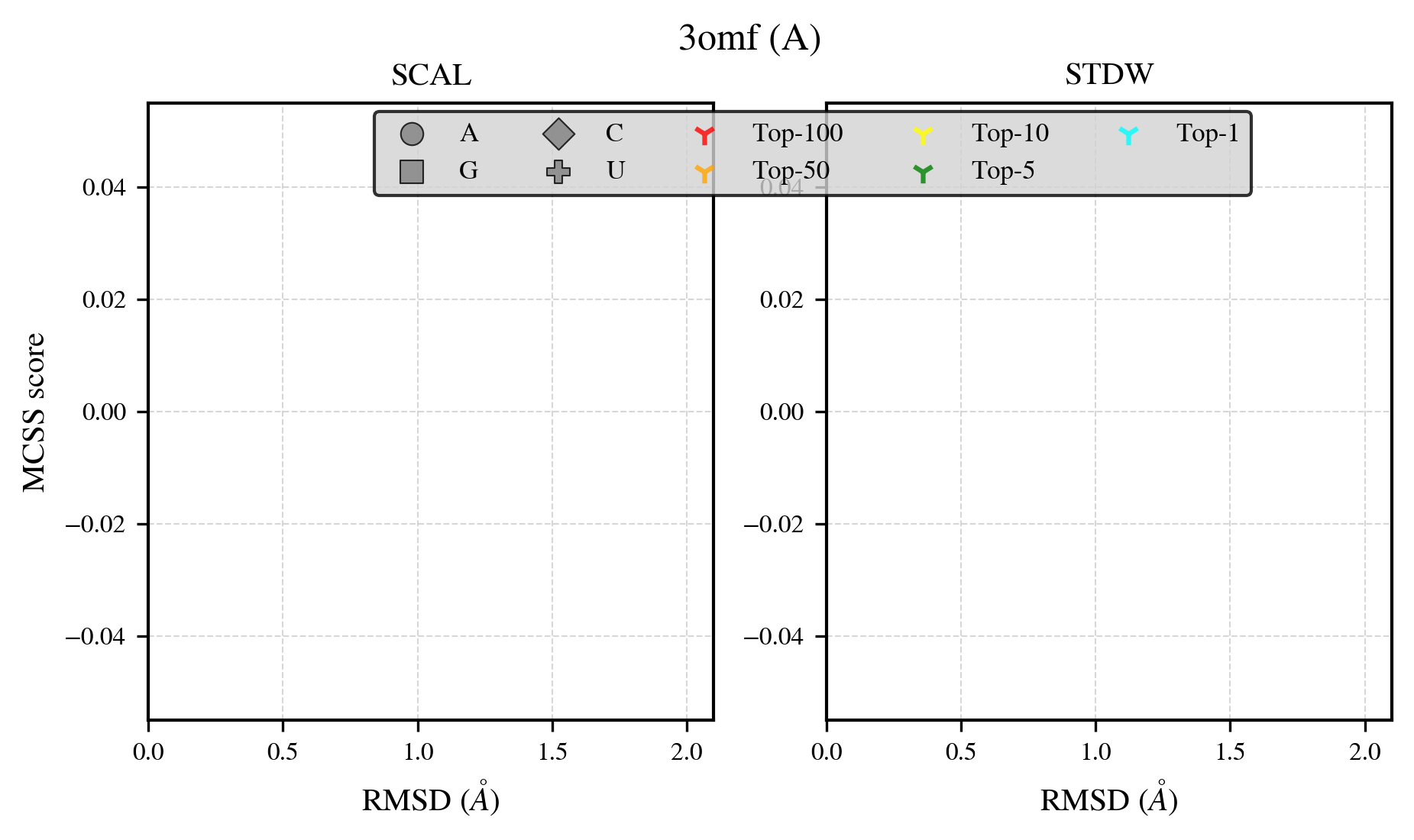

### 3pln_benchmark-121_selectivity_details.png

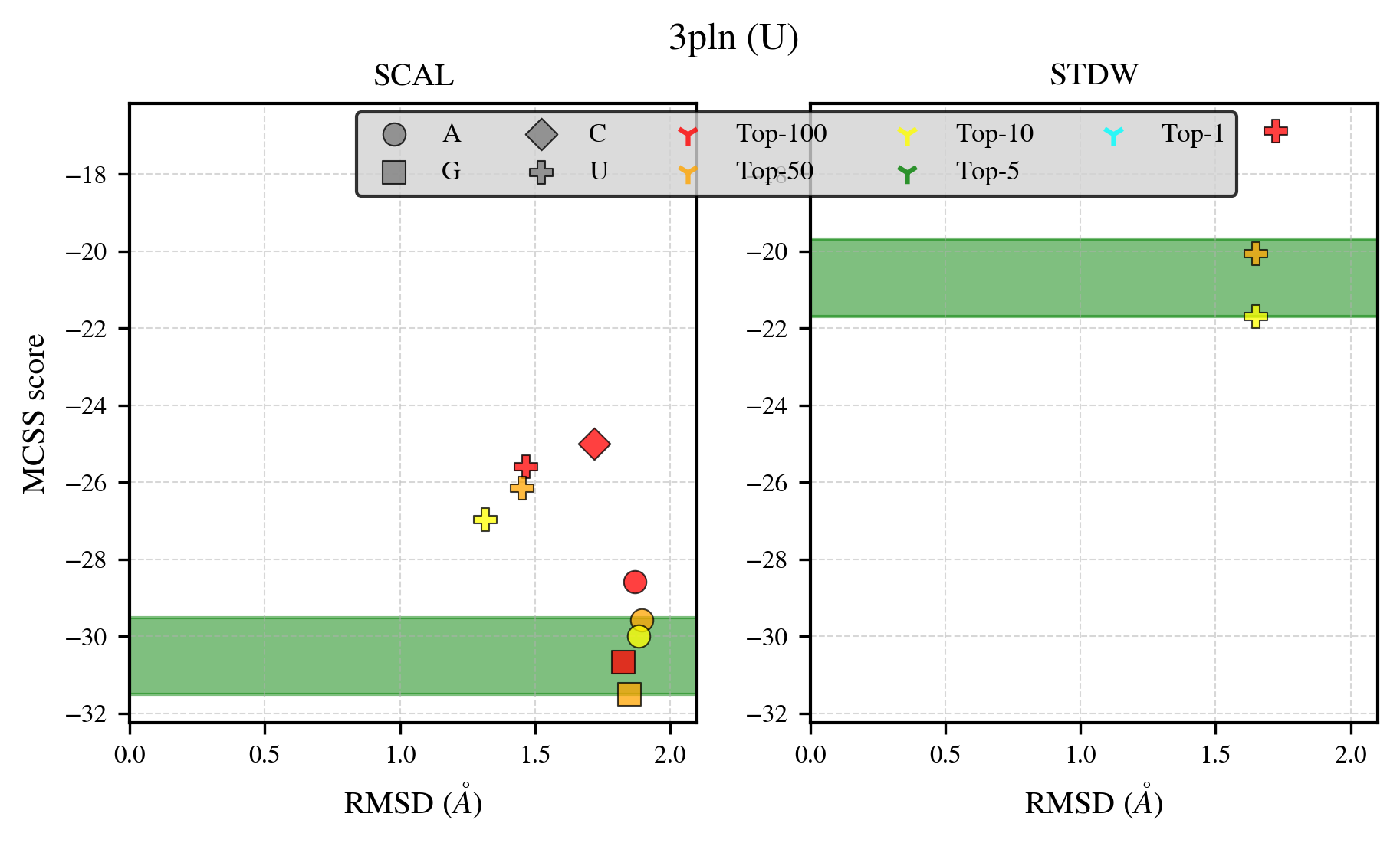

### 3rl4_benchmark-121_selectivity_details.png

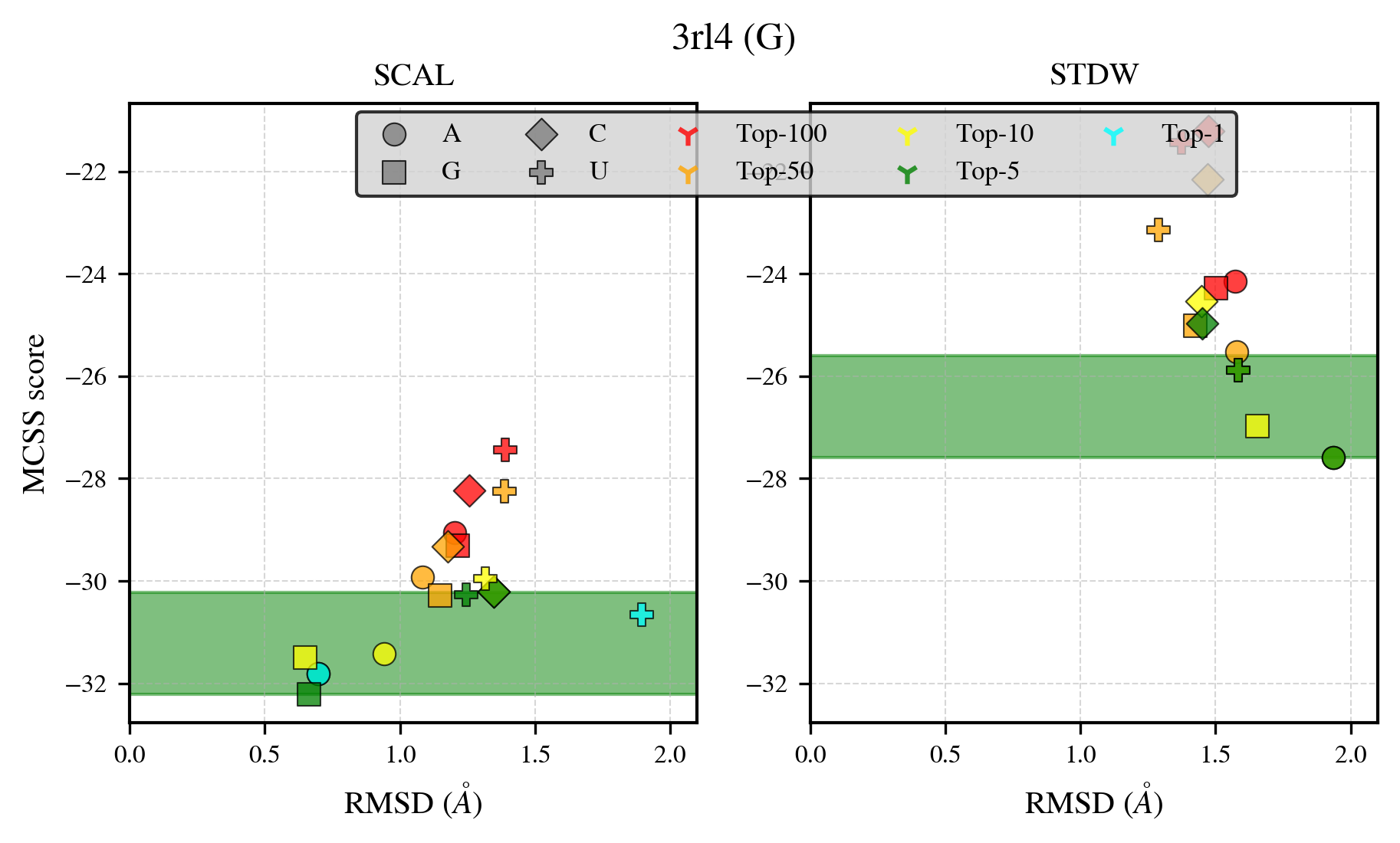

### 3rpz_benchmark-121_selectivity_details.png

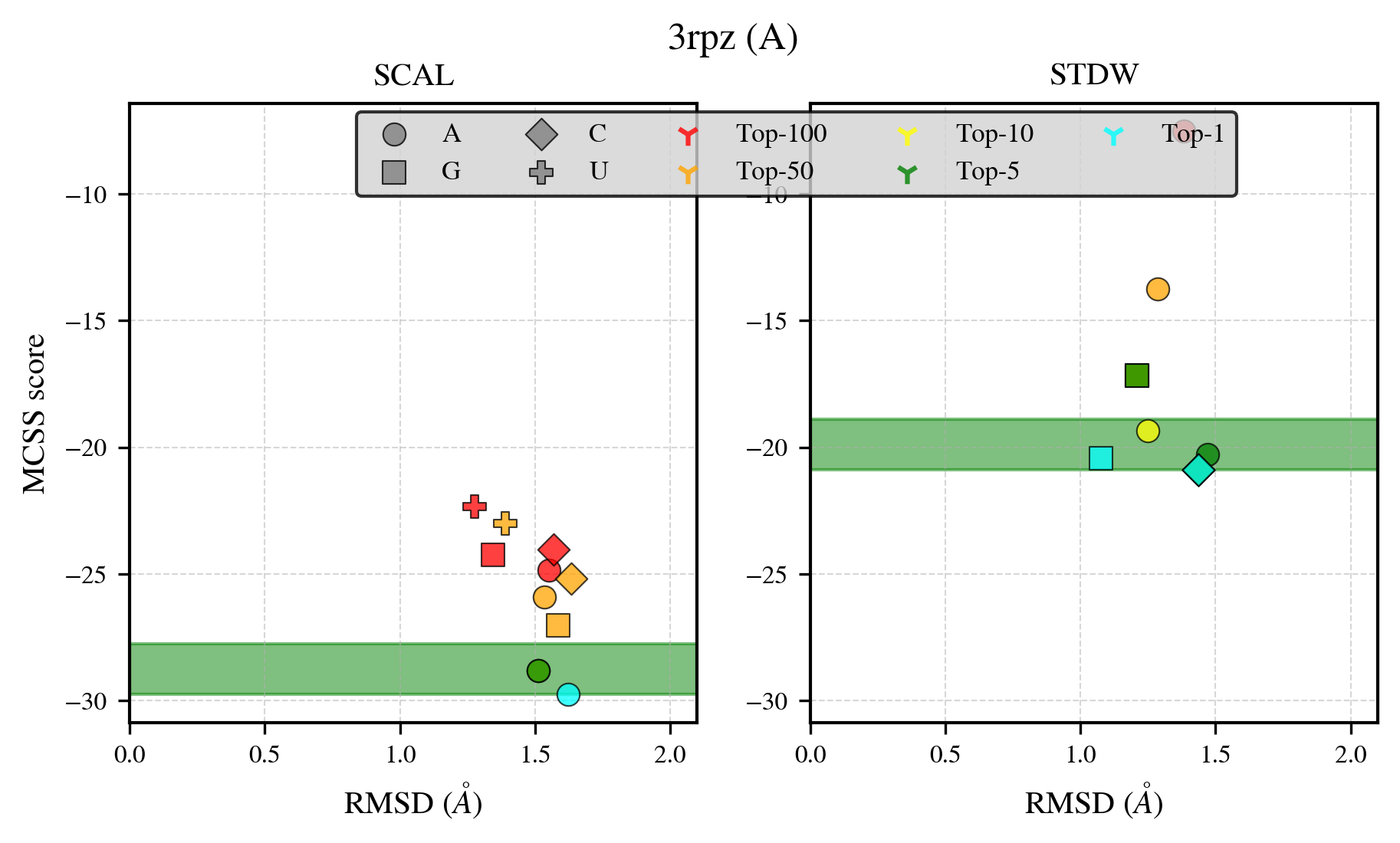

### 3uq8_benchmark-121_selectivity_details.png

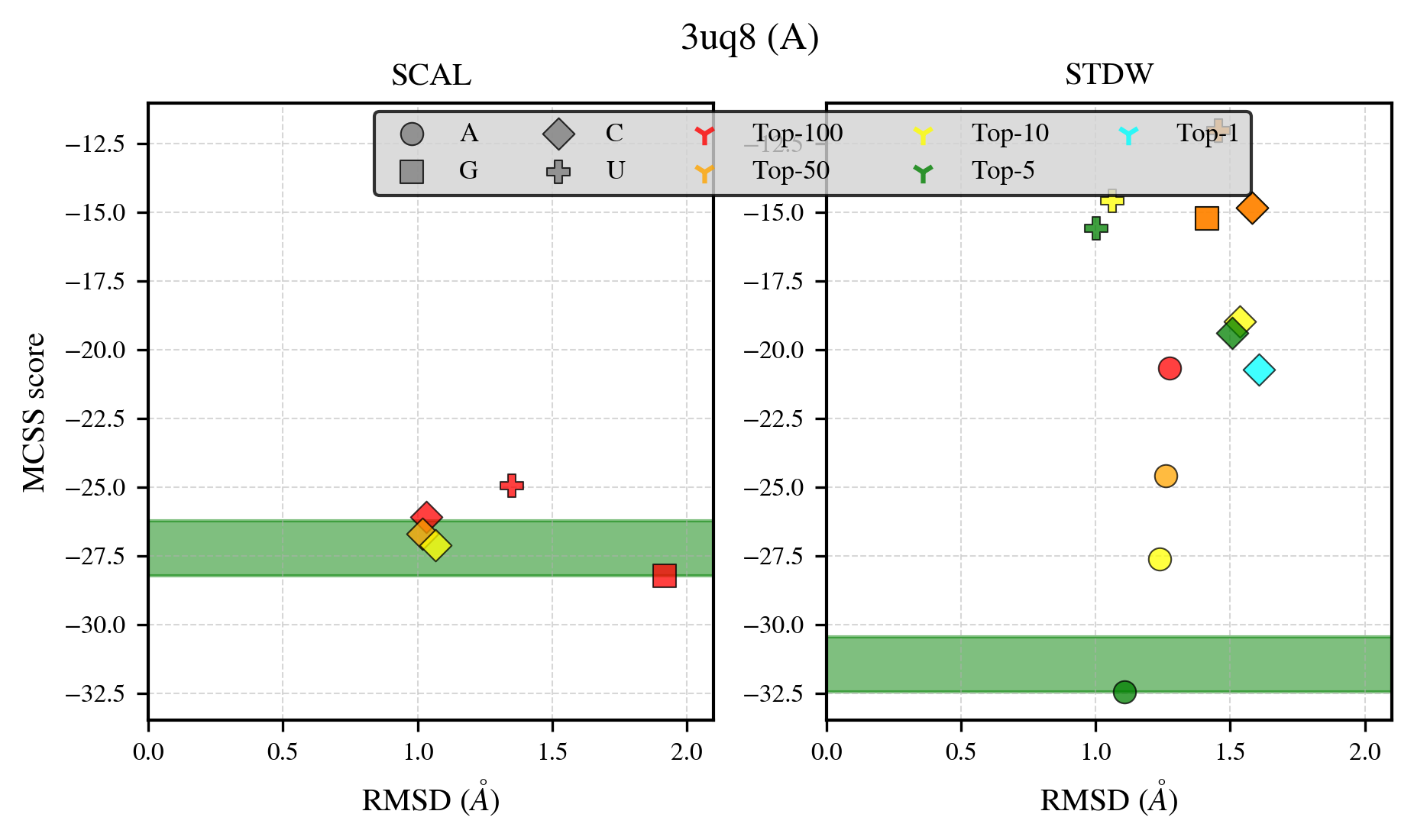

### 3uwq_benchmark-121_selectivity_details.png

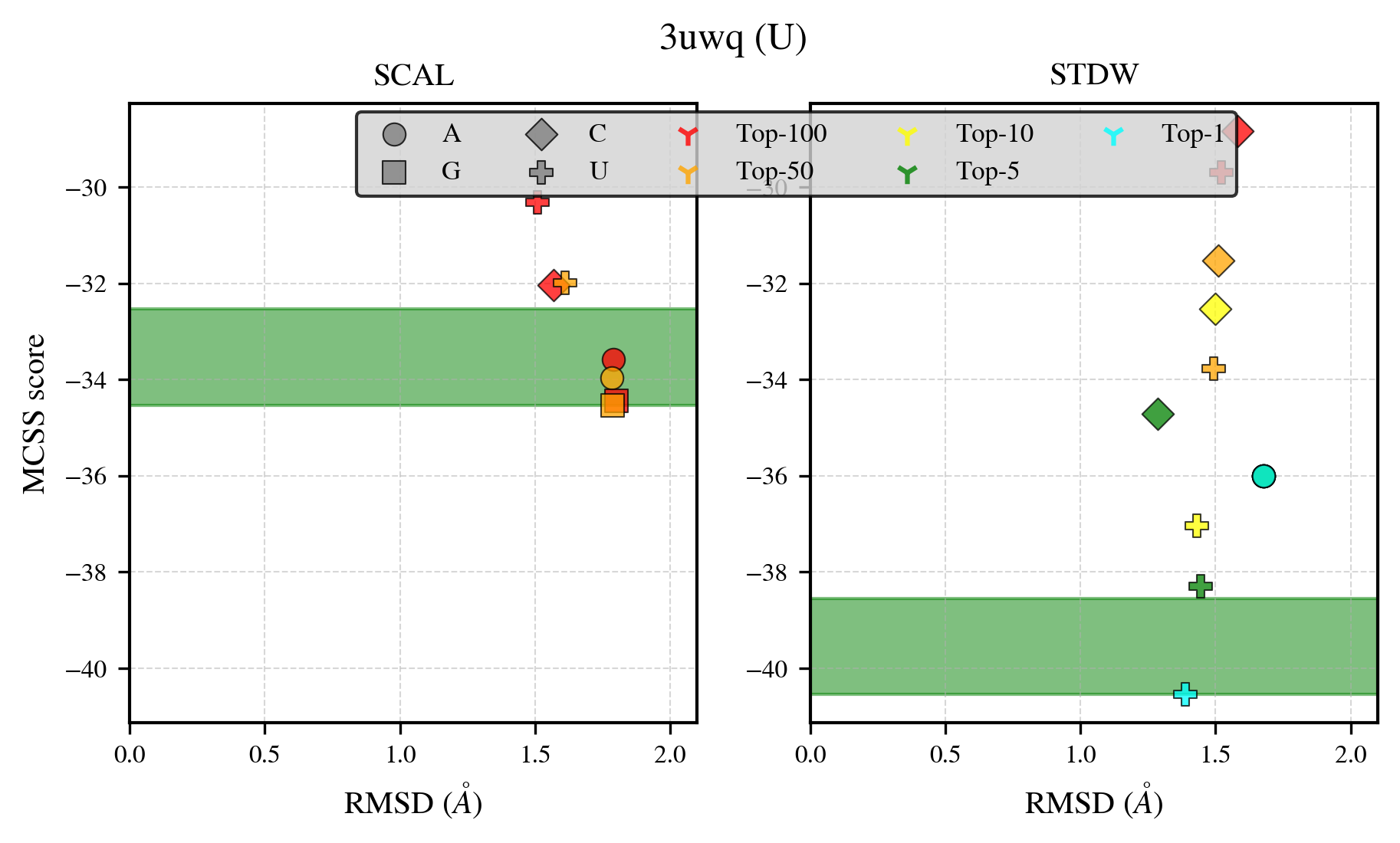

### 3w07_benchmark-121_selectivity_details.png

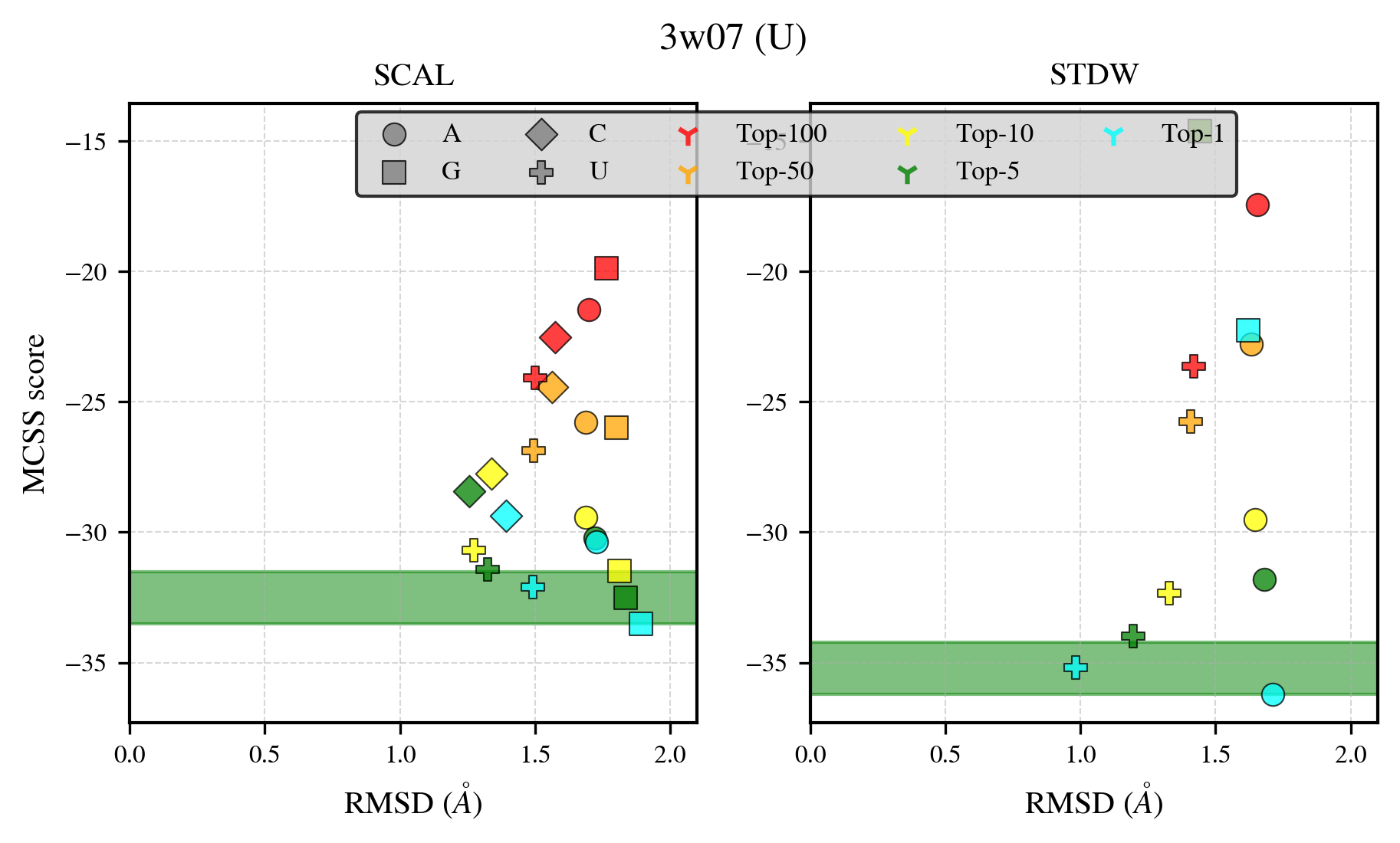

### 4blw_benchmark-121_selectivity_details.png

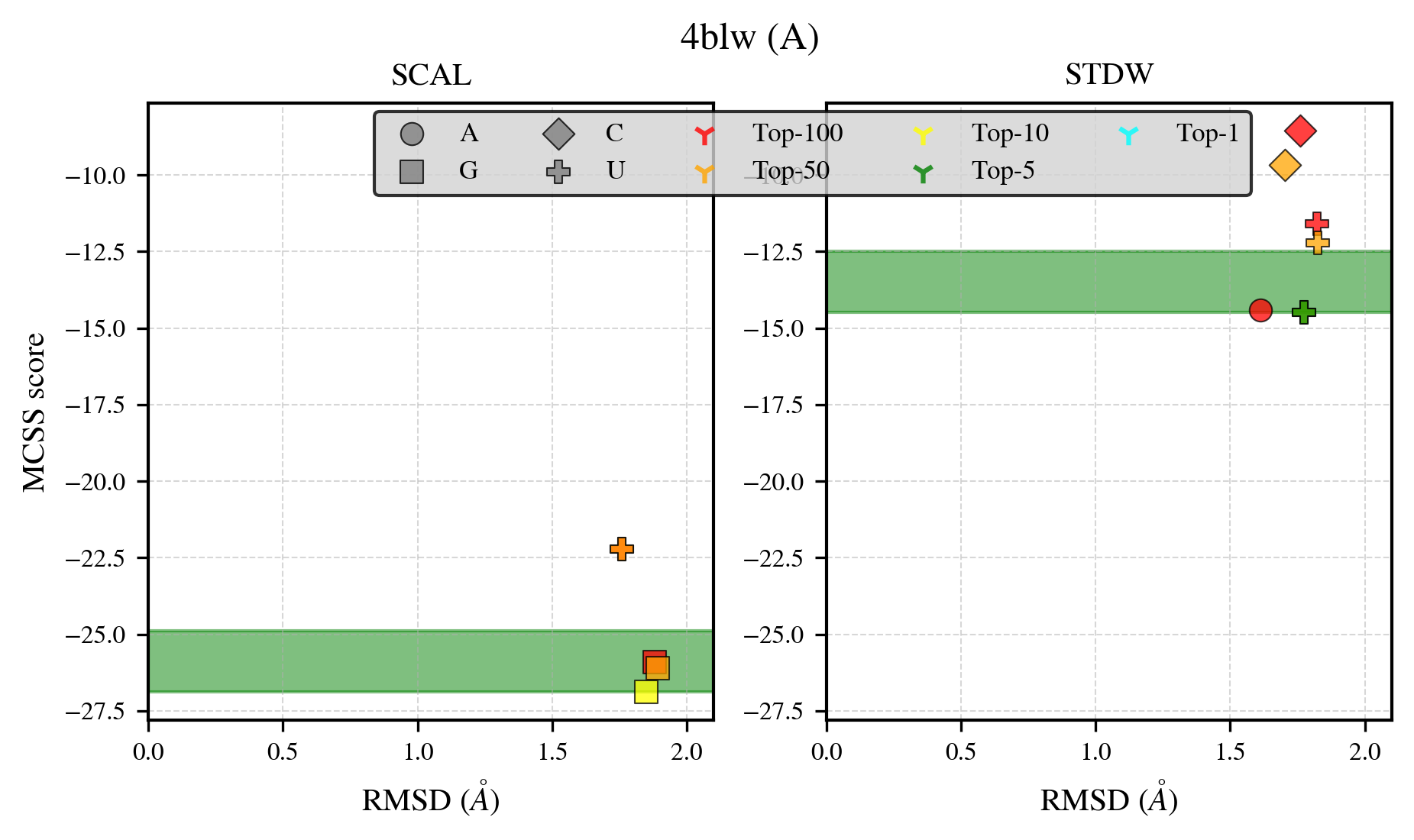

### 4brq_benchmark-121_selectivity_details.png

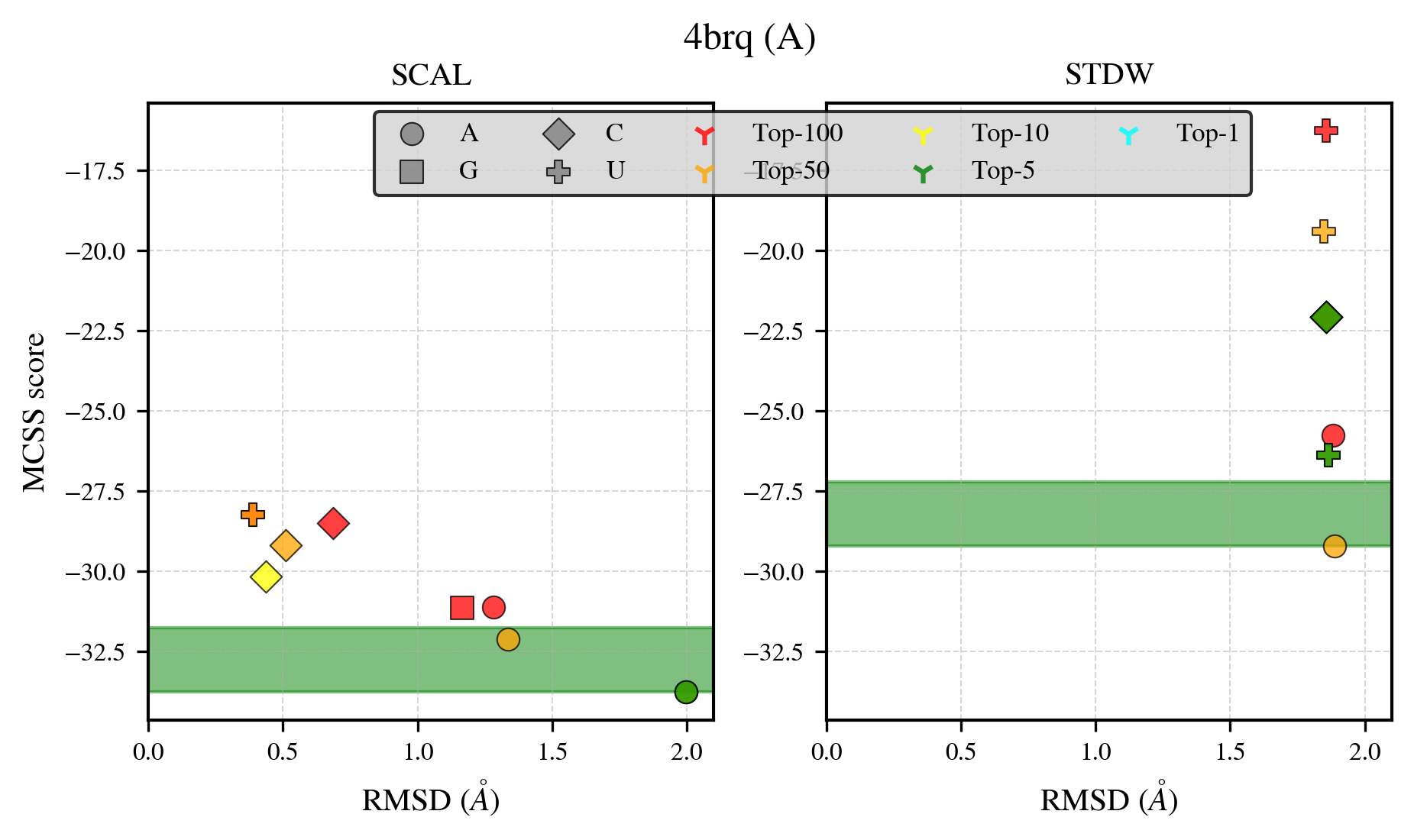

### 4cs3_benchmark-121_selectivity_details.png

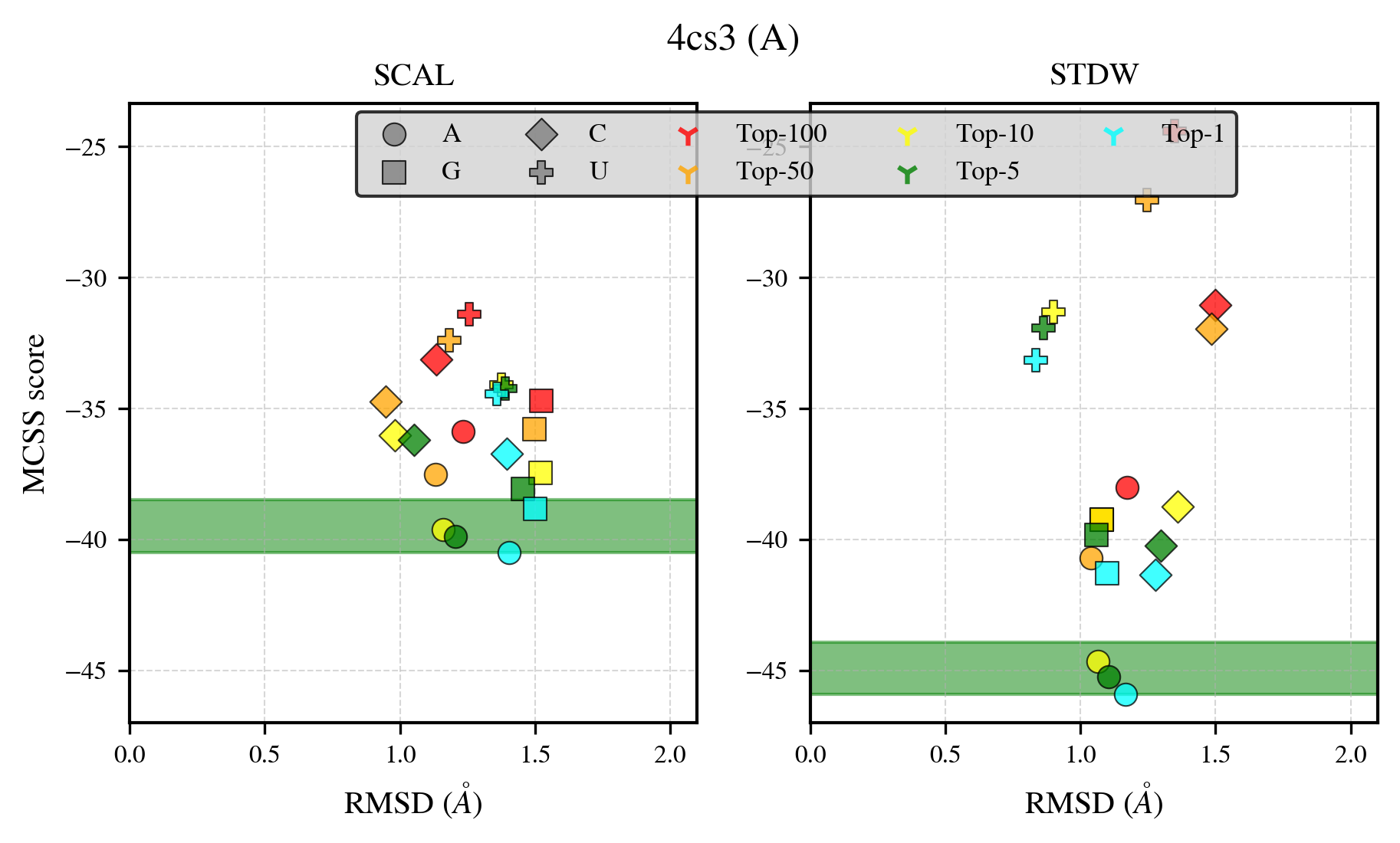

### 4d05_benchmark-121_selectivity_details.png

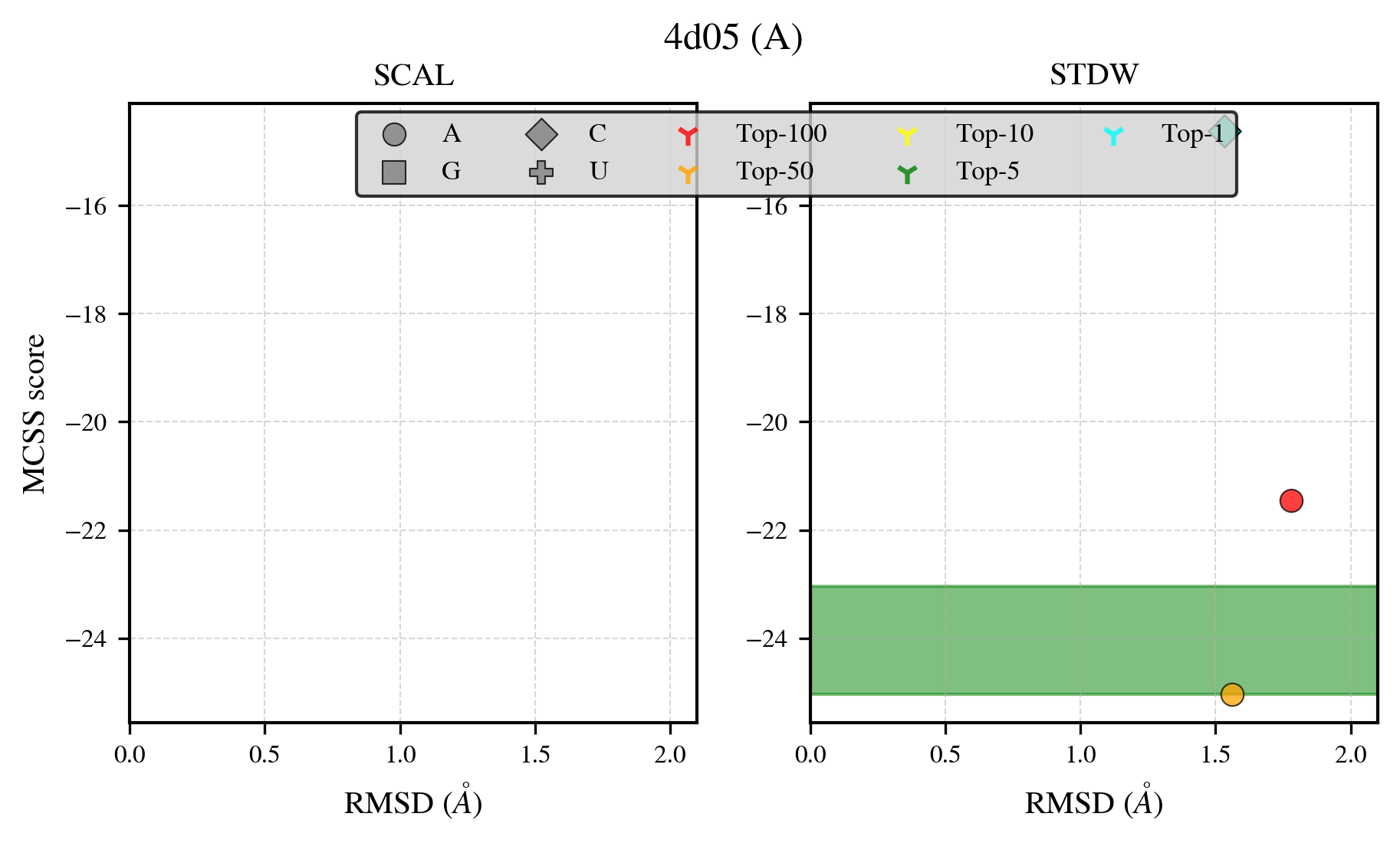

### 4d7a_benchmark-121_selectivity_details.png

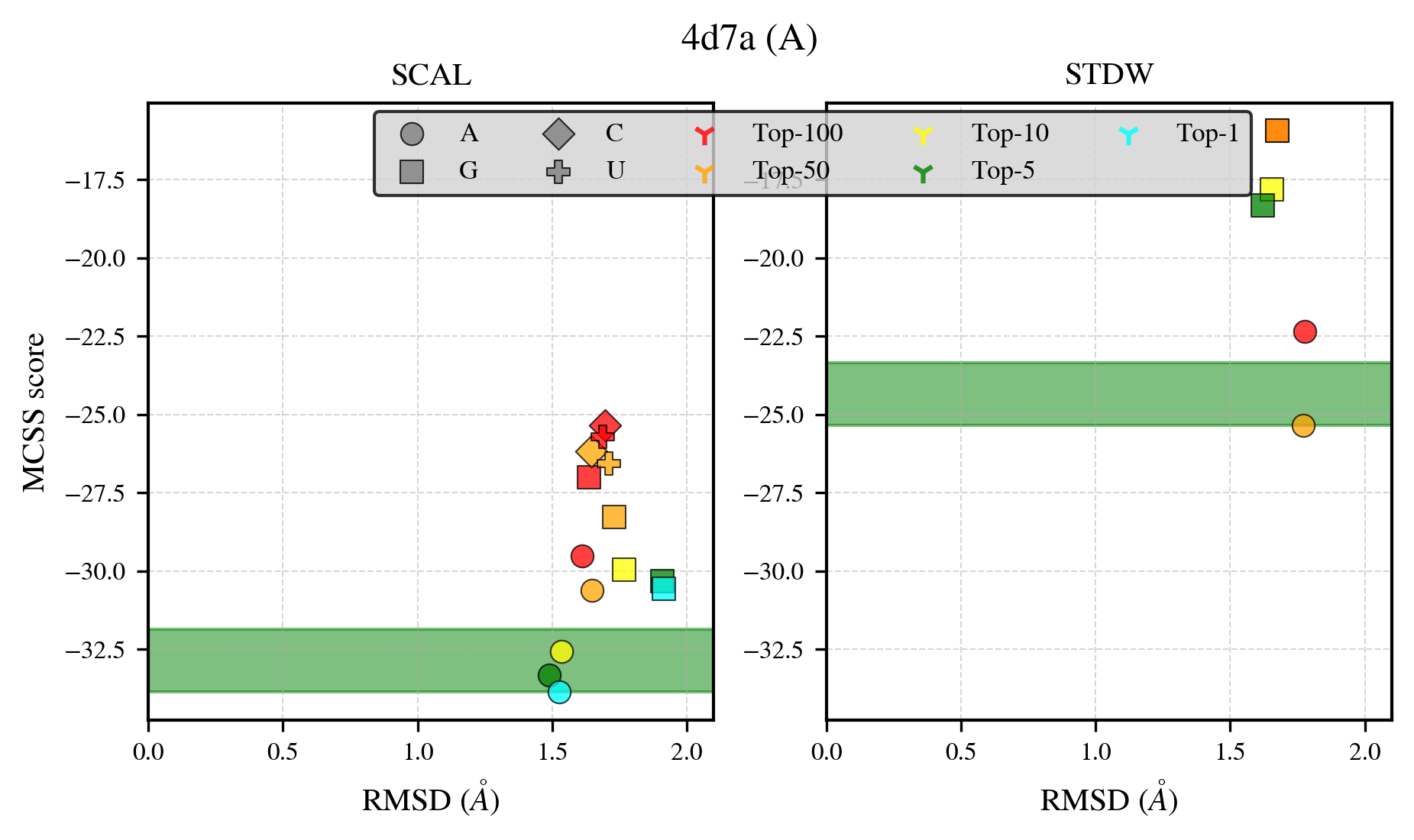

### 4eei_benchmark-121_selectivity_details.png

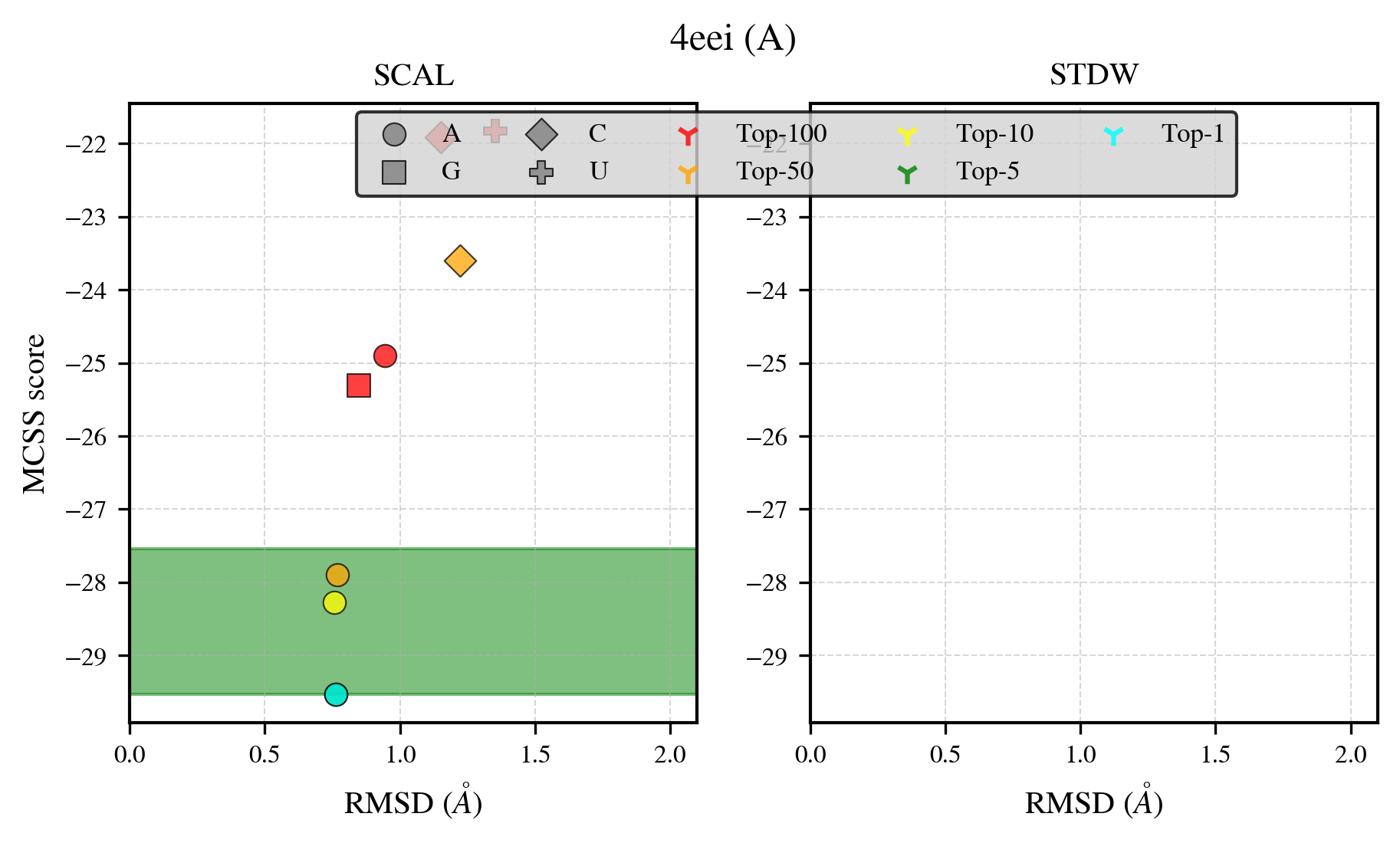

### 4emd_benchmark-121_selectivity_details.png

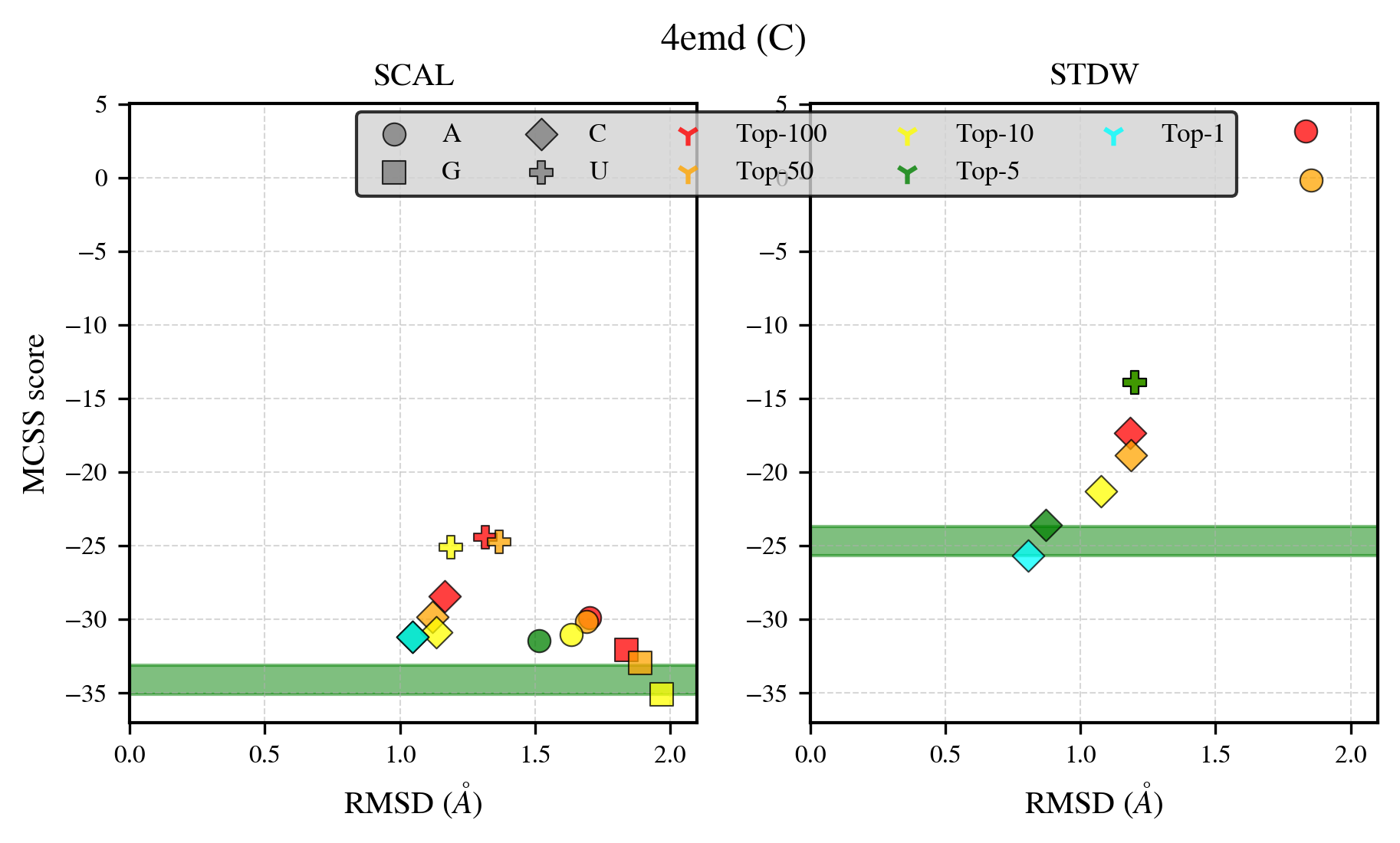

### 4eql_benchmark-121_selectivity_details.png

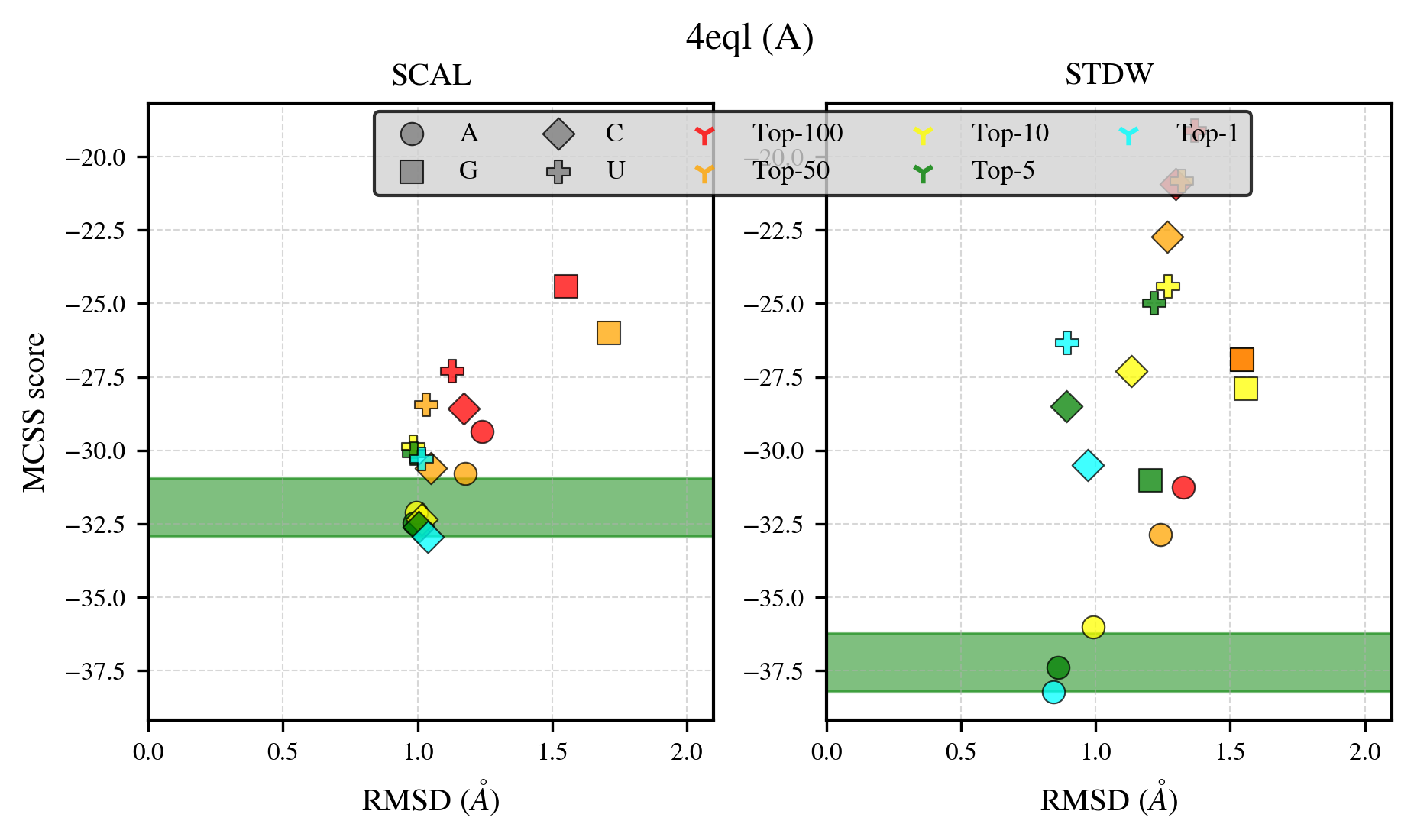

### 4fbc_benchmark-121_selectivity_details.png

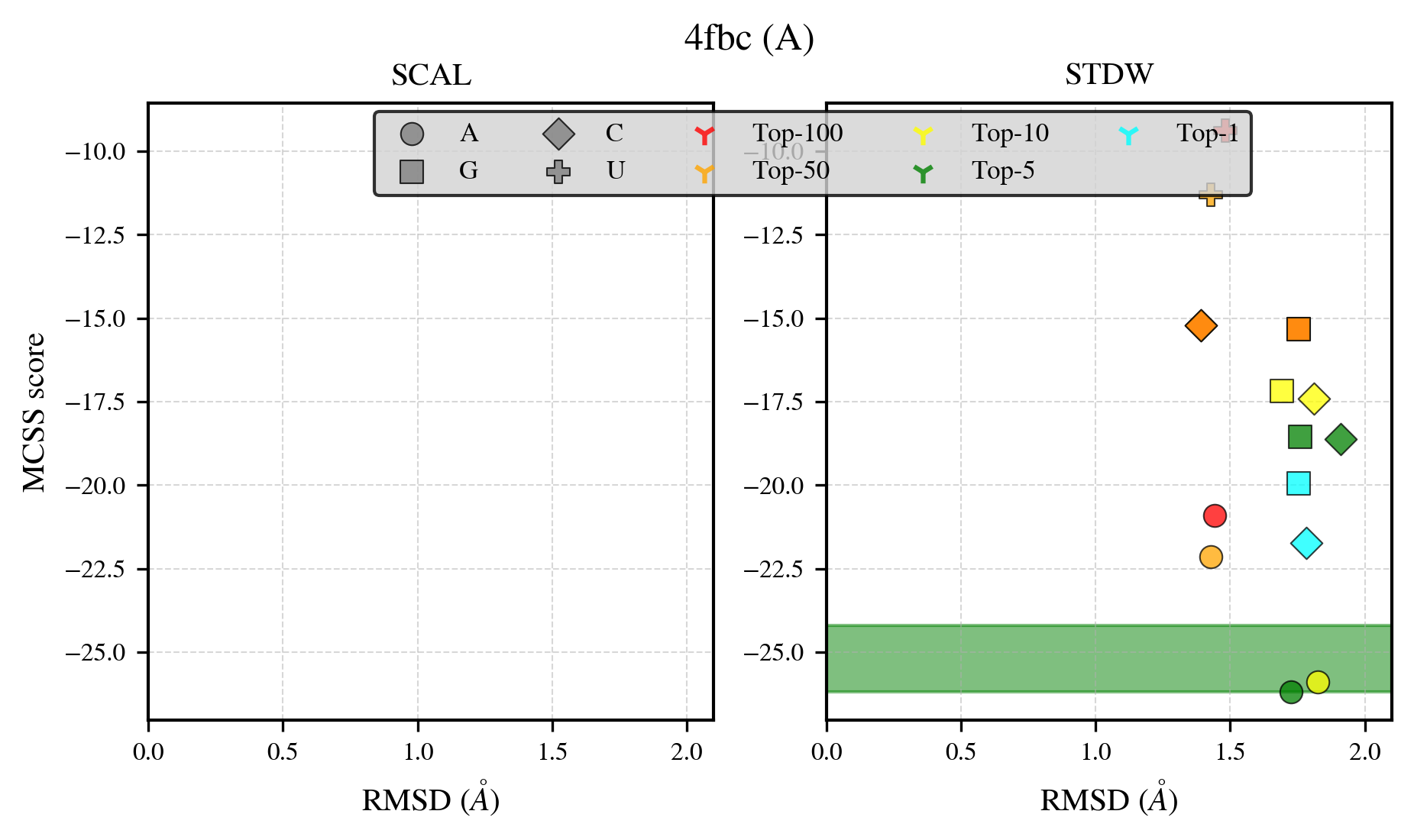

### 4fe3_benchmark-121_selectivity_details.png

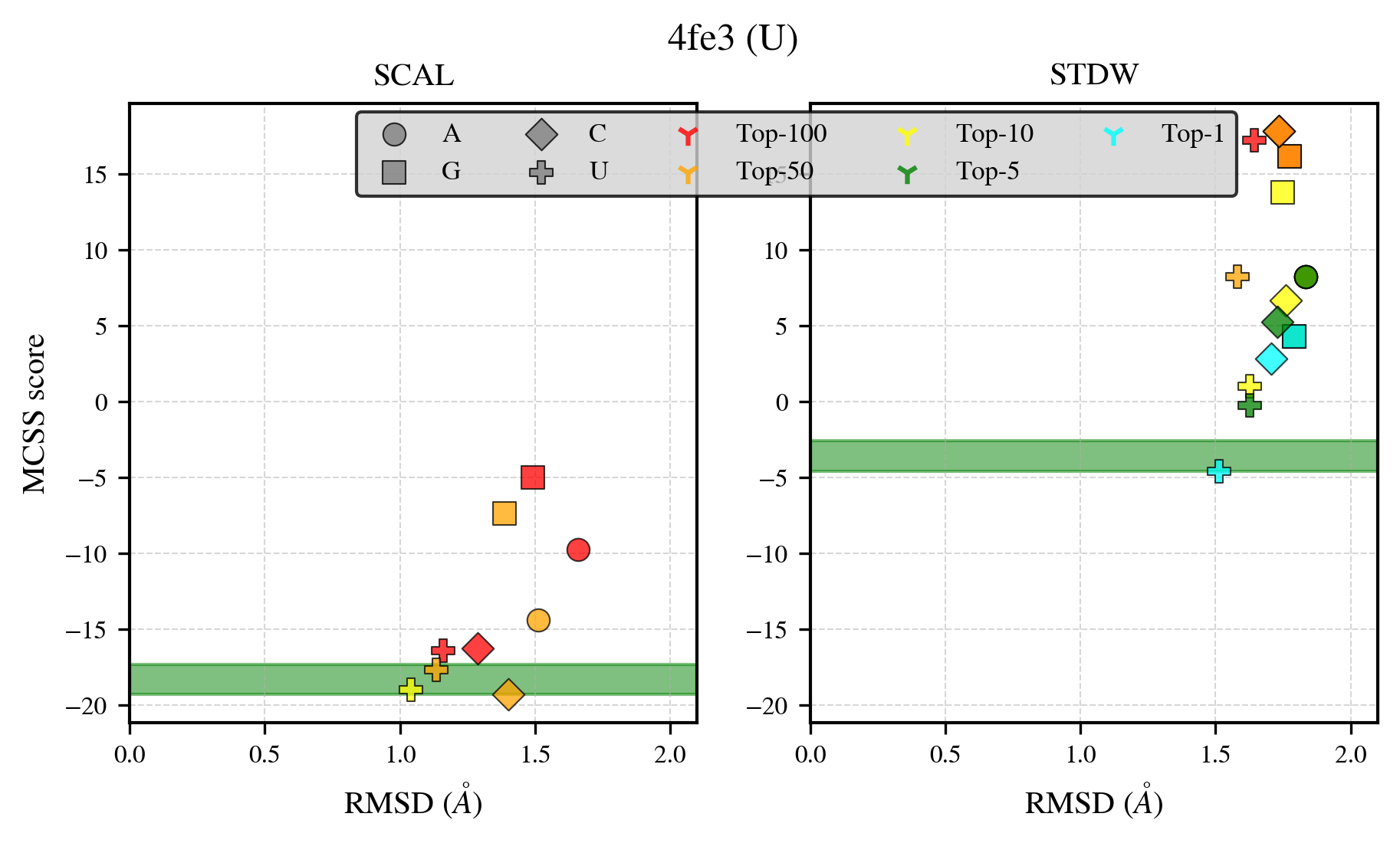

### 4h2w_benchmark-121_selectivity_details.png

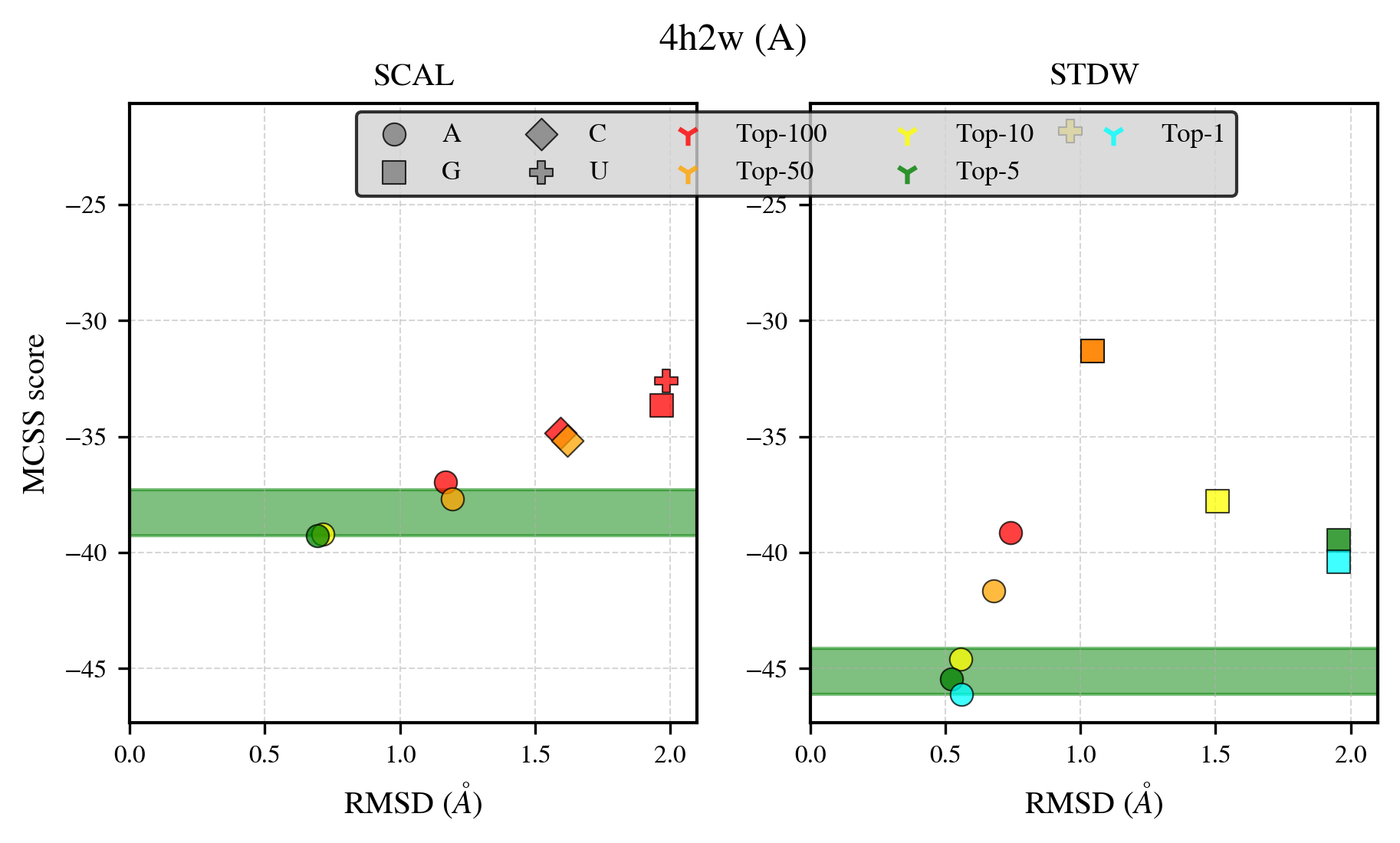

### 4he2_benchmark-121_selectivity_details.png

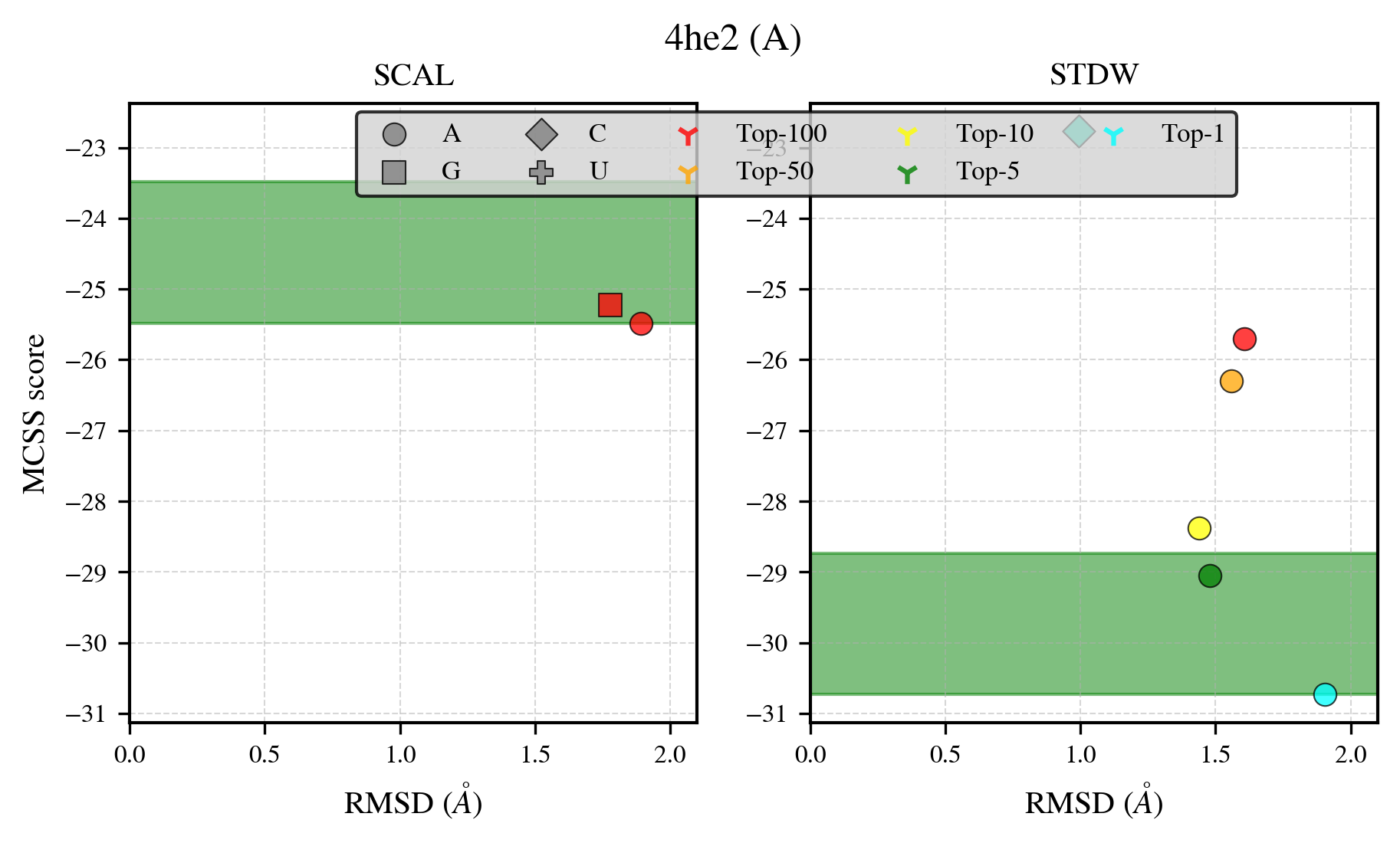

### 4ig1_benchmark-121_selectivity_details.png

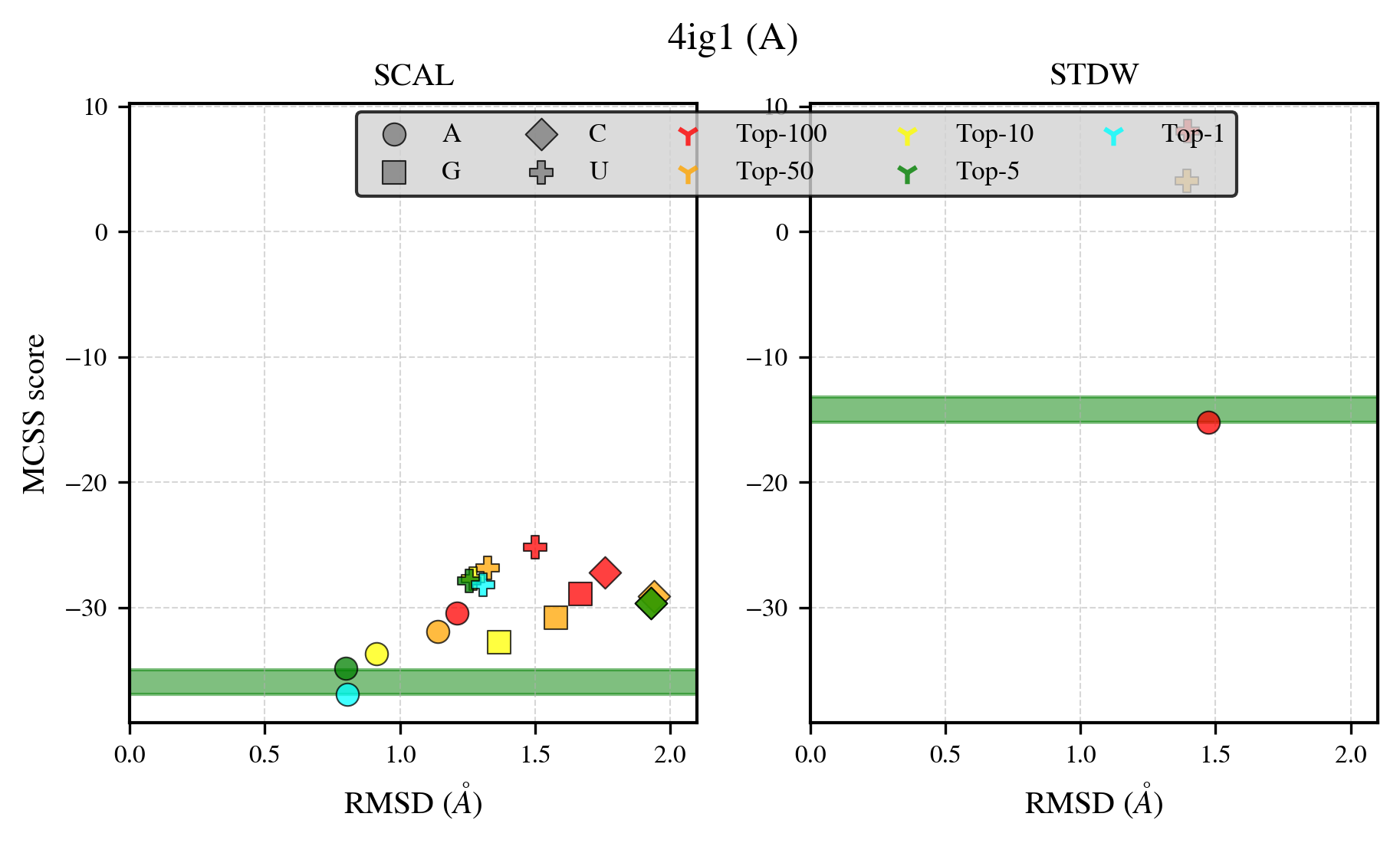

### 4ijn_benchmark-121_selectivity_details.png

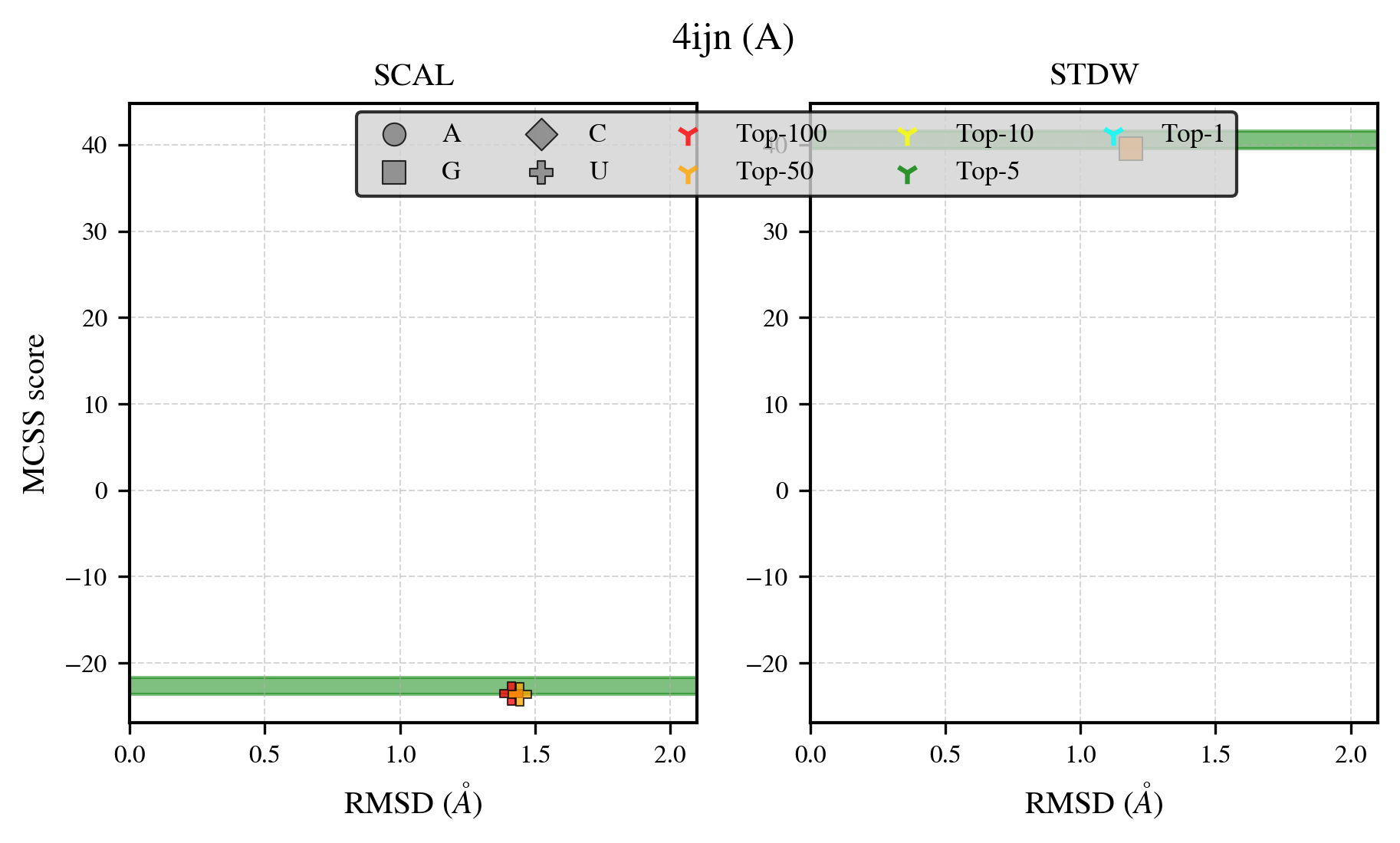

### 4ike_benchmark-121_selectivity_details.png

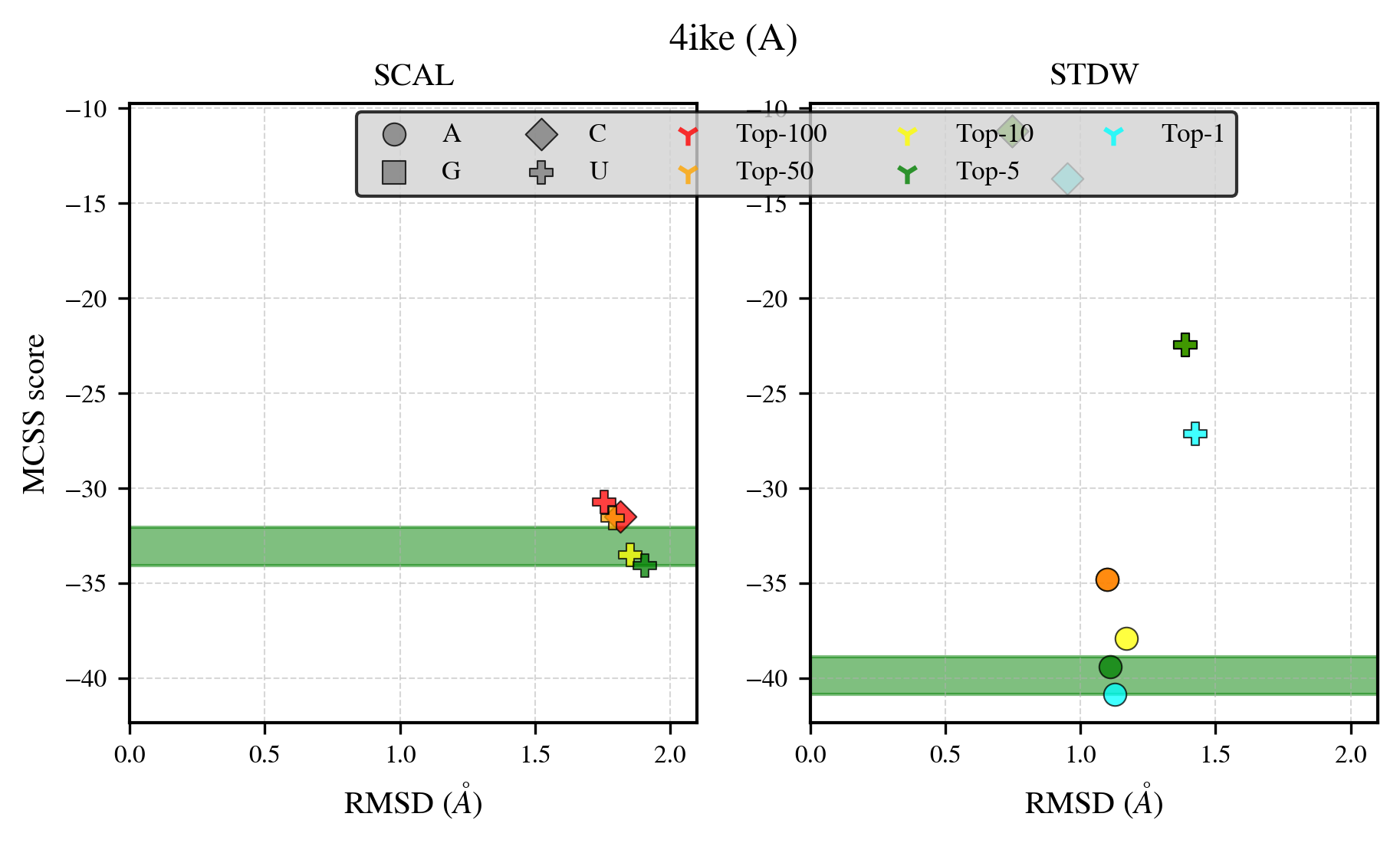

### 4jem_benchmark-121_selectivity_details.png

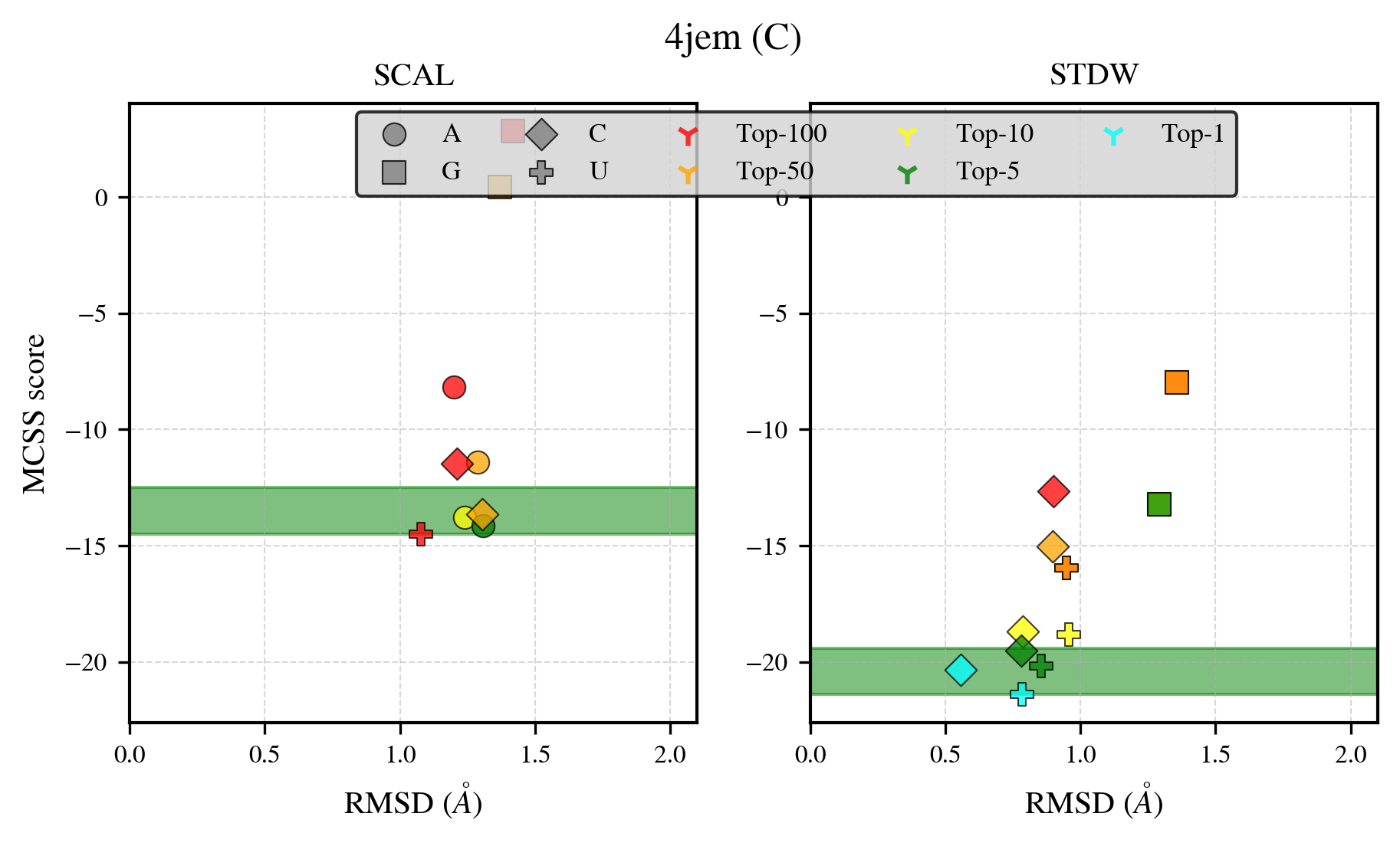

### 4kbf_benchmark-121_selectivity_details.png

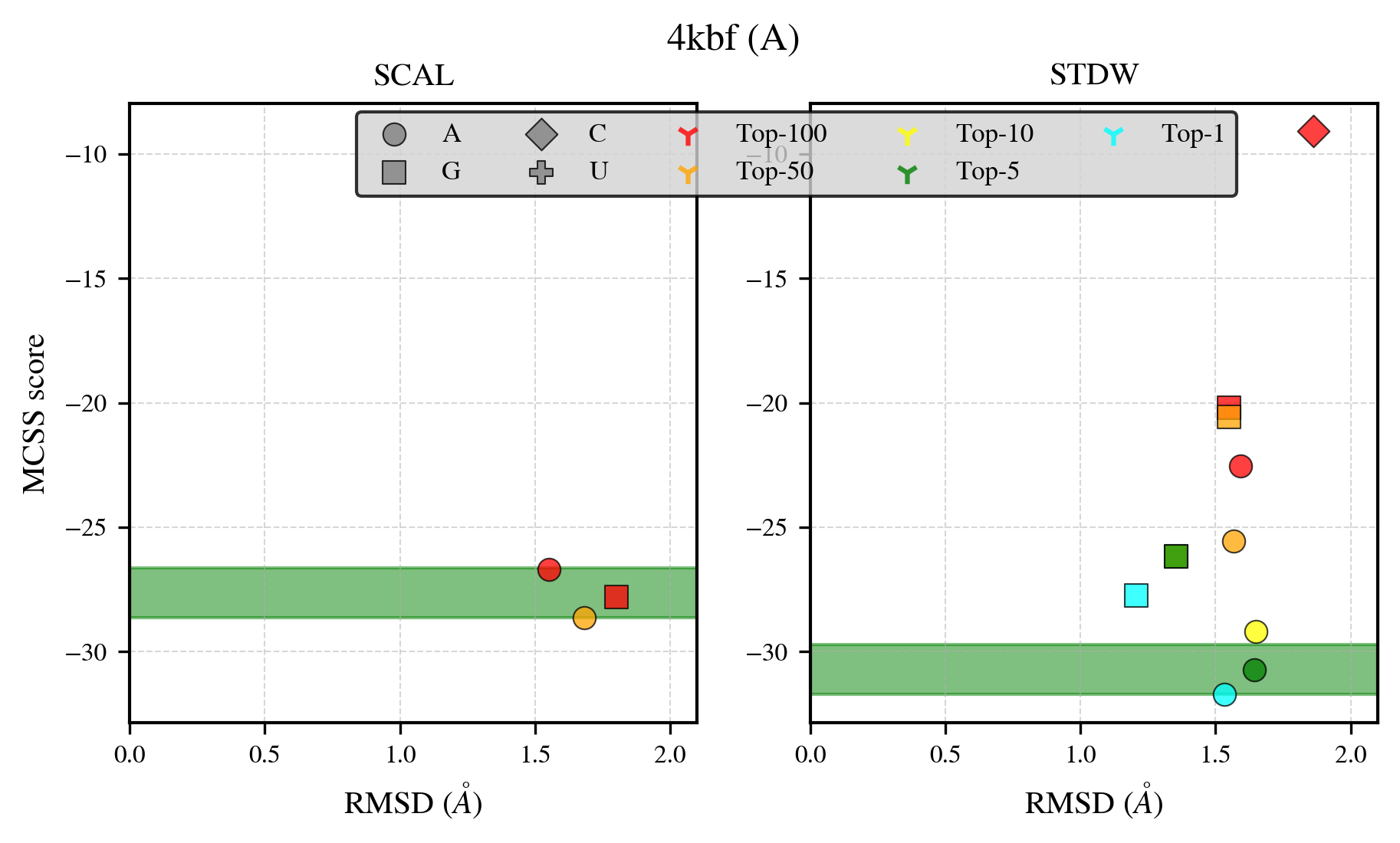

### 4m0k_benchmark-121_selectivity_details.png

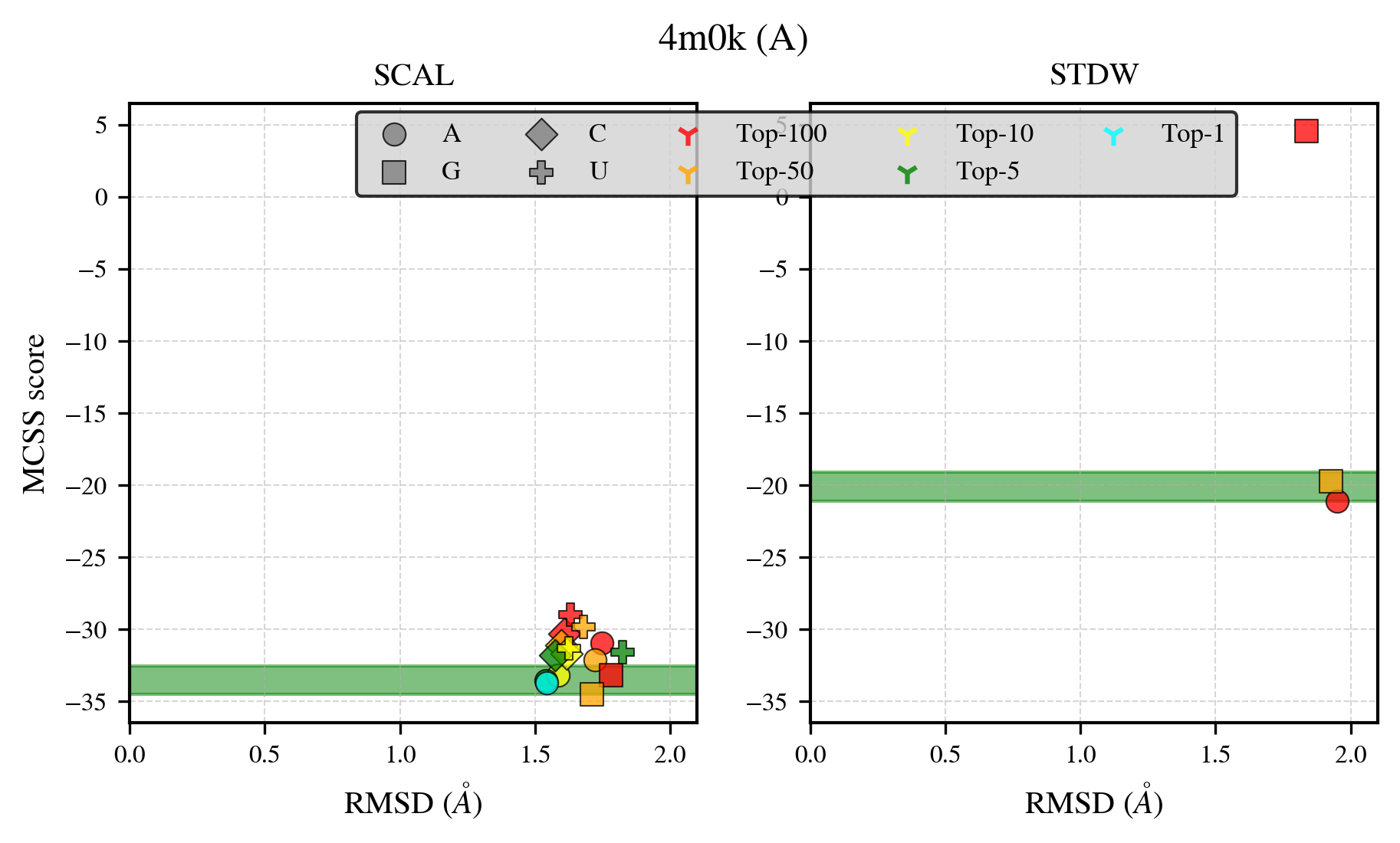

### 4m9d_benchmark-121_selectivity_details.png

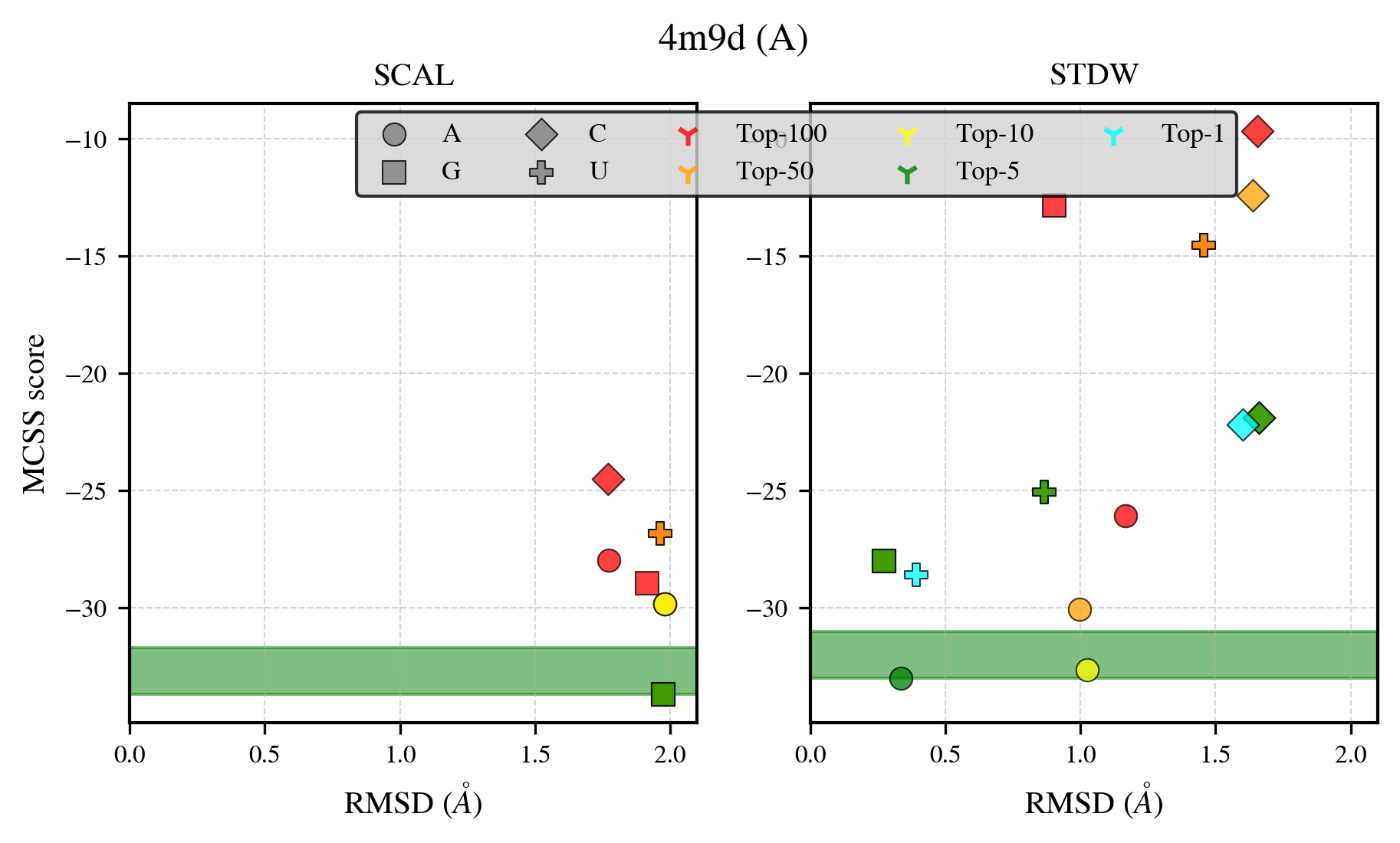

### 4ma0_benchmark-121_selectivity_details.png

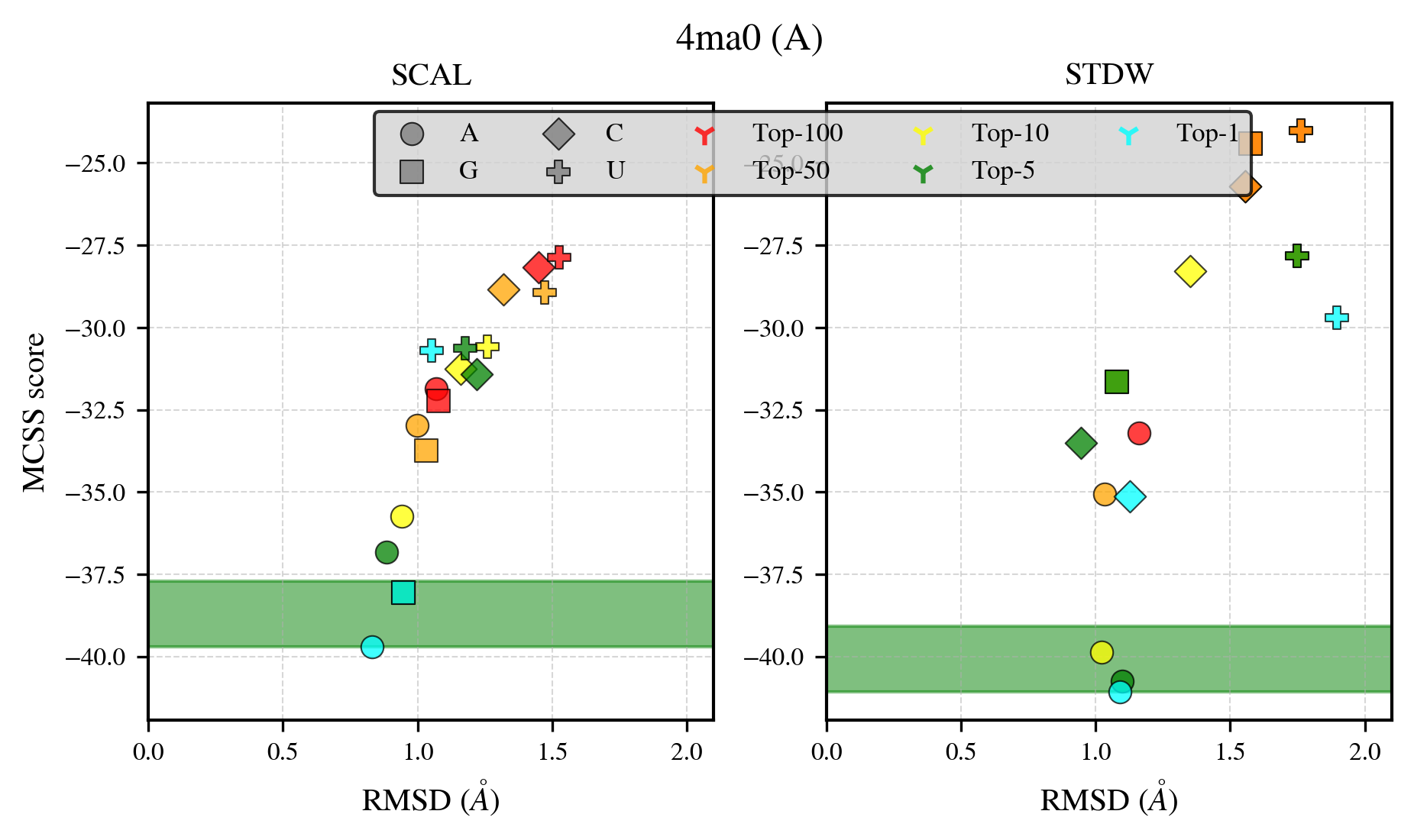

### 4mpo_benchmark-121_selectivity_details.png

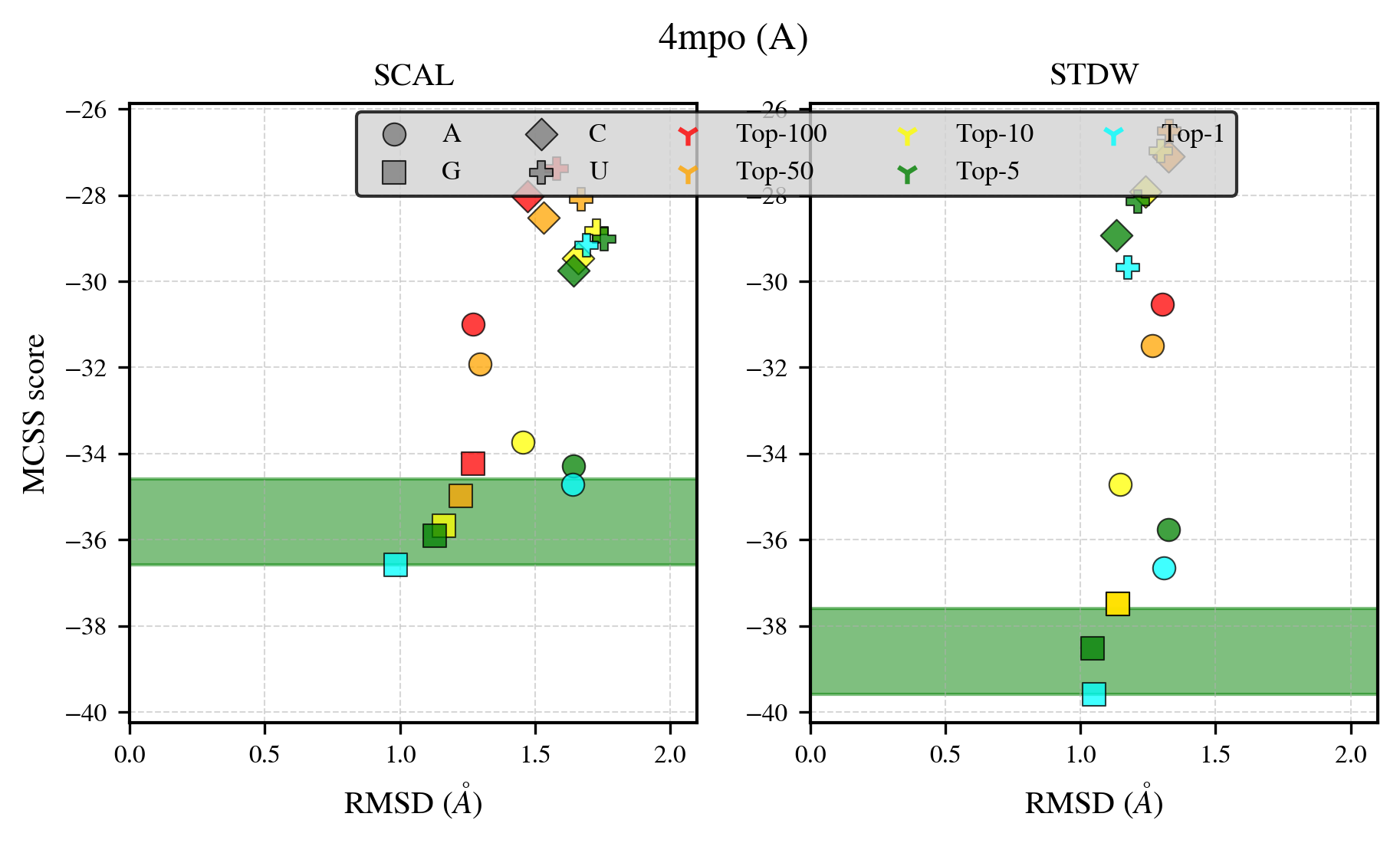

### 4ndf_benchmark-121_selectivity_details.png

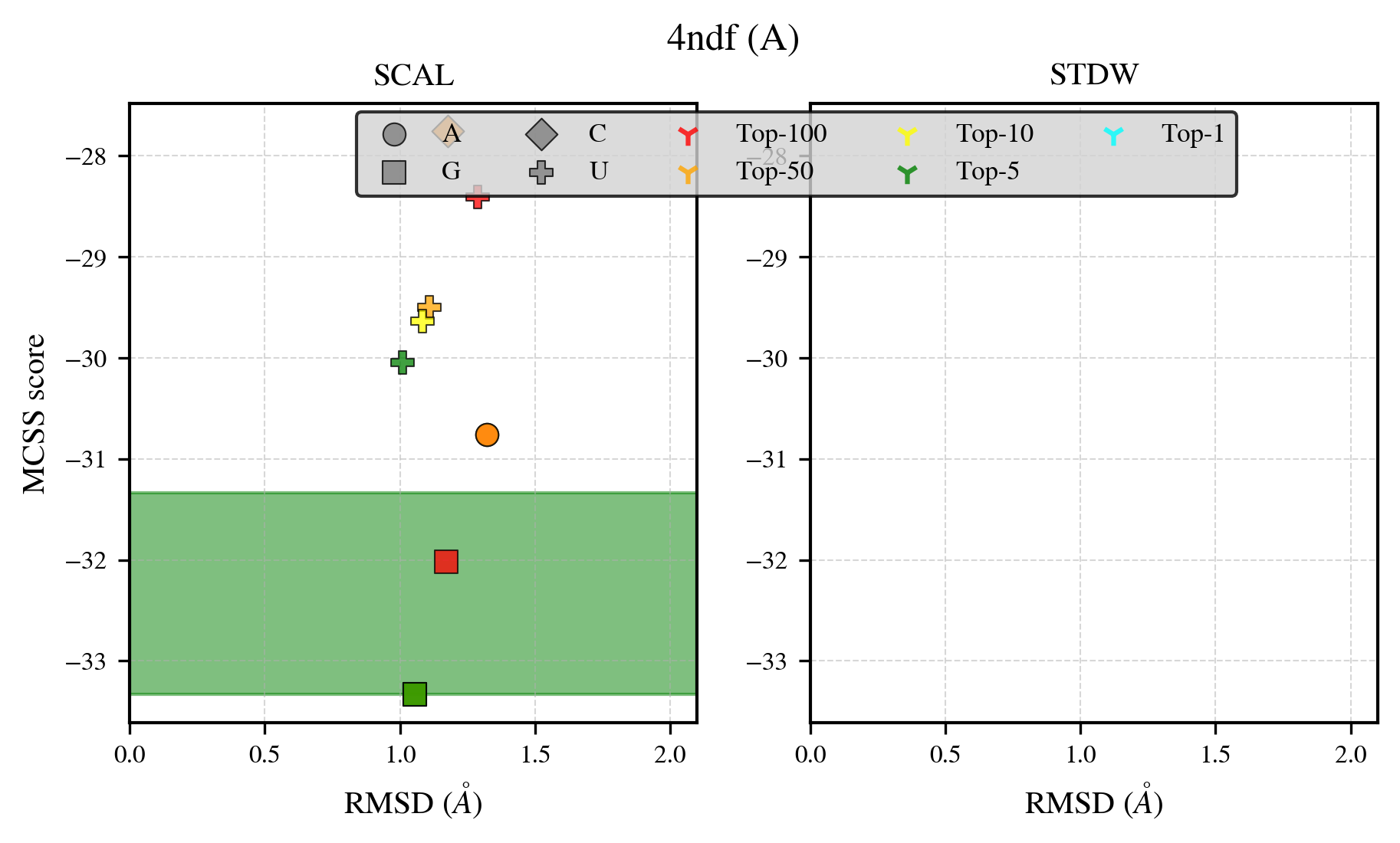

### 4o6m_benchmark-121_selectivity_details.png

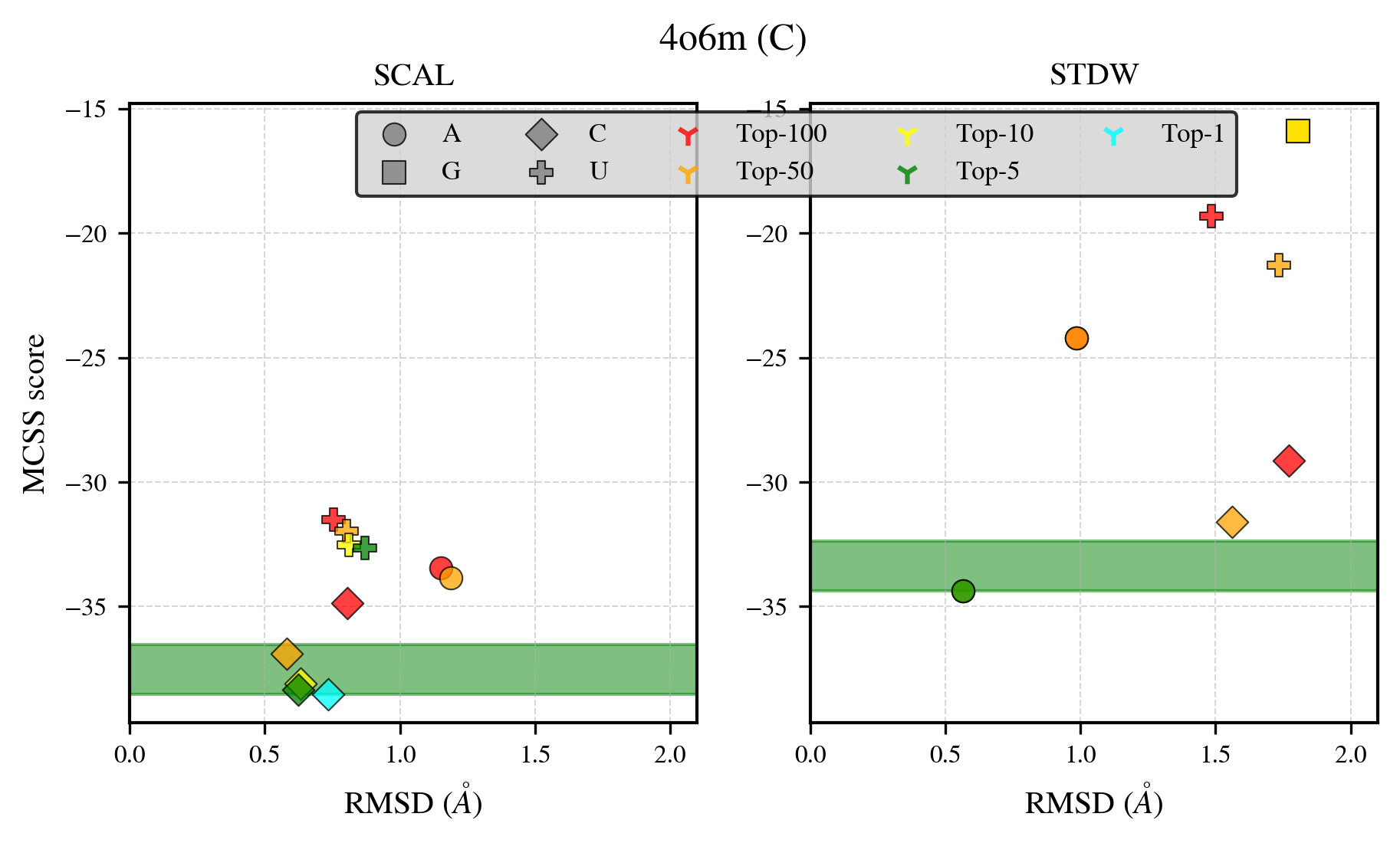

### 4ozl_benchmark-121_selectivity_details.png

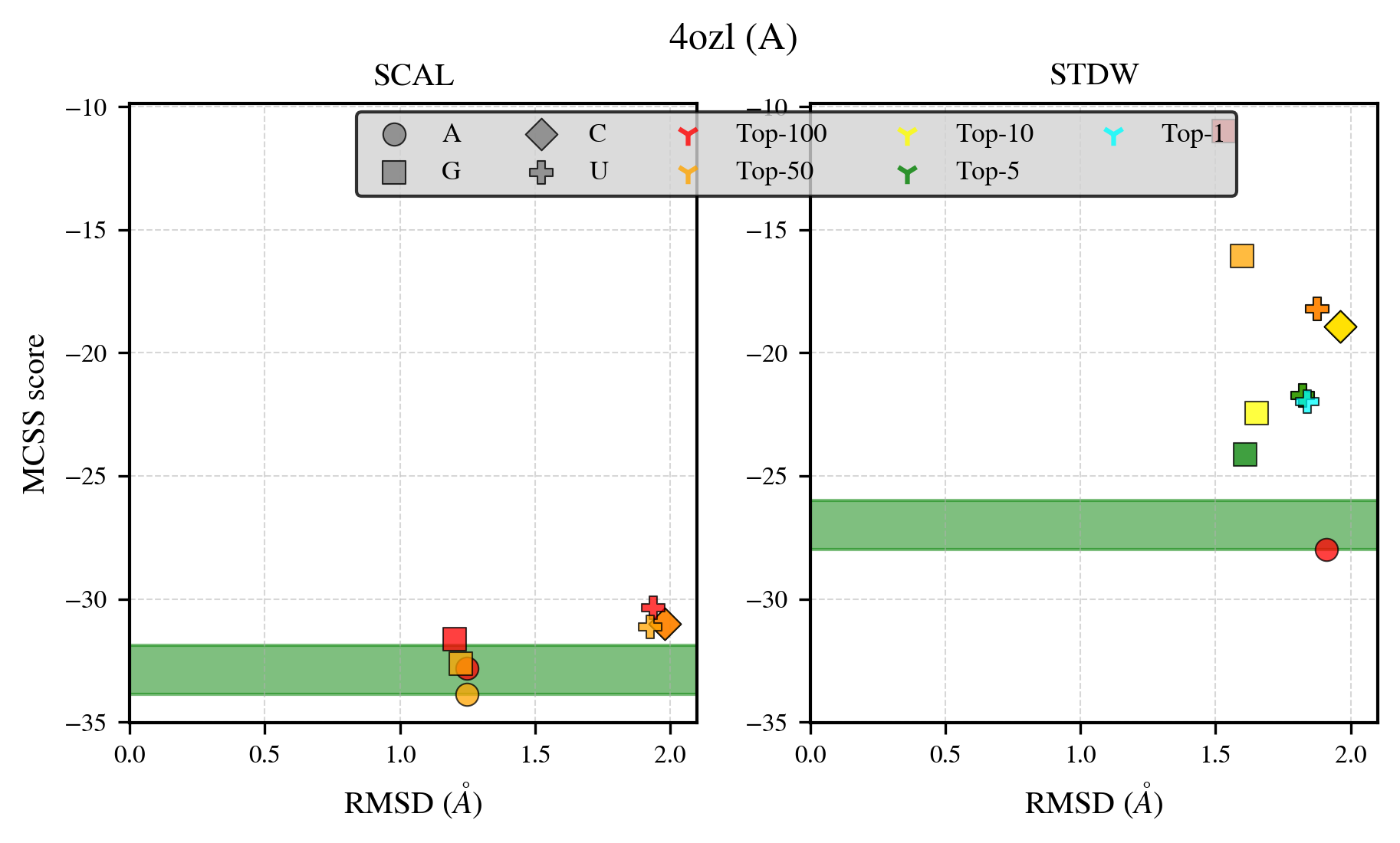

### 4r78_benchmark-121_selectivity_details.png

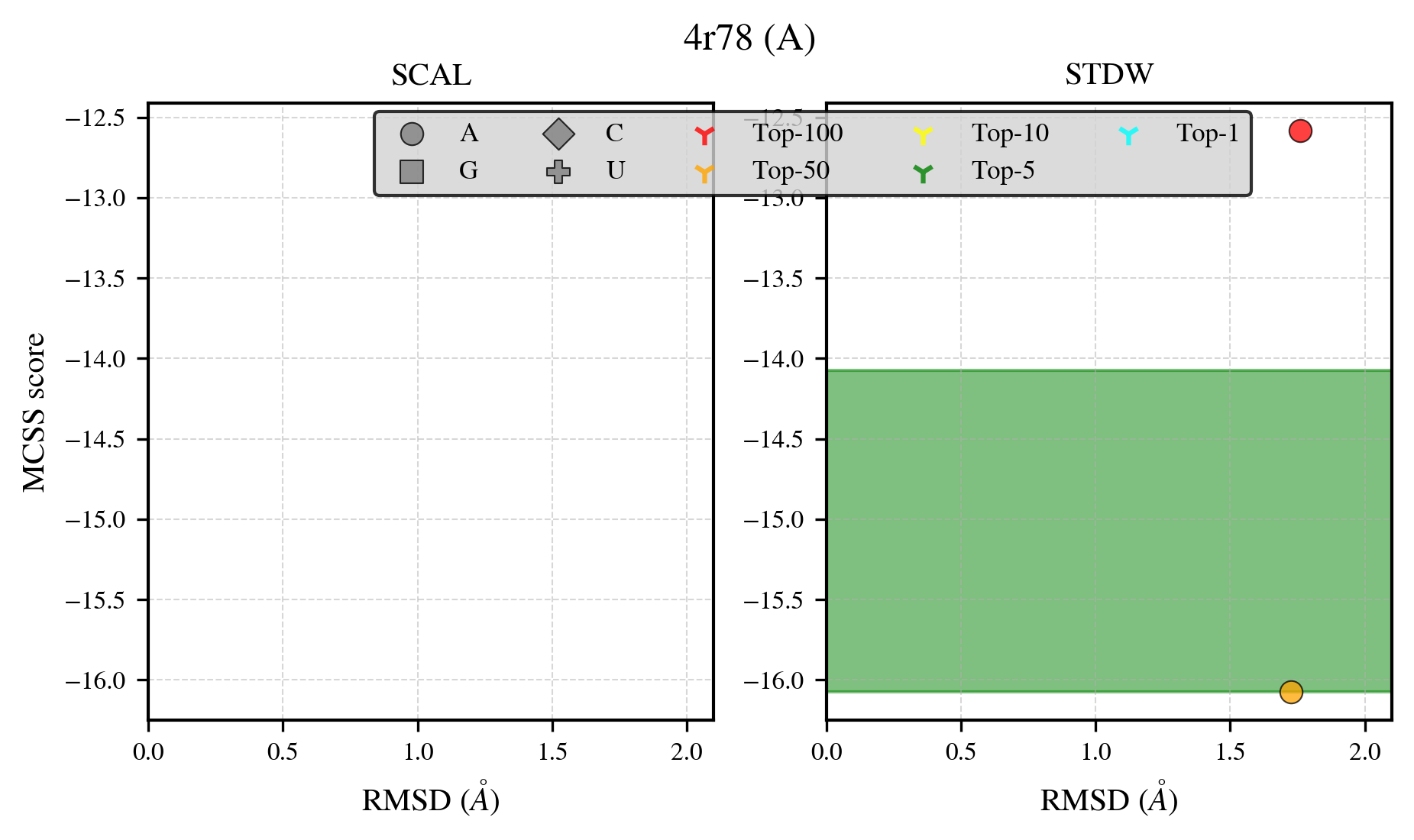

### 4ww7_benchmark-121_selectivity_details.png

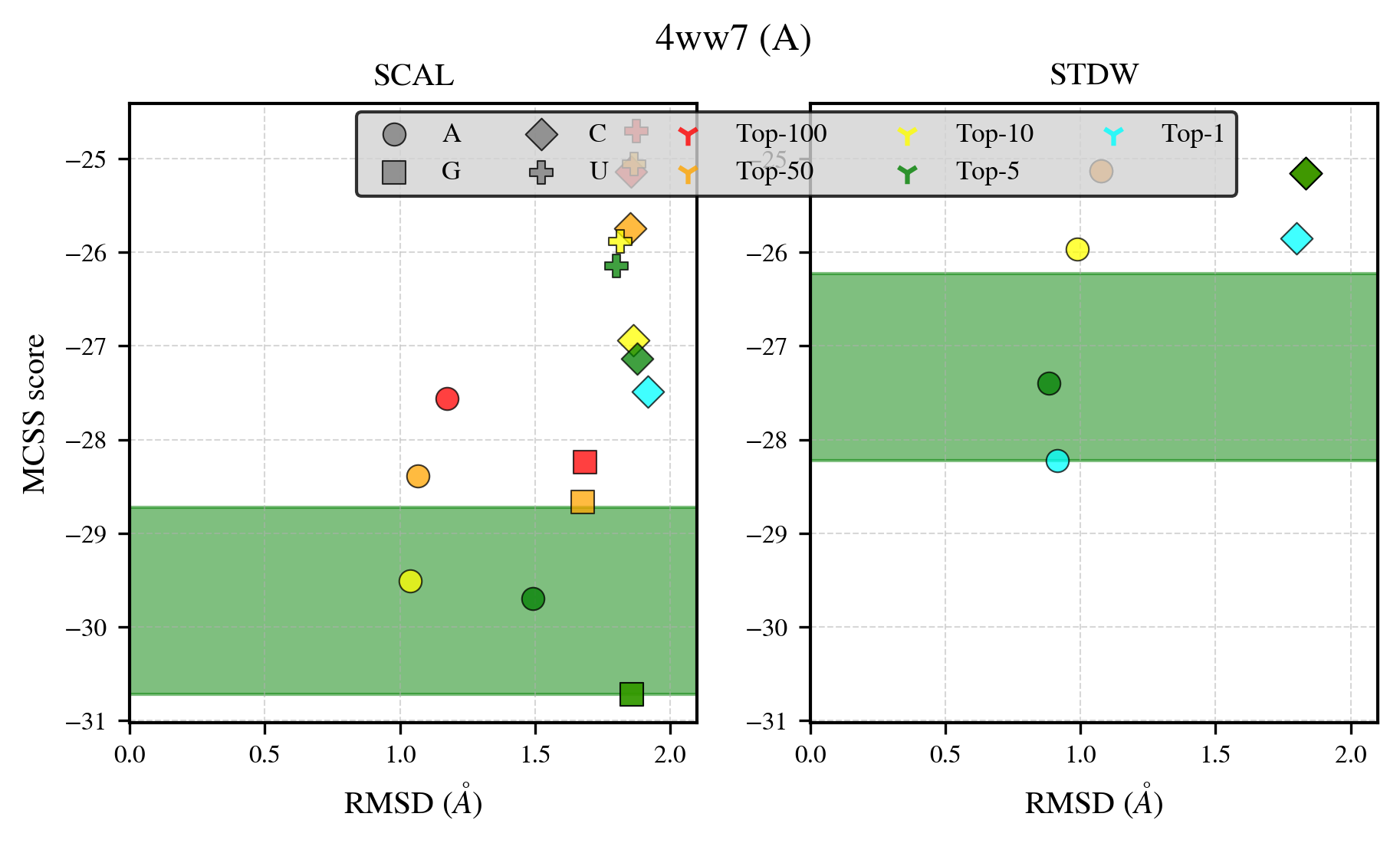

### 4x9d_benchmark-121_selectivity_details.png

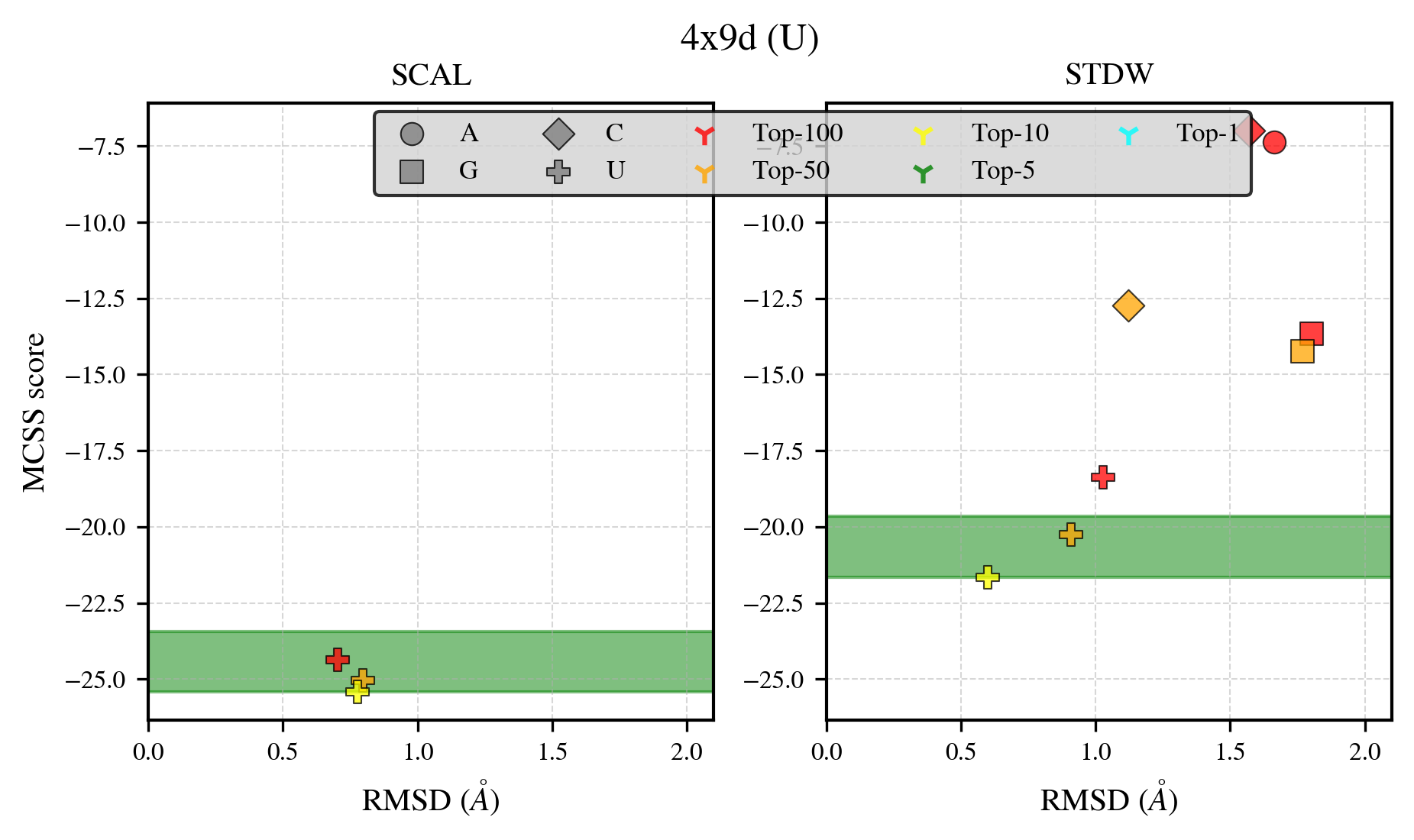

### 4zcp_benchmark-121_selectivity_details.png

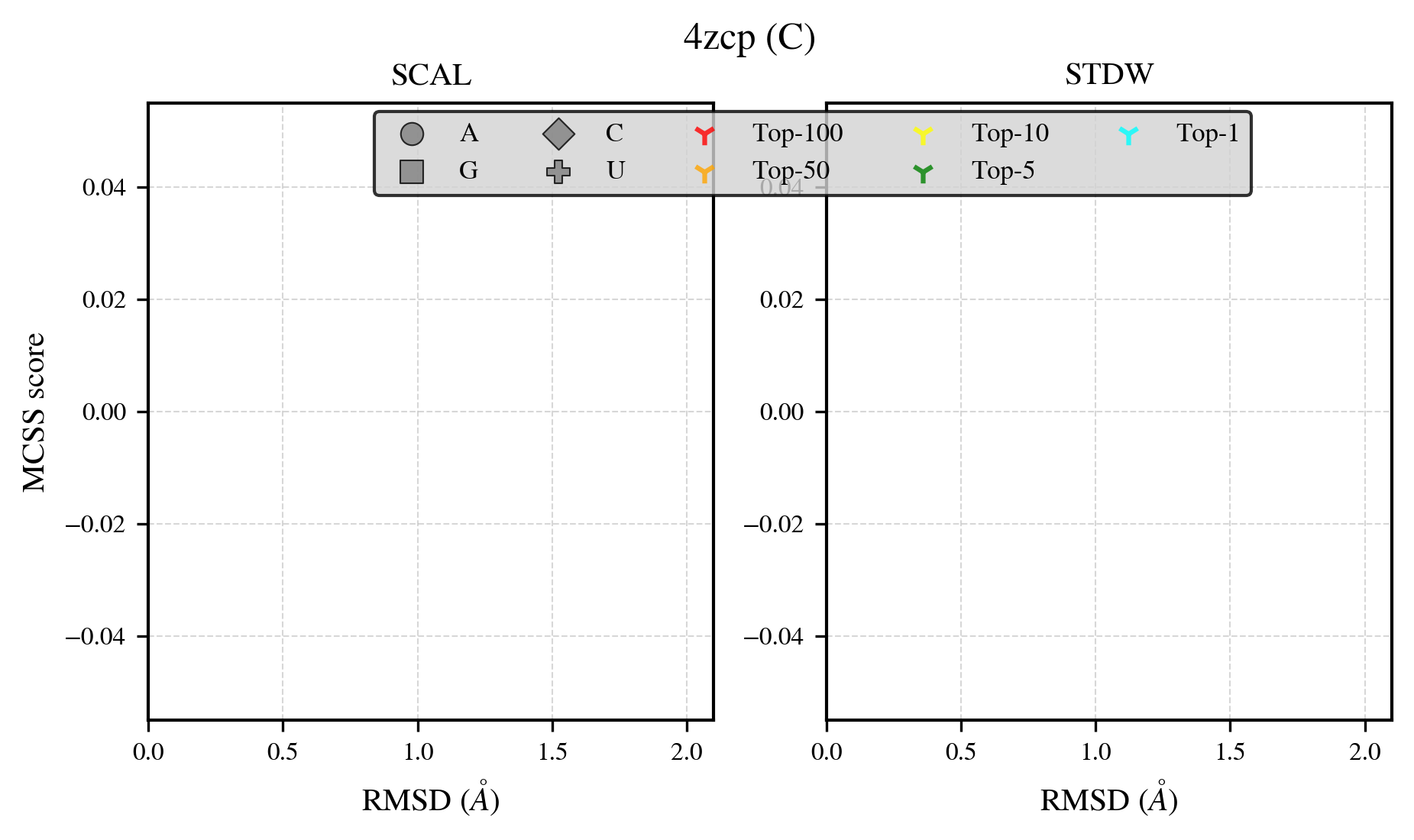

### 4zfn_benchmark-121_selectivity_details.png

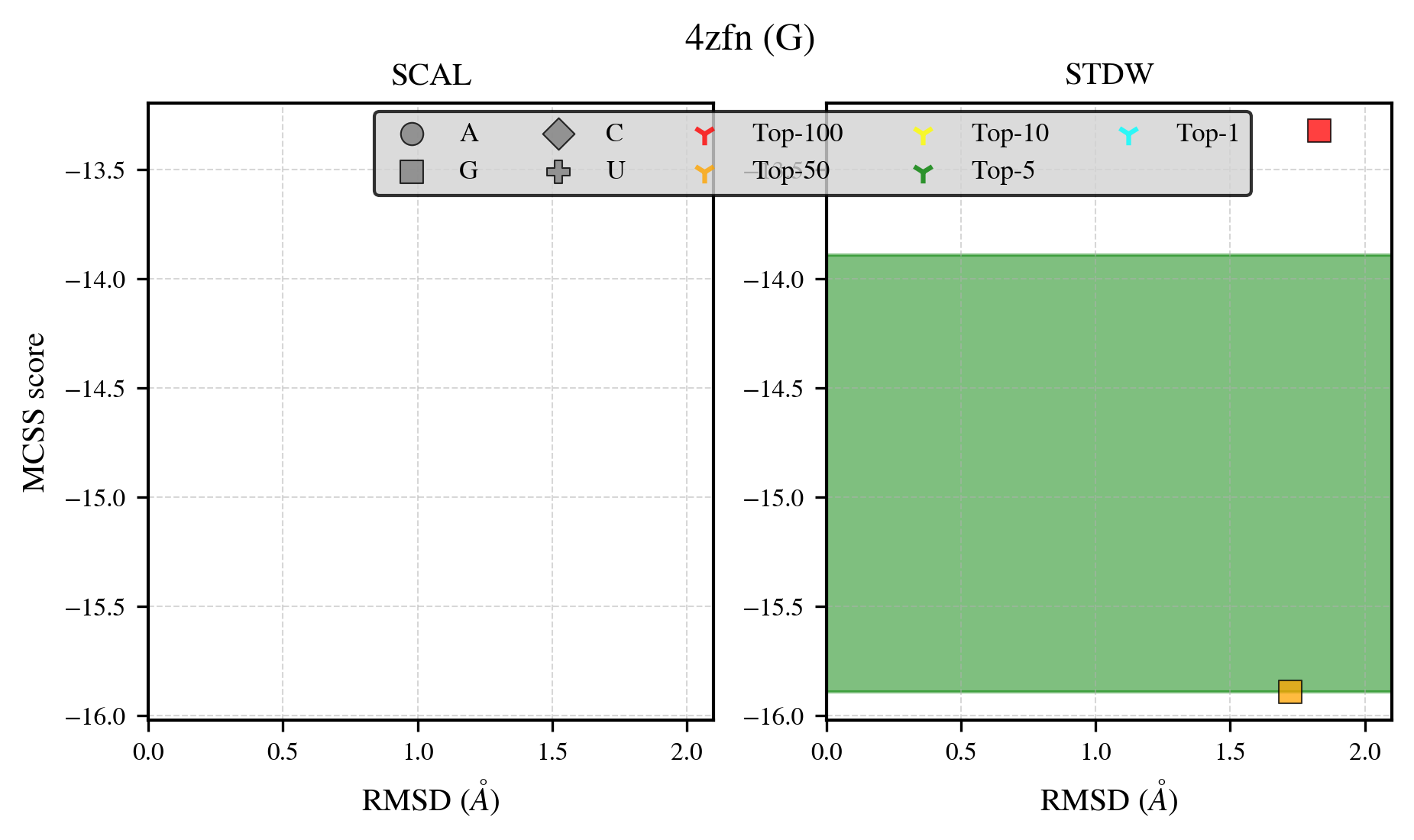

### 5b6d_benchmark-121_selectivity_details.png

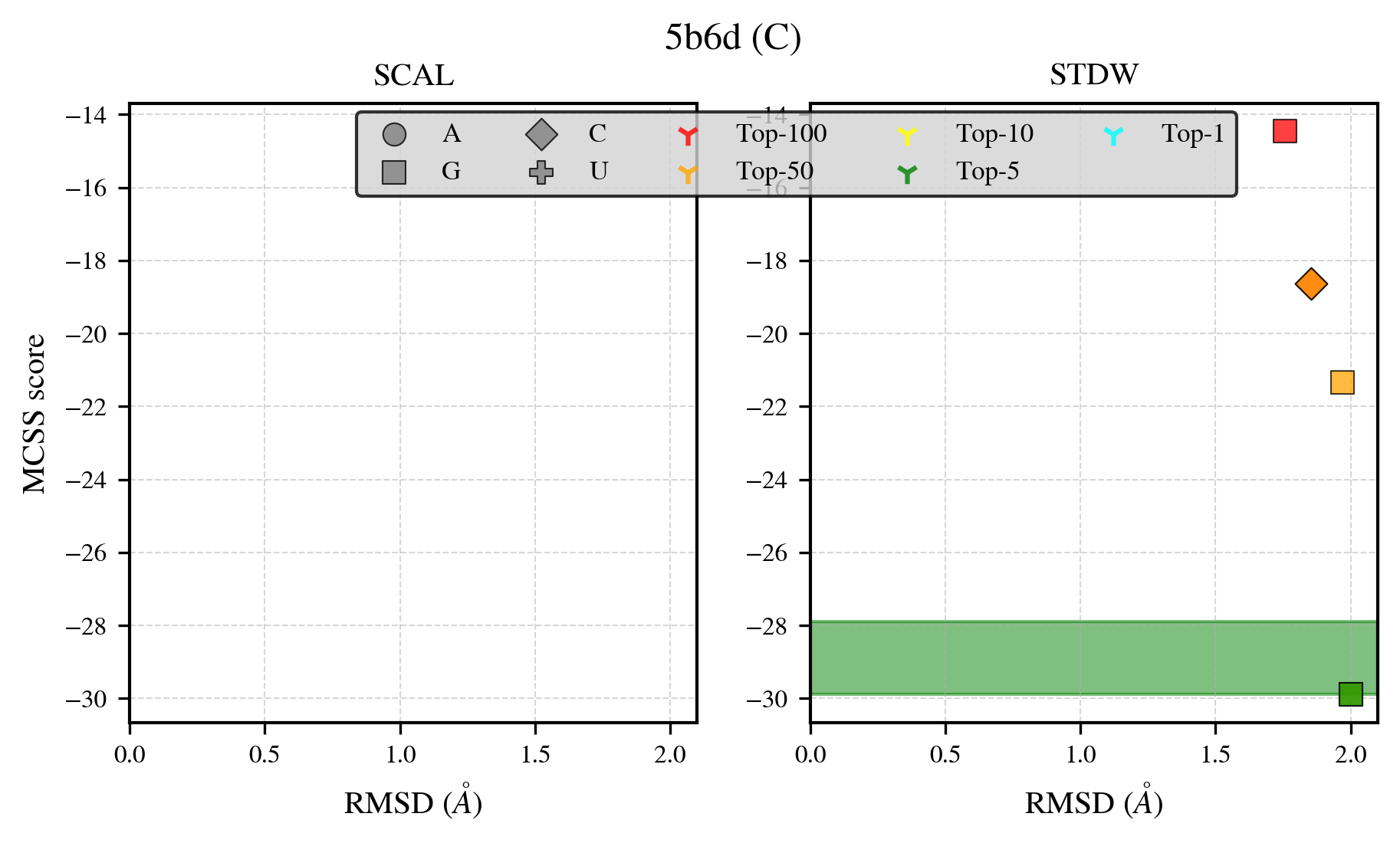

### 5b8f_benchmark-121_selectivity_details.png

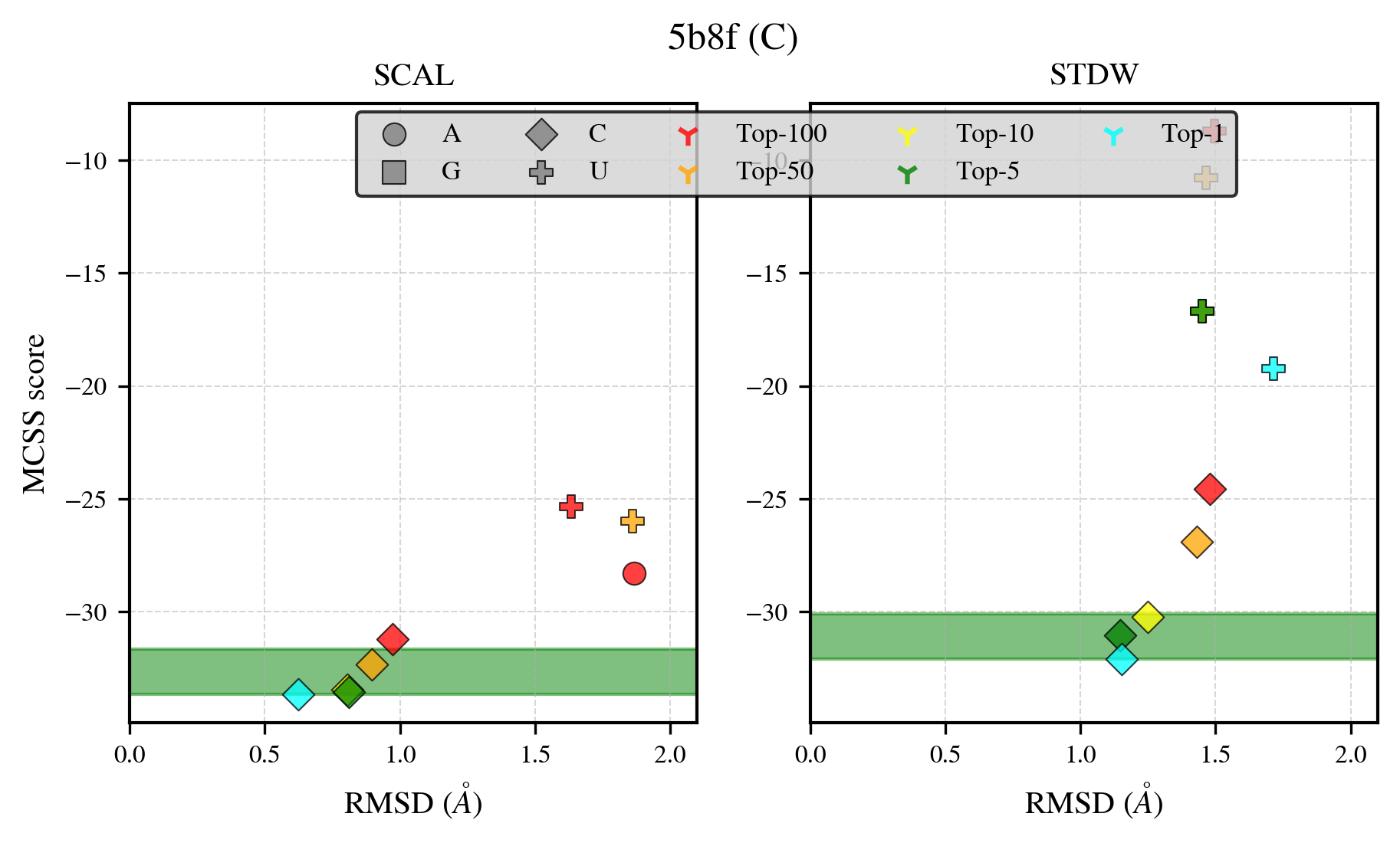

### 5bph_benchmark-121_selectivity_details.png

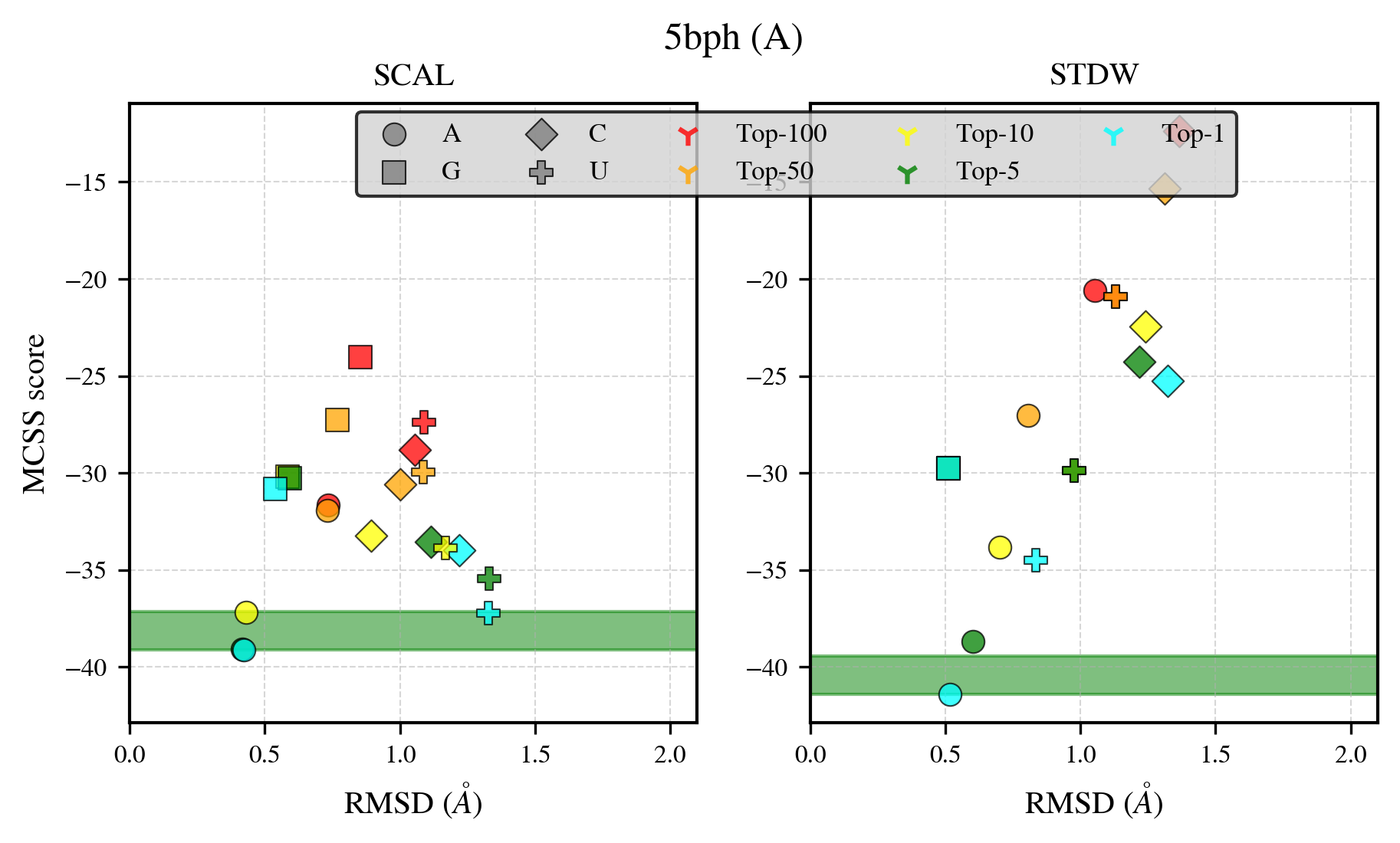

### 5cot_benchmark-121_selectivity_details.png

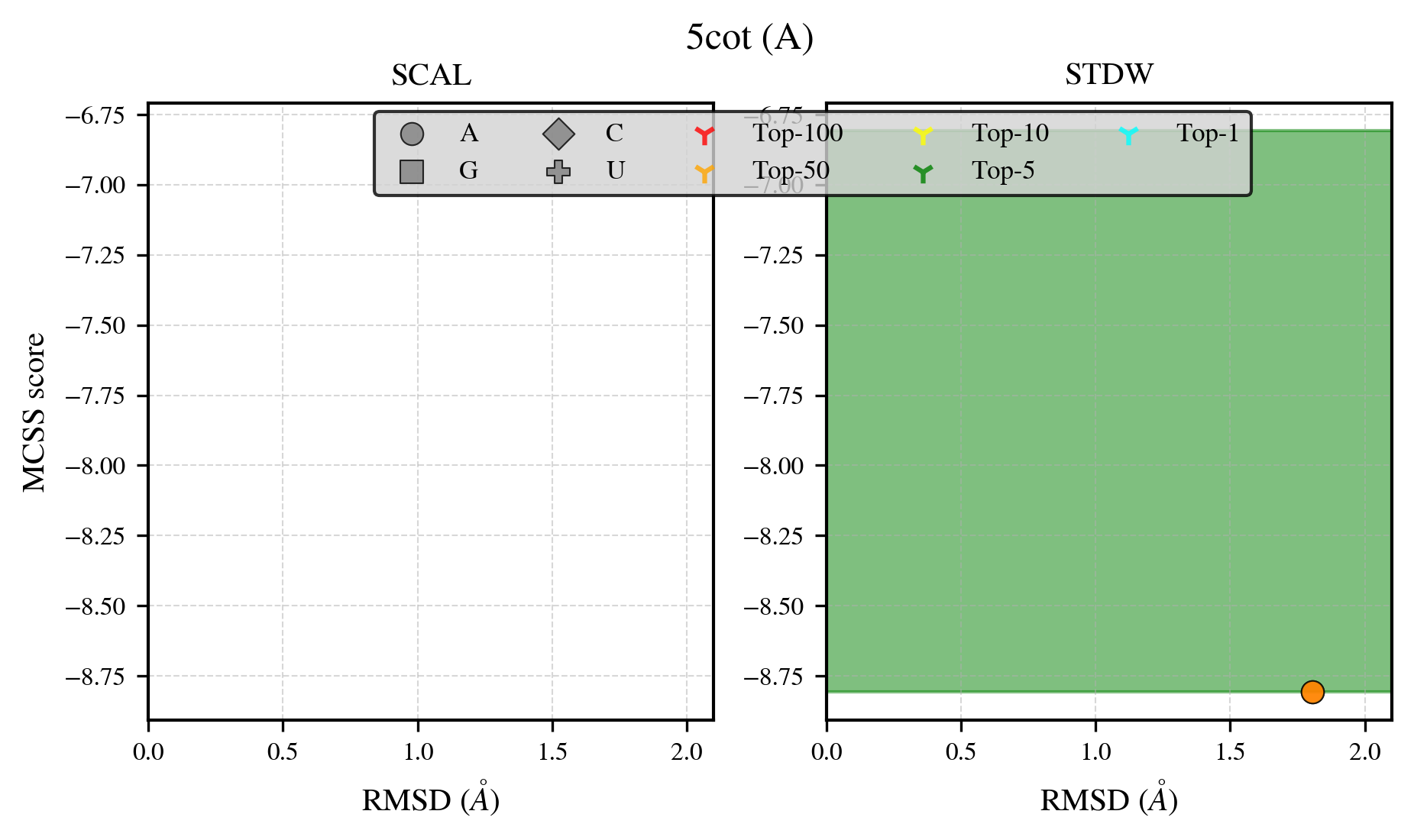

### 5d4n_benchmark-121_selectivity_details.png

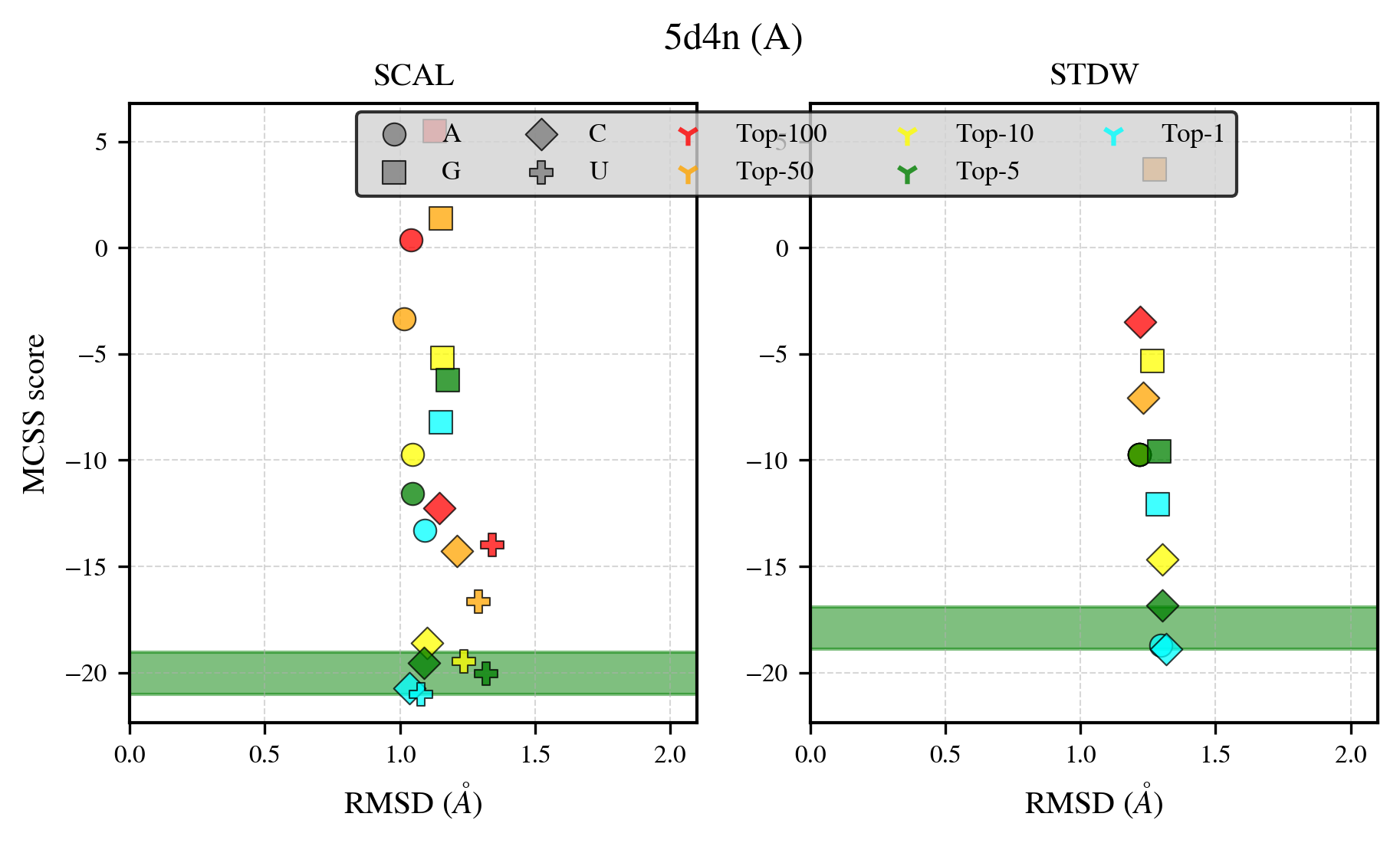

### 5ed3_benchmark-121_selectivity_details.png

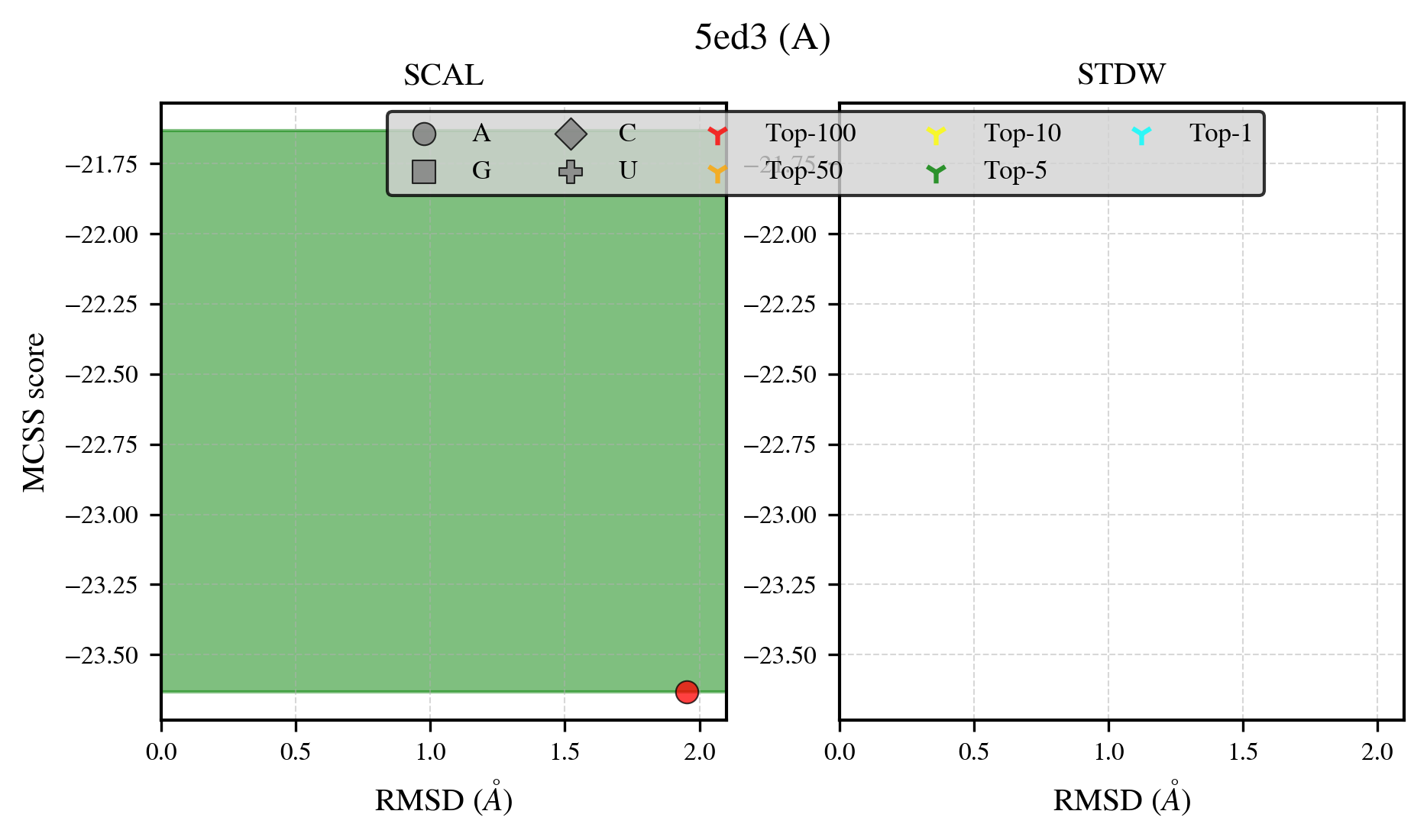

### 5gmd_benchmark-121_selectivity_details.png

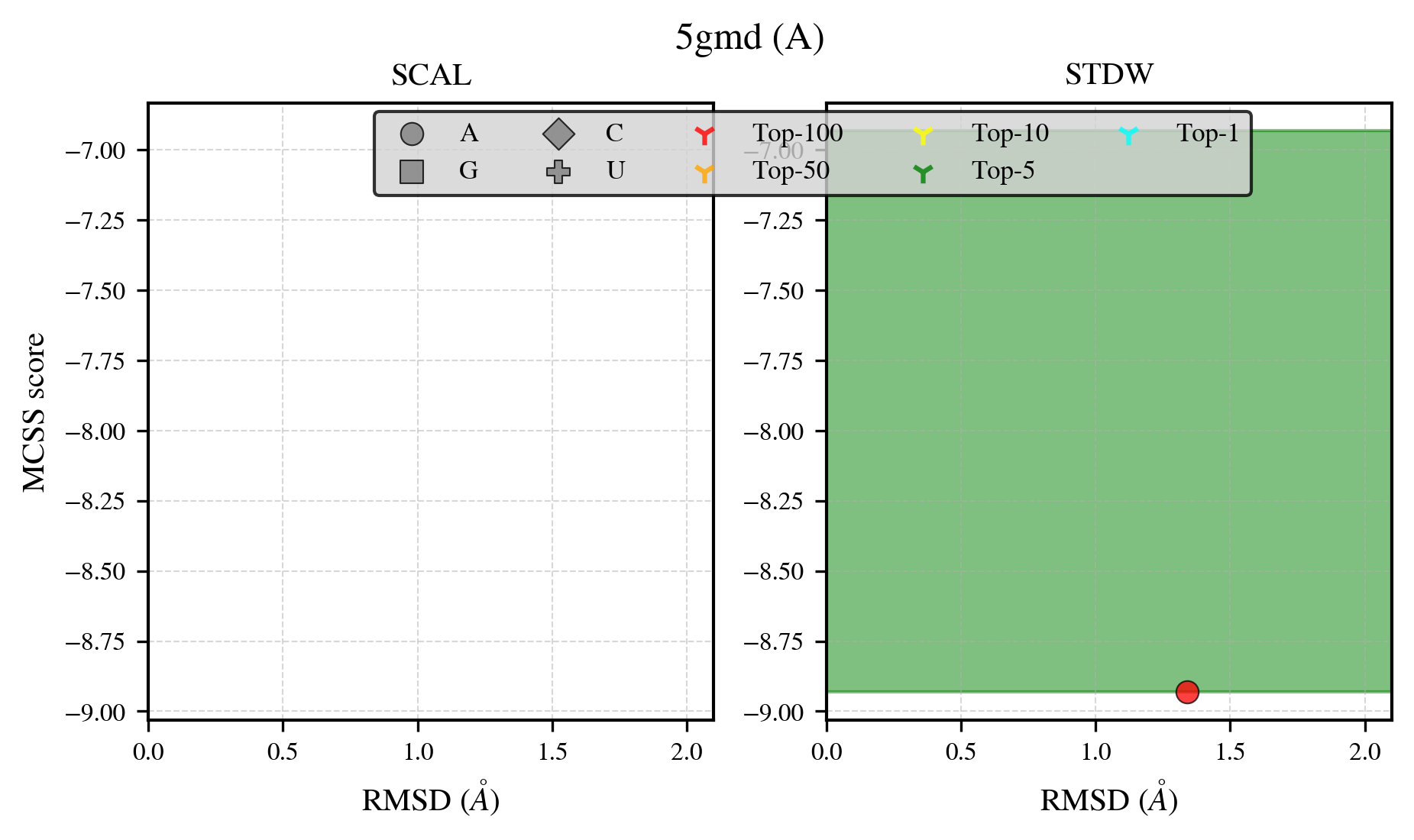

### 5k1d_benchmark-121_selectivity_details.png

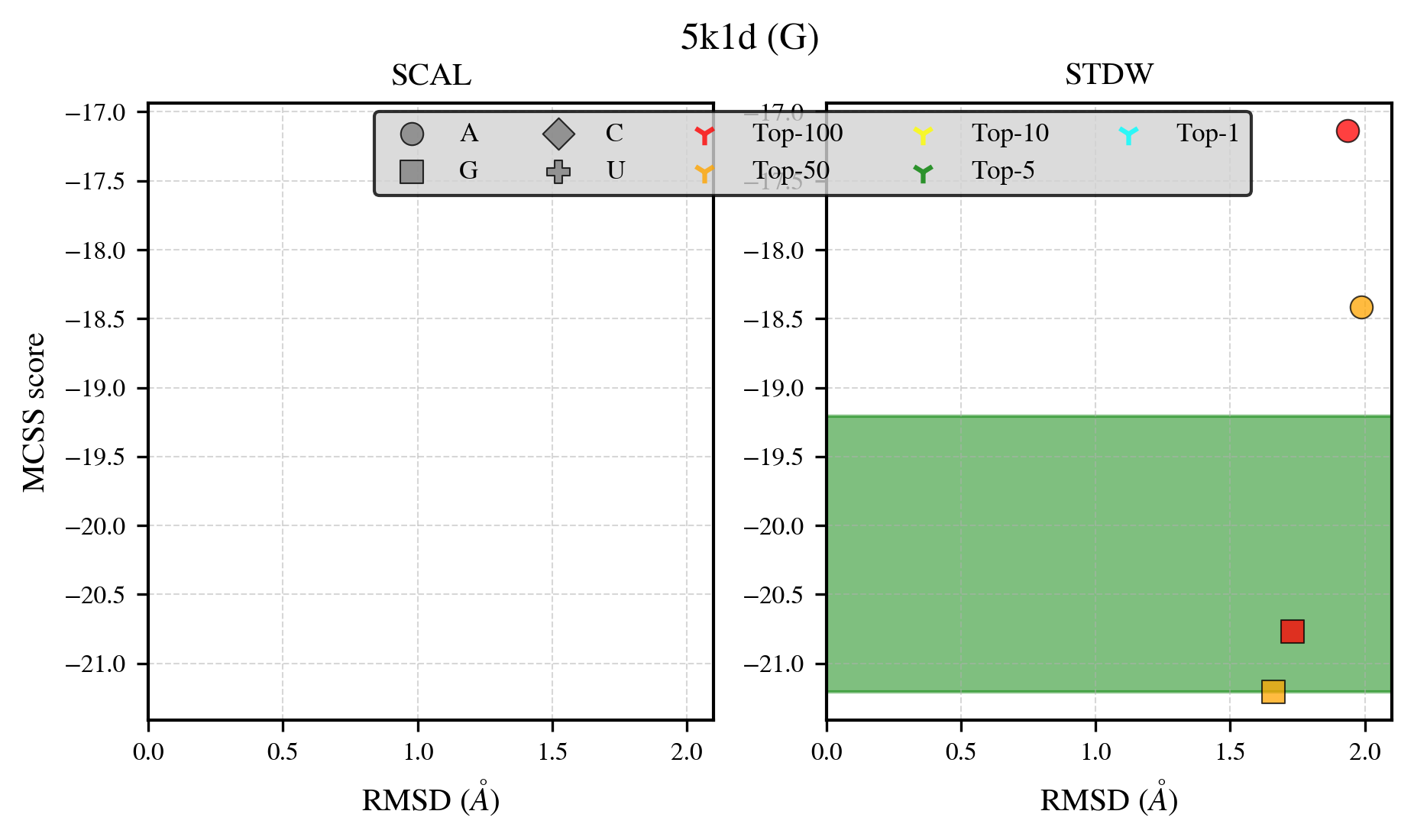

### 5v0i_benchmark-121_selectivity_details.png

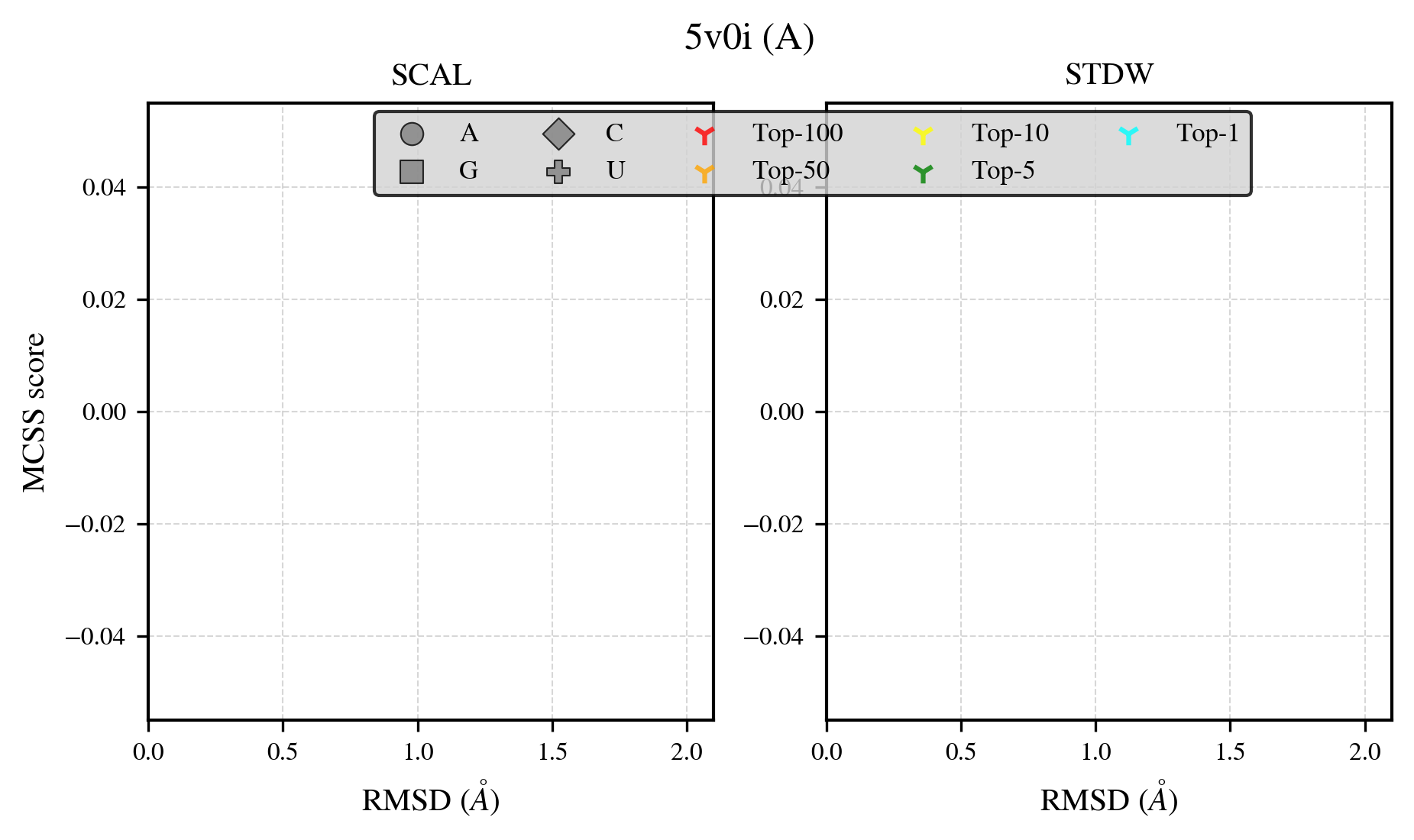

### 5v1m_benchmark-121_selectivity_details.png

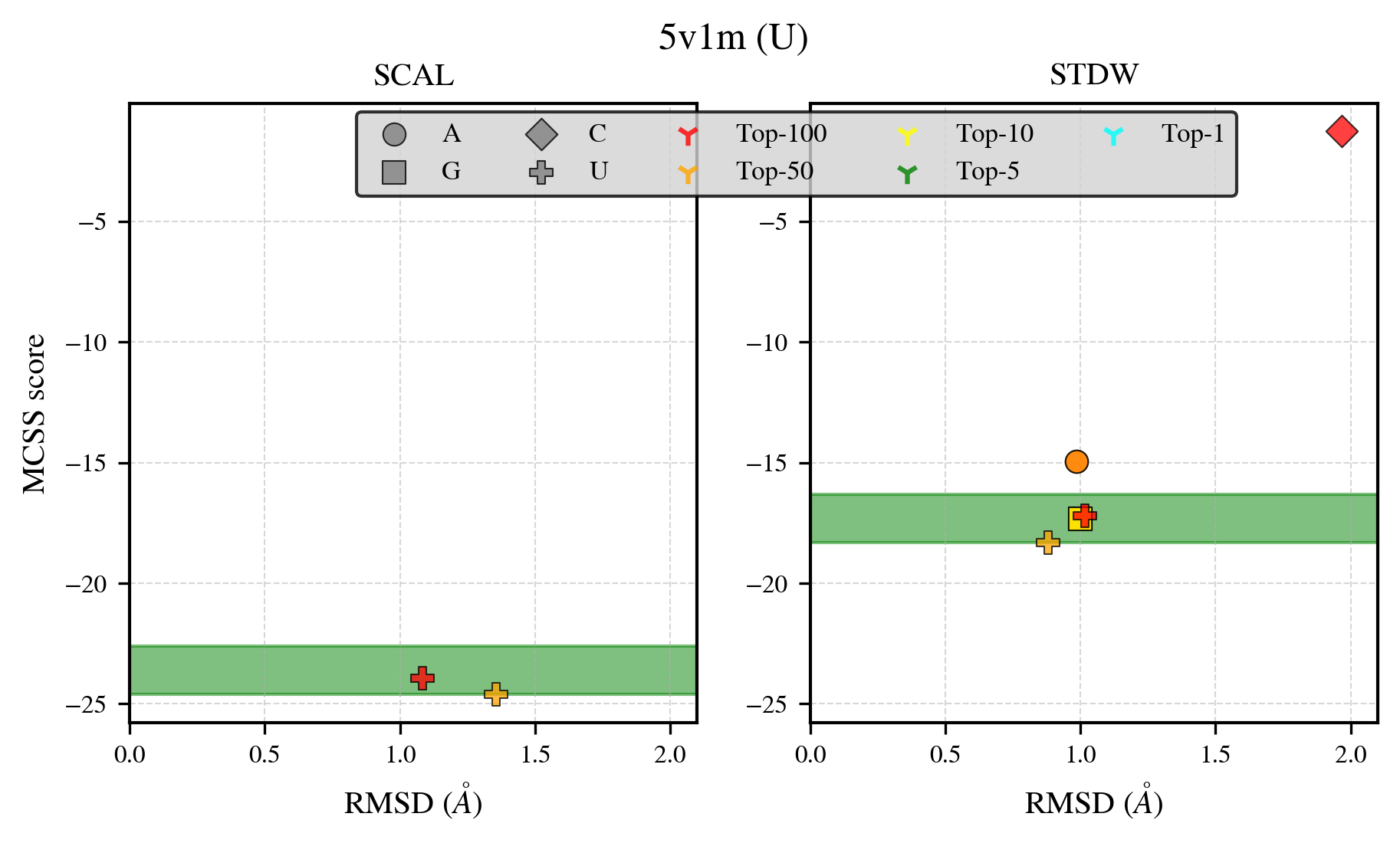

### 5x0j_benchmark-121_selectivity_details.png

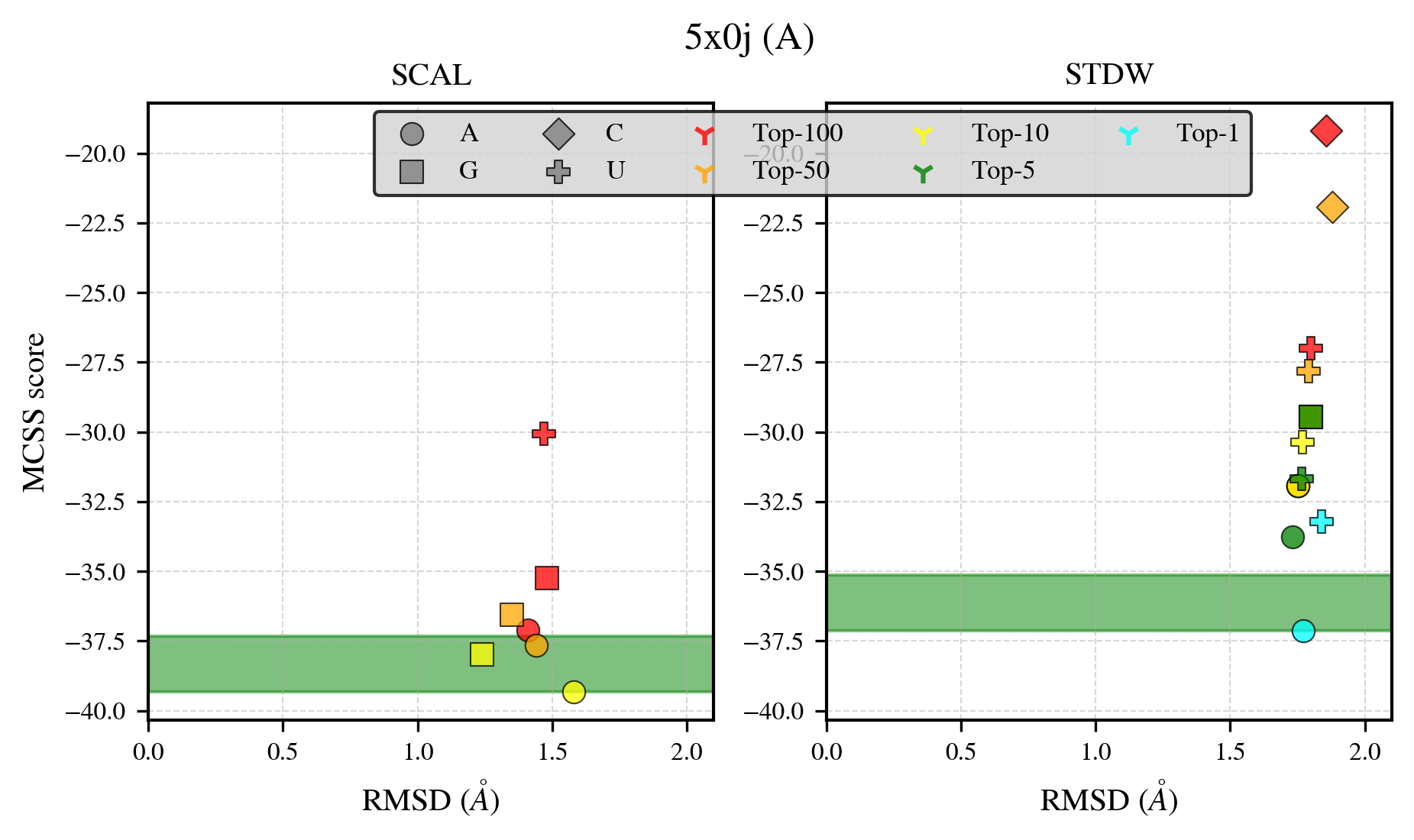
